# Human iPSC modeling reveals mutation-specific responses to gene therapy in Best disease

**DOI:** 10.1101/796581

**Authors:** Divya Sinha, Benjamin Steyer, Pawan K. Shahi, Katherine Mueller, Rasa Valiauga, Kimberly L. Edwards, Cole Bacig, Stephanie S. Steltzer, Sandhya Srinivasan, Amr Abdeen, Evan Cory, Viswesh Periyasamy, Alireza Fotuhi Siahpirani, Edwin M. Stone, Budd A Tucker, Sushmita Roy, Bikash R. Pattnaik, Krishanu Saha, David M. Gamm

## Abstract

Dominantly inherited disorders are not typically considered therapeutic candidates for gene augmentation. Here, we utilized patient-specific induced pluripotent stem cell-derived retinal pigment epithelium (iPSC-RPE) to test the potential of gene augmentation to treat Best disease, a dominant macular dystrophy caused by over 200 missense mutations in *BEST1*. Gene augmentation in iPSC-RPE fully restored BEST1 calcium-activated chloride channel activity and improved rhodopsin degradation in iPSC-RPE models of recessive bestrophinopathy and dominant Best disease caused by two different ion binding domain mutations. A dominant Best disease iPSC-RPE model that did not respond to gene augmentation showed normalization of BEST1 channel activity following CRISPR-Cas9 editing of the mutant allele. We then tested gene editing in all three dominant Best disease iPSC-RPE models, which produced premature stop codons exclusively within the mutant *BEST1* alleles. Single-cell profiling demonstrated no adverse perturbation of RPE transcriptional programs in any model, although off-target analysis detected a silent genomic alteration in one model. These results suggest that gene augmentation is a viable first-line approach for some dominant Best disease patients and that non-responders are candidates for alternate approaches such as genome editing. However, testing genome editing strategies for on-target efficiency and off-target events using patient-matched iPSC-RPE model systems is warranted. In summary, personalized iPSC-RPE models can be used to select among a growing list of gene therapy options to maximize safety and efficacy while minimizing time and cost. Similar scenarios likely exist for other genotypically diverse channelopathies, expanding the therapeutic landscape for affected patients.

**Significance:** Dominantly inherited disorders pose distinct challenges for gene therapies, particularly in the face of extreme mutational diversity. We tested whether a broad gene replacement strategy could reverse the cellular phenotype of Best disease, a dominant blinding condition that targets retinal pigment epithelium (RPE). Using RPE generated from patient-specific induced pluripotent stem cells (iPSCs), we show that gene replacement functionally overcomes some, but not all, of the tested mutations. In comparison, all dominant Best disease models tested were phenotypically corrected after mutation-specific genome editing, although one off-target genomic alteration was discovered. Our results support a two-tiered approach to gene therapy for Best disease, guided by safety and efficacy testing in iPSC-RPE models to maximize personal and public health value.

## Introduction

Genotypically heterogeneous dominant diseases pose significant challenges and opportunities for precision medicine (1). Among gene therapies, gene augmentation for recessive disorders is the most developed, having spurred multiple clinical trials (2–4) and FDA approval for one ocular disease (5). However, gene augmentation is generally ruled out as a stand-alone therapy for dominant disorders due to a perceived need to eliminate the deleterious effects of the mutant allele. Gene editing approaches to silence or repair mutant alleles hold promise in this regard (6–8), but testing safety and efficacy for every mutant allele-specific genome editor presents practical and economic challenges in diseases with high mutational diversity. Further, gene editing may not be able to target all mutations (6, 9, 10) and could lead to off-target mutagenesis—particularly within a heterozygous wildtype allele—or other adverse events (11). Another consideration for gene therapy development is the need for preclinical model systems with phenotypes and/or genotypes that are relevant to the human disease. This requirement is particularly challenging for genome editing strategies, which utilize sequence-specific tools and thus require human model systems to test safety and efficacy (12). Humanized animal models have also been employed for this purpose (13), although they cannot be used for genome-wide off-target analysis.

One disorder that faces a full array of these therapeutic obstacles is Best disease, a major cause of inherited macular degeneration that currently has no treatment options. Best disease exclusively targets the retinal pigment epithelium (RPE), a monolayer of cells essential for the survival and function of photoreceptors. Although Best disease is often diagnosed in early childhood based on its distinctive ophthalmological findings (14), its effects on central vision are generally mild at first. Vision loss occurs progressively and irreversibly over several decades, thus providing a wide time window for therapeutic intervention.

Best disease is a genotypically diverse disorder transmitted primarily in an autosomal dominant fashion, although rare cases of autosomal recessive bestrophinopathy (ARB) are known (15). Together, autosomal dominant Best disease (adBD) and ARB are linked to over 200 mutations in the *BEST1* gene, which encodes a putative homo-pentameric calcium-activated chloride channel (CaCC) found in the RPE. Recent elucidation of the high-resolution crystal structure of chicken Best1 reinforced its role as a CaCC and revealed that disease-associated mutations cluster within calcium or chloride ion binding sites or within structural regions of the channel (16).

A significant impediment to the development of therapies for adBD is the lack of model systems that adequately mimic the genotypic and phenotypic characteristics of the disorder. While canine models of ARB mirror the human ARB phenotype (14), no suitable animal models of adBD exist. To provide a therapeutic testing platform for adBD, we previously developed the first human iPSC-RPE models of the disease, which demonstrated relevant cellular dysfunction; most notably, delayed degradation of phagocytosed photoreceptor outer segment (POS) proteins (17, 18). These adBD iPSC-RPE models were then used to test the potential for selected pharmacological interventions to ameliorate the cellular phenotype of this disorder (18).

In the present study, we examined whether gene therapy could definitively correct the functional defects present in adBD iPSC-RPE. Given that BEST1 forms a homo-pentameric CaCC, we hypothesized that gene augmentation could potentially mitigate the cellular disease phenotype in adBD by increasing the ratio of wild-type to mutant BEST1 monomers available for channel assembly. This theory presumes that the deleterious effects of the mutant allele can be diluted sufficiently to restore CaCC function, preferably in a controlled manner without the risks associated with unregulated transgene expression.

To test our hypothesis, we employed three iPSC-RPE models of adBD, along with one iPSC-RPE model of ARB as a control. Importantly, the iPSC lines were generated from patients with *BEST1* mutations in different functional regions of the channel (*i.e.*, calcium binding, chloride binding, and structural) (16). We then ectopically expressed wildtype *BEST1* in iPSC-RPE using a viral vector that incorporated the native *BEST1* promoter, *VMD2*, in order to maintain RPE specificity and to keep transgene expression levels in check. Using this strategy, we obtained a >3-fold increase in wildtype BEST1 protein expression across all adBD iPSC-RPE models. Single cell electrophysiology and cell population-based assays revealed that two of the adBD mutations were exceedingly responsive to gene augmentation alone. Indeed, the correction of the cellular disease phenotype observed in these adBD iPSC-RPE models following gene augmentation was on par with that seen in the ARB iPSC-RPE model.

To address the adBD mutation that failed to respond to gene augmentation, as well as others that may also be refractory to this broad therapeutic strategy, we examined whether CRISPR-Cas9 gene editing could specifically target the mutant *BEST1* allele, leaving the normal allele intact. We found that gene editing was highly efficient at eliminating mutant allele expression and restoring iPSC-RPE CaCC activity in all three adBD models. These results bode well for the use of CRISPR-Cas9 to treat adBD mutations that are not candidates for gene augmentation, contingent on the availability of suitable guide RNAs. We then investigated whether gene editing caused untoward effects on the RPE transcriptome or induced off-target genome alterations in any of the adBD models. While no transcriptomic perturbations were detected, a single significant—albeit functionally silent—off-target site contained genomic insertions and deletion mutations (indels) in one adBD model. Based on our findings, we propose a two-tiered approach to adBD gene therapy that uses iPSC-RPE testing to first determine which mutations are likely to respond to frontline treatment with gene augmentation. *BEST1* mutant iPSC-RPE models that do not demonstrate phenotypic correction with gene augmentation would then undergo next-level safety and efficacy testing to assess candidacy for customized genome editing.

## Results

### BEST1 mutations decrease CaCC activity in patient-specific iPSC-RPE

In addition to the N926H and A146K adBD iPSC lines previously reported (17, 18), we generated iPSCs from a third adBD patient with an R218C mutation and an ARB patient with compound heterozygous mutations (R141H/A195V) (**Figure 1A**). Based on the crystallographic studies, each of these mutations lies within a different functional region of the BEST1 channel (**Figure 1B**) (16). We also employed two control iPSC lines: a wildtype (WT) iPSC line and an isogenic iPSC line generated via CRISPR-based gene correction of R218C adBD iPSCs (R218C>WT) (19). All six iPSC lines were tested for pluripotency, differentiated to RPE, and characterized (**Figures 1C-D**, and **S1A-D**). iPSC-RPE monolayers for all adBD and control lines, but not the ARB line, showed robust levels of BEST1 protein expression (**Figure 1D**). The profoundly decreased BEST1 level in our ARB cultures is consistent with reports using heterologous expression or iPSC-RPE systems that showed low or undetectable levels of R141H or A195V BEST1 (20, 21). As a measurement of CaCC activity, single-cell patch-clamp recordings of calcium-activated chloride current density were performed and found to be greatly diminished in all patient-specific iPSC-RPE relative to WT control iPSC-RPE (**Figures 1E** and S1E-I). Gene-corrected R218C>WT isogenic iPSC-RPE control showed CaCC current density at levels similar to native WT control lines (**Figures 1E and S1J**), indicating that the decreased CaCC activity was indeed the result of the BEST1 mutation.

**Figure 1.**
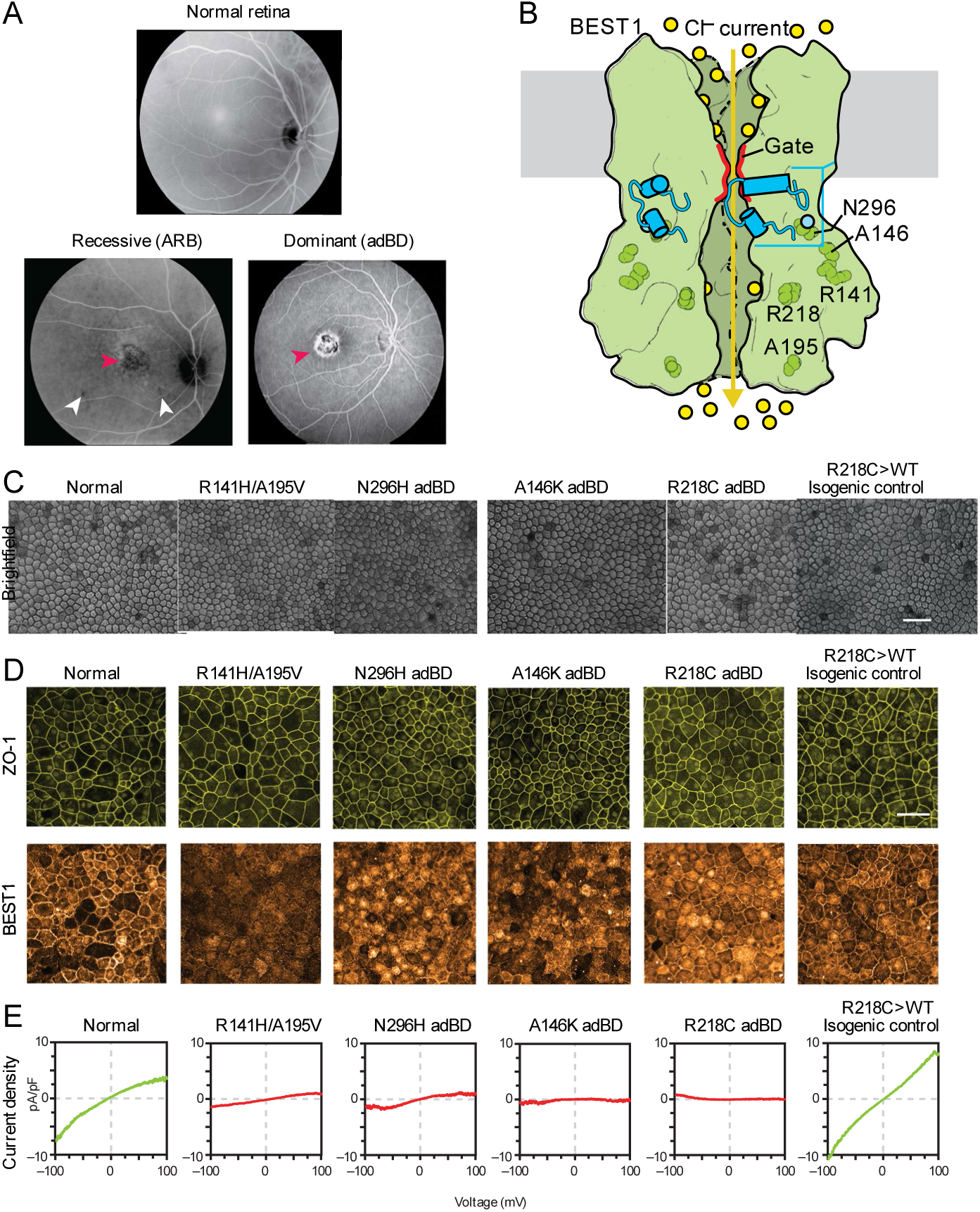
*BEST1* mutations reduce CaCC current in Best disease iPSC-RPE. **(A)** *top*, image (in grayscale) of a normal fundus; *bottom left*, fundus image of an ARB patient with R141H/A195V compound heterozygous mutations in *BEST1* showing a vitelliform lesion in the macula (*red* arrowhead) as well as small lesions outside the macula (*white* arrowheads); *bottom right*, fundus image showing a vitelliform macular lesion (*red* arrowhead) in an adBD patient with a heterozygous R218C encoding mutation in *BEST1*. **(B)** A fully functional homo-pentameric BEST1 channel is formed by assembly of WT subunits (*green*), allowing movement of chloride ions (*yellow* circles) upon binding of calcium ions (*light blue* circle) (based on the eukaryotic Best1 crystal structure (16)). **(C)** Light microscopic images of normal, patient-specific, and isogenic control iPSC-RPE used in this study. Scale bar = 50 µm (applies to all images in C). **(D)** Immunocytochemical analyses of ZO-1 and BEST1 protein expression in iPSC-RPE cells. Scale bar = 50 µm (applies to all images in D). **(E)** CaCC current density-voltage plots from WT, R141H/A195V ARB, or adBD iPSC-RPE cells, as determined by calculating the difference in average chloride currents in the presence or absence of calcium (Figure S1). For +calcium: n = 6 cells for WT, 12 cells for R141H/A195V ARB, 7 cells for N296H adBD, 5 cells for A146K adBD, 5 cells for R218C adBD, and 10 cells for R218C>WT isogenic control; for no calcium: n = 8 cells for WT, 12 cells for R141H/A195V ARB, 8 cells for N296H adBD, 7 cells for A146K adBD, 8 cells for R218C adBD, and 9 cells for R218C>WT isogenic control (data combined from at least two replicates).

### *BEST1* augmentation restores CaCC activity and enhances rhodopsin degradation in ARB iPSC-RPE

We next sought to confirm that ectopic expression of WT human BEST1 (hBEST1) could ameliorate the disease phenotype of R141H/A195V ARB iPSC-RPE, analogous to gene augmentation studies using ARB canines or other iPSC-RPE model systems for ARB (22, 23). Single-cell patch clamp recordings of calcium-activated chloride current density were used as a readout of efficacy in iPSC-RPE cells. In addition, we monitored degradation of rhodopsin following POS feeding as an assay of intact RPE monolayer function.

For gene augmentation we used a lentivirus construct (*hVMD2*-*hBEST1-T2A*-*GFP*) designed to co-express hBEST1 and green fluorescent protein (GFP) under control of the human *BEST1* promoter (*hVMD2*), which assures both RPE-specific expression and *BEST1*-specific gene regulation (**Figures 2A, B**). Lentivirus was chosen for transgene delivery based on its safe use in human retinal gene therapy trials (24) (ClinicalTrials.gov Identifiers: NCT01367444, NCT01736592) and its superior transduction efficiency in cultured human RPE (17, 25). GFP expression was observed in ARB iPSC-RPE cells post-transduction, and immunocytochemical (ICC) and western blot analysis confirmed enhanced expression of BEST1 in treated cultures (**Figures 2C, S2A-C**). By ≥4 weeks post-transduction, CaCC current density in ARB iPSC-RPE increased significantly, reaching levels comparable to WT iPSC-RPE (**Figures 2D, E and S2E**). Furthermore, transduced monolayers of ARB iPSC-RPE demonstrated enhanced degradation of rhodopsin following POS feeding (**Figure 2F and S2I**). These findings, together with those reported by Guziewicz et al. (22) and Li et al. (23), support *hBEST1* gene augmentation as a treatment for ARB.

**Figure 2.**
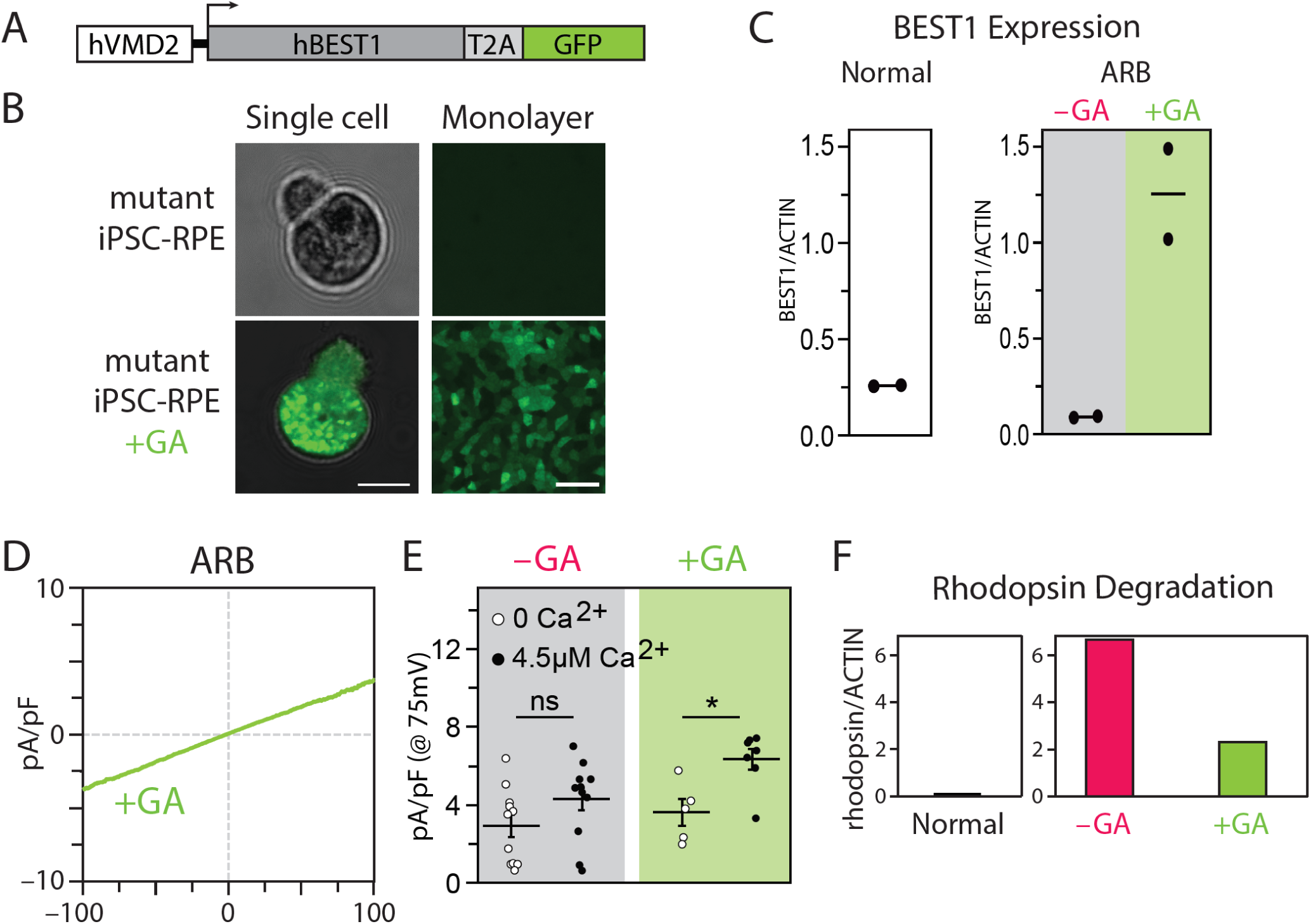
Gene augmentation rescues the ARB iPSC-RPE cell phenotype. **(A)** Construct used for *BEST1* gene augmentation (GA). **(B)** Presence or absence of GFP fluorescence in a single dissociated iPSC-RPE cell (*left*) or iPSC-RPE monolayers (*right*) before (*top*) or after (*bottom*) gene augmentation. Scale bar = 10 µm (*left*); 50 µm (*right*). **(C)** Western blot-based quantification of BEST1 protein levels (normalized to ACTIN) in WT iPSC-RPE, ARB iPSC-RPE, and ARB iPSC-RPE after *BEST1* augmentation. **(D)** CaCC current density-voltage plots after gene augmentation in ARB iPSC-RPE. n = 7 cells for +calcium and 5 cells for no calcium (data combined from two replicates (Figure S2)). **(E)** CaCC conductance for individual ARB iPSC-RPE cells at 75 mV before or after gene augmentation. The number of cells is the same as for panels 1E and 2D. Error bars represent mean ± SEM; ns = p ≥0.05, * for p <0.05. **(F)** Western blot-based quantification of rhodopsin levels 120 hr after photoreceptor outer segment (POS) feeding in WT iPSC-RPE or in ARB iPSC-RPE with or without WT *BEST1* gene augmentation.

### *BEST1* augmentation restores CaCC activity and enhances rhodopsin degradation in R218C and N296H adBD iPSC-RPE, but not in A146K adBD iPSC-RPE

Although not as intuitive, we suspected that gene augmentation might also be a viable solo therapeutic strategy for adBD-causing *BEST1* mutations. More specifically, we hypothesized that CaCC activity could be restored by increasing the intracellular ratio of wildtype to mutant BEST1 monomers available to form the homo-pentameric channel.

The same *hVMD2*-*hBEST1-T2A-GFP* lentiviral construct that was tested in ARB iPSC-RPE was used to transduce iPSC-RPE from all three adBD patients (**Figure S2D**). Following gene augmentation, BEST1 levels in each adBD iPSC-RPE model were comparable to those achieved in gene augmented ARB-iPSC-RPE and >3-fold higher than BEST1 levels present in parallel cultures of untreated adBD iPSC-RPE (**Figure 3A and S2C**). At ≥4 weeks post-transduction, CaCC activity was fully restored in the R218C and N296H adBD iPSC-RPE models, whereas the A146K adBD iPSC-RPE model remained unresponsive (**Figure 3B-D and S2F-H**) despite displaying the highest fold increase in BEST1 expression (**Figure 3A**). Consistent with these single-cell electrophysiological findings, gene augmentation improved rhodopsin degradation in R218C and N296H iPSC-RPE, but not in A146K iPSC-RPE (**Figure 3E and S2J-L**).

**Figure 3.**
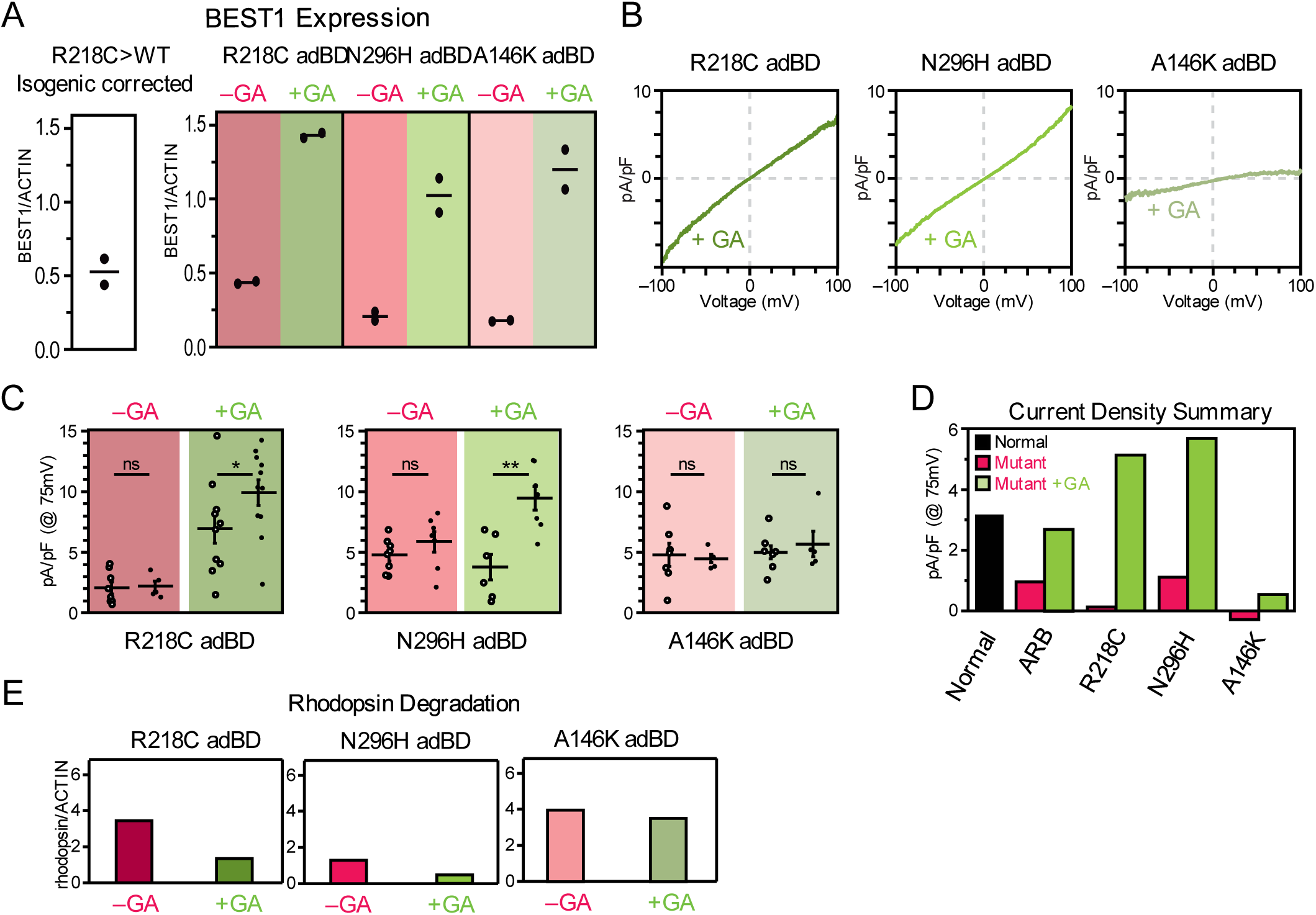
Gene augmentation rescues the cell phenotype in some, but not all, adBD iPSC-RPE models. **(A)** Western blot-based quantification of BEST1 protein levels (normalized to ACTIN) in WT iPSC-RPE and in the adBD iPSC-RPE models before and after *BEST1* gene augmentation (GA). **(B)** CaCC current density-voltage plots after gene augmentation in adBD iPSC-RPE. For +calcium: n = 11 cells for R218C, 7 cells for N296H, and 5 cells for A146K; for no calcium: n = 9 cells for R218C, 6 cells for N296H, and 8 cells for A146K (data combined from two replicates). **(C)** CaCC conductance for individual adBD iPSC-RPE cells at 75 mV before and after gene augmentation. The number of cells is the same as for panels 1E and 3B. Error bars represent mean ± SEM; ns = p ≥0.05, * for p <0.05, ** for p <0.01. **(D)** Mean CaCC conductance at 75 mV before or after gene augmentation for all iPSC-RPE tested. **(E)** Rhodopsin levels 48 hours after feeding POS to adBD iPSC-RPE with or without WT *BEST1* gene augmentation.

### Gene editing specifically targets the mutant allele in A146K adBD iPSC-RPE and restores CaCC activity

To determine whether A146K iPSC-RPE would respond to an alternative therapeutic approach, we tested gene editing as a means to eliminate expression of the mutant *BEST1* allele. Gene editing with CRISPR-Cas9 creates targeted double strand breaks in genomic DNA that are primarily repaired by endogenous non-homologous end joining (NHEJ) (26), leading to indels. These indels can cause transcriptional frameshifts that lead to premature termination codons, activation of intrinsic nonsense-mediated decay (NMD) pathways, and degradation of transcription products (27, 28).

An sgRNA sequence targeting specifically the A146K locus in the mutant *BEST1* allele was cloned into a lentiviral plasmid that encoded both the sgRNA (expressed via a U6 promoter) and a human codon optimized *Streptococcus pyogenes Cas9* (*spCas9)-T2A-GFP* transcript (expressed via a *hVMD2* promoter) (**Figures 4A and 4B**). We also cloned a sgRNA sequence targeting the *AAVS1* safe harbor locus (29) into the same lentiviral plasmid backbone to serve as an experimental control.

**Figure 4.**
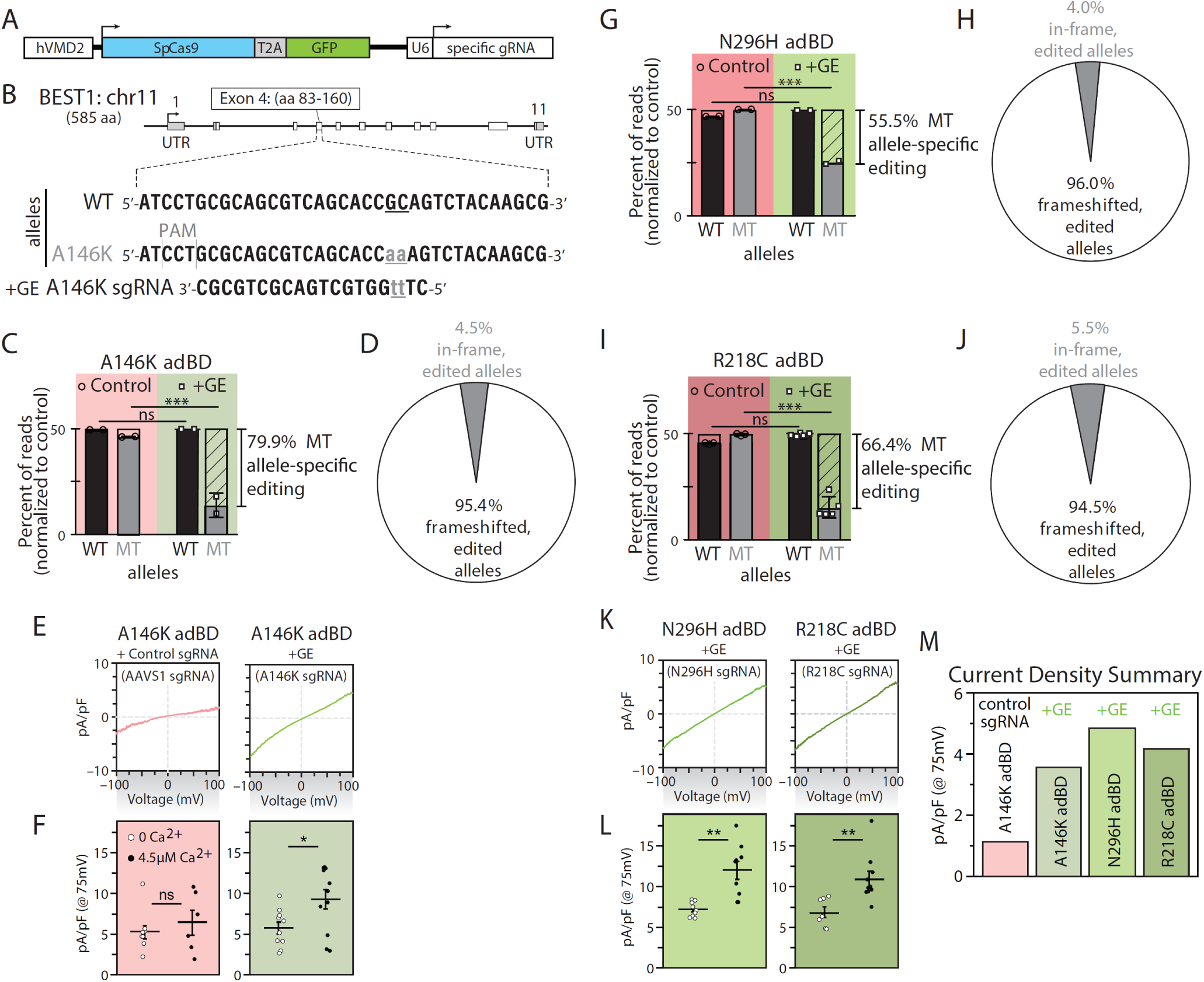
Gene editing specifically and efficiently introduces frameshifts within the mutant allele in adBD iPSC-RPE and rescues CaCC activity. **(A)** Lentiviral genome editing construct expressing *sp*Cas9 and mutant allele-targeted sgRNAs. **(B)** Diagram showing the heterozygous base pair substitutions in A146K adBD and the design of the A146K sgRNA. The wildtype (WT) allele is shown above, while the A146K adBD allele is shown below, with the mutated bases indicated in lower case and underlined. **(C)** Percentage of WT and mutant (MT; unedited and edited) allele sequencing reads in A146K iPSC-RPE treated with A146K sgRNA lentiviral genome editor (“+GE”), respectively, normalized to control (“Control”, genome edited with safe harbor *AAVS1*-targeting sgRNA). **(D)** Indel frameshift and in-frame frequency for mutant allele-edited reads from A146K adBD iPSC-RPE (corresponds to 4C). **(E)** CaCC current density-voltage plots and **(F)** CaCC conductance for individual iPSC-RPE cells from single-cell patch clamp experiments for A146K iPSC-RPE treated with control (*AAVS1*) or mutant allele-targeted sgRNA lentiviral genome editor. **(G-J)** Percentage of WT and mutant (MT; unedited and edited) allele sequencing reads in N296H (G) or R218C (I) adBD iPSC-RPE treated with N296H or R218C sgRNA lentiviral genome editor, respectively, normalized to control (*AAVS1* sgRNA). Indel frameshift and in-frame frequency in N296H (H) or R218C (J) adBD iPSC-RPE treated with N296H or R218C sgRNA lentiviral genome editor, respectively (correspond to 4G and 4I, respectively). **(K)** CaCC current density-voltage plots and **(L)** CaCC conductance for individual iPSC-RPE cells from single-cell patch clamp experiments for N296H or R218C adBD iPSC-RPE treated with respective mutant allele-targeted sgRNA lentiviral genome editor. **(M)** Mean CaCC conductance at 75 mV for each adBD iPSC-RPE model. The number of cells is the same as 4E and 4K. For gene editing experiments (4C,D and G-J), n = 2 (A146K IPSC-RPE and N296H RPE) and n = 5 (R218C iPSC-RPE). For electrophysiology experiments (4E,F and K-M), +calcium: n = 6 cells for *AAVS1*, 11 cells for A146K, 9 cells for N296H, 10 cells for R218C; no calcium: n = 9 cells for *AAVS1*, 10 cells for A146K, 9 cells for N296H, 7 cells for R218C (data combined from two replicates). Error bars in 4C,G,I represent mean ± SD; ns = p ≥0.05, *** for p <0.001. Error bars in 4F and 4L represent mean ± SEM; ns = p ≥0.05, * for p <0.05, ** for p <0.01.

Two weeks after transduction of A146K adBD iPSC-RPE with A146K sgRNA or control (*AAVS1* sgRNA) lentiviral genome editor, we quantified the average frequency of deep sequencing reads corresponding to WT, mutant, and edited alleles in genomic DNA. We detected a nearly 80% editing frequency of the A146K mutant allele with no decrease in WT allele frequency post-editing (**Figure 4C**). Together, these results reflect efficient editing with high specificity for the A146K mutant allele over the WT *BEST1* allele.

Using deep sequencing, we next examined specific indels that were introduced into A146K iPSC-RPE two weeks post-transduction with the A146K sgRNA genome editor (**SI data file A**). An average of 95.4% of the edited alleles resulted in a frameshift mutation (**Figure 4D and SI data file A**), which is higher than the percentage of out-of-frame indels predicted by a recent machine learning algorithm (**SI data file A**) (30). This finding indicates a high likelihood that indels resulting from gene editing at the A146K locus in the mutant *BEST1* allele will trigger NMD of the transcribed RNA, effectively knocking out expression of the mutant allele in the vast majority of edited RPE cells.

We next assessed functional rescue of BEST1 channel activity in AAVS1 control versus A146K mutant allele gene-edited iPSC-RPE. Single-cell patch-clamp experiments revealed restoration of CaCC activity in gene-edited A146K iPSC-RPE, but not in control *AAVS1* sgRNA treated A146K iPSC-RPE (**Figure 4E, F and S3**).

### Mutant allele-specific gene editing restores CaCC activity in all tested adBD iPSC-RPE

While the gene editing results obtained in the A146K adBD iPSC-RPE model were highly encouraging, it is possible that this locus is unique in its potential to be targeted by a mutant allele-specific sgRNA. To extend this investigation, we also evaluated the specificity and efficacy of gene mutant allele editing in the N296H and R218C adBD iPSC-RPE models. N296H and R218C mutant allele-targeted sgRNAs were designed and cloned into separate lentiviral plasmids as described for the A146K sgRNA. N296H iPSC-RPE and R218C iPSC-RPE were transduced with lentiviral genome editors encoding either control (*AAVS1*) or corresponding allele-targeted sgRNA and editing outcomes were measured via deep sequencing of genomic DNA (**SI data file A**). Quantification of WT and mutant allele frequency revealed efficient targeting of the N296H and R218C mutant alleles with their respective sgRNAs (55.5% and 66.4%, respectively) with no demonstrable targeting of the WT alleles (**Figure 4G, I**). A high proportion of editing in these two models resulted in out-of-frame indels (96.0% and 94.5% for N296H and R218C iPSC-RPE, respectively) (**Figure 4H, J**). Subsequent single-cell patch-clamp measurements of CaCC current density confirmed restoration of channel activity post-gene editing in both R218C and N296H iPSC-RPE (**Figure 4K-M and S3**). Thus, while some variation in gene editing efficiency was observed using the three different sgRNAs (as expected), more than half of the mutant alleles were edited (with a high percentage of out-of-frame indels) in the three adBD iPSC-RPE models, with no editing of the WT allele.

### Mutant allele-specific gene editing does not perturb global iPSC-RPE transcriptional programs, although off-target editing can occur

Although the mutant allele-specific sgRNAs tested in the three adBD iPSC-RPE models did not target the fellow WT alleles in any of our experiments, the potential for off-target adverse effects elsewhere within the genome still exists. To detect untoward transcriptional effects from gene editing, we performed single-cell RNA sequencing (scRNA-seq) for 12,061 individual iPSC-RPE cells treated with genome editors. iPSC-RPE (R218C, N296H, A146K, or isogenic control R218C>WT) were edited with genome editors encoding either a mutant allele-targeted sgRNA or a control sgRNA targeting the *AAVS1* site, to generate a total of eight separate samples (**Figure S4A**).

Evaluation of t-Distributed Stochastic Neighbor Embedding (t-SNE) clustering of cells across all eight samples indicated that, by virtue of using the *hVMD2* promoter, *spCas9-T2A-GFP* transcript levels closely corresponded with *BEST1* transcript levels (**Figure 5A**). Visual comparison of t-SNE clustering of each individual sample demonstrated that transcriptional signatures are grossly similar between iPSC-RPE lines, whether treated with mutant allele-targeted (+GE) or control (*AAVS1*) sgRNA (**Figure 5B *top***). This observation was supported quantitatively by non-negative matrix factorization (NMF). NMF analysis demonstrated that greater transcriptome variation exists between iPSC-RPE from different lines than between iPSC-RPE from the same line treated with mutant allele-targeted or control sgRNA (**Figure S4B**).

**Figure 5.**
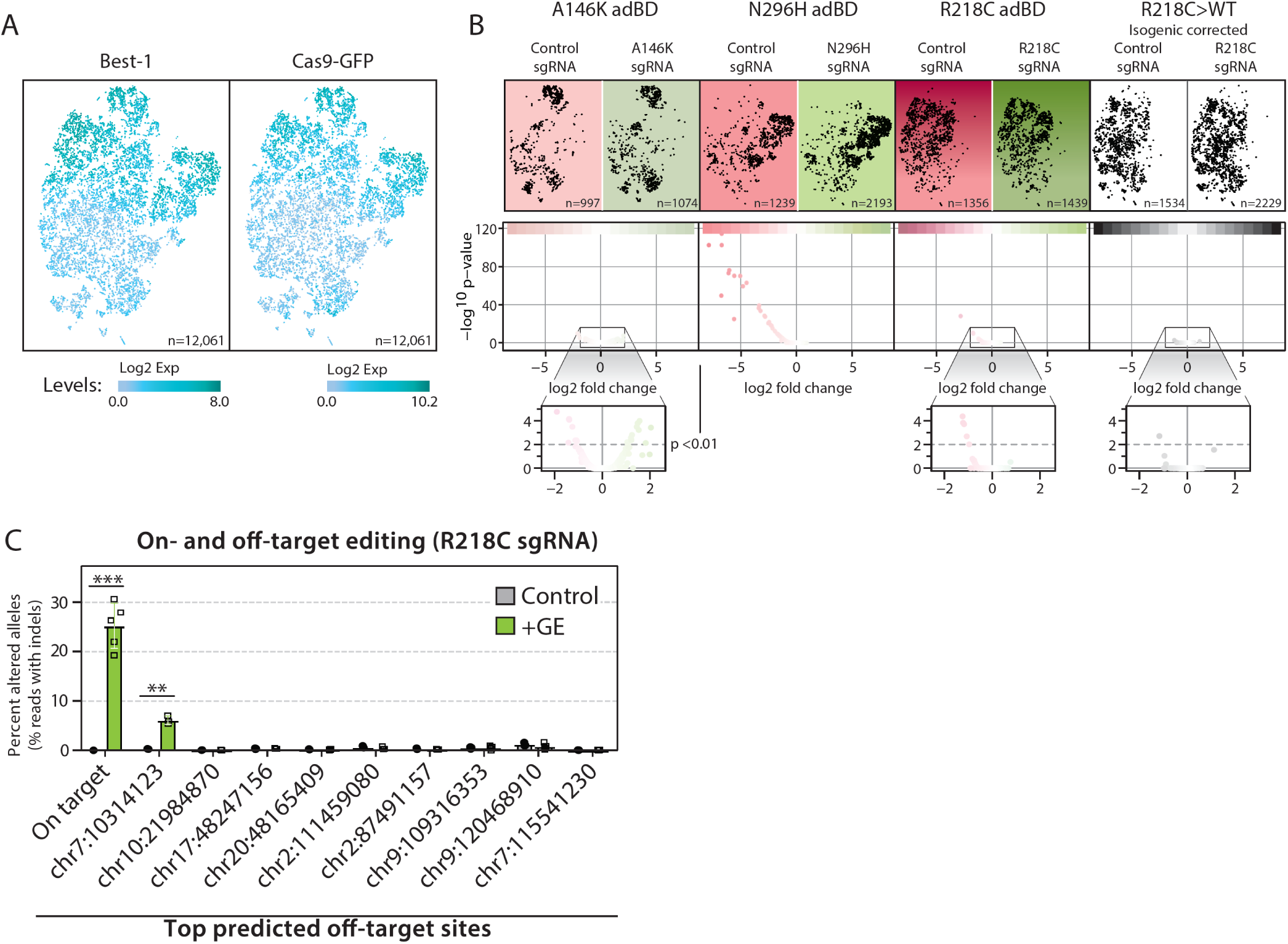
Gene editing did not disrupt iPSC-RPE transcriptional programs. **(A)** t-SNE plot of single iPSC-RPE cells across all 8 samples with relative expression of *BEST1* (*left*) and *spCas9-T2A-GFP* (*right*) depicted via increasing shades of *blue*. Total number of cells analyzed (n) is shown. **(B)** *Top*, t-SNE plot of single cells (*black* dots) from each treated sample. Number of cells analyzed (n) for each sample is shown. *Bottom*, Volcano plots of transcriptome-wide differences in expression of individual genes (*red* or *green* dots) between iPSC-RPE of the same genotype treated with mutant allele-targeted sgRNA (*green*) versus control (*AAVS1*, *red*) sgRNA lentiviral genome editor. p <0.01 was the threshold for determining significant versus non-significant changes in gene expression. **(C)** Frequency of edited alleles at on-target and top nine ranked off-target loci in R218C adBD iPSC-RPE treated with R218C sgRNA lentiviral genome editor (n=3 for control and n=5 for +GE, except n=3 at first *chr 7* off-target locus). Off-target sites are annotated by the location of the first base of the predicted off-target site (further detailed in SI Data File C). Error bars represent mean ± SD; ** for p <0.01, *** for p <0.001.

Additional analysis of global gene expression (**Figure 5B *bottom***) and of a focused set of genes related to negative or off-target effects (including cell cycle regulation, apoptosis, DNA damage response, or innate immune response; **Figure S4C, SI data file B**) did not reveal significant upregulation of those gene sets in mutant allele-targeted (+GE) versus control sgRNA-treated samples. However, examination of the top nine potential off-target sites for the R218C sgRNA revealed a low, yet significant percentage of editing at a single site within a non-coding region of chromosome 7 (**Figure 5C**). While this finding is not predicted to have a deleterious effect on RPE cell function, it emphasizes the importance of performing comprehensive on- and off-target genome editing analyses using a patient-specific model system.

## Discussion

The observation that a subset of adBD mutations may be amenable to gene augmentation greatly expands the Best disease patient population that might benefit from this therapeutic approach. Based on the crystallographic studies by Dickson et al. (16), the two mutations that responded to gene augmentation lie within calcium clasp (N296H) or chloride binding (R218C) sites within the BEST1 channel, whereas the mutation that failed to respond (A146K) localizes to a putative structural region. Among the over 200 known *BEST1* mutations, many are predicted to be directly or indirectly involved in ion binding (16, 31). Importantly, a recent study by Ji et al. using baculovirus supports our finding that chloride and calcium binding site mutations in BEST1 can be receptive to gene augmentation (32). However, the fact that not all adBD iPSC-RPE models respond to gene augmentation underscores the need to vet patient candidacy for gene augmentation carefully.

The mechanism underlying selective responsivity of adBD patients to gene augmentation cannot be due to traditional allelic haploinsufficiency, in which half the normal amount of WT protein and no mutant protein is produced, resulting in fewer (but fully WT) BEST1 channels. Such a situation exists in parents of ARB patients, who have no demonstrable disease phenotype. Rather, adBD mutant monomers must be incorporated alongside WT monomers in all (or nearly all) BEST1 channels (33). We propose that in the case of N296H and R218C, this commingling of WT and mutant monomers causes ion binding site insufficiency and channel impermeability, a condition that is surmountable by WT *BEST1* augmentation. In contrast, we hypothesize that *BEST1* mutations like A146K—which converts a nonpolar amino acid to a polar amino acid in a compact structural region of the protein—has more pervasive functional consequences, resulting in greater resistance to gene augmentation.

We did consider the possibility that mutation-specific resistance to gene augmentation was due to variability in transgene expression (*i.e.*, there was insufficient WT transgene expression in the non-responsive A146K adBD iPSC-RPE model). However, we found that BEST1 protein levels were similar in all models following gene augmentation. In fact, a slightly higher fold-increase in BEST1 levels was achieved in A146K adBD iPSC-RPE compared with the two adBD models that were rescued by gene augmentation (N296H and R218C). Thus, it is highly unlikely that the differences in functional response observed between R218C or N296H adBD iPSC-RPE and A146K adBD iPSC-RPE are due to variability in transgene expression. Resistance of the A146K mutation to functional recovery after gene augmentation also cannot be explained by occult artifacts inherent to the iPSC line or its RPE progeny, since gene editing was ultimately successful in restoring CaCC activity in the same differentiated A146K adBD iPSC-RPE population.

It is also notable that our lentiviral constructs employed the *hVMD2* promoter, which is ideal from a translational standpoint as it specifies expression in RPE and supports native regulation of *BEST1*. Use of alternative promoters poses risks of off-target cell effects and/or undesirably low (ineffectual) or high (toxic) levels of protein expression. For construct delivery, we selected lentivirus based on its excellent *in vitro* RPE transduction efficiency (17, 25) and its current use in RPE gene therapy trials (ClinicalTrials.gov Identifiers: NCT01367444, NCT01736592) (24). However, our findings are likely applicable across all *in vivo* transgene delivery platforms that possess comparable safety and transduction efficiency profiles. Indeed, Ji et al. observed improvement in CaCC activity in isolated R218H adBD iPSC-RPE cells following constitutive overexpression of WT BEST1 using an AAV delivery vector (32).

There is precedence for using patient-specific iPSCs as preclinical efficacy models for gene therapy clinical trials (34). Our work extends this utility by providing a framework for preclinical testing of mutation-specific responses in a genotypically heterogenous disease using the affected cell type. It remains to be determined whether separate adBD iPSC-RPE models will be required to assess suitability of gene augmentation versus gene editing for every mutation, or if a few models can sufficiently represent larger categories of mutations (*e.g.*, ion binding sites or structural regions) (35).

For adBD mutations like A146K that are not amenable to gene augmentation, we showed that targeted gene editing holds great promise as an alternative therapy. Indeed, there is a wide spectrum of *BEST1* mutations that could be treated by CRISPR-Cas9 by designing unique mutation-targeted sgRNAs (examples shown in **SI data file D**). While this approach would be costly and time-consuming if separate testing is required for each mutation-specific sgRNA, rapid advances in gene editing technologies and strategies may overcome such limitations. Other gene therapy strategies also exist for dominant ocular diseases; for example, knockdown of both wildtype and mutant allele transcripts with simultaneous introduction of a modified wildtype gene (36). Whether such an approach would be safe and effective for adBD mutations that fail to respond to straightforward gene augmentation is not known, but could be tested using the iPSC-RPE model systems employed here.

In our gene editing experiments, we observed higher efficiency out-of-frame editing in iPSC-RPE when compared to a prior study using undifferentiated iPSCs (19). This finding is consistent with recent reports of variable mutation bias across different cell types (30), and points to the importance of evaluating gene editing using the specific cell type(s) targeted by disease. In addition, editing at *BEST1* in iPSC-RPE did not provoke an increase in expression of genes associated with cell cycle regulation, apoptosis, DNA damage response, or innate immune response in comparison to editing at a well characterized safe-harbor locus (29) with a previously described sgRNA (37). Undesirable effects such as these have been reported in other cell types following Cas9-mediated gene editing (11, 38). Despite our reassuring findings, there remains the potential for off-target genomic alterations, as was observed at a single locus in a small percentage of iPSC-RPE cells in the R218C adBD model. While these particular off-target indels are in a non-coding region and are thus predicted to be functionally silent, their presence emphasizes the value of employing human model systems for preclinical genome editing safety studies. Interestingly, no off-target indels were detected in our prior study using the same sgRNA in undifferentiated R218C iPSCs (19), which further indicates the need to perform off-target analyses in iPSC-RPE and not in surrogate cell types.

Overall, our results provide a blueprint to guide gene therapy choice in the era of gene augmentation and gene editing (**Figure 6**). With its inherently larger target populations and established track record in patients, it is practical to utilize gene augmentation when possible, reserving gene editing for mutations that require allele repair or knockout or are otherwise untreatable by gene augmentation. It is noteworthy that the two adBD lines that demonstrated restoration of CaCC activity with gene augmentation or gene editing did so with equal efficacy, underscoring the suitability of either approach. Other desirable characteristics of Best disease as a clinical candidate for gene therapy include 1) a wide time window for gene therapy intervention, 2) accessibility of RPE using standard surgical techniques, 3) a small (∼5.5 mm diameter) treatment area, 4) availability of noninvasive retinal imaging and functional assessment tools, and 5) growing patient safety data from other RPE-based gene therapy trials (2–4). As such, Best disease is well-positioned to become the first genotypically heterogeneous disorder with dominant and recessive inheritance patterns to have a full menu of therapeutics for all affected individuals. Furthermore, implications of this work likely extend beyond the eye and Best disease to other intractable monogenic conditions caused by mutations in multimeric ion channels, including congenital myasthenic syndromes and some forms of epilepsy (39–41).

**Figure 6.**
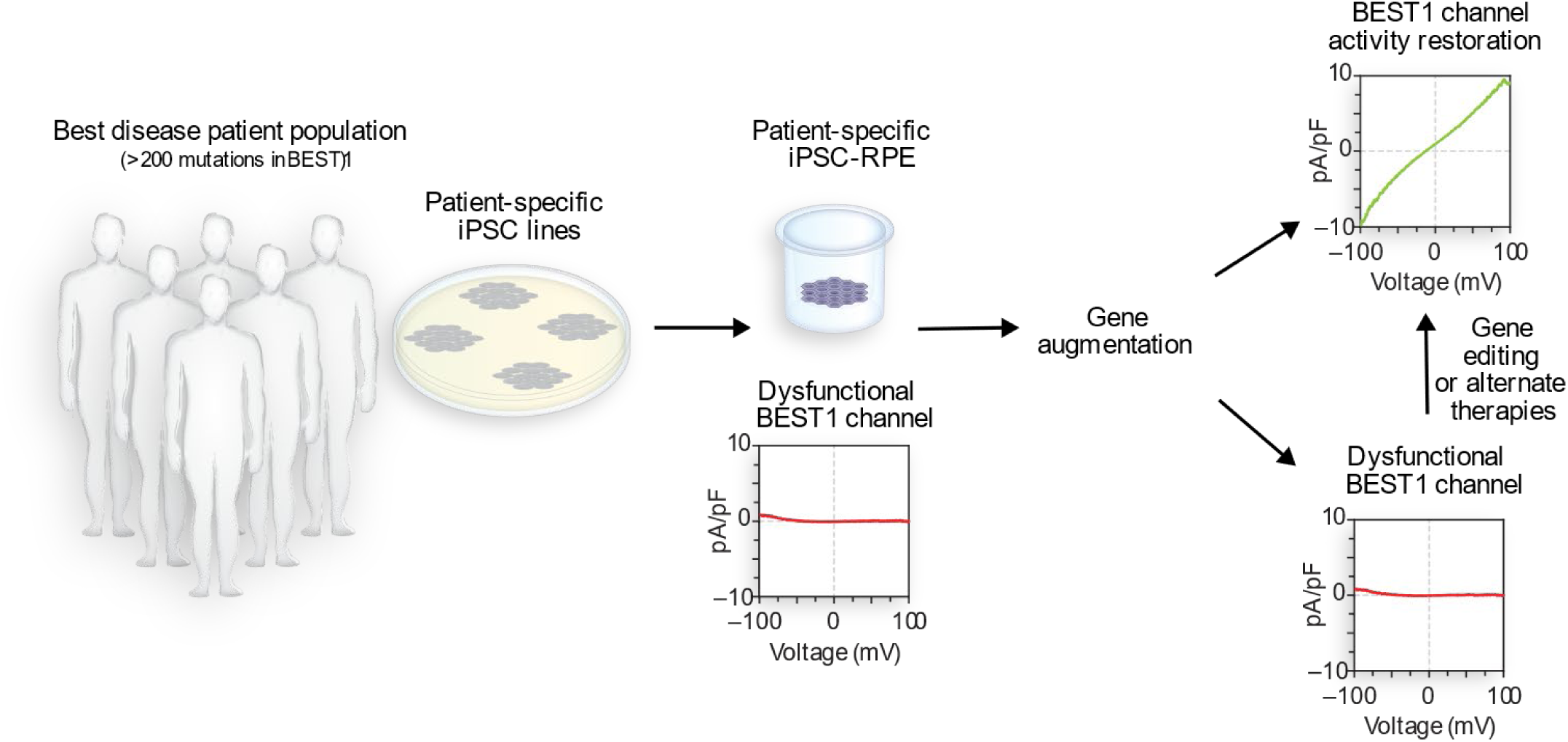
In vitro gene therapy testing strategy for adBD. The amenability of adBD mutations to correction via gene augmentation can be evaluated for efficacy and safety in a dish using patient iPSC-RPE models. Those patients with mutations that fail to respond to gene augmentation would then undergo further testing for genome editing (or another alternative strategy) using the same adBD iPSC-RPE model systems.

## Acknowledgements

The authors thank Alfred Lewin (U. Florida) for the *hVMD2*-*hBEST1*-*T2A-GFP* plasmid construct; the Cellular Imaging and Analysis core at the University of Wisconsin-Madison Waisman Center; and Andrew Thliveris for helpful discussions. This work was supported by NIH R01EY024588 (to D.M.G., B.A.T., and E.M.S.); Foundation Fighting Blindness, Research to Prevent Blindness, Retina Research Foundation Emmett Humble Chair, and McPherson Eye Research Institute Sandra Lemke Trout Chair in Eye Research (to D.M.G.); NSF CBET-1350178 and CBET-1645123, NIH R35GM119644, Brain & Behavior Research Foundation, Burroughs Wellcome Fund, and Retina Research Foundation Kathryn and Latimer Murfee Chair (to K.S.); NIH R01EY024995 and Retina Research Foundation M.D. Mathews Professorship (to B.R.P.); NIH T32HG002760 and F30EY027699, VitreoRetinal Surgery Foundation (to B.S.); DGE-1747503 (to K.M.). This study was supported in part by a core grant to the Waisman Center (NICHHD U54 HD090256). We thank members of the Gamm and Saha labs for comments on the manuscript, plasmid depositors to Addgene, the University of Wisconsin-Madison Biotechnology Center for DNA sequencing, and the University of Wisconsin-Madison Skin Disease Research Center for assistance with virus preparation.

## Author contributions

D.S. and D.M.G. designed the gene augmentation experiments. B.S., D.S., D.M.G. and K.S. designed the gene editing experiments. P.K.S. and B.R.P. performed and analyzed the electrophysiology experiments. D.S. and B.S. performed all other experiments with contributions from R.V., K.L.E., C.B, S.S.S., A.A., and E.C. K.M., S.S., V.P., A.F.S., and S.R. were primarily responsible for the scRNA-seq analysis. B.A.T. and E.M.S. provided and characterized the ARB iPSC line, D.S., B.S., K.S., and D.M.G. wrote the manuscript and analyzed data with input from all authors. D.M.G., K.S., and B.R.P. supervised research.

## Materials and Methods

### iPSC lines

A total of 6 iPSC lines, 2 control and 4 patient-specific, were used in this study. In addition to one control iPSC line (normal) and two adBD patient-specific iPSC lines (A146K adBD and N296H adBD) previously used by our group for Best disease modeling (17), we used three new iPSC lines. Two of the new iPSC lines harbored patient specific mutations: R218C for adBD and R141H/A195V for ARB. One isogenic control iPSC line was obtained by CRISPR/Cas9-based gene correction of the patient-specific R218C adBD iPSC line (19). All iPSC lines were cultured either on mouse embryonic fibroblasts (MEFs) or on Matrigel. Lines cultured on MEFs were maintained using iPS media (Dulbecco’s Modified Eagle’s Medium (DMEM)/F12 (1:1), 20% Knockout Serum Replacement (KOSR), 1% MEM non-essential amino acids, 1% L-glutamine, 0.2 mM β-mercaptoethanol, 100 ng/ml FGF-2), and iPSCs cultured on Matrigel were cultured with either mTeSR1 or StemFlex media. MEFs, FGF-2, and Matrigel were purchased from WiCell (Madison, WI). All other cell culture reagents were purchased from ThermoFisher Scientific. Karyotype analysis was performed as a quality control. The manuscript does not contain human subject or animal studies, and all work with iPSC lines was carried out in accordance with institutional, national, and international guidelines and approved by the Stem Cell Research Oversight Committee at the University of Wisconsin-Madison.

### Differentiation of iPSC lines to RPE

Differentiation of iPSCs to RPE was performed as previously described (17, 42). Briefly, iPSCs were enzymatically lifted (1 mg/ml dispase for cells cultured on MEFs; 2 mg/ml dispase or 1 ml ReLeSR for cells cultured on Matrigel) to form aggregates, also referred to as embryoid bodies (EBs). EBs were maintained in suspension culture either in EB media (iPS media without FGF-2) and then switched to neural induction media (NIM) on day 4, or gradually weaned off mTeSR1/StemFlex and transitioned to NIM by day 4. NIM is composed of 500 ml DMEM/F12 (1:1), 1% N2 supplement, 1% MEM non-essential amino acids, 1% L-glutamine, 2 µg/ml heparin. EBs were plated on laminin (Cat# 23017015) coated 6-well plates (Nunc; Thermo Fisher Scientific) on day 7. On day 16, neural rosettes were mechanically lifted, leaving adherent cells behind that were maintained in retinal differentiation media (RDM; DMEM:F12 (3:1), 2% B27 without retinoic acid, 1% antibiotic-antimycotic solution). For the first four media changes, RDM was supplemented with 10 µM SU5402 and 3 µM CHIR99021.

After 60 days of differentiation, pigmented patches of RPE were micro-dissected, dissociated using Trypsin-EDTA (0.25%), and plated on laminin coated surfaces in RDM with 10% FBS and Rho kinase inhibitor (ROCKi; Y-27632). After 2 days, the media was changed to RDM with 2% FBS, and eventually to RDM once the cells were fully confluent. There were no differences observed between RPE differentiated from iPSCs cultured on MEFs and Matrigel. Mutant and wildtype genotypes of iPSC-RPE were verified by Sanger sequencing periodically. Heparin (Cat# H-3149) and SU5402 (Cat# SML0443-25MG) were from Sigma-Aldrich, CHIR99021 (Cat# 4423) was from Tocris Bioscience, and ReLeSR was purchased from STEMCELL Technologies. All other differentiation reagents were purchased from ThermoFisher Scientific.

### Gene expression analysis

Reverse transcriptase-PCR was used to assess RPE-specific gene expression in RPE derived from different iPSC lines, as described previously (17). Primers used are listed in Table S1.

### Generation of lentiviral vectors

Lentiviral plasmid with the human *VMD2* promoter driving expression of *hBEST1-T2A-GFP* was provided by Alfred S. Lewin (University of Florida). LentiCRISPR v2 (LCv2) plasmid was purchased from Addgene (Cat# 52961). Lentiviral gene editing plasmids containing specific sgRNA sequences and the human *VMD2* promoter driving expression of *spCas9-T2A-GFP* were then generated as described hereafter (all primers and sgRNA sequences are listed in SI Tables). To begin, the ‘*T2A-GFP-WPRE*’ sequence was amplified from the *hVMD2*-*hBEST1*-*T2A-GFP* plasmid using LCv2-GFP.Gib.F and .R primers and Q5 2X MM (NEB, Cat# M0492L). The ‘*2A-Puro-WPRE*’ sequence was then removed from the LCv2 plasmid via restriction digestion with PmeI (NEB, Cat# R0560S) and BamHI (NEB, Cat# R3136S). The digestion product was resolved on a 0.7% agarose gel and the plasmid backbone was purified using the Monarch gel purification kit (NEB, Cat# T1020S). The ‘*T2A-GFP-WPRE*’ sequence was inserted into the digested backbone using the Gibson Assembly kit (SGI, Cat# GA1100) per the manufacturer’s instructions. The completed Gibson Assembly reaction was then amplified using chemically competent *E. coli* (NEB, Cat# C3040H) and Sanger sequenced to confirm insertion of ‘*T2A-GFP-WPRE*’ using LCv2-GFP.seq.L and LCv2-GFP.seq.R primers. This intermediate plasmid product (*pLCv2-GFP*) was digested with AfeI (NEB, Cat# R0652S) and EcoRI-HF (NEB, Cat R310S) to remove the constitutive EF-1 alpha core promoter. The desired digestion product was purified as described above. The *hVMD2* promoter was then PCR amplified from *hVMD2*-*hBEST1-T2A-GFP* using Q5 2X MM and VMD2.LCv2.GFP.Gib.F and .R primers, followed by insertion into the digested LCv2-GFP backbone via Gibson Assembly. Next, the completed Gibson reaction was transformed into chemically competent *E. coli* and the sequence of the final product *hVMD2*-*spCas9-T2A-GFP* was confirmed via Sanger sequencing using VMD2.LCv2.GFP.seq.L and .R primers. Subsequently, specific sgRNAs were cloned into *hVMD2*-*spCas9-T2A-GFP* using the restriction digest and Gibson Assembly protocol.

### Lentivirus production and cell transduction

Lentivirus stocks were generated by the Cell Culture Core of the UW Department of Dermatology Skin Disease Research Center (Madison, WI). Briefly, HEK293 cells cultured on 10-cm dishes were transfected with lentiviral plasmids—10 µg of sgRNA encoding lentiviral plasmid (*hVMD2*-*hBEST1-T2A-GFP or hVMD2*-*spCas9-T2A-GFP*); 5 µg of psPax2 (Addgene, Cat# 12260), and 2 µg of pMD2.G (Addgene, Cat# 12259)—using Lipofectamine (ThermoFisher; Cat# 11668019). After 15 hours, culture medium (DMEM with 10% FBS) was replaced with fresh media containing 1% Penicillin-Streptomycin. Media containing lentiviruses was collected the next day and viral titers were calculated using QuickTiter Lentivirus Titer Kit (Cell Biolabs, Cat# VPK-107). Titers for lentiviral stock were:

| Lentivirus | Titer (Transduction units/ml) |
| --- | --- |
| *hVMD2-hBEST1-T2A-GFP* | $22 \times 10^{6-7}$ |
| *hVMD2-spCas9-T2A-GFP* R218C sgRNA | $74.16 \times 10^{6-7}$ |
| *hVMD2-spCas9-T2A-GFP* A146K sgRNA | $74.26 \times 10^{6-7}$ |
| *hVMD2-spCas9-T2A-GFP* N296H sgRNA | $68.91 \times 10^{6-7}$ |
| *hVMD2-spCas9-T2A-GFP AAVS1* sgRNA | $74.01 \times 10^{6-7}$ |

For iPSC-RPE transduction, monolayers of iPSC-RPE on transwells were treated with 0, 5, 50, or 150 µl (Figure S3) or 150 µl alone of specified lentivirus preparation for all other experiments. Media was changed on day 2 to RDM, and cells were maintained in culture with media changes every 3 days until used for sequencing or other analyses.

### Transepithelial electrical resistance (TER) measurements

Monolayers of RPE cultured on transwell inserts (Corning, #3470) were used for all TER measurements. To perform the measurements, we employed an epithelial voltohmmeter (EVOM2) with chopstick electrodes (STX2) from World Precision Instruments (Sarasota, USA) according to manufacturer’s instructions. Electrodes were sterilized with ethanol, and then rinsed in sterile Milli-Q water followed by HBSS before measuring electrical resistance of RPE monolayers. Differences between TER values of transwells with cultured RPE monolayers versus background measurements of cell-free transwell inserts were multiplied by the surface area of the transwell membrane to obtain net TER values in Ω · cm^2^.

### Calcium-activated chloride channel current density measurements

All iPSC-RPE cells used for chloride current measurements were cultured as a monolayer on transwells. To singularize cells prior to measurement, transwells were washed twice with 0 Na-CMF solution (135 mM N-Methyl-D-glucamine (NMDG)-Cl, 5 mM KCl, 10 mM HEPES, 10 mM glucose, 2 mM EDTA-KOH, pH adjusted to 7.4) and then incubated with papain enzyme solution (0 Na-CMF solution containing 2.5 µl/ml papain (46 mg/ml, MP Biomedicals LLC, Cat#100921), 0.375 mg/ml adenosine, 0.3mg/ml L-cysteine, 0.25 mg/ml L-glutathione, and 0.05mg/ ml taurine) for 30 minutes at 37°C/5% CO_2_. To stop the reaction, 0.01% BSA was added to the enzymatic solution. After washing twice with 0 Na-CMF solution, cells were dispersed in extracellular solution containing 140 mM NaCl, 10 mM HEPES, 3 mM KCl, 2 mM CaCl_2_, 2 mM MgCl_2_, and 5.5 mM glucose adjusted to pH 7.4 with NaOH by gentle pipetting.

Cells with polarized RPE morphology post-dissociation (Figure 2B, *left*) were used to measure chloride currents. To test effects of gene augmentation or gene editing on *BEST1* mutant iPSC-RPE by single-cell patch clamp analysis, only cells with GFP fluorescence (from transduction with *hVMD2-hBEST1-T2A*-*GFP* for gene augmentation or *hVMD2-spCas9*-*T2A*-*GFP* encoding *AAVS1* sgRNA or mutant allele-targeted sgRNAs for gene editing) were used. Current recordings on these cells were performed using the conventional whole-cell patch clamp technique with an Axopatch 200A amplifier controlled by Clampex software program via the digidata 1550 data acquisition system (Axon Instruments, CA). Fire-polished borosilicate glass pipettes with 3-5 MΩ resistance were filled with pipette solution containing 4.5 µM calcium or no calcium.

Recordings were carried out at room temperature and current-voltage tracings were established using ramps from -100 to +100 mV for 1000 ms. The pipette solution with calcium was comprised of (in mM) 146 CsCl, 5 (Ca^2+^)-EGTA-NMDG, 2 MgCl_2_, 8 HEPES, and 10 sucrose at pH 7.3, adjusted with NMDG. Another pipette solution devoid of calcium was comprised of (in mM) 146 CsCl, 5 EGTA-NMDG, 2 MgCl_2_, 8 HEPES, and 10 Sucrose at pH 7.3, adjusted with NMDG. Both of these pipette solutions were mixed to make the solution containing 4.5 µM free calcium as described previously(43), which was then used for patch clamping.

Current density values were obtained by dividing current amplitude with cell capacitance measurements. CaCC current densities for iPSC-RPE are represented as differences between mean 4.5 µM calcium response and mean no calcium response from a total of at least five cells for each condition. At least two differentiations were used as replicates to obtain data for each line.

### Immunocytochemistry

iPSC-RPE cultured on transwell inserts were washed with PBS and fixed with 4% paraformaldehyde for 10 minutes at room temperature (RT). After washing fixed cells three times with PBS, transwell membranes were placed in blocking solution (10% normal donkey serum with 5% BSA, 1% fish gelatin and 0.5% Triton-X100 in PBS) for one hour at RT, and then incubated overnight at 4 °C in primary antibody (1:100 mouse anti-Bestrophin (Millipore, Cat# MAB5466); 1:100 rabbit anti-ZO-1 (ThermoFisher Scientific, Cat# 61-7300)) prepared in blocking solution. Cells were then washed three times in PBS and incubated for 30 minutes at RT in appropriate secondary antibody (ThermoFisher Scientific; 1:500 Donkey anti-Mouse IgG (Cat# A31571); 1:500 Donkey anti-Rabbit IgG (Cat# A10040)) prepared in blocking solution. Cells were again washed three times in PBS, incubated in DAPI (1:500; ThermoFisher; Cat# D1306) for 30 minutes, mounted using prolong gold with DAPI (ThermoFisher; Cat# P36931), and imaged using Nikon A1R confocal microscope with NIS Elements AR 5.0 software.

### Rhodopsin degradation assay

Photoreceptor outer segment (POS) feeding of iPSC-RPE was performed as described previously (17). Briefly, bovine POS (InVision BioResources (Seattle, WA)) were gently resuspended in DMEM. 100 µl media was then removed from each transwell insert, 6.25×10^6^ POS were added, and cells were incubated at 37 °C and 5% CO_2_ for 2 hours. Afterward, POS containing RDM was removed and each transwell was washed thoroughly three times using DPBS. Following the washes, cells were harvested (0 time point) or further incubated in fresh RDM for prescribed periods of time. At each time point, transwells were washed, 100 µl RIPA buffer (ThermoFisher; Cat# 89900) containing protease inhibitor cocktail (Sigma-Aldrich; Cat# P8340) was added, and cells were incubated on ice for 30 minutes to extract total cell protein. Protein quantification was performed using the DC Protein assay kit II (Bio-Rad, Cat# 5000112).

Western blots were then performed to monitor rhodopsin degradation as described (17, 18). Briefly, protein lysates were denatured in 1X Laemmli buffer (reducing) and kept on ice for 10 minutes. Protein samples were then separated on 4-20% mini-Protean TGX gels (Bio-Rad; Cat# 4568095) and electroblotted onto PVDF membranes (Millipore; IPFL10100). After blotting, membranes were dried at RT for 15 minutes, re-activated in methanol for 1 minute, and then incubated in blocking buffer (1:1 Odyssey blocking buffer (LI-COR Biosciences; Cat# 927-40000):PBS) for 1 hour. Post-blocking, blots were incubated in primary antibodies (1:500 mouse anti-rhodopsin (Millipore, Cat# MABN15); 0.1 µg/ml rabbit anti-beta actin (Abcam, Cat# ab8227)) in blocking buffer with 0.1% Tween-20 overnight, washed three times for 5 minutes each in PBS with 0.1% Tween-20, incubated for 1.5 hours at RT in appropriate secondary antibody (LI-COR Biosciences; 1:20,000 Donkey anti-Rabbit IgG (Cat# 926-32213); 1:20,000 Donkey anti-Mouse IgG (Cat# 926-68022)) in blocking buffer with 0.1% Tween-20 and 0.01% SDS, and then washed three times for 5 minutes each in PBS with 0.1% Tween-20. An Odyssey infrared Imager (LI-COR Biosciences) was used to image blots using Image Studio software. ImageJ was used for quantification of relevant protein bands. Samples from rhodopsin degradation assays were also used to assess levels of BEST1 protein before and after gene augmentation. Western blots were performed as described above, using 1:1000 rabbit anti-Bestrophin1 antibody (LAgen Laboratories; Cat# 016-Best1-01) and 1:1000 mouse anti-Actin antibody (Millipore; Cat# MAB1501) as primary antibodies.

### Deep sequencing analysis of DNA and RNA read frequency

Cells were singularized with TrypLE Express (Gibco, Cat# 12605010) per manufacturer’s instructions. Total DNA and/or RNA was extracted using QuickExtract DNA (Epicentre, Cat# QE09050) or QuickExtract RNA (Epicentre, Cat# QER090150), respectively. Both DNA and RNA extractions were performed per manufacturer’s instructions with the following minor modifications: 1) a ratio of 10,000-25,000 cells per 50 µl of QuickExtract solution was routinely used, and 2) an optional DNase 1 treatment was omitted from the RNA extraction protocol. All samples were stored at -80 °C until use. RNA was reverse transcribed to cDNA using the ProtoScript II First Strand synthesis kit (NEB, Cat# E6560S) and synthesis was performed with the “random primer” option included within the kit. 4 µl of crude RNA extract was added to each cDNA reaction.

In preparation for targeted deep sequencing, Illumina adapter sequences and sample-specific barcodes were appended to genomic or cDNA amplicons via overhang PCR as described (19). Purified amplicon libraries were assembled into 2 nM total DNA in DNAse/RNAse free H_2_O and sequenced using 150 nucleotide paired end reads using MiSeq (6M or 15M total reads) at the UW Biotech Center (Madison, WI) with the following loading condition: 8 pmol total DNA and 15% PhiX DNA. Raw FASTQ files were read and aligned to expected amplicons using a command line implementation of CRISPResso (v1.0.8) (44). Full commands used for analysis are available upon request. ‘Percent allele identity’ or ‘percent edited’ were determined using the software’s standard output table of individual read identities. Sequencing reads with counts <100 were not included in the analysis. All FASTQ files are available upon request.

### Single-cell RNA sequencing (scRNA-seq)

iPSC-RPE cultures derived from the A146K, N296H, and R218C adBD patient lines, and from an isogenic gene-corrected control line derived from the R218C line (R218C>WT) were transduced with 150 µl of *hVMD2*-*spCas9*-*T2A-GFP* encoding specific sgRNAs as described in the ‘Lentivirus production and cell transduction’ section. For each sample, sgRNAs were either targeted to mutant *BEST1* or to the *AAVS1* locus (control). On day 14, cells were dissociated from transwells with a papain dissociation kit (Worthington Biochemical, Cat# LK003150) and filtered using a Flowmi cell strainer (Bel-Art SP Scienceware, Cat# H13680-0040) to obtain single-cell suspension. Cells were then prepared for scRNA-seq with the droplet-based 10X Genomics GemCode platform according to the manufacturer’s instructions. In brief, singularized cells were encapsulated in oil beads containing a unique molecular identifier (UMI) barcode. The cells were then lysed and cDNA libraries were created featuring cell and transcript-specific molecular identifiers. Libraries were sequenced using an Illumina HiSeq2500 Rapid Run and reads were aligned to a custom reference genome consisting of the human hg19 GRCh38 genome and an added gene for the *spCas9*-*T2A-GFP* transcript.

### scRNA-seq data analysis

Gene edited iPSC-RPE were clustered based on their genome-wide transcriptome using the t-Distributed Stochastic Neighbor Embedding (t-SNE) algorithm with the 10X Genomics Loupe Cell Browser software (v2.0.0). Reads for each pair of samples (*BEST1* mutant allele-targeted sgRNA vs *AAVS1* sgRNA control) were aligned, analyzed, clustered with Cell Ranger v2.1.1, and compared to detect significant differences in gene expression, with p values adjusted using the Benjamini-Hochberg correction for multiple tests. P <0.01 was used as the significance threshold for all analyses. Cell Ranger using the aggregate feature was run to concatenate each pair of samples with the same genotype, and differential gene expression within each pair (with gene editing at either the *AAVS1* or *BEST1* locus) was then analyzed. Potential adverse events were probed using gene lists curated from gene ontology terms associated with the cell cycle, apoptosis, DNA damage response, and the innate immune response, as well as a list of 149 validated marker genes associated with human RPE (45) (**SI data file B**; gene ontology sets are available on the Molecular Signatures Database <http://software.broadinstitute.org/gsea/msigdb>). Differentially-expressed genes with p <0.01 were deemed to be significant. All significantly differentially-expressed genes per cluster are reported, with the exception of genes identified by Cell Ranger as having low average UMI counts. Volcano plots were generated in RStudio (v.1.1.456) using the ggplot2 package.

### Non-negative matrix factorization-based comparison of scRNA-seq datasets

Non-negative matrix factorization (NMF) followed by clustering of genes using the NMF factors was used for Figure S4 to project each dataset into a gene group. The input data for this analysis were a set of gene barcode matrices generated using the Cell Ranger 2.1.1 algorithm. The matrices were filtered to remove background barcodes in order to include only detected cellular barcodes, and then further filtered to exclude cells expressing fewer than 2000 total counts, followed by depth normalization.

To enable comparison of transcriptional signatures from each sample, NMF (46) was applied to each scRNA-seq dataset. NMF is a popular dimensionality reduction and clustering approach that is used to project data into low dimensional non-negative factors, and thus can be used to derive a clustering of cells and genes. NMF with k=10 factors was applied with a total of five NMF runs. Next, the similarity of NMF results was compared between two samples using the average best Jaccard coefficient between clusters of one versus another sample. 1-average Jaccard coefficient was then used as the distance to apply hierarchical clustering on the samples. This procedure was repeated five times and the tree that appeared most often was used. The trees learned in different iterations were largely similar and always grouped the patient-specific lines first before grouping different lines together.

### Quantification and statistical analysis

Unless otherwise specified, all analyses were performed using GraphPad Prism (v.8.0.1) and error bars represent mean ± SD; ns = p ≥0.05, * for p <0.05, ** for p <0.01, *** for p <0.001, **** for p <0.0001. Further detail for each analysis is provided here. Statistical analyses for Figures 2E, 2I and 4B were performed using Origin 2018b. Student’s *t*-test was performed to measure the significance between the groups. P values <0.05 were considered statistically significant. Statistical significance for Figure 4D and S3C was determined using the Holm-Sidak method with alpha = 0.05. Each row was analyzed individually, without assuming a consistent SD (number of *t* tests = 10 and 2 for Figure 4D, and S3C, respectively). Statistical significance for differential gene expression in Figures 4F and Figure S4G was determined using the Cell Ranger 2.1.1 algorithm. Sample pairs with each genotype were analyzed and clustered with individual Cell Ranger runs for each pair and analyzed using the Loupe Cell Browser (v.2.0.0). Differential expression was calculated using a negative binomial exact test, and p values were adjusted using the Benjamini-Hochberg correction for multiple tests. P <0.01 was used as the threshold for assigning significant versus non-significant changes in gene expression. Volcano plots were generated in RStudio (v 1.1.456) using the ggplot2 package. For Figures 3K, L, M, and S3B, discovery was determined using the two-stage linear step-up procedure of Benjamini, Krieger, and Yekutieli with Q = 1%. Each row was analyzed individually, without assuming a consistent SD (number of *t* tests = 3).

### Data and Software availability

Upon acceptance, scRNA-seq data will be posted to an accession database. Raw targeted sequencing files for DNA and RNA sequencing data will be deposited to the NCBI Trace and Short-Read Archive. Raw patch clamp data are available upon request. Other experimental data are provided in Supplemental files and all source data are available upon request.

## Supporting Information

**Figure S1.**
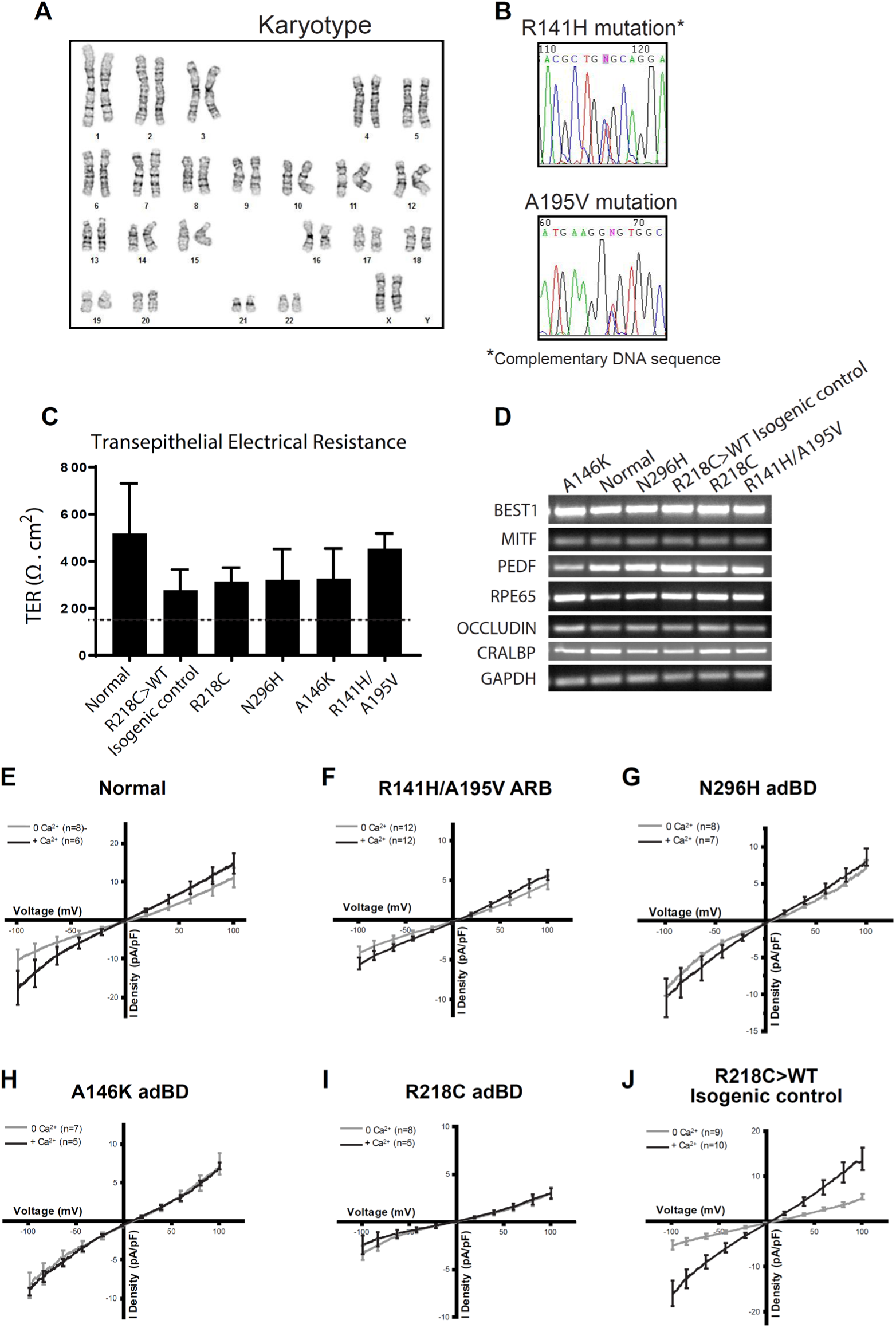
Characterization of iPSC-RPE. **(A)** Karyotype analysis for ARB iPSCs. **(B)** DNA sequencing confirming R141H and A195V encoding mutations in ARB iPSCs. **(C)** Net transepithelial electrical resistance (TER) (Ω · cm^2^) for iPSC-RPE from all six lines. The dashed line demarcates the minimum expected TER (150 Ω · cm^2^). Replicates: n=12 for each line (4 transwells from 3 replicates each), error bars represent mean ± SD. **(D)** Gene expression analysis (RT-PCR) of selected RPE-specific markers in all six lines. **(E-J)** Chloride current traces, measured in the presence (*black*) or absence (*gray*) of calcium over a voltage ramp (-100 to +100 mV), that were used to generate CaCC current density plots in Figure 1E. 4.5 µM calcium was used for +calcium conditions. The number (n) of individual cells patch clamped in the presence or absence of calcium in order to calculate CaCC current densities is shown in the top left corner of each graph. Data were obtained from at least two replicates.

**Figure S2.**
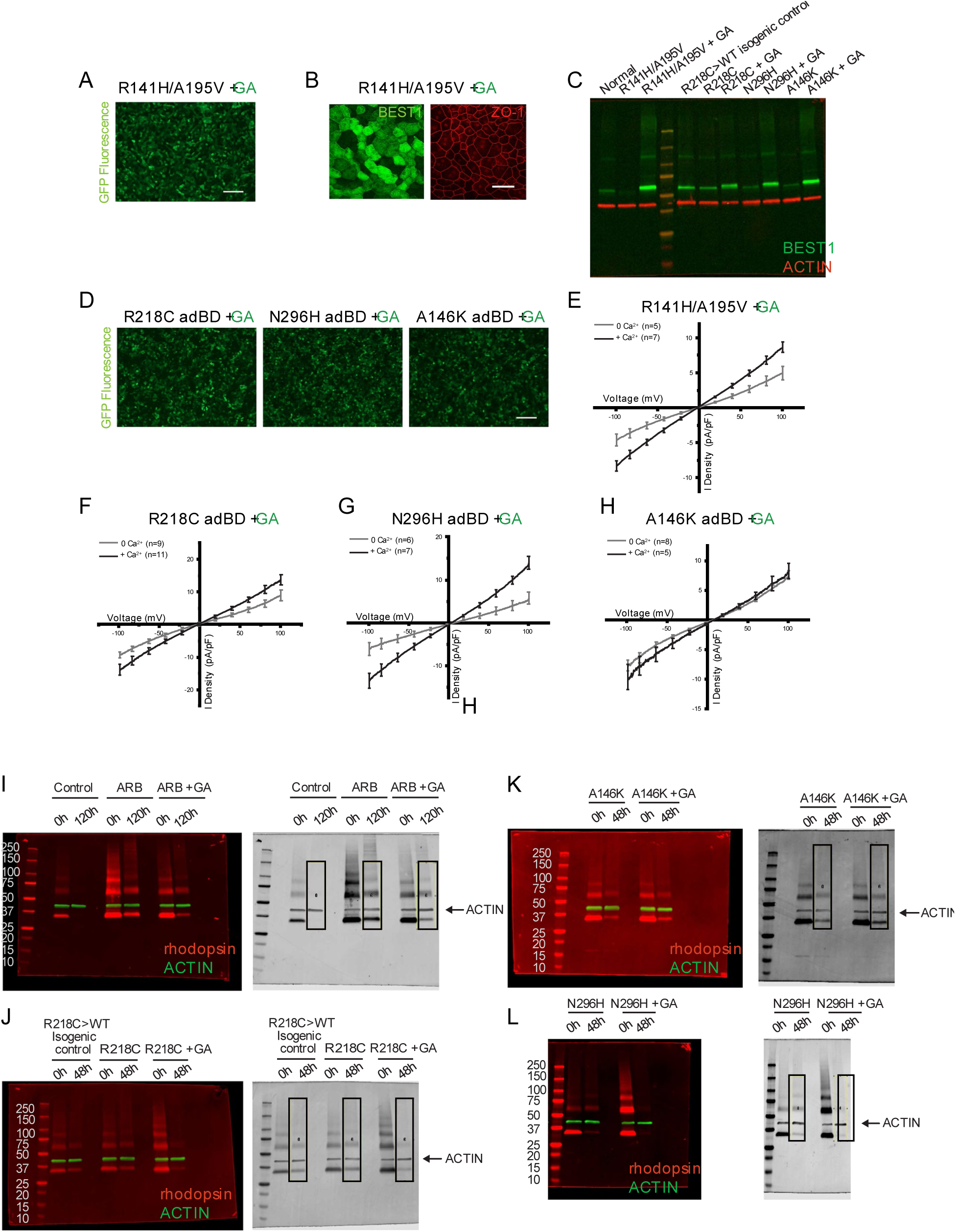
Gene augmentation (GA) restores CaCC function in ARB iPSC-RPE and R218C and N296H adBD iPSC-RPE, but not in A146K adBD iPSC-RPE. **(A)** GFP fluorescence in R141H/A195V ARB iPSC-RPE transduced with lentivirus expressing BEST1. Scale bar = 100 µm. **(B)** ICC analysis of BEST1 and ZO-1 expression in R141H/A195V iPSC-RPE transduced with lentivirus expressing BEST1. Increased BEST1 expression is observed in R141H/A195V iPSC-RPE cells following gene augmentation. Scale bar = 50 µm (applies to both images). **(C)** Representative western blot showing levels of BEST1 in iPSC-RPE. Protein samples from the rhodopsin degradation assays were used to assess BEST1 levels. **(D)** GFP fluorescence in adBD iPSC-RPE transduced with lentivirus expressing hBEST1. Scale bar = 100 µm (applies to all three images). **(E)** Chloride current traces of R141H/A195V iPSC-RPE after gene augmentation measured in the presence (*black*) or absence (*gray*) of calcium. **(F-H)** Chloride current traces for adBD iPSC-RPE after gene augmentation, measured in the presence (*black*) or absence (*gray*) of calcium over a voltage ramp (-100 to +100 mV), that were used to obtain CaCC current density. 4.5 µM calcium was used for +calcium conditions. Cells with green fluorescence were used for all patch clamp measurements after gene augmentation. The number (n) of individual cells patch clamped in the presence or absence of calcium (in order to calculate CaCC current densities) is shown in the top left corner of each graph. Data were obtained from at least two replicates. (I-L) *left*, Western blots used for the rhodopsin degradation assay, *right*, and corresponding grayscale images of western blots used to quantify levels of rhodopsin shown in Figures 2 and 3 (boxes represent areas used for quantification). For each lane, the boxed area was selected to include bands corresponding to fully denatured rhodopsin and its aggregated forms.

**Figure S3.**
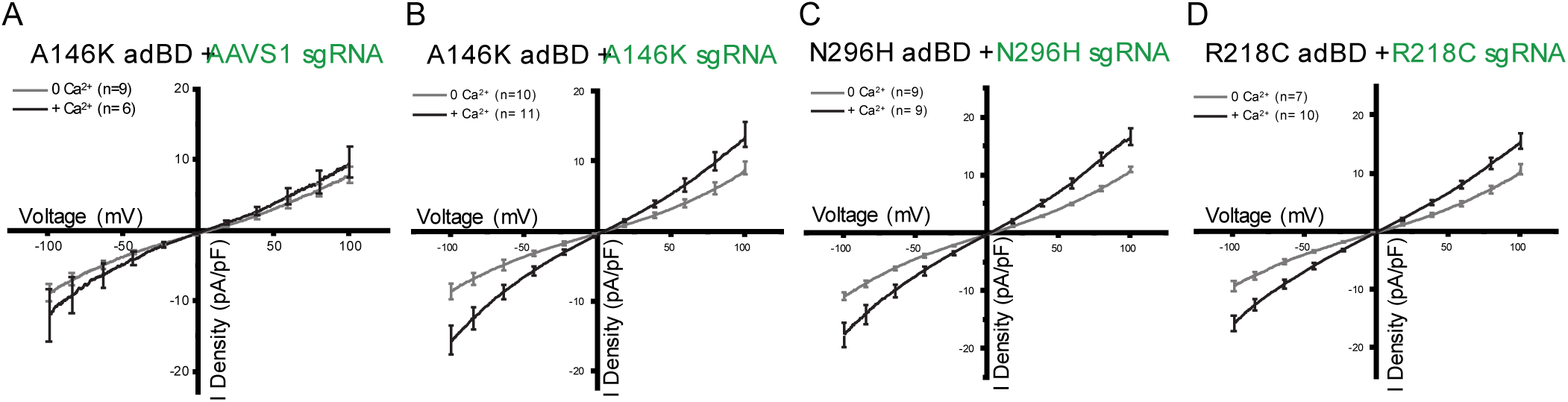
Gene editing (GE) restores CaCC activity in iPSC-RPE from all tested adBD lines. **(A-D)** Chloride current traces, measured in the presence (*black*) or absence (*gray*) of calcium over a voltage ramp (-100 to +100 mV), that were used to calculate CaCC current density plots after gene editing of adBD iPSC-RPE. iPSC-RPE was edited using lentiviral genome editors encoding sgRNA targeting **(A)** *AAVS1* site in A146K adBD iPSC-RPE, **(B)** A146K mutation in A146K adBD iPSC-RPE, **(C)** N296H mutation in N296H adBD iPSC-RPE, or **(D)** R218C mutation in R218C adBD iPSC-RPE. Cells with GFP fluorescence were used for whole cell patch clamp measurements and 4.5 µM calcium was used for +calcium conditions. The number (n) of individual cells patch clamped with or without calcium is shown at the top left corner of each graph. Data were obtained from two replicates.

**Figure S4.**
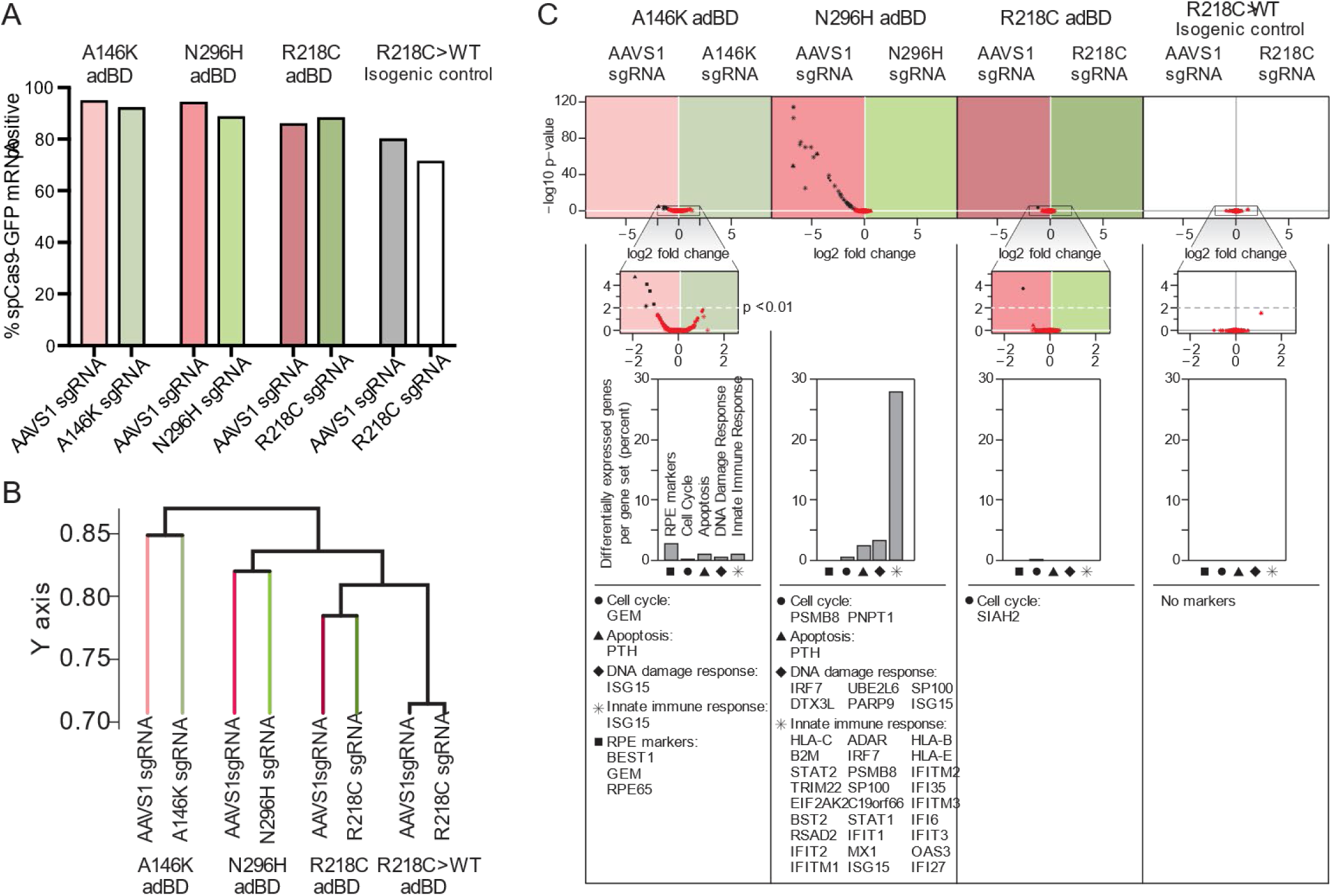
Single cell transcriptome analysis in gene-edited adBD iPSC-RPE. **(A)** Percent of analyzed cells per sample for which *spCas9-T2A-GFP* transcripts were captured using scRNA-seq. **(B)** Dendrogram tree depicting relative similarity between samples. Non-negative matrix factorization comparison across samples indicates that greater transcriptional variability exists between iPSC-RPE lines than in the same iPSC-RPE line treated with lentiviral genome editors (*AAVS1* lentiviral genome editor versus *BEST1* mutant allele-targeted lentiviral genome editor). The dendrogram shows the similarity of the transcriptomes from each sample, derived from the average Jaccard coefficient between gene clusters from one sample and those from another sample. The y-axis denotes 1-average Jaccard coefficient and indicates the distance between different samples (tree tips) as well as between groups of samples (internal nodes). **(C)** Differential gene expression in 5 curated gene sets associated with cell cycle regulation (*circles*), apoptosis (*triangles*), DNA damage response (*diamonds*), innate immune response (*asterisks*), or RPE-identity (*squares*) in control (*AAVS1*) lentiviral genome editor versus mutant allele-targeted lentiviral genome editor treated samples. For one sample pair (N296H iPSC-RPE), genes associated with a potential adverse treatment effect were upregulated in control lentiviral genome editor-treated sample compared to the mutant allele-targeted lentiviral genome editor sgRNA-treated sample.

**SI Tables** (attached below):

Table S1: RPE-specific RT-PCR primers used.

Table S2. List of GE vectors used.

Table S3. List of primers for lentiviral plasmid generation.

Table S4. List of sgRNAs.

Table S5. Primers for deep sequencing of DNA and cDNA.

**SI data files** (available for download):

SI Data File A. Frameshift analysis of iPSC-RPE+GE.

SI Data File B. Curated gene sets used to assess differences in gene expression between control (*AAVS1*) and mutant *BEST1* allele-targeted sgRNA.

SI Data File C. Ranked off-target sites for sgRNAs used in this study.

SI Data File D. Analysis of additional adBD mutations for amenability to allele-specific editing or scarless base editing.

## SI Tables

**Table S1:**
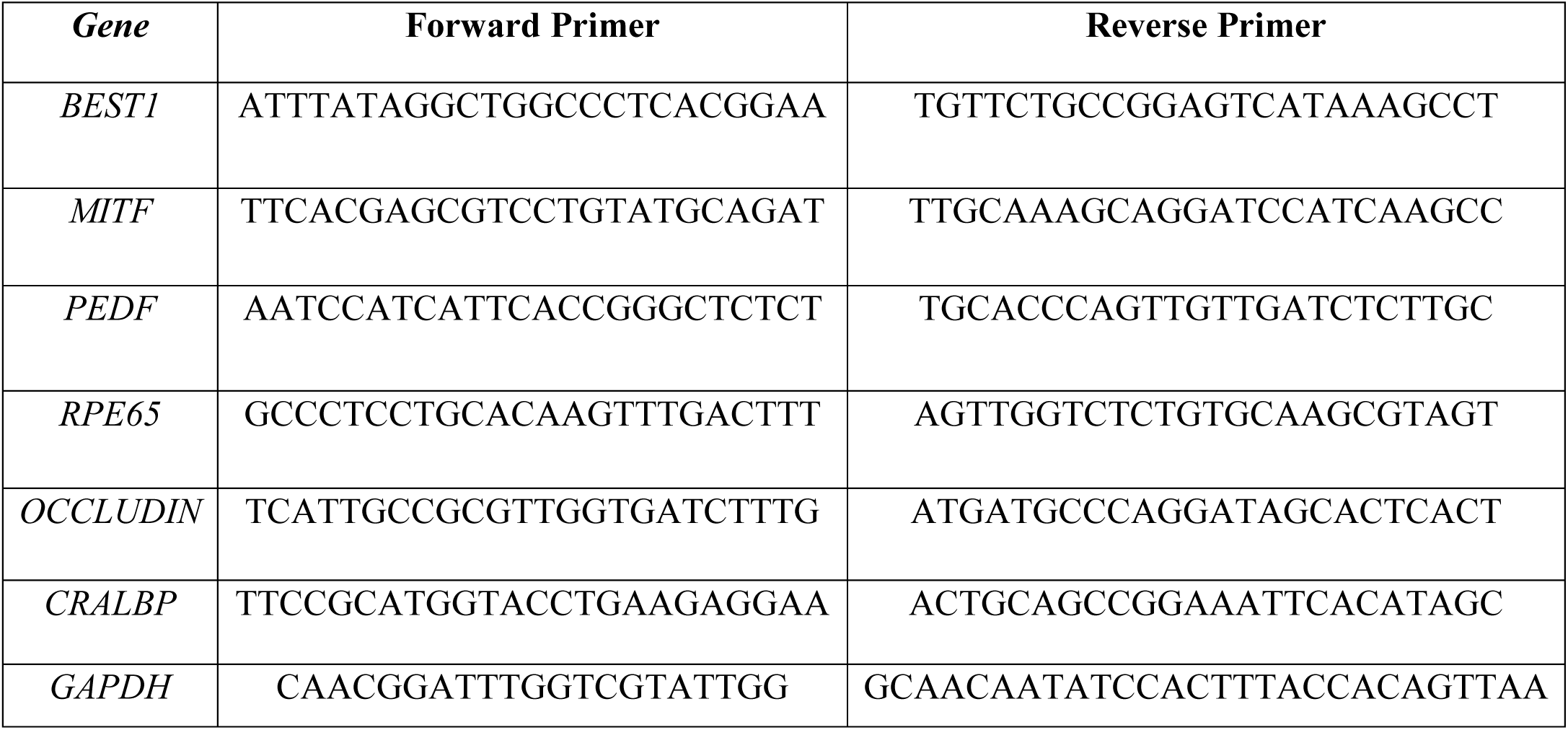
RPE-specific RT-PCR primers used.

**Table S2.**
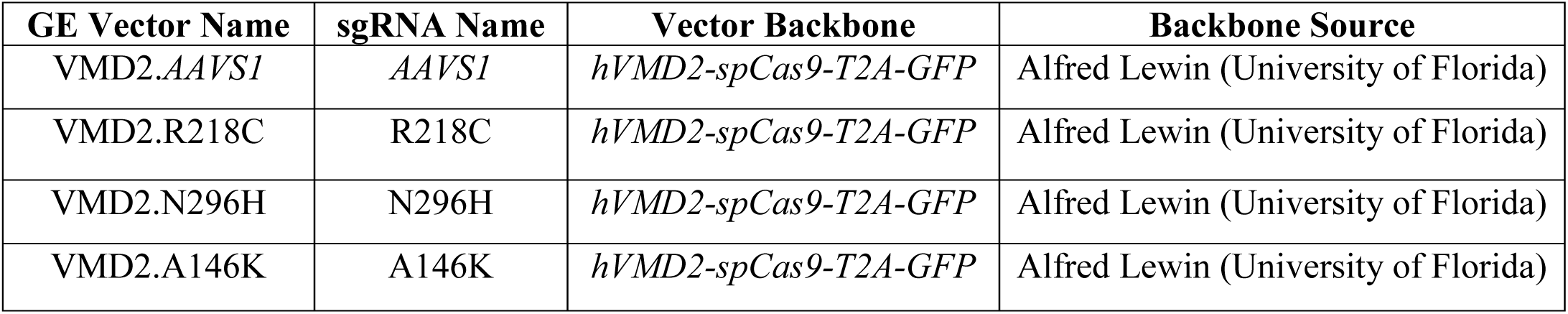
List of gene editing vectors used.

**Table S3.**
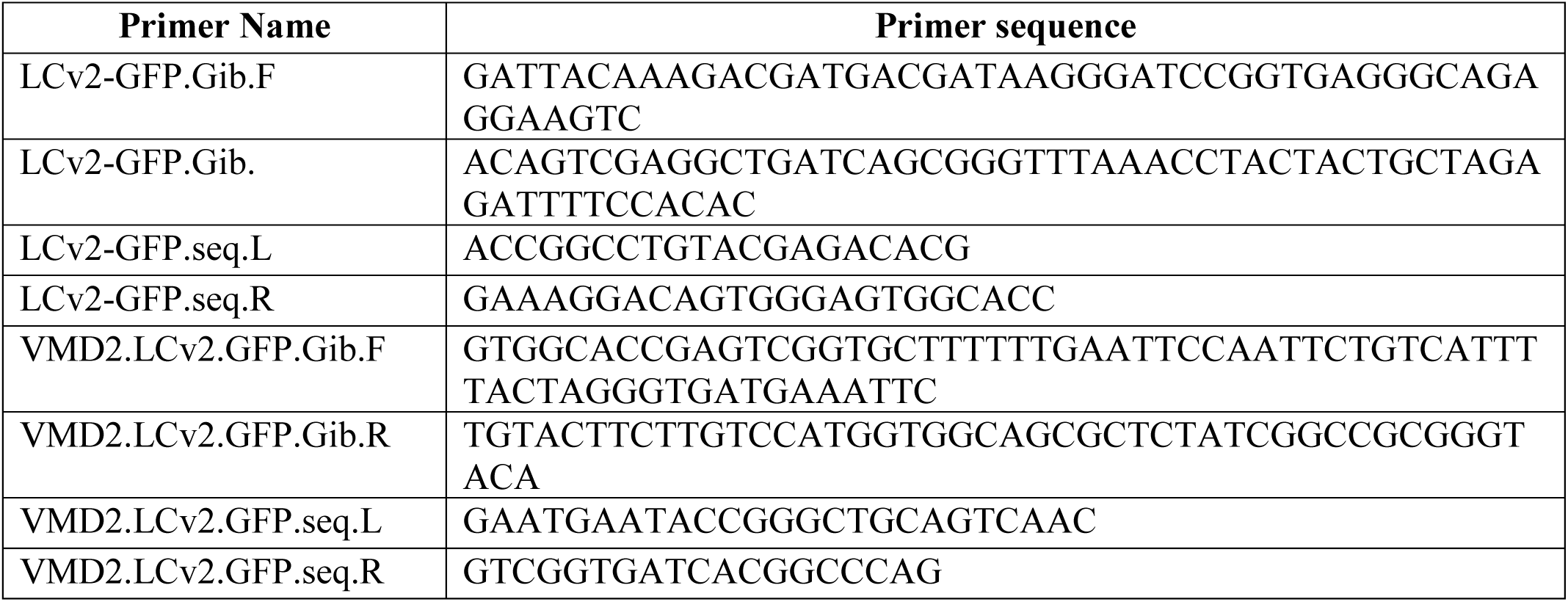
List of primers for lentiviral plasmid generation.

**Table S4.**
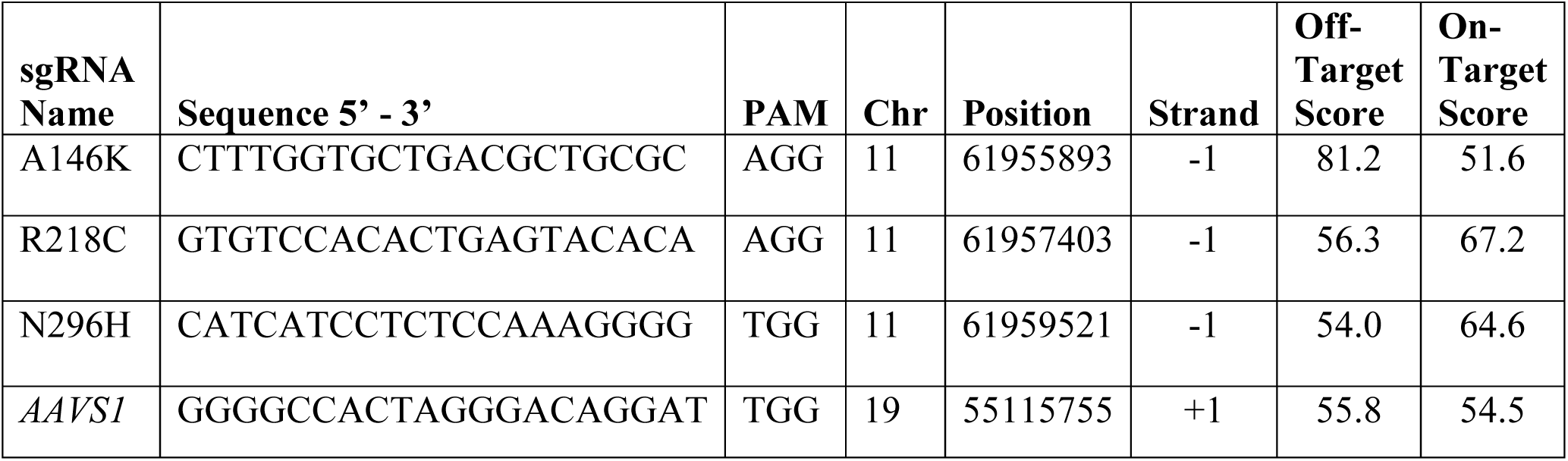
List of sgRNAs. Off-target (47) and on-target (48) scores are also presented. Scores range from 0-100 with higher scores being better for both scoring systems. Highest ranked off-target cut sites for each sgRNA are available in SI Data File C.

**Table S5.**
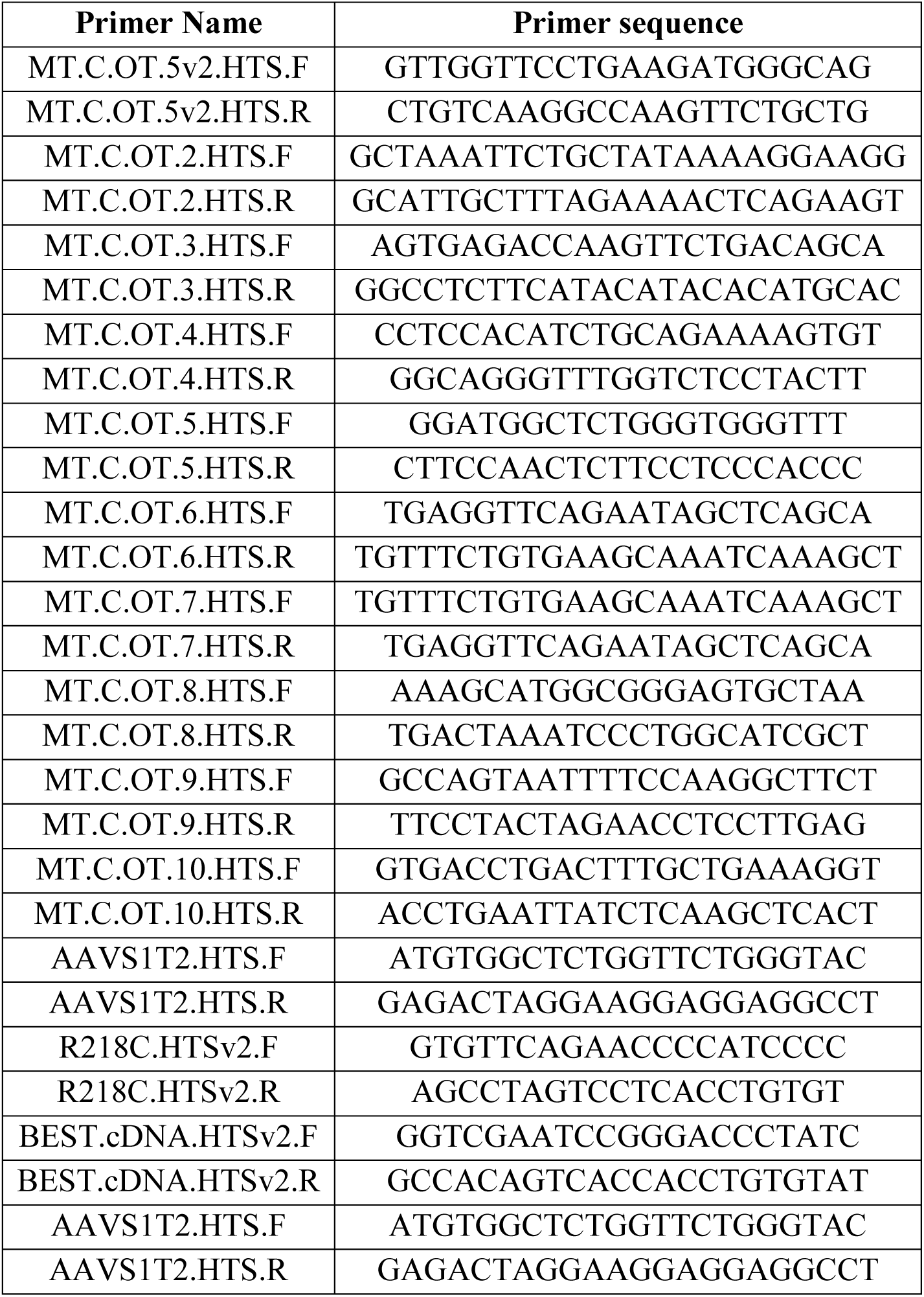
Primers for deep sequencing of DNA and cDNA.

