## Supplementary material for "Human iPSC modeling reveals mutation-specific responses to gene therapy in Best disease": SI Data File A

SI Data File A: Frameshift analysis of iPSC-RPE+GE

R218C MTC sgRNA in R218C RPE

| Seq Batch | Sample # | Sample Name |
|---|---|---|
| 180531_G256N | Sample 8 | 6M150RD |

* these reads are presented in Figure 3f (bottom)

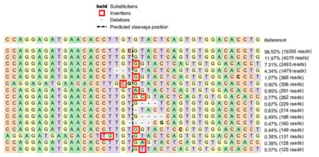


| Aligned_Sequence | Reference_Sequence | Allele | Allele Edited | %Reads | #Reads |
|---|---|---|---|---|---|
| CCAGGAGATGAACACCTTGCGTACTCAGTGTGGACAC | CCAGGAGATGAACACCTTGCGTACTCAGTGTGGACACCTG | WT | N/A | 56.520464 | 19265 |
| CCAGGAGATGAACACCTTGTGTACTCAGTGTGGACAC | CCAGGAGATGAACACCTTGCGTACTCAGTGTGGACACCTG | R218C | N/A | 11.967141 | 4079 |
| CCAGGAGATGAACACCTTGTGGTACTCAGTGTGGACA | CCAGGAGATGAACACCTTGCGTACTCAGTGTGGACACCTG | OUT | R218C | 7.3140678 | 2493 |
| CCAGGAGATGAACACCTTGT-ACTCAGTGTGGACACC | CCAGGAGATGAACACCTTGCGTACTCAGTGTGGACACCTG | OUT | R218C | 4.3391521 | 1479 |
| CCAGGAGATGAACACCTTGTGGTACTCAGTGTGGACC | CCAGGAGATGAACACCTTGCGTACTCAGTGTGGACACCTG | OUT | R218C | 1.0737861 | 366 |
| CCAGGAGATGAACACCTTGCGGTACTCAGTGTGGACA | CCAGGAGATGAACACCTTGCGTACTCAGTGTGGACACCTG | OUT | WT | 0.8977556 | 306 |
| CCAGGAGATGAACACCTTGTAGTACTCAGTGTGGACA | CCAGGAGATGAACACCTTGCGTACTCAGTGTGGACACCTG | OUT | R218C | 0.853748 | 291 |
| CCAGGAGATGAACACCTTGTGCGTACTCAGTGTGGAC | CCAGGAGATGAACACCTTGCGTACTCAGTGTGGACACCTG | OUT | R218C | 0.7686666 | 262 |
| CCAGGAGATGAACACCTTGTG---TCAGTGTGGACACCT | CCAGGAGATGAACACCTTGCGTACTCAGTGTGGACACCTG | IN | R218C | 0.6718498 | 229 |
| CCAGGAGATGAACACCTTGT-TACTCAGTGTGGACACC | CCAGGAGATGAACACCTTGCGTACTCAGTGTGGACACCTG | OUT | R218C | 0.6278422 | 214 |
| CCAGGAGATGAACACCTTGTG--CAGTGTGGACACCT | CCAGGAGATGAACACCTTGCGTACTCAGTGTGGACACCTG | OUT | R218C | 0.4870177 | 166 |
| CCAGGAGATGAACACCTTGCG--CAGTGTGGACACCT | CCAGGAGATGAACACCTTGCGTACTCAGTGTGGACACCTG | OUT | Unknown | 0.4694147 | 160 |
| CCAGGAGATGAACACCTTGTGGTACTCCGTGTGGACA | CCAGGAGATGAACACCTTGCGTACTCAGTGTGGACACCTG | OUT | R218C | 0.4371424 | 149 |
| AGGAGATGAACACCTTGTGTGTACTCAGTGTGGACAC | CCAGGAGATGAACACCTTGCGTACTCAGTGTGGACACCTG | OUT | R218C | 0.3843333 | 131 |
| CCAGGAGATGAACACCTTGTGAGTACTCAGTGTGGAC | CCAGGAGATGAACACCTTGT-GTACTCAGTGTGGACACC | OUT | R218C | 0.3755318 | 128 |
| CCAGGAGATGAACACCTTGTGTTACTCAGTGTGGACA | CCAGGAGATGAACACCTTGTG-TACTCAGTGTGGACACCT | OUT | R218C | 0.3667302 | 125 |

| Allele | % (of total) | percentage | | of indels |
|---|---|---|---|---|
| WT | 56.5% | 56.52 | | |
| R218C | 12.0% | 11.97 | | |
| Indel | 19.1% | 19.07 | | |
| Out of Frame | 18.4% | 18.40 | Out of Frame | 96.48% |
| In Frame | 0.7% | 0.67 | In Frame | 3.52% |

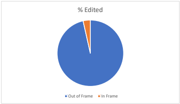


| Seq Batch | Sample # | Sample Name |
|---|---|---|
| 180718_G2GC6 | Sample 3 | 6VM1 |

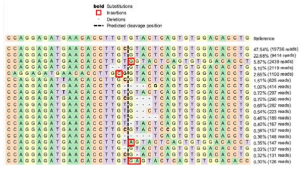


| Aligned_Sequence | Reference_Sequence | Allele | %Reads | #Reads |
|---|---|---|---|---|
| CCAGGAGATGAACACCTTGCGTACTCAGTGTGGACAC | CCAGGAGATGAACACCTTGTGTACTCAGTGTGGACACCTG | WT | 47.5371535 | 19736 |
| CCAGGAGATGAACACCTTGTGTACTCAGTGTGGACAC | CCAGGAGATGAACACCTTGTGTACTCAGTGTGGACACCTG | R218C | 22.6750488 | 9414 |
| CCAGGAGATGAACACCTTGTGGTACTCAGTGTGGACA | CCAGGAGATGAACACCTTGT-GTACTCAGTGTGGACACCT | OUT | 5.87470193 | 2439 |
| CCAGGAGATGAACACCT-TGTACTCAGTGTGGACACC | CCAGGAGATGAACACCTTGTGTACTCAGTGTGGACACCTG | OUT | 5.10393333 | 2119 |
| CAGGAGATGAACACCTTGCGGTACTCAGTGTGGACAC | CAGGAGATGAACACCTTG-TGTACTCAGTGTGGACACCTG | OUT | 2.64951707 | 1100 |
| CCAGGAGATTAACACCTTGCGTACTCAGTGTGGACAC | CCAGGAGATGAACACCTTGTGTACTCAGTGTGGACACCTG | WT | 1.50540742 | 625 |
| [illegible] | [illegible] | OUT | 0.99718188 | 414 |
| [illegible] | [illegible] | MT | 0.71536961 | 297 |
| [illegible] | [illegible] | IN | 0.69850904 | 290 |
| [illegible] | [illegible] | IN | 0.67923983 | 282 |
| [illegible] | [illegible] | OUT | 0.53712937 | 223 |
| [illegible] | [illegible] | OUT | 0.4552352 | 189 |
| [illegible] | [illegible] | OUT | 0.40224486 | 167 |
| [illegible] | [illegible] | WT | 0.37815834 | 157 |
| [illegible] | [illegible] | OUT | 0.35648048 | 148 |
| [illegible] | [illegible] | OUT | 0.35407183 | 147 |
| [illegible] | [illegible] | OUT | 0.32998531 | 137 |
| [illegible] | [illegible] | OUT | 0.3155334 | 131 |
| [illegible] | [illegible] | OUT | 0.30349014 | 126 |

| Allele | % | percentage | | of indels |
|---|---|---|---|---|
| WT | 49.4% | 49.42 | | |
| R218C | 23.4% | 23.39 | | |
| Edited | 19.1% | 19.06 | | |
| Out of Frame | 17.7% | 17.68 | Out of Frame | 92.77% |
| In Frame | 1.4% | 1.38 | In Frame | 7.79% |

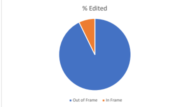


| Seq Batch | Sample # | Sample Name |
|---|---|---|
| 180718_G2GC6 | Sample 4 | 6VM2 |

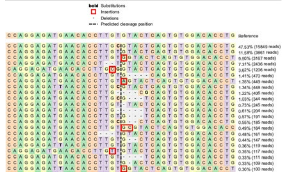


| Aligned_Sequence | Reference_Sequence | Allele | %Reads | #Reads |
|---|---|---|---|---|
| [illegible] | [illegible] | WT | 47.534641 | 15849 |
| [illegible] | [illegible] | R218C | 11.5799892 | 3861 |
| [illegible] | [illegible] | OUT | 9.49853038 | 3167 |
| [illegible] | [illegible] | OUT | 7.30610041 | 2436 |
| [illegible] | [illegible] | OUT | 3.61705956 | 1206 |
| [illegible] | [illegible] | OUT | 1.4096335 | 470 |
| [illegible] | [illegible] | OUT | 1.34664987 | 449 |
| [illegible] | [illegible] | WT | 1.34365065 | 448 |
| [illegible] | [illegible] | OUT | 1.2176834 | 406 |
| [illegible] | [illegible] | IN | 1.03173175 | 344 |
| [illegible] | [illegible] | OUT | 0.73480895 | 245 |
| [illegible] | [illegible] | OUT | 0.61184092 | 204 |
| [illegible] | [illegible] | IN | 0.57285106 | 191 |
| [illegible] | [illegible] | OUT | 0.55485574 | 185 |
| [illegible] | [illegible] | OUT | 0.49187211 | 164 |
| [illegible] | [illegible] | IN | 0.48287445 | 161 |
| [illegible] | [illegible] | OUT | 0.44088537 | 147 |
| [illegible] | [illegible] | MT | 0.3569072 | 119 |
| [illegible] | [illegible] | OUT | 0.35090876 | 117 |
| [illegible] | [illegible] | OUT | 0.33291344 | 111 |
| [illegible] | [illegible] | OUT | 0.326915 | 109 |
| [illegible] | [illegible] | OUT | 0.29992202 | 100 |

| Allele | % | percentage | | of indels |
|---|---|---|---|---|
| WT | 48.9% | 48.88 | | |
| R218C | 11.9% | 11.94 | | |
| Edited | 30.6% | 30.63 | | |
| Out of Frame | 28.5% | 28.54 | Out of Frame | 93.18% |
| In Frame | 2.1% | 2.09 | In Frame | 7.31% |

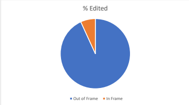


| Seq Batch | Sample # | Sample Name |
|---|---|---|
| 180718_G2GC6 | Sample 1 | 6LM1 |

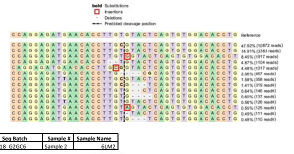


| Aligned_Sequence | Reference_Sequence | Allele | %Reads | #Reads |
|---|---|---|---|---|
| [illegible] | [illegible] | WT | 47.9196051 | 10872 |
| [illegible] | [illegible] | R218C | 14.9109661 | 3383 |
| [illegible] | [illegible] | OUT | 8.44940056 | 1917 |
| [illegible] | [illegible] | OUT | 4.86600846 | 1104 |
| [illegible] | [illegible] | OUT | 4.48254584 | 1017 |
| [illegible] | [illegible] | OUT | 2.05835684 | 467 |
| [illegible] | [illegible] | WT | 1.57792666 | 358 |
| [illegible] | [illegible] | IN | 1.40602962 | 319 |
| [illegible] | [illegible] | OUT | 0.64351199 | 146 |
| [illegible] | [illegible] | OUT | 0.60384344 | 137 |
| [illegible] | [illegible] | R218C | 0.55535966 | 126 |
| [illegible] | [illegible] | OUT | 0.55095205 | 125 |
| [illegible] | [illegible] | OUT | 0.48924542 | 111 |
| [illegible] | [illegible] | OUT | 0.4848378 | 110 |

| Allele | % | percentage | | of indels |
|---|---|---|---|---|
| WT | 49.5% | 49.50 | | |
| R218C | 15.5% | 15.47 | | |
| Edited | 24.0% | 24.03 | | |
| Out of Frame | 22.1% | 22.14 | Out of Frame | 92.13% |
| In Frame | 1.9% | 1.89 | In Frame | 8.54% |

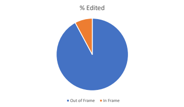


| Seq Batch | Sample # | Sample Name |
|---|---|---|
| 180718_G2GC6 | Sample 2 | 6LM2 |

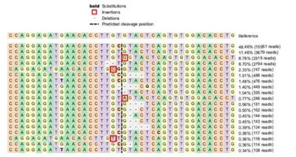


| Aligned_Sequence | Reference_Sequence | Allele | %Reads | #Reads |
|---|---|---|---|---|
| [illegible] | [illegible] | WT | 48.4600293 | 15561 |
| [illegible] | [illegible] | R218C | 11.4571331 | 3679 |
| [illegible] | [illegible] | out | 8.77892311 | 2819 |
| [illegible] | [illegible] | out | 8.70106817 | 2794 |
| [illegible] | [illegible] | out | 2.32630563 | 747 |
| [illegible] | [illegible] | in | 1.51350005 | 486 |
| [illegible] | [illegible] | WT | 1.48235807 | 476 |
| [illegible] | [illegible] | out | 1.39827473 | 449 |
| [illegible] | [illegible] | out | 1.04325621 | 335 |
| [illegible] | [illegible] | out | 0.76609262 | 246 |
| [illegible] | [illegible] | in | 0.56366977 | 181 |
| [illegible] | [illegible] | out | 0.50450002 | 162 |
| [illegible] | [illegible] | out | 0.44844446 | 144 |
| [illegible] | [illegible] | out | 0.44533026 | 143 |
| [illegible] | [illegible] | out | 0.38616051 | 124 |
| [illegible] | [illegible] | wt | 0.36436112 | 117 |
| [illegible] | [illegible] | out | 0.36124692 | 116 |
| [illegible] | [illegible] | out | 0.35501853 | 114 |
| [illegible] | [illegible] | MT | 0.33633334 | 108 |

| Allele | % | percentage | | of indels |
|---|---|---|---|---|
| WT | 50.3% | 50.31 | | |
| R218C | 11.8% | 11.79 | | |
| Edited | 27.6% | 27.59 | | |
| Out of Frame | 27.0% | 27.03 | Out of Frame | 97.96% |
| In Frame | 0.6% | 0.56 | In Frame | 2.09% |

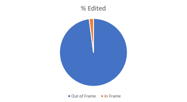


A146K sgRNA in A146K RPE

| Seq Batch | Sample # | Sample Name |
|---|---|---|
| 2VMTA-1-44044 | Sample 1 | 2VMTA1 |

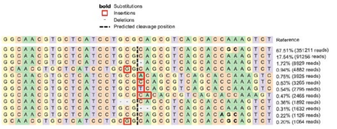


| Aligned_Sequence | Reference_Sequence | Allele | %Reads | #Reads |
|---|---|---|---|---|
| [illegible] | [illegible] | WT | 67.5124035 | 351211 |
| [illegible] | [illegible] | A146K | 17.5419104 | 91256 |
| [illegible] | [illegible] | OUT | 1.71639912 | 8929 |
| [illegible] | [illegible] | OUT | 0.93845453 | 4882 |
| [illegible] | [illegible] | OUT | 0.75449284 | 3925 |
| [illegible] | [illegible] | OUT | 0.6276227 | 3265 |
| [illegible] | [illegible] | OUT | 0.53727579 | 2795 |
| [illegible] | [illegible] | OUT | 0.47384072 | 2465 |
| [illegible] | [illegible] | OUT | 0.36369438 | 1892 |
| [illegible] | [illegible] | OUT | 0.31371524 | 1632 |
| [illegible] | [illegible] | IN | 0.21644814 | 1126 |
| [illegible] | [illegible] | OUT | 0.20453003 | 1064 |

| Allele | % | percentage | | of indels |
|---|---|---|---|---|
| WT | 67.5% | 67.51 | | |
| A146K | 17.5% | 17.54 | | |
| Edited | 6.1% | 6.15 | | |
| Out of Frame | 5.9% | 5.93 | Out of Frame | 96.48% |
| In Frame | 0.2% | 0.22 | In Frame | 3.65% |

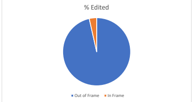


2VMTA-2-44044

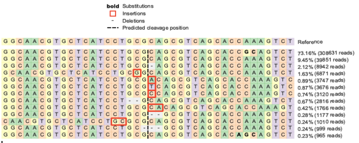


| Aligned_Sequence | Reference_Sequence | Allele | %Reads | #Reads |
|---|---|---|---|---|
| [illegible] | [illegible] | WT | 73.1611398 | 308631 |
| [illegible] | [illegible] | A146K | 9.44670038 | 39851 |
| [illegible] | [illegible] | OUT | 2.11970577 | 8942 |
| [illegible] | [illegible] | OUT | 1.62877414 | 6871 |
| [illegible] | [illegible] | OUT | 0.88822831 | 3747 |
| [illegible] | [illegible] | OUT | 0.87139772 | 3676 |
| [illegible] | [illegible] | OUT | 0.73959763 | 3120 |
| [illegible] | [illegible] | OUT | 0.66753427 | 2816 |
| [illegible] | [illegible] | OUT | 0.41863122 | 1766 |
| [illegible] | [illegible] | OUT | 0.27900847 | 1177 |
| [illegible] | [illegible] | OUT | 0.23942103 | 1010 |
| [illegible] | [illegible] | IN | 0.23681347 | 999 |
| [illegible] | [illegible] | IN | 0.22875375 | 965 |

| Allele | % | percentage | | of indels |
|---|---|---|---|---|
| WT | 73.2% | 73.16 | | |
| A146K | 9.4% | 9.45 | | |
| Edited | 8.3% | 8.32 | | |
| Out of Frame | 7.9% | 7.85 | Out of Frame | 94.40% |
| In Frame | 0.5% | 0.47 | In Frame | 5.93% |

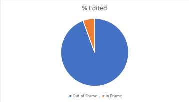


A146K MTB sgRNA in A146K RPE

2VMTB-1-44044

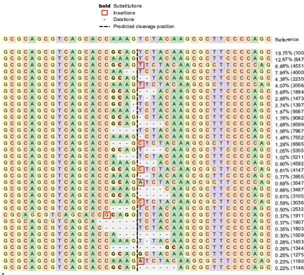


| Aligned_Sequence | Reference_Sequence | Allele | %Reads | #Reads |
|---|---|---|---|---|
| [illegible] | [illegible] | WT | 19.747621 | 100812 |
| [illegible] | [illegible] | A146K | 12.6747789 | 64705 |
| [illegible] | [illegible] | OUT | 8.87636092 | 45314 |
| [illegible] | [illegible] | OUT | 7.83640417 | 40005 |
| [illegible] | [illegible] | OUT | 4.37980654 | 22359 |
| [illegible] | [illegible] | OUT | 4.02819186 | 20564 |
| [illegible] | [illegible] | IN | 3.69146448 | 18845 |
| [illegible] | [illegible] | OUT | 2.88637459 | 14735 |
| [illegible] | [illegible] | OUT | 2.73750152 | 13975 |
| [illegible] | [illegible] | OUT | 1.87403771 | 9567 |
| [illegible] | [illegible] | OUT | 1.77903319 | 9082 |
| [illegible] | [illegible] | OUT | 1.68442043 | 8599 |
| [illegible] | [illegible] | OUT | 1.56062072 | 7967 |
| [illegible] | [illegible] | OUT | 1.47932819 | 7552 |
| [illegible] | [illegible] | OUT | 1.28598909 | 6565 |
| [illegible] | [illegible] | IN | 1.04896749 | 5355 |
| [illegible] | [illegible] | IN | 1.02075996 | 5211 |
| [illegible] | [illegible] | IN | 0.89950676 | 4592 |
| [illegible] | [illegible] | OUT | 0.81233766 | 4147 |
| [illegible] | [illegible] | OUT | 0.77472762 | 3955 |
| [illegible] | [illegible] | OUT | 0.69480629 | 3547 |
| [illegible] | [illegible] | OUT | 0.68305315 | 3487 |
| [illegible] | [illegible] | OUT | 0.60822485 | 3105 |
| [illegible] | [illegible] | OUT | 0.59470874 | 3036 |
| [illegible] | [illegible] | IN | 0.49598239 | 2532 |
| [illegible] | [illegible] | OUT | 0.37433742 | 1911 |
| [illegible] | [illegible] | IN | 0.37355387 | 1907 |
| [illegible] | [illegible] | OUT | 0.35318177 | 1803 |
| [illegible] | [illegible] | OUT | 0.2955914 | 1509 |
| [illegible] | [illegible] | OUT | 0.2846218 | 1453 |
| [illegible] | [illegible] | OUT | 0.26327027 | 1344 |
| [illegible] | [illegible] | IN | 0.24975416 | 1275 |
| [illegible] | [illegible] | OUT | 0.23114503 | 1180 |
| [illegible] | [illegible] | IN | 0.22448492 | 1146 |

| Allele | % | percentage | | of indels |
|---|---|---|---|---|
| WT | 19.7% | 19.75 | | |
| A146K | 12.7% | 12.67 | | |
| Edited | 54.4% | 54.38 | | |
| Out of Frame | 46.4% | 46.38 | Out of Frame | 85.28% |
| In Frame | 8.0% | 8.00 | In Frame | 17.26% |

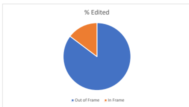


2VMTB-2-44044

bold Substitutions

Insertions

| Aligned_Sequence | Reference_Sequence | Allele | %Reads | #Reads |
|---|---|---|---|---|
| [illegible] | [illegible] | WT | 17.6273825 | 89007 |
| [illegible] | [illegible] | A146K | 9.32989527 | 47110 |

| Allele | % | percentage |
|---|---|---|
| WT | 17.6% | 17.63 |
| A146K | 9.3% | 9.33 |

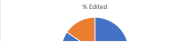

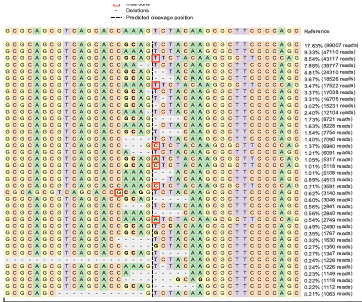

| | | | | |
|---|---|---|---|---|
| GCGCAGCGTCAGCACCGCAGTTCTACAAGCGCTTCCCCAG | GCGCAGCGTCAGCACCAAAG-TCTACAAGCGCTTCCCCAG | OUT | 8.53910199 | 43117 |
| GCGCAGCGTCAGCACCAA--TCTACAAGCGCTTCCCCAGC | GCGCAGCGTCAGCACCAAAGTCTACAAGCGCTTCCCCAGC | OUT | 7.87763202 | 39777 |
| GCGCAGCGTCAGCACCGCAG--TACAAGCGCTTCCCCAGC | GCGCAGCGTCAGCACCAAAGTCTACAAGCGCTTCCCCAGC | OUT | 4.81447154 | 24310 |
| GCGCAGCGTCAGCACCGCAG---ACAAGCGCTTCCCCAGC | GCGCAGCGTCAGCACCAAAGTCTACAAGCGCTTCCCCAGC | IN | 3.66897983 | 18526 |
| GCGCAGCGTCAGCACCAAAGTTCTACAAGCGCTTCCCCAG | GCGCAGCGTCAGCACCAAAG-TCTACAAGCGCTTCCCCAG | OUT | 3.47014275 | 17522 |
| GCGCAGCGTCAGCACCGCA-TCTACAAGCGCTTCCCCAGC | GCGCAGCGTCAGCACCAAAGTCTACAAGCGCTTCCCCAGC | OUT | 3.37428902 | 17038 |
| GCGCAGCGTCAGCACCGCAGT-TACAAGCGCTTCCCCAGC | GCGCAGCGTCAGCACCAAAGTCTACAAGCGCTTCCCCAGC | OUT | 3.30834007 | 16705 |
| GCGCAGCGTCAGCACCGCAG-CTACAAGCGCTTCCCCAGC | GCGCAGCGTCAGCACCAAAGTCTACAAGCGCTTCCCCAGC | OUT | 3.01642188 | 15231 |
| GCGCAGCGTCAGCACCAAA-TCTACAAGCGCTTCCCCAGC | GCGCAGCGTCAGCACCAAAGTCTACAAGCGCTTCCCCAGC | OUT | 2.39713548 | 12104 |
| GCGCAGCGTCAGCACCGCAG----CAAGCGCTTCCCCAGC | GCGCAGCGTCAGCACCAAAGTCTACAAGCGCTTCCCCAGC | OUT | 1.72714958 | 8721 |
| GCGCAGCGTCAGCACCAAAG-CTACAAGCGCTTCCCCAGC | GCGCAGCGTCAGCACCAAAGTCTACAAGCGCTTCCCCAGC | OUT | 1.62951344 | 8228 |
| GCGCAGCGTCAGCACCGCAGT---CAAGCGCTTCCCCAGC | GCGCAGCGTCAGCACCAAAGTCTACAAGCGCTTCCCCAGC | IN | 1.53564016 | 7754 |
| GCGCAGCGTCAGCACC----TCTACAAGCGCTTCCCCAGC | GCGCAGCGTCAGCACCAAAGTCTACAAGCGCTTCCCCAGC | OUT | 1.40413835 | 7090 |
| GCGCAGCGTCAGCACC---GCTCTACAAGCGCTTCCCCAG | GCGCAGCGTCAGCACCAAAG-TCTACAAGCGCTTCCCCAG | OUT | 1.37443161 | 6940 |
| GCGCAGCGTCAGCACCA---TCTACAAGCGCTTCCCCAGC | GCGCAGCGTCAGCACCAAAGTCTACAAGCGCTTCCCCAGC | IN | 1.20629149 | 6091 |
| GCGCAGCGTCAGCACCGCAGATCTACAAGCGCTTCCCCAG | GCGCAGCGTCAGCACCAAAG-TCTACAAGCGCTTCCCCAG | OUT | 1.05300474 | 5317 |
| GCGCAGCGTCAGCACCGCAGCTCTACAAGCGCTTCCCCAG | GCGCAGCGTCAGCACCAAAG-TCTACAAGCGCTTCCCCAG | OUT | 1.0135938 | 5118 |
| GCGCAGCGTCAGCACCAAAG---ACAAGCGCTTCCCCAGC | GCGCAGCGTCAGCACCAAAGTCTACAAGCGCTTCCCCAGC | IN | 1.01161335 | 5108 |
| GCGCAGCGTCAGCACCAAAG--TACAAGCGCTTCCCCAGC | GCGCAGCGTCAGCACCAAAGTCTACAAGCGCTTCCCCAGC | OUT | 0.89377664 | 4513 |
| GCGCAGCGTCAGCACCAAAGCTCTACAAGCGCTTCCCCAG | GCGCAGCGTCAGCACCAAAG-TCTACAAGCGCTTCCCCAG | OUT | 0.71117924 | 3591 |
| CGCAGCGTCAGCACCGCAGGTCTACAAGCGCTTCCCCAGC | CGCAGCGTCAGCACC-AAAGTCTACAAGCGCTTCCCCAGC | OUT | 0.62186099 | 3140 |
| GCGCAGCGTCAGCACCGCAG-----AAGCGCTTCCCCAGC | GCGCAGCGTCAGCACCAAAGTCTACAAGCGCTTCCCCAGC | OUT | 0.60324477 | 3046 |
| GCGCAGCGTCAGCACC---GTCTACAAGCGCTTCCCCAGC | GCGCAGCGTCAGCACCAAAGTCTACAAGCGCTTCCCCAGC | IN | 0.56264556 | 2841 |
| GCGCAGCGTCAGCACCAAAG----CAAGCGCTTCCCCAGC | GCGCAGCGTCAGCACCAAAGTCTACAAGCGCTTCCCCAGC | OUT | 0.56244752 | 2840 |
| GCGCAGCGTCAGCACCAAAGATCTACAAGCGCTTCCCCAG | GCGCAGCGTCAGCACCAAAG-TCTACAAGCGCTTCCCCAG | OUT | 0.54442543 | 2749 |
| GCGCAGCGTCAGCACCGCAGTCCACAAGCGCTTCCCCAGC | GCGCAGCGTCAGCACCAAAGTCTACAAGCGCTTCCCCAGC | IN | 0.4931318 | 2490 |
| GCGCAGCGTCAGCACCGCAGGCTACAAGCGCTTCCCCAGC | GCGCAGCGTCAGCACCAAAGTCTACAAGCGCTTCCCCAGC | IN | 0.34994534 | 1767 |
| GCGCAGCGTCAGCAC-----TCTACAAGCGCTTCCCCAGC | GCGCAGCGTCAGCACCAAAGTCTACAAGCGCTTCCCCAGC | OUT | 0.32281319 | 1630 |
| GCGCAGCGTCAGCACC----GCTACAAGCGCTTCCCCAGC | GCGCAGCGTCAGCACCAAAGTCTACAAGCGCTTCCCCAGC | OUT | 0.26736062 | 1350 |
| GCGCAGCGTCAGCACCGCAGT----AAGCGCTTCCCCAGC | GCGCAGCGTCAGCACCAAAGTCTACAAGCGCTTCCCCAGC | OUT | 0.26676648 | 1347 |
| --------------------TCTACAAGCGCTTCCCCAGC | GCGCAGCGTCAGCACCAAAGTCTACAAGCGCTTCCCCAGC | OUT | 0.24319914 | 1228 |
| GCGCAGCGTCAGCACCAAAGT-TACAAGCGCTTCCCCAGC | GCGCAGCGTCAGCACCAAAGTCTACAAGCGCTTCCCCAGC | OUT | 0.24280305 | 1226 |
| GCGCAGCGTCAGCACCA--------AAGCGCTTCCCCAGC | GCGCAGCGTCAGCACCAAAGTCTACAAGCGCTTCCCCAGC | OUT | 0.22755359 | 1149 |
| GCGCAGCGTCAGCACC-------GCAGGCGCTTCCCCAGC | GCGCAGCGTCAGCACCAAAGTCTACAAGCGCTTCCCCAGC | IN | 0.2214142 | 1118 |
| GCGCAGCGTCAGCACCGCAG------AGCGCTTCCCCAGC | GCGCAGCGTCAGCACCAAAGTCTACAAGCGCTTCCCCAGC | IN | 0.22022593 | 1112 |

| | | | | |
|---|---|---|---|---|
| **Edited** | 59.8% | 59.78 | | **of indels** |
| Out of Frame | 50.5% | 50.51 | Out of Frame | 84.49% |
| In Frame | 9.3% | 9.27 | In Frame | 18.35% |

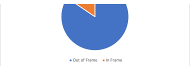


AAVS1 sgRNA in A146K RPE

2vAAVS-1-44044

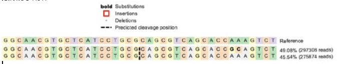

| Aligned_Sequence | Reference_Sequence | Allele | %Reads | #Reads |
|---|---|---|---|---|
| GGCAACGTGCTCATCCTGCGCAGCGTCAGCACCGCAGTCT | GGCAACGTGCTCATCCTGCGCAGCGTCAGCACCAAAGTCT | WT | 49.0784624 | 297308 |
| GGCAACGTGCTCATCCTGCGCAGCGTCAGCACCAAAGTCT | GGCAACGTGCTCATCCTGCGCAGCGTCAGCACCAAAGTCT | A146K | 45.54022 | 275874 |

| Allele | % | percentage | | |
|---|---|---|---|---|
| **WT** | 49.1% | 49.08 | | |
| **A146K** | 45.5% | 45.54 | | |
| **Edited** | 0.0% | 0.00 | | **of indels** |
| Out of Frame | 0.0% | 0.00 | Out of Frame | NA |
| In Frame | 0.0% | 0.00 | In Frame | NA |

2vAAVS-2-44044

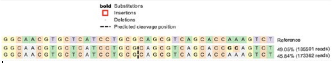

| Aligned_Sequence | Reference_Sequence | Allele | %Reads | #Reads |
|---|---|---|---|---|
| GGCAACGTGCTCATCCTGCGCAGCGTCAGCACCGCAGTCT | GGCAACGTGCTCATCCTGCGCAGCGTCAGCACCAAAGTCT | WT | 49.054613 | 185501 |
| GGCAACGTGCTCATCCTGCGCAGCGTCAGCACCAAAGTCT | GGCAACGTGCTCATCCTGCGCAGCGTCAGCACCAAAGTCT | A146K | 45.8445281 | 173362 |

| Allele | % | percentage | | |
|---|---|---|---|---|
| **WT** | 49.1% | 49.05 | | |
| **A146K** | 45.8% | 45.84 | | |
| **Edited** | 0.0% | 0.00 | | **of indels** |
| Out of Frame | 0.0% | 0.00 | Out of Frame | NA |
| In Frame | 0.0% | 0.00 | In Frame | NA |

18VMTA-1-L-44044

N296H sgRNA in N296H RPE

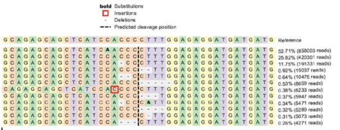

| Aligned_Sequence | Reference_Sequence | Allele | %Reads | #Reads |
|---|---|---|---|---|
| GCAGAGCAGCTCATCAACCCCTTTGGAGAGGATGATGATG | GCAGAGCAGCTCATCCACCCCTTTGGAGAGGATGATGATG | WT | 52.7051855 | 858003 |
| GCAGAGCAGCTCATCCACCCCTTTGGAGAGGATGATGATG | GCAGAGCAGCTCATCCACCCCTTTGGAGAGGATGATGATG | N296H | 25.8181407 | 420301 |
| GCAGAGCAGCTCATCCA-CCCTTTGGAGAGGATGATGATG | GCAGAGCAGCTCATCCACCCCTTTGGAGAGGATGATGATG | OUT | 11.753031 | 191331 |
| GCAGAGCAGCTCATCCACCC--TTGGAGAGGATGATGATG | GCAGAGCAGCTCATCCACCCCTTTGGAGAGGATGATGATG | OUT | 0.92368893 | 15037 |
| GCAGAGCAGCTCATCCA--CCTTTGGAGAGGATGATGATG | GCAGAGCAGCTCATCCACCCCTTTGGAGAGGATGATGATG | OUT | 0.64345558 | 10475 |
| GCAGAGCAGCTCATCCACCC-----GAGAGGATGATGATG | GCAGAGCAGCTCATCCACCCCTTTGGAGAGGATGATGATG | OUT | 0.5319028 | 8659 |
| CAGAGCAGCTCATCCACCCCCTTTGGAGAGGATGATGATG | CAGAGCAGCTCATCCA-CCCCTTTGGAGAGGATGATGATG | OUT | 0.38287911 | 6233 |
| GCAGAGCAGCTCATCCACCC---TGGAGAGGATGATGATG | GCAGAGCAGCTCATCCACCCCTTTGGAGAGGATGATGATG | IN | 0.36531077 | 5947 |
| GCAGAGCAGCTCATCCA-CCCATTGGAGAGGATGATGATG | GCAGAGCAGCTCATCCACCCCTTTGGAGAGGATGATGATG | OUT | 0.33607117 | 5471 |
| GCAGAGCAGCTCATCCACCC----GGAGAGGATGATGATG | GCAGAGCAGCTCATCCACCCCTTTGGAGAGGATGATGATG | OUT | 0.32489132 | 5289 |
| GCAGAGCAGCTCATCCACCCC-TTGGAGAGGATGATGATG | GCAGAGCAGCTCATCCACCCCTTTGGAGAGGATGATGATG | OUT | 0.31162293 | 5073 |
| GCAGAGCAGCTCATCCA---CTTTGGAGAGGATGATGATG | GCAGAGCAGCTCATCCACCCCTTTGGAGAGGATGATGATG | IN | 0.26235788 | 4271 |

| Allele | % | percentage | | |
|---|---|---|---|---|
| **WT** | 52.7% | 52.71 | | |
| **N296H** | 25.8% | 25.82 | | |
| **Edited** | 15.8% | 15.84 | | **of indels** |
| Out of Frame | 15.2% | 15.21 | Out of Frame | 96.04% |
| In Frame | 0.6% | 0.63 | In Frame | 4.13% |

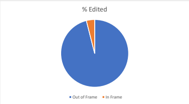


18VMTA-2-L-44044

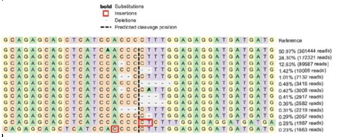

| Aligned_Sequence | Reference_Sequence | Allele | %Reads | #Reads |
|---|---|---|---|---|
| GCAGAGCAGCTCATCAACCCCTTTGGAGAGGATGATGATG | GCAGAGCAGCTCATCCACCCCTTTGGAGAGGATGATGATG | WT | 50.9685525 | 361444 |
| GCAGAGCAGCTCATCCACCCCTTTGGAGAGGATGATGATG | GCAGAGCAGCTCATCCACCCCTTTGGAGAGGATGATGATG | N296H | 24.2996203 | 172321 |
| GCAGAGCAGCTCATCCA-CCCTTTGGAGAGGATGATGATG | GCAGAGCAGCTCATCCACCCCTTTGGAGAGGATGATGATG | OUT | 12.6329935 | 89587 |
| GCAGAGCAGCTCATCCACCC--TTGGAGAGGATGATGATG | GCAGAGCAGCTCATCCACCCCTTTGGAGAGGATGATGATG | OUT | 1.41986685 | 10069 |
| GCAGAGCAGCTCATCCA--CCTTTGGAGAGGATGATGATG | GCAGAGCAGCTCATCCACCCCTTTGGAGAGGATGATGATG | OUT | 1.00570964 | 7132 |
| GCAGAGCAGCTCATCCACCC-----GAGAGGATGATGATG | GCAGAGCAGCTCATCCACCCCTTTGGAGAGGATGATGATG | OUT | 0.48170277 | 3416 |
| GCAGAGCAGCTCATCCA-CCCATTGGAGAGGATGATGATG | GCAGAGCAGCTCATCCACCCCTTTGGAGAGGATGATGATG | OUT | 0.42416918 | 3008 |
| GCAGAGCAGCTCATCCACCC---TGGAGAGGATGATGATG | GCAGAGCAGCTCATCCACCCCTTTGGAGAGGATGATGATG | IN | 0.41133694 | 2917 |
| GCAGAGCAGCTCATCCACCC----GGAGAGGATGATGATG | GCAGAGCAGCTCATCCACCCCTTTGGAGAGGATGATGATG | OUT | 0.36409735 | 2582 |
| GCAGAGCAGCTCATCCA---CTTTGGAGAGGATGATGATG | GCAGAGCAGCTCATCCACCCCTTTGGAGAGGATGATGATG | IN | 0.31290938 | 2219 |
| GCAGAGCAGCTCATCCACCCC-TTGGAGAGGATGATGATG | GCAGAGCAGCTCATCCACCCCTTTGGAGAGGATGATGATG | OUT | 0.29006516 | 2057 |
| GCAGAGCAGCTCATCCACCCCTCTTTGGAGAGGATGATGA | GCAGAGCAGCTCATCCACCC--CTTTGGAGAGGATGATGA | OUT | 0.2801942 | 1987 |
| CAGAGCAGCTCATCCACCCCCTTTGGAGAGGATGATGATG | CAGAGCAGCTCATCCA-CCCCTTTGGAGAGGATGATGATG | OUT | 0.23450577 | 1663 |

| Allele | % | percentage | | |
|---|---|---|---|---|
| **WT** | 51.0% | 50.97 | | |
| **N296H** | 24.3% | 24.30 | | |
| **Edited** | 17.9% | 17.86 | | **of indels** |
| Out of Frame | 17.1% | 17.13 | Out of Frame | 95.94% |
| In Frame | 0.7% | 0.72 | In Frame | 4.23% |

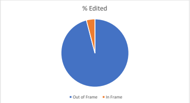


18vAAVS-1-L-44044

AAVS1 sgRNA in N296H RPE

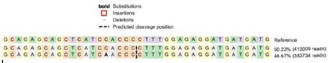

| Aligned_Sequence | Reference_Sequence | Allele | %Reads | #Reads |
|---|---|---|---|---|
| GCAGAGCAGCTCATCCACCCCTTTGGAGAGGATGATGATG | GCAGAGCAGCTCATCCACCCCTTTGGAGAGGATGATGATG | N296H | 50.2290663 | 413009 |
| GCAGAGCAGCTCATCAACCCCTTTGGAGAGGATGATGATG | GCAGAGCAGCTCATCCACCCCTTTGGAGAGGATGATGATG | WT | 46.6687179 | 383734 |

| Allele | % | percentage | | |
|---|---|---|---|---|
| **WT** | 46.7% | 46.67 | | |
| **N296H** | 50.2% | 50.23 | | |
| **Edited** | 0.0% | 0.00 | | **of indels** |
| Out of Frame | 0.0% | 0.00 | Out of Frame | NA |
| In Frame | 0.0% | 0.00 | In Frame | NA |

18vAAVS-2-L-44044

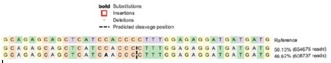

| Aligned_Sequence | Reference_Sequence | Allele | %Reads | #Reads |
|---|---|---|---|---|
| GCAGAGCAGCTCATCCACCCCTTTGGAGAGGATGATGATG | GCAGAGCAGCTCATCCACCCCTTTGGAGAGGATGATGATG | N296H | 50.1330909 | 654675 |
| GCAGAGCAGCTCATCAACCCCTTTGGAGAGGATGATGATG | GCAGAGCAGCTCATCCACCCCTTTGGAGAGGATGATGATG | WT | 46.6152937 | 608737 |

| Allele | % | percentage | | |
|---|---|---|---|---|
| **WT** | 46.6% | 46.62 | | |
| **A146K** | 50.1% | 50.13 | | |
| **Edited** | 0.0% | 0.00 | | **of indels** |
| Out of Frame | 0.0% | 0.00 | Out of Frame | NA |
| In Frame | 0.0% | 0.00 | In Frame | NA |

Averaged in and out of frame editing pie charts

| **R218C** | rep 1 | rep 2 | rep 3 | rep 4 | rep 5 | avg |
|---|---|---|---|---|---|---|
| out of fram | 96.48 | 92.77 | 93.18 | 92.13 | 97.96 | 94.504 |
| In frame | 3.52 | 7.23 | 6.82 | 7.87 | 2.04 | 5.496 |

| **R218C** | |
|---|---|
| avg out of frame | 94.5% |
| avg In frame | 5.50% |

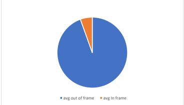


| **A146K** | rep 1 | rep 2 | avg |
|---|---|---|---|
| out of fram | 96.48 | 94.4 | 95.44 |
| In frame | 3.52 | 5.6 | 4.56 |

| **A146K** | |
|---|---|
| avg out of frame | 95.4% |
| avg In frame | 4.56% |

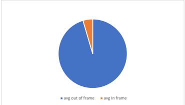


| **N296H** | rep 1 | rep 2 | avg |
|---|---|---|---|
| out of fram | 96.04 | 95.94 | 95.99 |
| In frame | 3.96 | 4.06 | 4.01 |

| **N296H** | |
|---|---|
| avg out of frame | 96.0% |
| avg In frame | 4.01% |

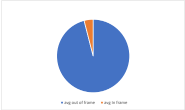

**Comparison of experimental indel frequency outcomes in iPSC-RPE+GE to outcomes predicted by inDelphi tool**

| gRNA name | gRNA sequence | Celltype | 1-bp ins frequency | Frameshift frequency | Frame +0 frequency | Frame +1 frequency | Frame +2 frequency | URL link to analysis |
|---|---|---|---|---|---|---|---|---|
| R218C sgRNA | GGTGTCCACACTGAGTACACA | mESC | 8.60384578 | 84.21743046 | 15.78256954 | 61.8101643 | 22.40726616 | /single_mESC_ILy6sOiJgBxepTEAlNgQFSbfduffLygYK_C_50 |
| R218C sgRNA | GGTGTCCACACTGAGTACACA | HEK293 | 21.5628124 | 86.45522474 | 13.54477526 | 67.22506179 | 19.23016296 | /single_HEK293_ILy6sOiJgBxepTEAlNgQFSbfduffLygYK_C_50 |
| R218C sgRNA | GGTGTCCACACTGAGTACACA | U2OS | 38.47271826 | 89.3752794 | 10.6247206 | 74.29085719 | 15.0844222 | /single_U2OS_ILy6sOiJgBxepTEAlNgQFSbfduffLygYK_C_50 |
| R218C sgRNA | GGTGTCCACACTGAGTACACA | HCT116 | 24.19476211 | 86.90971793 | 13.09028207 | 68.32482061 | 18.58489732 | /single_HCT116_ILy6sOiJgBxepTEAlNgQFSbfduffLygYK_C_50 |
| R218C sgRNA | GGTGTCCACACTGAGTACACA | K562 | 13.66236759 | 85.09095159 | 14.90904841 | 63.92386501 | 21.16708657 | /single_K562_ILy6sOiJgBxepTEAlNgQFSbfduffLygYK_C_50 |
| A146K sgRNA | ACTTTGGTGCTGACGCTGCGC | U2OS* | 19.01305833 | 82.92588141 | 17.07411859 | 75.67184442 | 7.254036991 | /single_U2OS_K3Gf7Acy5vLHv6vP74dfKR2tgbC9MPbkI_G_50 |
| N296H sgRNA | TCATCATCCTCTCCAAAGGGG | U2OS* | 19.80291099 | 78.05645198 | 21.94354802 | 48.09345543 | 29.96299654 | /single_U2OS_qIIGKdv4J3D3IoPFhndfNcHTNC5I9lT0Z_C_50 |

*Note: only predicted oucome in U2OS line data shown for A146K sgRNA and N296H sgRNA as U2OS celltype most closely correlated with % frameshift observed expirimentally for the R218C sgRNA in iPSC-RPE. Results for other lines can be viewed by following URL.

| inDelphi analysis (all celltypes) | R218C sgRNA in iPSC-RPE (this study) | |
|---|---|---|
| **Predicted (% frameshift)** | **Experimental (% frameshift)** | |
| 84.2% | 96.5% | |
| 86.5% | 92.8% | |
| 89.4% | 93.2% | |
| 86.9% | 92.1% | |
| 85.1% | 98.0% | |
| 86.4% | 94.5% | avg frameshift |
| 13.6% | 5.50% | avg non-frameshift |

| | frameshift | non-frameshift |
|---|---|---|
| Predicted | 86.4% | 13.6% |

| | frameshift | non-frameshift |
|---|---|---|
| Experimental | 94.5% | 5.50% |

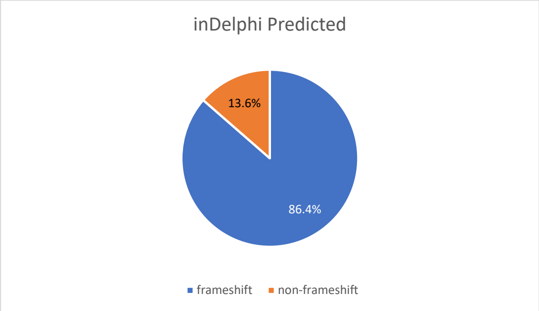


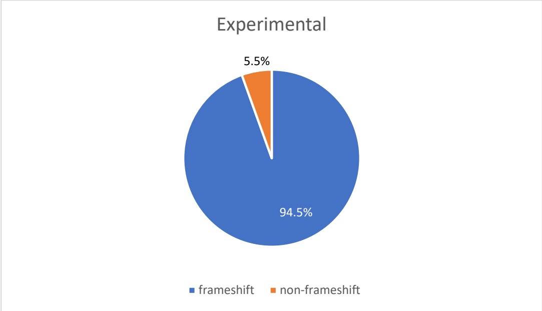

| | Phi | Precision | 1-bp ins frequency | MH del frequency | MHless del frequency | Frameshift frequency | Frame +0 frequency | Frame +1 frequency | Frame +2 frequency | Highest outcome frequency | Highest del frequency | Highest ins frequency | Expected indel length | Reference sequence | Cutsite | gRNA | gRNA orientation | Celltype | URL |
|---|---|---|---|---|---|---|---|---|---|---|---|---|---|---|---|---|---|---|---|
| 0 | 1.920438927 | 0.4794201 | 8.60384578 | 76.50094355 | 14.89521067 | 84.21743046 | 15.78256954 | 61.8101643 | 22.40726616 | 29.37686903 | 29.37686903 | 6.332992795 | 6.635332581 | CAGTGGGATACTAATCCAGTCGTAGGCATACAGGTGTCCACACTGA<br>GTACACAAGGTGTTCATCTCCTGGGGGGCAGAAGAGGGGGATGGG<br>GTTCTGAAC | 50 | GGTGTCCACACTGAGTACACA | + | mESC | /single_mESC_ILv6sOiJgBxepTEAlNgOFSbfduffLvgYK_C_50 |
| | Phi | Precision | 1-bp ins frequency | MH del frequency | MHless del frequency | Frameshift frequency | Frame +0 frequency | Frame +1 frequency | Frame +2 frequency | Highest outcome frequency | Highest del frequency | Highest ins frequency | Expected indel length | Reference sequence | Cutsite | gRNA | gRNA orientation | Celltype | URL |
| 0 | 1.920438927 | 0.4752583 | 21.5628124 | 65.65395352 | 12.78323408 | 86.45522474 | 13.54477526 | 67.22506179 | 19.23016296 | 25.21155301 | 25.21155301 | 12.96817239 | 5.836304576 | CAGTGGGATACTAATCCAGTCGTAGGCATACAGGTGTCCACACTGA<br>GTACACAAGGTGTTCATCTCCTGGGGGGCAGAAGAGGGGGATGGG<br>GTTCTGAAC | 50 | GGTGTCCACACTGAGTACACA | + | HEK293 | /single_HEK293_ILv6sOiJgBxepTEAlNgOFSbfduffLvgYK_C_50 |
| | Phi | Precision | 1-bp ins frequency | MH del frequency | MHless del frequency | Frameshift frequency | Frame +0 frequency | Frame +1 frequency | Frame +2 frequency | Highest outcome frequency | Highest del frequency | Highest ins frequency | Expected indel length | Reference sequence | Cutsite | gRNA | gRNA orientation | Celltype | URL |
| 0 | 1.920438927 | 0.5291017 | 38.47271826 | 51.49992521 | 10.02735653 | 89.3752794 | 10.6247206 | 74.29085719 | 15.0844222 | 30.18562563 | 19.77631239 | 30.18562563 | 4.793668326 | CAGTGGGATACTAATCCAGTCGTAGGCATACAGGTGTCCACACTGA<br>GTACACAAGGTGTTCATCTCCTGGGGGGCAGAAGAGGGGGATGGG<br>GTTCTGAAC | 50 | GGTGTCCACACTGAGTACACA | + | U2OS | /single_U2OS_ILv6sOiJgBxepTEAlNgOFSbfduffLvgYK_C_50 |
| | Phi | Precision | 1-bp ins frequency | MH del frequency | MHless del frequency | Frameshift frequency | Frame +0 frequency | Frame +1 frequency | Frame +2 frequency | Highest outcome frequency | Highest del frequency | Highest ins frequency | Expected indel length | Reference sequence | Cutsite | gRNA | gRNA orientation | Celltype | URL |
| 0 | 1.920438927 | 0.4768167 | 24.19476211 | 63.45094358 | 12.35429431 | 86.90971793 | 13.09028207 | 68.32482061 | 18.58489732 | 24.3655826 | 24.3655826 | 14.29884564 | 5.674023 | CAGTGGGATACTAATCCAGTCGTAGGCATACAGGTGTCCACACTGA<br>GTACACAAGGTGTTCATCTCCTGGGGGGCAGAAGAGGGGGATGGG<br>GTTCTGAAC | 50 | GGTGTCCACACTGAGTACACA | + | HCT116 | /single_HCT116_ILv6sOiJgBxepTEAlNgOFSbfduffLvgYK_C_50 |
| | Phi | Precision | 1-bp ins frequency | MH del frequency | MHless del frequency | Frameshift frequency | Frame +0 frequency | Frame +1 frequency | Frame +2 frequency | Highest outcome frequency | Highest del frequency | Highest ins frequency | Expected indel length | Reference sequence | Cutsite | gRNA | gRNA orientation | Celltype | URL |
| 0 | 1.920438927 | 0.4835021 | 13.66236759 | 72.26683004 | 14.07080237 | 85.09095159 | 14.90904841 | 63.92386501 | 21.16708657 | 27.75094139 | 27.75094139 | 11.37887222 | 6.323432666 | CAGTGGGATACTAATCCAGTCGTAGGCATACAGGTGTCCACACTGA<br>GTACACAAGGTGTTCATCTCCTGGGGGGCAGAAGAGGGGGATGGG<br>GTTCTGAAC | 50 | GGTGTCCACACTGAGTACACA | + | K562 | /single_K562_ILv6sOiJgBxepTEAlNgOFSbfduffLvgYK_C_50 |
| | Phi | Precision | 1-bp ins frequency | MH del frequency | MHless del frequency | Frameshift frequency | Frame +0 frequency | Frame +1 frequency | Frame +2 frequency | Highest outcome frequency | Highest del frequency | Highest ins frequency | Expected indel length | Reference sequence | Cutsite | gRNA | gRNA orientation | Celltype | URL |
| 0 | 2.581564899 | 0.538107 | 19.01305833 | 68.24190564 | 12.74503603 | 82.92588141 | 17.07411859 | 75.67184442 | 7.254036991 | 40.92112462 | 40.92112462 | 7.858485908 | 5.466906119 | CCAGGTGCTGGGCGCTGGGGAAGCGCTTGTAGACTTTGGTGCTGAC<br>GCTGCGCAGGATGAGCACGTTGCCCAGGTTGGCGTAGCGGATGAGC<br>GTGCGCCG | 50 | ACTTTGGTGCTGACGCTGCGC | + | U2OS | /single_U2OS_K3Gf7Acv5vLHv6vP74dfKR2tgbC9MPbkI_G_50 |
| | Phi | Precision | 1-bp ins frequency | MH del frequency | MHless del frequency | Frameshift frequency | Frame +0 frequency | Frame +1 frequency | Frame +2 frequency | Highest outcome frequency | Highest del frequency | Highest ins frequency | Expected indel length | Reference sequence | Cutsite | gRNA | gRNA orientation | Celltype | URL |
| 0 | 1.032456493 | 0.449326 | 19.80291099 | 58.00280371 | 22.19428529 | 78.05645198 | 21.94354802 | 48.09345543 | 29.96299654 | 18.908798 | 18.908798 | 13.30944276 | 4.825191232 | TCCTGTCGACAATCCAGTTGGTCTCAAAATCATCATCATCCTCTCCAA<br>AGGGGTGGATGAGCTGCTCTGCCACCTTGGGGACAGGTGAGGCTCT<br>GTAAAC | 50 | TCATCATCCTCTCCAAAGGGG | + | U2OS | /single_U2OS_aIIGKdv4J3D3IoPFhndfNcHTNC5I9lT0Z_C_50 |
