## Supplementary material for "Human iPSC modeling reveals mutation-specific responses to gene therapy in Best disease": SI Data File B

SI Data File B: Curated gene sets used to assess differences in gene expression between control (*AAVS1* ) and mutant *BEST1* allel-targeted sgRNA

| List | Name |
|---|---|
| Strunnikova_2010_RPE_MARKERS | RPE65 |
| Strunnikova_2010_RPE_MARKERS | TTR |
| Strunnikova_2010_RPE_MARKERS | CRX |
| Strunnikova_2010_RPE_MARKERS | DCT |
| Strunnikova_2010_RPE_MARKERS | BEST1 |
| Strunnikova_2010_RPE_MARKERS | SIX3 |
| Strunnikova_2010_RPE_MARKERS | CHRNA3 |
| Strunnikova_2010_RPE_MARKERS | TRPM1 |
| Strunnikova_2010_RPE_MARKERS | LHX2 |
| Strunnikova_2010_RPE_MARKERS | SFRP5 |
| Strunnikova_2010_RPE_MARKERS | SILV |
| Strunnikova_2010_RPE_MARKERS | CSPG5 |
| Strunnikova_2010_RPE_MARKERS | APLP1 |
| Strunnikova_2010_RPE_MARKERS | RBP1 |
| Strunnikova_2010_RPE_MARKERS | TYRP1 |
| Strunnikova_2010_RPE_MARKERS | MYRIP |
| Strunnikova_2010_RPE_MARKERS | TFPI2 |
| Strunnikova_2010_RPE_MARKERS | MAB21L1 |
| Strunnikova_2010_RPE_MARKERS | PTGDS |
| Strunnikova_2010_RPE_MARKERS | FRZB |
| Strunnikova_2010_RPE_MARKERS | SLC6A15 |
| Strunnikova_2010_RPE_MARKERS | SERPINF1 |
| Strunnikova_2010_RPE_MARKERS | DUSP4 |
| Strunnikova_2010_RPE_MARKERS | GEM |
| Strunnikova_2010_RPE_MARKERS | BMP4 |
| Strunnikova_2010_RPE_MARKERS | SLC6A20 |
| Strunnikova_2010_RPE_MARKERS | CDO1 |
| Strunnikova_2010_RPE_MARKERS | TTLL4 |
| Strunnikova_2010_RPE_MARKERS | CDH3 |
| Strunnikova_2010_RPE_MARKERS | ALDH1A3 |
| Strunnikova_2010_RPE_MARKERS | CLCN4 |
| Strunnikova_2010_RPE_MARKERS | ENPP2 |
| Strunnikova_2010_RPE_MARKERS | COL8A2 |
| Strunnikova_2010_RPE_MARKERS | GPR143 |
| Strunnikova_2010_RPE_MARKERS | SOSTDC1 |
| Strunnikova_2010_RPE_MARKERS | PDPN |
| Strunnikova_2010_RPE_MARKERS | GPNMB |
| Strunnikova_2010_RPE_MARKERS | SORBS2 |
| Strunnikova_2010_RPE_MARKERS | SULF1 |
| Strunnikova_2010_RPE_MARKERS | EFEMP1 |
| Strunnikova_2010_RPE_MARKERS | FOXD1 |
| Strunnikova_2010_RPE_MARKERS | MPDZ |
| Strunnikova_2010_RPE_MARKERS | SDC2 |

| | |
|---|---|
| Strunnikova_2010_RPE_MARKERS | GJA1 |
| Strunnikova_2010_RPE_MARKERS | LIMCH1 |
| Strunnikova_2010_RPE_MARKERS | FGFR2 |
| Strunnikova_2010_RPE_MARKERS | PLCB4 |
| Strunnikova_2010_RPE_MARKERS | GAS1 |
| Strunnikova_2010_RPE_MARKERS | PKNOX2 |
| Strunnikova_2010_RPE_MARKERS | PITPNA |
| Strunnikova_2010_RPE_MARKERS | ARMC9 |
| Strunnikova_2010_RPE_MARKERS | BHLHB3 |
| Strunnikova_2010_RPE_MARKERS | MFAP3L |
| Strunnikova_2010_RPE_MARKERS | ITGAV |
| Strunnikova_2010_RPE_MARKERS | LOXL1 |
| Strunnikova_2010_RPE_MARKERS | NAV3 |
| Strunnikova_2010_RPE_MARKERS | VEGFA |
| Strunnikova_2010_RPE_MARKERS | SLC16A1 |
| Strunnikova_2010_RPE_MARKERS | SLC16A1 |
| Strunnikova_2010_RPE_MARKERS | RDH11 |
| Strunnikova_2010_RPE_MARKERS | SGK3 |
| Strunnikova_2010_RPE_MARKERS | RAB38 |
| Strunnikova_2010_RPE_MARKERS | SCAMP1 |
| Strunnikova_2010_RPE_MARKERS | TIMP3 |
| Strunnikova_2010_RPE_MARKERS | LSR |
| Strunnikova_2010_RPE_MARKERS | SLC16A4 |
| Strunnikova_2010_RPE_MARKERS | MET |
| Strunnikova_2010_RPE_MARKERS | KLHL21 |
| Strunnikova_2010_RPE_MARKERS | MPHOSPH9 |
| Strunnikova_2010_RPE_MARKERS | GPM6B |
| Strunnikova_2010_RPE_MARKERS | ARL6IP1 |
| Strunnikova_2010_RPE_MARKERS | BDH2 |
| Strunnikova_2010_RPE_MARKERS | SLC4A2 |
| Strunnikova_2010_RPE_MARKERS | FADS1 |
| Strunnikova_2010_RPE_MARKERS | SMAD6 |
| Strunnikova_2010_RPE_MARKERS | RHOBTB3 |
| Strunnikova_2010_RPE_MARKERS | RRAGD |
| Strunnikova_2010_RPE_MARKERS | SLC24A1 |
| Strunnikova_2010_RPE_MARKERS | EID1 |
| Strunnikova_2010_RPE_MARKERS | EFHC1 |
| Strunnikova_2010_RPE_MARKERS | NEDD4L |
| Strunnikova_2010_RPE_MARKERS | GRAMD3 |
| Strunnikova_2010_RPE_MARKERS | ASAH1 |
| Strunnikova_2010_RPE_MARKERS | DZIP1 |
| Strunnikova_2010_RPE_MARKERS | LAPTM4B |
| Strunnikova_2010_RPE_MARKERS | NDC80 |
| Strunnikova_2010_RPE_MARKERS | COX15 |
| Strunnikova_2010_RPE_MARKERS | PHACTR2 |
| Strunnikova_2010_RPE_MARKERS | SLC39A6 |
| Strunnikova_2010_RPE_MARKERS | WWC2 |

| | |
|---|---|
| Strunnikova_2010_RPE_MARKERS | SEMA3C |
| Strunnikova_2010_RPE_MARKERS | TAX1BP1 |
| Strunnikova_2010_RPE_MARKERS | GULP1 |
| Strunnikova_2010_RPE_MARKERS | ADCY9 |
| Strunnikova_2010_RPE_MARKERS | SIL1 |
| Strunnikova_2010_RPE_MARKERS | RNF13 |
| Strunnikova_2010_RPE_MARKERS | DHPA |
| Strunnikova_2010_RPE_MARKERS | MED8 |
| Strunnikova_2010_RPE_MARKERS | DNAJB14 |
| Strunnikova_2010_RPE_MARKERS | UBL3 |
| Strunnikova_2010_RPE_MARKERS | KLHL24 |
| Strunnikova_2010_RPE_MARKERS | CRIM1 |
| Strunnikova_2010_RPE_MARKERS | CTBP2 |
| Strunnikova_2010_RPE_MARKERS | DMXL1 |
| Strunnikova_2010_RPE_MARKERS | C20orf19 |
| Strunnikova_2010_RPE_MARKERS | NRIP1 |
| Strunnikova_2010_RPE_MARKERS | USP34 |
| Strunnikova_2010_RPE_MARKERS | PRNP |
| Strunnikova_2010_RPE_MARKERS | SMC3 |
| Strunnikova_2010_RPE_MARKERS | LIN7C |
| Strunnikova_2010_RPE_MARKERS | OSTM1 |
| Strunnikova_2010_RPE_MARKERS | PCYOX1 |
| Strunnikova_2010_RPE_MARKERS | PLAG1 |
| Strunnikova_2010_RPE_MARKERS | WWTR1 |
| Strunnikova_2010_RPE_MARKERS | C1orf108 |
| Strunnikova_2010_RPE_MARKERS | PTPRG |
| Strunnikova_2010_RPE_MARKERS | DIXDC1 |
| Strunnikova_2010_RPE_MARKERS | DAP3 |
| Strunnikova_2010_RPE_MARKERS | STCH |
| Strunnikova_2010_RPE_MARKERS | FAM18B |
| Strunnikova_2010_RPE_MARKERS | ANKRD12 |
| Strunnikova_2010_RPE_MARKERS | PLOD2 |
| Strunnikova_2010_RPE_MARKERS | ADAM9 |
| Strunnikova_2010_RPE_MARKERS | MAP9 |
| Strunnikova_2010_RPE_MARKERS | LAMP2 |
| Strunnikova_2010_RPE_MARKERS | STAM2 |
| Strunnikova_2010_RPE_MARKERS | DCUN1D4 |
| Strunnikova_2010_RPE_MARKERS | PDZD8 |
| Strunnikova_2010_RPE_MARKERS | CYP20A1 |
| Strunnikova_2010_RPE_MARKERS | PSME4 |
| Strunnikova_2010_RPE_MARKERS | BAT2D1 |
| Strunnikova_2010_RPE_MARKERS | SPAST |
| Strunnikova_2010_RPE_MARKERS | CALU |
| Strunnikova_2010_RPE_MARKERS | MBNL2 |
| Strunnikova_2010_RPE_MARKERS | DEGS1 |
| Strunnikova_2010_RPE_MARKERS | IGF2BP2 |
| Strunnikova_2010_RPE_MARKERS | IFT74 |

| | |
|---|---|
| Strunnikova_2010_RPE_MARKERS | PAK1IP1 |
| Strunnikova_2010_RPE_MARKERS | MANEA |
| Strunnikova_2010_RPE_MARKERS | GOLPH3L |
| Strunnikova_2010_RPE_MARKERS | LGALS8 |
| Strunnikova_2010_RPE_MARKERS | ATF1 |
| Strunnikova_2010_RPE_MARKERS | BCLAF1 |
| Strunnikova_2010_RPE_MARKERS | HSP90B1 |
| Strunnikova_2010_RPE_MARKERS | NUDT4 |
| Strunnikova_2010_RPE_MARKERS | CDH1 |
| Strunnikova_2010_RPE_MARKERS | RBM34 |
| Strunnikova_2010_RPE_MARKERS | AHR |
| Strunnikova_2010_RPE_MARKERS | WASL |
| GO_CELL_CYCLE | AAAS |
| GO_CELL_CYCLE | ABCB1 |
| GO_CELL_CYCLE | ABL1 |
| GO_CELL_CYCLE | ACTR1A |
| GO_CELL_CYCLE | ACTR2 |
| GO_CELL_CYCLE | ACTR3 |
| GO_CELL_CYCLE | ACTR8 |
| GO_CELL_CYCLE | ACVR1 |
| GO_CELL_CYCLE | ACVR1B |
| GO_CELL_CYCLE | ADCY3 |
| GO_CELL_CYCLE | AHCTF1 |
| GO_CELL_CYCLE | AHR |
| GO_CELL_CYCLE | AK1 |
| GO_CELL_CYCLE | AKAP8 |
| GO_CELL_CYCLE | AKAP8L |
| GO_CELL_CYCLE | AKAP9 |
| GO_CELL_CYCLE | ALKBH4 |
| GO_CELL_CYCLE | ALMS1 |
| GO_CELL_CYCLE | ANAPC1 |
| GO_CELL_CYCLE | ANAPC10 |
| GO_CELL_CYCLE | ANAPC11 |
| GO_CELL_CYCLE | ANAPC13 |
| GO_CELL_CYCLE | ANAPC16 |
| GO_CELL_CYCLE | ANAPC2 |
| GO_CELL_CYCLE | ANAPC4 |
| GO_CELL_CYCLE | ANAPC5 |
| GO_CELL_CYCLE | ANAPC7 |
| GO_CELL_CYCLE | ANK3 |
| GO_CELL_CYCLE | ANKLE2 |
| GO_CELL_CYCLE | ANLN |
| GO_CELL_CYCLE | ANXA11 |
| GO_CELL_CYCLE | APBB1 |
| GO_CELL_CYCLE | APBB2 |
| GO_CELL_CYCLE | APC |
| GO_CELL_CYCLE | APEX2 |

| | |
|---|---|
| GO_CELL_CYCLE | APITD1 |
| GO_CELL_CYCLE | APPL1 |
| GO_CELL_CYCLE | APPL2 |
| GO_CELL_CYCLE | ARAP1 |
| GO_CELL_CYCLE | ARF6 |
| GO_CELL_CYCLE | ARHGEF10 |
| GO_CELL_CYCLE | ARHGEF11 |
| GO_CELL_CYCLE | ARHGEF2 |
| GO_CELL_CYCLE | ARID3A |
| GO_CELL_CYCLE | ARL2 |
| GO_CELL_CYCLE | ARL3 |
| GO_CELL_CYCLE | ARL8A |
| GO_CELL_CYCLE | ARL8B |
| GO_CELL_CYCLE | ARPP19 |
| GO_CELL_CYCLE | ASPM |
| GO_CELL_CYCLE | ASZ1 |
| GO_CELL_CYCLE | ATF2 |
| GO_CELL_CYCLE | ATM |
| GO_CELL_CYCLE | ATR |
| GO_CELL_CYCLE | ATRIP |
| GO_CELL_CYCLE | AURKA |
| GO_CELL_CYCLE | AURKB |
| GO_CELL_CYCLE | AURKC |
| GO_CELL_CYCLE | AVPI1 |
| GO_CELL_CYCLE | AZI1 |
| GO_CELL_CYCLE | AZI2 |
| GO_CELL_CYCLE | B9D2 |
| GO_CELL_CYCLE | BABAM1 |
| GO_CELL_CYCLE | BACH1 |
| GO_CELL_CYCLE | BAG6 |
| GO_CELL_CYCLE | BANF1 |
| GO_CELL_CYCLE | BANP |
| GO_CELL_CYCLE | BARD1 |
| GO_CELL_CYCLE | BAX |
| GO_CELL_CYCLE | BBS4 |
| GO_CELL_CYCLE | BCAT1 |
| GO_CELL_CYCLE | BCCIP |
| GO_CELL_CYCLE | BCL2L1 |
| GO_CELL_CYCLE | BCL2L11 |
| GO_CELL_CYCLE | BECN1 |
| GO_CELL_CYCLE | BEX2 |
| GO_CELL_CYCLE | BIN3 |
| GO_CELL_CYCLE | BIRC2 |
| GO_CELL_CYCLE | BIRC3 |
| GO_CELL_CYCLE | BIRC5 |
| GO_CELL_CYCLE | BIRC6 |
| GO_CELL_CYCLE | BIRC7 |

| | |
|---|---|
| GO_CELL_CYCLE | BIRC8 |
| GO_CELL_CYCLE | BLCAP |
| GO_CELL_CYCLE | BLM |
| GO_CELL_CYCLE | BOD1 |
| GO_CELL_CYCLE | BOD1P |
| GO_CELL_CYCLE | BOLL |
| GO_CELL_CYCLE | BORA |
| GO_CELL_CYCLE | BRCA1 |
| GO_CELL_CYCLE | BRCA2 |
| GO_CELL_CYCLE | BRCC3 |
| GO_CELL_CYCLE | BRD4 |
| GO_CELL_CYCLE | BRD7 |
| GO_CELL_CYCLE | BRDT |
| GO_CELL_CYCLE | BRE |
| GO_CELL_CYCLE | BRIP1 |
| GO_CELL_CYCLE | BRSK1 |
| GO_CELL_CYCLE | BRSK2 |
| GO_CELL_CYCLE | BTG2 |
| GO_CELL_CYCLE | BTG4 |
| GO_CELL_CYCLE | BTRC |
| GO_CELL_CYCLE | BUB1 |
| GO_CELL_CYCLE | BUB1B |
| GO_CELL_CYCLE | BUB3 |
| GO_CELL_CYCLE | C10orf46 |
| GO_CELL_CYCLE | C11orf20 |
| GO_CELL_CYCLE | C11orf51 |
| GO_CELL_CYCLE | C11orf80 |
| GO_CELL_CYCLE | C11orf82 |
| GO_CELL_CYCLE | C11orf85 |
| GO_CELL_CYCLE | C12orf11 |
| GO_CELL_CYCLE | C12orf32 |
| GO_CELL_CYCLE | C13orf15 |
| GO_CELL_CYCLE | C15orf23 |
| GO_CELL_CYCLE | C15orf42 |
| GO_CELL_CYCLE | C15orf43 |
| GO_CELL_CYCLE | C15orf60 |
| GO_CELL_CYCLE | C16orf73 |
| GO_CELL_CYCLE | C17orf87 |
| GO_CELL_CYCLE | C19orf21 |
| GO_CELL_CYCLE | C19orf46 |
| GO_CELL_CYCLE | C1orf135 |
| GO_CELL_CYCLE | C1orf96 |
| GO_CELL_CYCLE | C2CD3 |
| GO_CELL_CYCLE | C2orf29 |
| GO_CELL_CYCLE | C2orf40 |
| GO_CELL_CYCLE | C2orf65 |
| GO_CELL_CYCLE | C7orf11 |

| | |
|---|---|
| GO_CELL_CYCLE | C7orf59 |
| GO_CELL_CYCLE | C9orf114 |
| GO_CELL_CYCLE | C9orf69 |
| GO_CELL_CYCLE | CAB39 |
| GO_CELL_CYCLE | CAB39L |
| GO_CELL_CYCLE | CABLES1 |
| GO_CELL_CYCLE | CABLES2 |
| GO_CELL_CYCLE | CALM1 |
| GO_CELL_CYCLE | CALM2 |
| GO_CELL_CYCLE | CALM3 |
| GO_CELL_CYCLE | CALR |
| GO_CELL_CYCLE | CAMK1 |
| GO_CELL_CYCLE | CAMK2A |
| GO_CELL_CYCLE | CAMK2D |
| GO_CELL_CYCLE | CAMK2G |
| GO_CELL_CYCLE | CAPN3 |
| GO_CELL_CYCLE | CARM1 |
| GO_CELL_CYCLE | CASC5 |
| GO_CELL_CYCLE | CASP2 |
| GO_CELL_CYCLE | CASP8AP2 |
| GO_CELL_CYCLE | CCAR1 |
| GO_CELL_CYCLE | CCDC124 |
| GO_CELL_CYCLE | CCDC155 |
| GO_CELL_CYCLE | CCDC67 |
| GO_CELL_CYCLE | CCDC79 |
| GO_CELL_CYCLE | CCDC99 |
| GO_CELL_CYCLE | CCNA1 |
| GO_CELL_CYCLE | CCNA2 |
| GO_CELL_CYCLE | CCNB1 |
| GO_CELL_CYCLE | CCNB1IP1 |
| GO_CELL_CYCLE | CCNB2 |
| GO_CELL_CYCLE | CCNB3 |
| GO_CELL_CYCLE | CCND1 |
| GO_CELL_CYCLE | CCND2 |
| GO_CELL_CYCLE | CCND3 |
| GO_CELL_CYCLE | CCNDBP1 |
| GO_CELL_CYCLE | CCNE1 |
| GO_CELL_CYCLE | CCNE2 |
| GO_CELL_CYCLE | CCNF |
| GO_CELL_CYCLE | CCNG1 |
| GO_CELL_CYCLE | CCNG2 |
| GO_CELL_CYCLE | CCNH |
| GO_CELL_CYCLE | CCNK |
| GO_CELL_CYCLE | CCNO |
| GO_CELL_CYCLE | CCNT1 |
| GO_CELL_CYCLE | CCNT2 |
| GO_CELL_CYCLE | CCNY |

| | |
|---|---|
| GO_CELL_CYCLE | CCP110 |
| GO_CELL_CYCLE | CCPG1 |
| GO_CELL_CYCLE | CD2AP |
| GO_CELL_CYCLE | CDC123 |
| GO_CELL_CYCLE | CDC14A |
| GO_CELL_CYCLE | CDC14B |
| GO_CELL_CYCLE | CDC16 |
| GO_CELL_CYCLE | CDC20 |
| GO_CELL_CYCLE | CDC23 |
| GO_CELL_CYCLE | CDC25A |
| GO_CELL_CYCLE | CDC25B |
| GO_CELL_CYCLE | CDC25C |
| GO_CELL_CYCLE | CDC26 |
| GO_CELL_CYCLE | CDC27 |
| GO_CELL_CYCLE | CDC34 |
| GO_CELL_CYCLE | CDC45 |
| GO_CELL_CYCLE | CDC5L |
| GO_CELL_CYCLE | CDC6 |
| GO_CELL_CYCLE | CDC7 |
| GO_CELL_CYCLE | CDC73 |
| GO_CELL_CYCLE | CDCA2 |
| GO_CELL_CYCLE | CDCA3 |
| GO_CELL_CYCLE | CDCA5 |
| GO_CELL_CYCLE | CDCA8 |
| GO_CELL_CYCLE | CDH13 |
| GO_CELL_CYCLE | CDK1 |
| GO_CELL_CYCLE | CDK11A |
| GO_CELL_CYCLE | CDK11B |
| GO_CELL_CYCLE | CDK14 |
| GO_CELL_CYCLE | CDK2 |
| GO_CELL_CYCLE | CDK20 |
| GO_CELL_CYCLE | CDK2AP1 |
| GO_CELL_CYCLE | CDK3 |
| GO_CELL_CYCLE | CDK4 |
| GO_CELL_CYCLE | CDK5 |
| GO_CELL_CYCLE | CDK5RAP2 |
| GO_CELL_CYCLE | CDK5RAP3 |
| GO_CELL_CYCLE | CDK6 |
| GO_CELL_CYCLE | CDK7 |
| GO_CELL_CYCLE | CDKN1A |
| GO_CELL_CYCLE | CDKN1B |
| GO_CELL_CYCLE | CDKN1C |
| GO_CELL_CYCLE | CDKN2A |
| GO_CELL_CYCLE | CDKN2B |
| GO_CELL_CYCLE | CDKN2C |
| GO_CELL_CYCLE | CDKN2D |
| GO_CELL_CYCLE | CDKN3 |

| | |
|---|---|
| GO_CELL_CYCLE | CDT1 |
| GO_CELL_CYCLE | CECR2 |
| GO_CELL_CYCLE | CENPA |
| GO_CELL_CYCLE | CENPC1 |
| GO_CELL_CYCLE | CENPE |
| GO_CELL_CYCLE | CENPF |
| GO_CELL_CYCLE | CENPH |
| GO_CELL_CYCLE | CENPI |
| GO_CELL_CYCLE | CENPJ |
| GO_CELL_CYCLE | CENPK |
| GO_CELL_CYCLE | CENPL |
| GO_CELL_CYCLE | CENPM |
| GO_CELL_CYCLE | CENPN |
| GO_CELL_CYCLE | CENPO |
| GO_CELL_CYCLE | CENPP |
| GO_CELL_CYCLE | CENPQ |
| GO_CELL_CYCLE | CENPT |
| GO_CELL_CYCLE | CENPV |
| GO_CELL_CYCLE | CENPW |
| GO_CELL_CYCLE | CEP120 |
| GO_CELL_CYCLE | CEP135 |
| GO_CELL_CYCLE | CEP152 |
| GO_CELL_CYCLE | CEP164 |
| GO_CELL_CYCLE | CEP192 |
| GO_CELL_CYCLE | CEP250 |
| GO_CELL_CYCLE | CEP290 |
| GO_CELL_CYCLE | CEP41 |
| GO_CELL_CYCLE | CEP55 |
| GO_CELL_CYCLE | CEP57 |
| GO_CELL_CYCLE | CEP57L1 |
| GO_CELL_CYCLE | CEP63 |
| GO_CELL_CYCLE | CEP70 |
| GO_CELL_CYCLE | CEP72 |
| GO_CELL_CYCLE | CEP76 |
| GO_CELL_CYCLE | CEP78 |
| GO_CELL_CYCLE | CETN1 |
| GO_CELL_CYCLE | CETN2 |
| GO_CELL_CYCLE | CETN3 |
| GO_CELL_CYCLE | CGREF1 |
| GO_CELL_CYCLE | CGRRF1 |
| GO_CELL_CYCLE | CHAF1A |
| GO_CELL_CYCLE | CHAF1B |
| GO_CELL_CYCLE | CHAMP1 |
| GO_CELL_CYCLE | CHD3 |
| GO_CELL_CYCLE | CHD4 |
| GO_CELL_CYCLE | CHEK1 |
| GO_CELL_CYCLE | CHEK2 |

| | |
|---|---|
| GO_CELL_CYCLE | CHFR |
| GO_CELL_CYCLE | CHMP1A |
| GO_CELL_CYCLE | CHMP1B |
| GO_CELL_CYCLE | CHMP2A |
| GO_CELL_CYCLE | CHMP2B |
| GO_CELL_CYCLE | CHMP3 |
| GO_CELL_CYCLE | CHMP4A |
| GO_CELL_CYCLE | CHMP4B |
| GO_CELL_CYCLE | CHMP4C |
| GO_CELL_CYCLE | CHMP5 |
| GO_CELL_CYCLE | CHMP6 |
| GO_CELL_CYCLE | CHMP7 |
| GO_CELL_CYCLE | CHTF18 |
| GO_CELL_CYCLE | CHTF8 |
| GO_CELL_CYCLE | CIB1 |
| GO_CELL_CYCLE | CINP |
| GO_CELL_CYCLE | CIT |
| GO_CELL_CYCLE | CKAP2 |
| GO_CELL_CYCLE | CKAP5 |
| GO_CELL_CYCLE | CKS1B |
| GO_CELL_CYCLE | CKS2 |
| GO_CELL_CYCLE | CLASP1 |
| GO_CELL_CYCLE | CLASP2 |
| GO_CELL_CYCLE | CLIP1 |
| GO_CELL_CYCLE | CLOCK |
| GO_CELL_CYCLE | CLSPN |
| GO_CELL_CYCLE | CLTC |
| GO_CELL_CYCLE | CLTCL1 |
| GO_CELL_CYCLE | CNOT1 |
| GO_CELL_CYCLE | CNOT10 |
| GO_CELL_CYCLE | CNOT2 |
| GO_CELL_CYCLE | CNOT3 |
| GO_CELL_CYCLE | CNOT4 |
| GO_CELL_CYCLE | CNOT6 |
| GO_CELL_CYCLE | CNOT6L |
| GO_CELL_CYCLE | CNOT7 |
| GO_CELL_CYCLE | CNOT8 |
| GO_CELL_CYCLE | CNTD1 |
| GO_CELL_CYCLE | CNTRL |
| GO_CELL_CYCLE | CNTROB |
| GO_CELL_CYCLE | CRADD |
| GO_CELL_CYCLE | CREBL2 |
| GO_CELL_CYCLE | CRLF3 |
| GO_CELL_CYCLE | CROCC |
| GO_CELL_CYCLE | CSNK1A1 |
| GO_CELL_CYCLE | CSNK1D |
| GO_CELL_CYCLE | CSNK1E |

| | |
|---|---|
| GO_CELL_CYCLE | CSNK2A1 |
| GO_CELL_CYCLE | CSNK2A2 |
| GO_CELL_CYCLE | CSRP2BP |
| GO_CELL_CYCLE | CTCFL |
| GO_CELL_CYCLE | CTDNEP1 |
| GO_CELL_CYCLE | CTDP1 |
| GO_CELL_CYCLE | CTNNB1 |
| GO_CELL_CYCLE | CUL1 |
| GO_CELL_CYCLE | CUL2 |
| GO_CELL_CYCLE | CUL3 |
| GO_CELL_CYCLE | CUL4A |
| GO_CELL_CYCLE | CUL4B |
| GO_CELL_CYCLE | CUL5 |
| GO_CELL_CYCLE | CUL7 |
| GO_CELL_CYCLE | CUZD1 |
| GO_CELL_CYCLE | CYLD |
| GO_CELL_CYCLE | CYP26B1 |
| GO_CELL_CYCLE | CYP27B1 |
| GO_CELL_CYCLE | DAB2IP |
| GO_CELL_CYCLE | DAPK3 |
| GO_CELL_CYCLE | DAZL |
| GO_CELL_CYCLE | DBC1 |
| GO_CELL_CYCLE | DBF4 |
| GO_CELL_CYCLE | DBF4B |
| GO_CELL_CYCLE | DCLRE1A |
| GO_CELL_CYCLE | DCLRE1B |
| GO_CELL_CYCLE | DCTN1 |
| GO_CELL_CYCLE | DCTN2 |
| GO_CELL_CYCLE | DCTN3 |
| GO_CELL_CYCLE | DDIT3 |
| GO_CELL_CYCLE | DDX11 |
| GO_CELL_CYCLE | DDX12P |
| GO_CELL_CYCLE | DDX4 |
| GO_CELL_CYCLE | DGKZ |
| GO_CELL_CYCLE | DHCR24 |
| GO_CELL_CYCLE | DHFR |
| GO_CELL_CYCLE | DHFRP1 |
| GO_CELL_CYCLE | DIAPH2 |
| GO_CELL_CYCLE | DIS3L2 |
| GO_CELL_CYCLE | DIXDC1 |
| GO_CELL_CYCLE | DLG1 |
| GO_CELL_CYCLE | DLGAP5 |
| GO_CELL_CYCLE | DMC1 |
| GO_CELL_CYCLE | DMRTC2 |
| GO_CELL_CYCLE | DMTF1 |
| GO_CELL_CYCLE | DNA2 |
| GO_CELL_CYCLE | DNM2 |

| | |
|---|---|
| GO_CELL_CYCLE | DNMT3A |
| GO_CELL_CYCLE | DPEP3 |
| GO_CELL_CYCLE | DSCC1 |
| GO_CELL_CYCLE | DSN1 |
| GO_CELL_CYCLE | DST |
| GO_CELL_CYCLE | DTL |
| GO_CELL_CYCLE | DTYMK |
| GO_CELL_CYCLE | DUSP1 |
| GO_CELL_CYCLE | DUSP13 |
| GO_CELL_CYCLE | DYNC1H1 |
| GO_CELL_CYCLE | DYNC1I2 |
| GO_CELL_CYCLE | DYNC1LI1 |
| GO_CELL_CYCLE | DYNLL1 |
| GO_CELL_CYCLE | DYNLT1 |
| GO_CELL_CYCLE | DYNLT3 |
| GO_CELL_CYCLE | E2F1 |
| GO_CELL_CYCLE | E2F2 |
| GO_CELL_CYCLE | E2F3 |
| GO_CELL_CYCLE | E2F4 |
| GO_CELL_CYCLE | E2F6 |
| GO_CELL_CYCLE | E2F7 |
| GO_CELL_CYCLE | E2F8 |
| GO_CELL_CYCLE | E4F1 |
| GO_CELL_CYCLE | ECT2 |
| GO_CELL_CYCLE | EGFL6 |
| GO_CELL_CYCLE | EID1 |
| GO_CELL_CYCLE | EIF2AK4 |
| GO_CELL_CYCLE | EIF4E |
| GO_CELL_CYCLE | EIF4EBP1 |
| GO_CELL_CYCLE | EIF4G2 |
| GO_CELL_CYCLE | EMD |
| GO_CELL_CYCLE | EML1 |
| GO_CELL_CYCLE | EML4 |
| GO_CELL_CYCLE | ENSA |
| GO_CELL_CYCLE | EP300 |
| GO_CELL_CYCLE | EPS8 |
| GO_CELL_CYCLE | ERBB2IP |
| GO_CELL_CYCLE | ERCC4 |
| GO_CELL_CYCLE | ERCC6L |
| GO_CELL_CYCLE | EREG |
| GO_CELL_CYCLE | ERF |
| GO_CELL_CYCLE | ERH |
| GO_CELL_CYCLE | ERN1 |
| GO_CELL_CYCLE | ESCO1 |
| GO_CELL_CYCLE | ESCO2 |
| GO_CELL_CYCLE | ESPL1 |
| GO_CELL_CYCLE | EVI5 |

| | |
|---|---|
| GO_CELL_CYCLE | EXD1 |
| GO_CELL_CYCLE | EXO1 |
| GO_CELL_CYCLE | EYA1 |
| GO_CELL_CYCLE | EZH2 |
| GO_CELL_CYCLE | EZR |
| GO_CELL_CYCLE | FAM175A |
| GO_CELL_CYCLE | FAM175B |
| GO_CELL_CYCLE | FAM32A |
| GO_CELL_CYCLE | FAM5B |
| GO_CELL_CYCLE | FAM5C |
| GO_CELL_CYCLE | FAM64A |
| GO_CELL_CYCLE | FAM83D |
| GO_CELL_CYCLE | FANCA |
| GO_CELL_CYCLE | FANCD2 |
| GO_CELL_CYCLE | FANCG |
| GO_CELL_CYCLE | FANCI |
| GO_CELL_CYCLE | FANCM |
| GO_CELL_CYCLE | FAP |
| GO_CELL_CYCLE | FBXL15 |
| GO_CELL_CYCLE | FBXL7 |
| GO_CELL_CYCLE | FBXO31 |
| GO_CELL_CYCLE | FBXO43 |
| GO_CELL_CYCLE | FBXO5 |
| GO_CELL_CYCLE | FBXO6 |
| GO_CELL_CYCLE | FBXW11 |
| GO_CELL_CYCLE | FBXW5 |
| GO_CELL_CYCLE | FBXW7 |
| GO_CELL_CYCLE | FER |
| GO_CELL_CYCLE | FGF10 |
| GO_CELL_CYCLE | FGFR1OP |
| GO_CELL_CYCLE | FIGN |
| GO_CELL_CYCLE | FKBP6 |
| GO_CELL_CYCLE | FLNA |
| GO_CELL_CYCLE | FMN2 |
| GO_CELL_CYCLE | FOXM1 |
| GO_CELL_CYCLE | FOXN3 |
| GO_CELL_CYCLE | FOXO4 |
| GO_CELL_CYCLE | FSD1 |
| GO_CELL_CYCLE | FZR1 |
| GO_CELL_CYCLE | GADD45A |
| GO_CELL_CYCLE | GADD45GIP1 |
| GO_CELL_CYCLE | GAK |
| GO_CELL_CYCLE | GAS1 |
| GO_CELL_CYCLE | GAS2 |
| GO_CELL_CYCLE | GAS2L1 |
| GO_CELL_CYCLE | GAS2L2 |
| GO_CELL_CYCLE | GAS2L3 |

| | |
|---|---|
| GO_CELL_CYCLE | GAS6 |
| GO_CELL_CYCLE | GAS7 |
| GO_CELL_CYCLE | GEM |
| GO_CELL_CYCLE | GFI1 |
| GO_CELL_CYCLE | GIGYF2 |
| GO_CELL_CYCLE | GINS1 |
| GO_CELL_CYCLE | GINS2 |
| GO_CELL_CYCLE | GML |
| GO_CELL_CYCLE | GMNC |
| GO_CELL_CYCLE | GMNN |
| GO_CELL_CYCLE | GNAI1 |
| GO_CELL_CYCLE | GNAI2 |
| GO_CELL_CYCLE | GNAI3 |
| GO_CELL_CYCLE | GNB2L1 |
| GO_CELL_CYCLE | GOLGA2 |
| GO_CELL_CYCLE | GORASP1 |
| GO_CELL_CYCLE | GPER |
| GO_CELL_CYCLE | GPR132 |
| GO_CELL_CYCLE | GPS1 |
| GO_CELL_CYCLE | GPS2 |
| GO_CELL_CYCLE | GPSM2 |
| GO_CELL_CYCLE | GSG2 |
| GO_CELL_CYCLE | GSK3B |
| GO_CELL_CYCLE | GSPT1 |
| GO_CELL_CYCLE | GSPT2 |
| GO_CELL_CYCLE | GTPBP8 |
| GO_CELL_CYCLE | GTSE1 |
| GO_CELL_CYCLE | H1FOO |
| GO_CELL_CYCLE | H2AFX |
| GO_CELL_CYCLE | HACE1 |
| GO_CELL_CYCLE | HAUS1 |
| GO_CELL_CYCLE | HAUS2 |
| GO_CELL_CYCLE | HAUS3 |
| GO_CELL_CYCLE | HAUS4 |
| GO_CELL_CYCLE | HAUS5 |
| GO_CELL_CYCLE | HAUS6 |
| GO_CELL_CYCLE | HAUS7 |
| GO_CELL_CYCLE | HAUS8 |
| GO_CELL_CYCLE | HBP1 |
| GO_CELL_CYCLE | HBXIP |
| GO_CELL_CYCLE | HCFC1 |
| GO_CELL_CYCLE | HDAC3 |
| GO_CELL_CYCLE | HDAC8 |
| GO_CELL_CYCLE | HELLS |
| GO_CELL_CYCLE | HEPACAM |
| GO_CELL_CYCLE | HEPACAM2 |
| GO_CELL_CYCLE | HFM1 |

| | |
|---|---|
| GO_CELL_CYCLE | HGF |
| GO_CELL_CYCLE | HINFP |
| GO_CELL_CYCLE | HJURP |
| GO_CELL_CYCLE | HMG20B |
| GO_CELL_CYCLE | HMGA2 |
| GO_CELL_CYCLE | HMMR |
| GO_CELL_CYCLE | HORMAD1 |
| GO_CELL_CYCLE | HORMAD2 |
| GO_CELL_CYCLE | HRAS |
| GO_CELL_CYCLE | HRSP12 |
| GO_CELL_CYCLE | HSF1 |
| GO_CELL_CYCLE | HSP90AA1 |
| GO_CELL_CYCLE | HSPA2 |
| GO_CELL_CYCLE | HTT |
| GO_CELL_CYCLE | HUS1 |
| GO_CELL_CYCLE | HUS1B |
| GO_CELL_CYCLE | ID2 |
| GO_CELL_CYCLE | ID4 |
| GO_CELL_CYCLE | IDAS |
| GO_CELL_CYCLE | IFNG |
| GO_CELL_CYCLE | IFNW1 |
| GO_CELL_CYCLE | IKZF1 |
| GO_CELL_CYCLE | IL12A |
| GO_CELL_CYCLE | IL12B |
| GO_CELL_CYCLE | IL8 |
| GO_CELL_CYCLE | ILK |
| GO_CELL_CYCLE | INCENP |
| GO_CELL_CYCLE | ING1 |
| GO_CELL_CYCLE | ING2 |
| GO_CELL_CYCLE | ING4 |
| GO_CELL_CYCLE | INHA |
| GO_CELL_CYCLE | INHBA |
| GO_CELL_CYCLE | INO80 |
| GO_CELL_CYCLE | INSM1 |
| GO_CELL_CYCLE | INTS3 |
| GO_CELL_CYCLE | INTS7 |
| GO_CELL_CYCLE | IQGAP3 |
| GO_CELL_CYCLE | IRF1 |
| GO_CELL_CYCLE | IRF6 |
| GO_CELL_CYCLE | IST1 |
| GO_CELL_CYCLE | ITGB1 |
| GO_CELL_CYCLE | ITGB3BP |
| GO_CELL_CYCLE | JAG2 |
| GO_CELL_CYCLE | JMJD5 |
| GO_CELL_CYCLE | JMY |
| GO_CELL_CYCLE | JTB |
| GO_CELL_CYCLE | JUB |

| | |
|---|---|
| GO_CELL_CYCLE | KAT2B |
| GO_CELL_CYCLE | KATNA1 |
| GO_CELL_CYCLE | KATNB1 |
| GO_CELL_CYCLE | KCTD11 |
| GO_CELL_CYCLE | KHDRBS1 |
| GO_CELL_CYCLE | KIAA0196 |
| GO_CELL_CYCLE | KIAA0430 |
| GO_CELL_CYCLE | KIAA0753 |
| GO_CELL_CYCLE | KIAA1377 |
| GO_CELL_CYCLE | KIAA1383 |
| GO_CELL_CYCLE | KIAA1967 |
| GO_CELL_CYCLE | KIF11 |
| GO_CELL_CYCLE | KIF13A |
| GO_CELL_CYCLE | KIF14 |
| GO_CELL_CYCLE | KIF15 |
| GO_CELL_CYCLE | KIF18A |
| GO_CELL_CYCLE | KIF18B |
| GO_CELL_CYCLE | KIF20A |
| GO_CELL_CYCLE | KIF20B |
| GO_CELL_CYCLE | KIF22 |
| GO_CELL_CYCLE | KIF23 |
| GO_CELL_CYCLE | KIF25 |
| GO_CELL_CYCLE | KIF2A |
| GO_CELL_CYCLE | KIF2B |
| GO_CELL_CYCLE | KIF2C |
| GO_CELL_CYCLE | KIF3B |
| GO_CELL_CYCLE | KIF4A |
| GO_CELL_CYCLE | KIF4B |
| GO_CELL_CYCLE | KIFC1 |
| GO_CELL_CYCLE | KLF11 |
| GO_CELL_CYCLE | KLHDC3 |
| GO_CELL_CYCLE | KLHDC5 |
| GO_CELL_CYCLE | KLHL13 |
| GO_CELL_CYCLE | KLHL21 |
| GO_CELL_CYCLE | KLHL22 |
| GO_CELL_CYCLE | KLHL9 |
| GO_CELL_CYCLE | KLK10 |
| GO_CELL_CYCLE | KLLN |
| GO_CELL_CYCLE | KNTC1 |
| GO_CELL_CYCLE | KPNB1 |
| GO_CELL_CYCLE | KRT18 |
| GO_CELL_CYCLE | LAMTOR1 |
| GO_CELL_CYCLE | LAMTOR2 |
| GO_CELL_CYCLE | LAMTOR3 |
| GO_CELL_CYCLE | LATS1 |
| GO_CELL_CYCLE | LATS2 |
| GO_CELL_CYCLE | LEPREL4 |

| | |
|---|---|
| GO_CELL_CYCLE | LFNG |
| GO_CELL_CYCLE | LIG1 |
| GO_CELL_CYCLE | LIG3 |
| GO_CELL_CYCLE | LIG4 |
| GO_CELL_CYCLE | LIN37 |
| GO_CELL_CYCLE | LIN52 |
| GO_CELL_CYCLE | LIN54 |
| GO_CELL_CYCLE | LIN9 |
| GO_CELL_CYCLE | LLGL1 |
| GO_CELL_CYCLE | LLGL2 |
| GO_CELL_CYCLE | LMLN |
| GO_CELL_CYCLE | LMNA |
| GO_CELL_CYCLE | LOC389493 |
| GO_CELL_CYCLE | LOC728637 |
| GO_CELL_CYCLE | LPIN1 |
| GO_CELL_CYCLE | LRRCC1 |
| GO_CELL_CYCLE | LZTS1 |
| GO_CELL_CYCLE | LZTS2 |
| GO_CELL_CYCLE | MACF1 |
| GO_CELL_CYCLE | MAD1L1 |
| GO_CELL_CYCLE | MAD2L1 |
| GO_CELL_CYCLE | MAD2L1BP |
| GO_CELL_CYCLE | MAD2L2 |
| GO_CELL_CYCLE | MAEA |
| GO_CELL_CYCLE | MAEL |
| GO_CELL_CYCLE | MAGI2 |
| GO_CELL_CYCLE | MAP2K1 |
| GO_CELL_CYCLE | MAP2K6 |
| GO_CELL_CYCLE | MAP3K8 |
| GO_CELL_CYCLE | MAP4 |
| GO_CELL_CYCLE | MAP9 |
| GO_CELL_CYCLE | MAPK1 |
| GO_CELL_CYCLE | MAPK12 |
| GO_CELL_CYCLE | MAPK13 |
| GO_CELL_CYCLE | MAPK14 |
| GO_CELL_CYCLE | MAPK3 |
| GO_CELL_CYCLE | MAPK4 |
| GO_CELL_CYCLE | MAPK6 |
| GO_CELL_CYCLE | MAPK7 |
| GO_CELL_CYCLE | MAPKAPK2 |
| GO_CELL_CYCLE | MAPRE1 |
| GO_CELL_CYCLE | MAPRE2 |
| GO_CELL_CYCLE | MAPRE3 |
| GO_CELL_CYCLE | MARK4 |
| GO_CELL_CYCLE | MARVELD1 |
| GO_CELL_CYCLE | MASTL |
| GO_CELL_CYCLE | MAU2 |

| | |
|---|---|
| GO_CELL_CYCLE | MCM10 |
| GO_CELL_CYCLE | MCM2 |
| GO_CELL_CYCLE | MCM3 |
| GO_CELL_CYCLE | MCM4 |
| GO_CELL_CYCLE | MCM5 |
| GO_CELL_CYCLE | MCM6 |
| GO_CELL_CYCLE | MCM7 |
| GO_CELL_CYCLE | MCM8 |
| GO_CELL_CYCLE | MCMBP |
| GO_CELL_CYCLE | MCPH1 |
| GO_CELL_CYCLE | MCTS1 |
| GO_CELL_CYCLE | MDC1 |
| GO_CELL_CYCLE | MDM2 |
| GO_CELL_CYCLE | MDM4 |
| GO_CELL_CYCLE | MEI1 |
| GO_CELL_CYCLE | MELK |
| GO_CELL_CYCLE | MEN1 |
| GO_CELL_CYCLE | METTL11A |
| GO_CELL_CYCLE | MFN2 |
| GO_CELL_CYCLE | MIS12 |
| GO_CELL_CYCLE | MIS18A |
| GO_CELL_CYCLE | MIS18BP1 |
| GO_CELL_CYCLE | MITD1 |
| GO_CELL_CYCLE | MKI67 |
| GO_CELL_CYCLE | MLF1 |
| GO_CELL_CYCLE | MLF1IP |
| GO_CELL_CYCLE | MLH1 |
| GO_CELL_CYCLE | MLH3 |
| GO_CELL_CYCLE | MLL5 |
| GO_CELL_CYCLE | MLST8 |
| GO_CELL_CYCLE | MNAT1 |
| GO_CELL_CYCLE | MND1 |
| GO_CELL_CYCLE | MNS1 |
| GO_CELL_CYCLE | MOS |
| GO_CELL_CYCLE | MOV10L1 |
| GO_CELL_CYCLE | MRE11A |
| GO_CELL_CYCLE | MRPL41 |
| GO_CELL_CYCLE | MSH2 |
| GO_CELL_CYCLE | MSH3 |
| GO_CELL_CYCLE | MSH4 |
| GO_CELL_CYCLE | MSH5 |
| GO_CELL_CYCLE | MSH6 |
| GO_CELL_CYCLE | MST4 |
| GO_CELL_CYCLE | MTBP |
| GO_CELL_CYCLE | MTOR |
| GO_CELL_CYCLE | MUC1 |
| GO_CELL_CYCLE | MYBL2 |

| | |
|---|---|
| GO_CELL_CYCLE | MYC |
| GO_CELL_CYCLE | MYH10 |
| GO_CELL_CYCLE | MYH9 |
| GO_CELL_CYCLE | MYOG |
| GO_CELL_CYCLE | NAA50 |
| GO_CELL_CYCLE | NAE1 |
| GO_CELL_CYCLE | NASP |
| GO_CELL_CYCLE | NBN |
| GO_CELL_CYCLE | NCAPD2 |
| GO_CELL_CYCLE | NCAPD3 |
| GO_CELL_CYCLE | NCAPG |
| GO_CELL_CYCLE | NCAPG2 |
| GO_CELL_CYCLE | NCAPH |
| GO_CELL_CYCLE | NCOR1 |
| GO_CELL_CYCLE | NDC80 |
| GO_CELL_CYCLE | NDE1 |
| GO_CELL_CYCLE | NDEL1 |
| GO_CELL_CYCLE | NEDD1 |
| GO_CELL_CYCLE | NEDD9 |
| GO_CELL_CYCLE | NEK1 |
| GO_CELL_CYCLE | NEK11 |
| GO_CELL_CYCLE | NEK2 |
| GO_CELL_CYCLE | NEK3 |
| GO_CELL_CYCLE | NEK4 |
| GO_CELL_CYCLE | NEK6 |
| GO_CELL_CYCLE | NEK7 |
| GO_CELL_CYCLE | NEK9 |
| GO_CELL_CYCLE | NES |
| GO_CELL_CYCLE | NINL |
| GO_CELL_CYCLE | NIPBL |
| GO_CELL_CYCLE | NKX3-1 |
| GO_CELL_CYCLE | NLRC4 |
| GO_CELL_CYCLE | NOLC1 |
| GO_CELL_CYCLE | NOTCH2 |
| GO_CELL_CYCLE | NOX5 |
| GO_CELL_CYCLE | NPAT |
| GO_CELL_CYCLE | NPM1 |
| GO_CELL_CYCLE | NR2C2 |
| GO_CELL_CYCLE | NSL1 |
| GO_CELL_CYCLE | NSMCE2 |
| GO_CELL_CYCLE | NSUN2 |
| GO_CELL_CYCLE | NUDC |
| GO_CELL_CYCLE | NUF2 |
| GO_CELL_CYCLE | NUMA1 |
| GO_CELL_CYCLE | NUP107 |
| GO_CELL_CYCLE | NUP133 |
| GO_CELL_CYCLE | NUP153 |

| | |
|---|---|
| GO_CELL_CYCLE | NUP155 |
| GO_CELL_CYCLE | NUP160 |
| GO_CELL_CYCLE | NUP188 |
| GO_CELL_CYCLE | NUP205 |
| GO_CELL_CYCLE | NUP210 |
| GO_CELL_CYCLE | NUP214 |
| GO_CELL_CYCLE | NUP35 |
| GO_CELL_CYCLE | NUP37 |
| GO_CELL_CYCLE | NUP43 |
| GO_CELL_CYCLE | NUP50 |
| GO_CELL_CYCLE | NUP54 |
| GO_CELL_CYCLE | NUP62 |
| GO_CELL_CYCLE | NUP85 |
| GO_CELL_CYCLE | NUP88 |
| GO_CELL_CYCLE | NUP93 |
| GO_CELL_CYCLE | NUP98 |
| GO_CELL_CYCLE | NUPL1 |
| GO_CELL_CYCLE | NUPL2 |
| GO_CELL_CYCLE | NUSAP1 |
| GO_CELL_CYCLE | OBFC2A |
| GO_CELL_CYCLE | OBFC2B |
| GO_CELL_CYCLE | ODF2 |
| GO_CELL_CYCLE | OFD1 |
| GO_CELL_CYCLE | OIP5 |
| GO_CELL_CYCLE | OPTN |
| GO_CELL_CYCLE | ORC1 |
| GO_CELL_CYCLE | ORC2 |
| GO_CELL_CYCLE | ORC3 |
| GO_CELL_CYCLE | ORC4 |
| GO_CELL_CYCLE | ORC5 |
| GO_CELL_CYCLE | ORC6 |
| GO_CELL_CYCLE | OSGIN2 |
| GO_CELL_CYCLE | OVOL1 |
| GO_CELL_CYCLE | PA2G4 |
| GO_CELL_CYCLE | PAFAH1B1 |
| GO_CELL_CYCLE | PAK1 |
| GO_CELL_CYCLE | PAK2 |
| GO_CELL_CYCLE | PAK3 |
| GO_CELL_CYCLE | PAK4 |
| GO_CELL_CYCLE | PAK6 |
| GO_CELL_CYCLE | PAK7 |
| GO_CELL_CYCLE | PAPD5 |
| GO_CELL_CYCLE | PAPD7 |
| GO_CELL_CYCLE | PARD3 |
| GO_CELL_CYCLE | PARD3B |
| GO_CELL_CYCLE | PARD6A |
| GO_CELL_CYCLE | PARD6B |

| | |
|---|---|
| GO_CELL_CYCLE | PARD6G |
| GO_CELL_CYCLE | PAX6 |
| GO_CELL_CYCLE | PBK |
| GO_CELL_CYCLE | PBRM1 |
| GO_CELL_CYCLE | PCBP4 |
| GO_CELL_CYCLE | PCM1 |
| GO_CELL_CYCLE | PCNA |
| GO_CELL_CYCLE | PCNP |
| GO_CELL_CYCLE | PCNT |
| GO_CELL_CYCLE | PDCD2L |
| GO_CELL_CYCLE | PDCD6IP |
| GO_CELL_CYCLE | PDS5A |
| GO_CELL_CYCLE | PDS5B |
| GO_CELL_CYCLE | PEA15 |
| GO_CELL_CYCLE | PELO |
| GO_CELL_CYCLE | PFDN1 |
| GO_CELL_CYCLE | PHB2 |
| GO_CELL_CYCLE | PHF13 |
| GO_CELL_CYCLE | PHF8 |
| GO_CELL_CYCLE | PHGDH |
| GO_CELL_CYCLE | PHLDA1 |
| GO_CELL_CYCLE | PIBF1 |
| GO_CELL_CYCLE | PIDD |
| GO_CELL_CYCLE | PIK3C3 |
| GO_CELL_CYCLE | PIM1 |
| GO_CELL_CYCLE | PIM2 |
| GO_CELL_CYCLE | PIM3 |
| GO_CELL_CYCLE | PIN1 |
| GO_CELL_CYCLE | PINX1 |
| GO_CELL_CYCLE | PIWIL1 |
| GO_CELL_CYCLE | PIWIL2 |
| GO_CELL_CYCLE | PIWIL3 |
| GO_CELL_CYCLE | PIWIL4 |
| GO_CELL_CYCLE | PKD1 |
| GO_CELL_CYCLE | PKD2 |
| GO_CELL_CYCLE | PKMYT1 |
| GO_CELL_CYCLE | PKN2 |
| GO_CELL_CYCLE | PLAGL1 |
| GO_CELL_CYCLE | PLCB1 |
| GO_CELL_CYCLE | PLD6 |
| GO_CELL_CYCLE | PLK1 |
| GO_CELL_CYCLE | PLK1S1 |
| GO_CELL_CYCLE | PLK2 |
| GO_CELL_CYCLE | PLK3 |
| GO_CELL_CYCLE | PLK4 |
| GO_CELL_CYCLE | PLK5 |
| GO_CELL_CYCLE | PMF1 |

| | |
|---|---|
| GO_CELL_CYCLE | PML |
| GO_CELL_CYCLE | PNPT1 |
| GO_CELL_CYCLE | POC5 |
| GO_CELL_CYCLE | POGZ |
| GO_CELL_CYCLE | POLA1 |
| GO_CELL_CYCLE | POLA2 |
| GO_CELL_CYCLE | POLE |
| GO_CELL_CYCLE | POLE2 |
| GO_CELL_CYCLE | POM121 |
| GO_CELL_CYCLE | POM121C |
| GO_CELL_CYCLE | PPAT |
| GO_CELL_CYCLE | PPM1A |
| GO_CELL_CYCLE | PPM1D |
| GO_CELL_CYCLE | PPM1G |
| GO_CELL_CYCLE | PPP1CA |
| GO_CELL_CYCLE | PPP1CB |
| GO_CELL_CYCLE | PPP1CC |
| GO_CELL_CYCLE | PPP1R12A |
| GO_CELL_CYCLE | PPP1R12B |
| GO_CELL_CYCLE | PPP1R15A |
| GO_CELL_CYCLE | PPP1R1C |
| GO_CELL_CYCLE | PPP1R9B |
| GO_CELL_CYCLE | PPP2CA |
| GO_CELL_CYCLE | PPP2R1A |
| GO_CELL_CYCLE | PPP2R2A |
| GO_CELL_CYCLE | PPP2R2D |
| GO_CELL_CYCLE | PPP2R3B |
| GO_CELL_CYCLE | PPP2R4 |
| GO_CELL_CYCLE | PPP2R5C |
| GO_CELL_CYCLE | PPP3CA |
| GO_CELL_CYCLE | PPP5C |
| GO_CELL_CYCLE | PPP6C |
| GO_CELL_CYCLE | PRC1 |
| GO_CELL_CYCLE | PRCC |
| GO_CELL_CYCLE | PRDM5 |
| GO_CELL_CYCLE | PRDM9 |
| GO_CELL_CYCLE | PRIM1 |
| GO_CELL_CYCLE | PRIM2 |
| GO_CELL_CYCLE | PRKAA1 |
| GO_CELL_CYCLE | PRKAA2 |
| GO_CELL_CYCLE | PRKAB1 |
| GO_CELL_CYCLE | PRKAB2 |
| GO_CELL_CYCLE | PRKACA |
| GO_CELL_CYCLE | PRKAG1 |
| GO_CELL_CYCLE | PRKAG2 |
| GO_CELL_CYCLE | PRKAG3 |
| GO_CELL_CYCLE | PRKAR1A |

| | |
|---|---|
| GO_CELL_CYCLE | PRKAR2B |
| GO_CELL_CYCLE | PRKCA |
| GO_CELL_CYCLE | PRKCB |
| GO_CELL_CYCLE | PRKCD |
| GO_CELL_CYCLE | PRKCE |
| GO_CELL_CYCLE | PRKDC |
| GO_CELL_CYCLE | PRMT1 |
| GO_CELL_CYCLE | PRNP |
| GO_CELL_CYCLE | PRPF19 |
| GO_CELL_CYCLE | PRPF40A |
| GO_CELL_CYCLE | PRR5 |
| GO_CELL_CYCLE | PSMA1 |
| GO_CELL_CYCLE | PSMA2 |
| GO_CELL_CYCLE | PSMA3 |
| GO_CELL_CYCLE | PSMA4 |
| GO_CELL_CYCLE | PSMA5 |
| GO_CELL_CYCLE | PSMA6 |
| GO_CELL_CYCLE | PSMA7 |
| GO_CELL_CYCLE | PSMB1 |
| GO_CELL_CYCLE | PSMB10 |
| GO_CELL_CYCLE | PSMB2 |
| GO_CELL_CYCLE | PSMB3 |
| GO_CELL_CYCLE | PSMB4 |
| GO_CELL_CYCLE | PSMB5 |
| GO_CELL_CYCLE | PSMB6 |
| GO_CELL_CYCLE | PSMB7 |
| GO_CELL_CYCLE | PSMB8 |
| GO_CELL_CYCLE | PSMB9 |
| GO_CELL_CYCLE | PSMC1 |
| GO_CELL_CYCLE | PSMC2 |
| GO_CELL_CYCLE | PSMC3 |
| GO_CELL_CYCLE | PSMC3IP |
| GO_CELL_CYCLE | PSMC4 |
| GO_CELL_CYCLE | PSMC5 |
| GO_CELL_CYCLE | PSMC6 |
| GO_CELL_CYCLE | PSMD1 |
| GO_CELL_CYCLE | PSMD10 |
| GO_CELL_CYCLE | PSMD11 |
| GO_CELL_CYCLE | PSMD12 |
| GO_CELL_CYCLE | PSMD13 |
| GO_CELL_CYCLE | PSMD14 |
| GO_CELL_CYCLE | PSMD2 |
| GO_CELL_CYCLE | PSMD3 |
| GO_CELL_CYCLE | PSMD4 |
| GO_CELL_CYCLE | PSMD5 |
| GO_CELL_CYCLE | PSMD6 |
| GO_CELL_CYCLE | PSMD7 |

| | |
|---|---|
| GO_CELL_CYCLE | PSMD8 |
| GO_CELL_CYCLE | PSMD9 |
| GO_CELL_CYCLE | PSME1 |
| GO_CELL_CYCLE | PSME2 |
| GO_CELL_CYCLE | PSME3 |
| GO_CELL_CYCLE | PSMF1 |
| GO_CELL_CYCLE | PSMG2 |
| GO_CELL_CYCLE | PSRC1 |
| GO_CELL_CYCLE | PTEN |
| GO_CELL_CYCLE | PTP4A1 |
| GO_CELL_CYCLE | PTPN11 |
| GO_CELL_CYCLE | PTPN6 |
| GO_CELL_CYCLE | PTPRC |
| GO_CELL_CYCLE | PTTG1 |
| GO_CELL_CYCLE | PTTG2 |
| GO_CELL_CYCLE | PTTG3P |
| GO_CELL_CYCLE | PYHIN1 |
| GO_CELL_CYCLE | RAB11A |
| GO_CELL_CYCLE | RAB11FIP3 |
| GO_CELL_CYCLE | RAB11FIP4 |
| GO_CELL_CYCLE | RAB1A |
| GO_CELL_CYCLE | RAB35 |
| GO_CELL_CYCLE | RAB6C |
| GO_CELL_CYCLE | RAB8A |
| GO_CELL_CYCLE | RABGAP1 |
| GO_CELL_CYCLE | RACGAP1 |
| GO_CELL_CYCLE | RAD1 |
| GO_CELL_CYCLE | RAD17 |
| GO_CELL_CYCLE | RAD21 |
| GO_CELL_CYCLE | RAD21L1 |
| GO_CELL_CYCLE | RAD50 |
| GO_CELL_CYCLE | RAD51 |
| GO_CELL_CYCLE | RAD51B |
| GO_CELL_CYCLE | RAD51C |
| GO_CELL_CYCLE | RAD51D |
| GO_CELL_CYCLE | RAD54B |
| GO_CELL_CYCLE | RAD54L |
| GO_CELL_CYCLE | RAD9A |
| GO_CELL_CYCLE | RAD9B |
| GO_CELL_CYCLE | RAE1 |
| GO_CELL_CYCLE | RALA |
| GO_CELL_CYCLE | RALB |
| GO_CELL_CYCLE | RAN |
| GO_CELL_CYCLE | RANBP1 |
| GO_CELL_CYCLE | RANBP2 |
| GO_CELL_CYCLE | RANGAP1 |
| GO_CELL_CYCLE | RASA1 |

| | |
|---|---|
| GO_CELL_CYCLE | RASSF1 |
| GO_CELL_CYCLE | RASSF2 |
| GO_CELL_CYCLE | RASSF4 |
| GO_CELL_CYCLE | RB1 |
| GO_CELL_CYCLE | RB1CC1 |
| GO_CELL_CYCLE | RBBP4 |
| GO_CELL_CYCLE | RBBP8 |
| GO_CELL_CYCLE | RBL1 |
| GO_CELL_CYCLE | RBL2 |
| GO_CELL_CYCLE | RBM38 |
| GO_CELL_CYCLE | RBM7 |
| GO_CELL_CYCLE | RCBTB1 |
| GO_CELL_CYCLE | RCC1 |
| GO_CELL_CYCLE | RCC2 |
| GO_CELL_CYCLE | REC8 |
| GO_CELL_CYCLE | RECQL5 |
| GO_CELL_CYCLE | REEP3 |
| GO_CELL_CYCLE | REEP4 |
| GO_CELL_CYCLE | RFWD3 |
| GO_CELL_CYCLE | RGS14 |
| GO_CELL_CYCLE | RGS2 |
| GO_CELL_CYCLE | RHEB |
| GO_CELL_CYCLE | RHOA |
| GO_CELL_CYCLE | RHOB |
| GO_CELL_CYCLE | RHOC |
| GO_CELL_CYCLE | RHOU |
| GO_CELL_CYCLE | RIF1 |
| GO_CELL_CYCLE | RINT1 |
| GO_CELL_CYCLE | RNF2 |
| GO_CELL_CYCLE | RNF212 |
| GO_CELL_CYCLE | RNF8 |
| GO_CELL_CYCLE | ROCK2 |
| GO_CELL_CYCLE | ROPN1B |
| GO_CELL_CYCLE | RPA1 |
| GO_CELL_CYCLE | RPA2 |
| GO_CELL_CYCLE | RPA3 |
| GO_CELL_CYCLE | RPA4 |
| GO_CELL_CYCLE | RPRM |
| GO_CELL_CYCLE | RPS27 |
| GO_CELL_CYCLE | RPS27A |
| GO_CELL_CYCLE | RPS27L |
| GO_CELL_CYCLE | RPS3 |
| GO_CELL_CYCLE | RPS6 |
| GO_CELL_CYCLE | RPS6KA1 |
| GO_CELL_CYCLE | RPS6KA3 |
| GO_CELL_CYCLE | RPS6KB1 |
| GO_CELL_CYCLE | RPTOR |

| | |
|---|---|
| GO_CELL_CYCLE | RQCD1 |
| GO_CELL_CYCLE | RRAGA |
| GO_CELL_CYCLE | RRAGB |
| GO_CELL_CYCLE | RRAGC |
| GO_CELL_CYCLE | RRAGD |
| GO_CELL_CYCLE | RRM1 |
| GO_CELL_CYCLE | RRM2 |
| GO_CELL_CYCLE | RRS1 |
| GO_CELL_CYCLE | RSPH1 |
| GO_CELL_CYCLE | RTEL1 |
| GO_CELL_CYCLE | RUVBL1 |
| GO_CELL_CYCLE | SAC3D1 |
| GO_CELL_CYCLE | SART1 |
| GO_CELL_CYCLE | SASS6 |
| GO_CELL_CYCLE | SBDS |
| GO_CELL_CYCLE | SDCCAG3 |
| GO_CELL_CYCLE | SDCCAG8 |
| GO_CELL_CYCLE | SEC13 |
| GO_CELL_CYCLE | SEH1L |
| GO_CELL_CYCLE | SENP5 |
| GO_CELL_CYCLE | 1-Sep |
| GO_CELL_CYCLE | 10-Sep |
| GO_CELL_CYCLE | 11-Sep |
| GO_CELL_CYCLE | 12-Sep |
| GO_CELL_CYCLE | 14-Sep |
| GO_CELL_CYCLE | 2-Sep |
| GO_CELL_CYCLE | 3-Sep |
| GO_CELL_CYCLE | 4-Sep |
| GO_CELL_CYCLE | 5-Sep |
| GO_CELL_CYCLE | 6-Sep |
| GO_CELL_CYCLE | 7-Sep |
| GO_CELL_CYCLE | 9-Sep |
| GO_CELL_CYCLE | SESN1 |
| GO_CELL_CYCLE | SETD8 |
| GO_CELL_CYCLE | SETDB2 |
| GO_CELL_CYCLE | SETMAR |
| GO_CELL_CYCLE | SFI1 |
| GO_CELL_CYCLE | SFN |
| GO_CELL_CYCLE | SGOL1 |
| GO_CELL_CYCLE | SGOL2 |
| GO_CELL_CYCLE | SGSM3 |
| GO_CELL_CYCLE | SIAH1 |
| GO_CELL_CYCLE | SIAH2 |
| GO_CELL_CYCLE | SIK1 |
| GO_CELL_CYCLE | SIRT2 |
| GO_CELL_CYCLE | SIRT7 |
| GO_CELL_CYCLE | SKA1 |

| | |
|---|---|
| GO_CELL_CYCLE | SKA2 |
| GO_CELL_CYCLE | SKA3 |
| GO_CELL_CYCLE | SKIL |
| GO_CELL_CYCLE | SKP1 |
| GO_CELL_CYCLE | SKP2 |
| GO_CELL_CYCLE | SLBP |
| GO_CELL_CYCLE | SLC26A8 |
| GO_CELL_CYCLE | SLC2A8 |
| GO_CELL_CYCLE | SLC39A5 |
| GO_CELL_CYCLE | SMAD3 |
| GO_CELL_CYCLE | SMARCA4 |
| GO_CELL_CYCLE | SMARCAD1 |
| GO_CELL_CYCLE | SMARCB1 |
| GO_CELL_CYCLE | SMC1A |
| GO_CELL_CYCLE | SMC1B |
| GO_CELL_CYCLE | SMC2 |
| GO_CELL_CYCLE | SMC3 |
| GO_CELL_CYCLE | SMC4 |
| GO_CELL_CYCLE | SMC5 |
| GO_CELL_CYCLE | SMPD3 |
| GO_CELL_CYCLE | SNX18 |
| GO_CELL_CYCLE | SNX33 |
| GO_CELL_CYCLE | SNX9 |
| GO_CELL_CYCLE | SON |
| GO_CELL_CYCLE | SOX11 |
| GO_CELL_CYCLE | SOX2 |
| GO_CELL_CYCLE | SOX4 |
| GO_CELL_CYCLE | SPAG5 |
| GO_CELL_CYCLE | SPANXA2-OT1 |
| GO_CELL_CYCLE | SPAST |
| GO_CELL_CYCLE | SPATA22 |
| GO_CELL_CYCLE | SPC24 |
| GO_CELL_CYCLE | SPC25 |
| GO_CELL_CYCLE | SPDYA |
| GO_CELL_CYCLE | SPDYC |
| GO_CELL_CYCLE | SPECC1L |
| GO_CELL_CYCLE | SPICE1 |
| GO_CELL_CYCLE | SPIN1 |
| GO_CELL_CYCLE | SPIN2A |
| GO_CELL_CYCLE | SPIN2B |
| GO_CELL_CYCLE | SPIRE1 |
| GO_CELL_CYCLE | SPIRE2 |
| GO_CELL_CYCLE | SPO11 |
| GO_CELL_CYCLE | SPRY1 |
| GO_CELL_CYCLE | SPRY2 |
| GO_CELL_CYCLE | SPTBN1 |
| GO_CELL_CYCLE | SRC |

| | |
|---|---|
| GO_CELL_CYCLE | SSNA1 |
| GO_CELL_CYCLE | SSSCA1 |
| GO_CELL_CYCLE | STAG1 |
| GO_CELL_CYCLE | STAG2 |
| GO_CELL_CYCLE | STAG3 |
| GO_CELL_CYCLE | STAMBP |
| GO_CELL_CYCLE | STARD9 |
| GO_CELL_CYCLE | STEAP3 |
| GO_CELL_CYCLE | STIL |
| GO_CELL_CYCLE | STK10 |
| GO_CELL_CYCLE | STK11 |
| GO_CELL_CYCLE | STK24 |
| GO_CELL_CYCLE | STMN1 |
| GO_CELL_CYCLE | STOX1 |
| GO_CELL_CYCLE | STRA13 |
| GO_CELL_CYCLE | STRA8 |
| GO_CELL_CYCLE | STRADA |
| GO_CELL_CYCLE | STRADB |
| GO_CELL_CYCLE | SUGT1 |
| GO_CELL_CYCLE | SUN1 |
| GO_CELL_CYCLE | SUN2 |
| GO_CELL_CYCLE | SUPT5H |
| GO_CELL_CYCLE | SUV39H1 |
| GO_CELL_CYCLE | SUV39H2 |
| GO_CELL_CYCLE | SYCE1 |
| GO_CELL_CYCLE | SYCE1L |
| GO_CELL_CYCLE | SYCE2 |
| GO_CELL_CYCLE | SYCE3 |
| GO_CELL_CYCLE | SYCP1 |
| GO_CELL_CYCLE | SYCP2 |
| GO_CELL_CYCLE | SYCP3 |
| GO_CELL_CYCLE | SYF2 |
| GO_CELL_CYCLE | TACC1 |
| GO_CELL_CYCLE | TADA3 |
| GO_CELL_CYCLE | TAF1 |
| GO_CELL_CYCLE | TAF1L |
| GO_CELL_CYCLE | TAF2 |
| GO_CELL_CYCLE | TAOK1 |
| GO_CELL_CYCLE | TAOK2 |
| GO_CELL_CYCLE | TAOK3 |
| GO_CELL_CYCLE | TBRG1 |
| GO_CELL_CYCLE | TBRG4 |
| GO_CELL_CYCLE | TCF7L2 |
| GO_CELL_CYCLE | TDRD1 |
| GO_CELL_CYCLE | TDRD12 |
| GO_CELL_CYCLE | TDRD9 |
| GO_CELL_CYCLE | TDRKH |

| | |
|---|---|
| GO_CELL_CYCLE | TERF1 |
| GO_CELL_CYCLE | TERF2 |
| GO_CELL_CYCLE | TET2 |
| GO_CELL_CYCLE | TEX11 |
| GO_CELL_CYCLE | TEX12 |
| GO_CELL_CYCLE | TEX14 |
| GO_CELL_CYCLE | TEX15 |
| GO_CELL_CYCLE | TEX19 |
| GO_CELL_CYCLE | TFDP1 |
| GO_CELL_CYCLE | TFDP2 |
| GO_CELL_CYCLE | TFDP3 |
| GO_CELL_CYCLE | TGFB1 |
| GO_CELL_CYCLE | TGFB2 |
| GO_CELL_CYCLE | TGFBR1 |
| GO_CELL_CYCLE | THAP1 |
| GO_CELL_CYCLE | THAP5 |
| GO_CELL_CYCLE | THBS1 |
| GO_CELL_CYCLE | TIMELESS |
| GO_CELL_CYCLE | TIPIN |
| GO_CELL_CYCLE | TIPRL |
| GO_CELL_CYCLE | TLK1 |
| GO_CELL_CYCLE | TLK2 |
| GO_CELL_CYCLE | TMEM188 |
| GO_CELL_CYCLE | TMEM48 |
| GO_CELL_CYCLE | TMPRSS11A |
| GO_CELL_CYCLE | TNKS |
| GO_CELL_CYCLE | TNKS1BP1 |
| GO_CELL_CYCLE | TOP2A |
| GO_CELL_CYCLE | TOP2B |
| GO_CELL_CYCLE | TOP3A |
| GO_CELL_CYCLE | TP53 |
| GO_CELL_CYCLE | TP53BP1 |
| GO_CELL_CYCLE | TP53BP2 |
| GO_CELL_CYCLE | TP53INP1 |
| GO_CELL_CYCLE | TP63 |
| GO_CELL_CYCLE | TP73 |
| GO_CELL_CYCLE | TPD52L1 |
| GO_CELL_CYCLE | TPR |
| GO_CELL_CYCLE | TPX2 |
| GO_CELL_CYCLE | TRIAP1 |
| GO_CELL_CYCLE | TRIM21 |
| GO_CELL_CYCLE | TRIM71 |
| GO_CELL_CYCLE | TRIOBP |
| GO_CELL_CYCLE | TRIP13 |
| GO_CELL_CYCLE | TRNP1 |
| GO_CELL_CYCLE | TSC1 |
| GO_CELL_CYCLE | TSC2 |

| | |
|---|---|
| GO_CELL_CYCLE | TSG101 |
| GO_CELL_CYCLE | TSPYL2 |
| GO_CELL_CYCLE | TTC19 |
| GO_CELL_CYCLE | TTC28 |
| GO_CELL_CYCLE | TTK |
| GO_CELL_CYCLE | TTN |
| GO_CELL_CYCLE | TTYH1 |
| GO_CELL_CYCLE | TUBA1A |
| GO_CELL_CYCLE | TUBA4A |
| GO_CELL_CYCLE | TUBB |
| GO_CELL_CYCLE | TUBB1 |
| GO_CELL_CYCLE | TUBB3 |
| GO_CELL_CYCLE | TUBB4A |
| GO_CELL_CYCLE | TUBB4B |
| GO_CELL_CYCLE | TUBE1 |
| GO_CELL_CYCLE | TUBG1 |
| GO_CELL_CYCLE | TUBGCP2 |
| GO_CELL_CYCLE | TUBGCP3 |
| GO_CELL_CYCLE | TUBGCP4 |
| GO_CELL_CYCLE | TUBGCP5 |
| GO_CELL_CYCLE | TUBGCP6 |
| GO_CELL_CYCLE | TUSC2 |
| GO_CELL_CYCLE | TXLNG |
| GO_CELL_CYCLE | TXNIP |
| GO_CELL_CYCLE | TXNL4A |
| GO_CELL_CYCLE | TXNL4B |
| GO_CELL_CYCLE | TYMS |
| GO_CELL_CYCLE | UBA3 |
| GO_CELL_CYCLE | UBA52 |
| GO_CELL_CYCLE | UBB |
| GO_CELL_CYCLE | UBC |
| GO_CELL_CYCLE | UBE2B |
| GO_CELL_CYCLE | UBE2C |
| GO_CELL_CYCLE | UBE2D1 |
| GO_CELL_CYCLE | UBE2E1 |
| GO_CELL_CYCLE | UBE2I |
| GO_CELL_CYCLE | UBE2L3 |
| GO_CELL_CYCLE | UBE2S |
| GO_CELL_CYCLE | UBR2 |
| GO_CELL_CYCLE | UHMK1 |
| GO_CELL_CYCLE | UHRF1 |
| GO_CELL_CYCLE | UHRF2 |
| GO_CELL_CYCLE | UIMC1 |
| GO_CELL_CYCLE | UPF1 |
| GO_CELL_CYCLE | URGCP |
| GO_CELL_CYCLE | USH1C |
| GO_CELL_CYCLE | USP16 |

| | |
|---|---|
| GO_CELL_CYCLE | USP17L2 |
| GO_CELL_CYCLE | USP2 |
| GO_CELL_CYCLE | USP22 |
| GO_CELL_CYCLE | USP28 |
| GO_CELL_CYCLE | USP3 |
| GO_CELL_CYCLE | USP33 |
| GO_CELL_CYCLE | USP37 |
| GO_CELL_CYCLE | USP39 |
| GO_CELL_CYCLE | USP44 |
| GO_CELL_CYCLE | USP8 |
| GO_CELL_CYCLE | USP9X |
| GO_CELL_CYCLE | UTP14C |
| GO_CELL_CYCLE | UVRAG |
| GO_CELL_CYCLE | VASH1 |
| GO_CELL_CYCLE | VCPIP1 |
| GO_CELL_CYCLE | VPS4A |
| GO_CELL_CYCLE | VPS4B |
| GO_CELL_CYCLE | VRK1 |
| GO_CELL_CYCLE | WAC |
| GO_CELL_CYCLE | WAPAL |
| GO_CELL_CYCLE | WASL |
| GO_CELL_CYCLE | WBP2NL |
| GO_CELL_CYCLE | WDR43 |
| GO_CELL_CYCLE | WDR6 |
| GO_CELL_CYCLE | WDR62 |
| GO_CELL_CYCLE | WDR81 |
| GO_CELL_CYCLE | WEE1 |
| GO_CELL_CYCLE | WEE2 |
| GO_CELL_CYCLE | WNT10B |
| GO_CELL_CYCLE | WNT9A |
| GO_CELL_CYCLE | WTAP |
| GO_CELL_CYCLE | XIAP |
| GO_CELL_CYCLE | XPC |
| GO_CELL_CYCLE | XPO1 |
| GO_CELL_CYCLE | XRCC2 |
| GO_CELL_CYCLE | XRCC3 |
| GO_CELL_CYCLE | YEATS4 |
| GO_CELL_CYCLE | YWHAE |
| GO_CELL_CYCLE | YWHAG |
| GO_CELL_CYCLE | ZAK |
| GO_CELL_CYCLE | ZBTB49 |
| GO_CELL_CYCLE | ZC3HC1 |
| GO_CELL_CYCLE | ZFHX3 |
| GO_CELL_CYCLE | ZFP42 |
| GO_CELL_CYCLE | ZFYVE19 |
| GO_CELL_CYCLE | ZFYVE26 |
| GO_CELL_CYCLE | ZMYND11 |

| | |
|---|---|
| GO_CELL_CYCLE | ZNF16 |
| GO_CELL_CYCLE | ZNF207 |
| GO_CELL_CYCLE | ZNF259 |
| GO_CELL_CYCLE | ZNF268 |
| GO_CELL_CYCLE | ZNF318 |
| GO_CELL_CYCLE | ZNF365 |
| GO_CELL_CYCLE | ZNF385A |
| GO_CELL_CYCLE | ZNF503 |
| GO_CELL_CYCLE | ZNF830 |
| GO_CELL_CYCLE | ZW10 |
| GO_CELL_CYCLE | ZWILCH |
| GO_CELL_CYCLE | ZWINT |
| GO_APOPTOTIC_SIGNALING_PATHWAY | ABL1 |
| GO_APOPTOTIC_SIGNALING_PATHWAY | ACVR1B |
| GO_APOPTOTIC_SIGNALING_PATHWAY | ADORA1 |
| GO_APOPTOTIC_SIGNALING_PATHWAY | AEN |
| GO_APOPTOTIC_SIGNALING_PATHWAY | AIFM1 |
| GO_APOPTOTIC_SIGNALING_PATHWAY | ANXA6 |
| GO_APOPTOTIC_SIGNALING_PATHWAY | APAF1 |
| GO_APOPTOTIC_SIGNALING_PATHWAY | APOPT1 |
| GO_APOPTOTIC_SIGNALING_PATHWAY | APPL1 |
| GO_APOPTOTIC_SIGNALING_PATHWAY | ARL6IP5 |
| GO_APOPTOTIC_SIGNALING_PATHWAY | ATF4 |
| GO_APOPTOTIC_SIGNALING_PATHWAY | ATM |
| GO_APOPTOTIC_SIGNALING_PATHWAY | ATP2A1 |
| GO_APOPTOTIC_SIGNALING_PATHWAY | BAD |
| GO_APOPTOTIC_SIGNALING_PATHWAY | BAG3 |
| GO_APOPTOTIC_SIGNALING_PATHWAY | BAG6 |
| GO_APOPTOTIC_SIGNALING_PATHWAY | BAK1 |
| GO_APOPTOTIC_SIGNALING_PATHWAY | BAX |
| GO_APOPTOTIC_SIGNALING_PATHWAY | BBC3 |
| GO_APOPTOTIC_SIGNALING_PATHWAY | BCL2 |
| GO_APOPTOTIC_SIGNALING_PATHWAY | BCL2A1 |
| GO_APOPTOTIC_SIGNALING_PATHWAY | BCL2L1 |
| GO_APOPTOTIC_SIGNALING_PATHWAY | BCL2L10 |
| GO_APOPTOTIC_SIGNALING_PATHWAY | BCL2L11 |
| GO_APOPTOTIC_SIGNALING_PATHWAY | BCL2L2 |
| GO_APOPTOTIC_SIGNALING_PATHWAY | BCL3 |
| GO_APOPTOTIC_SIGNALING_PATHWAY | BID |
| GO_APOPTOTIC_SIGNALING_PATHWAY | BLOC1S2 |
| GO_APOPTOTIC_SIGNALING_PATHWAY | BMF |
| GO_APOPTOTIC_SIGNALING_PATHWAY | BNIP3 |
| GO_APOPTOTIC_SIGNALING_PATHWAY | BNIP3L |
| GO_APOPTOTIC_SIGNALING_PATHWAY | BOK |
| GO_APOPTOTIC_SIGNALING_PATHWAY | BRCA1 |
| GO_APOPTOTIC_SIGNALING_PATHWAY | BRCA2 |
| GO_APOPTOTIC_SIGNALING_PATHWAY | BRSK2 |

| | |
|---|---|
| GO_APOPTOTIC_SIGNALING_PATHWAY | BTK |
| GO_APOPTOTIC_SIGNALING_PATHWAY | C16orf5 |
| GO_APOPTOTIC_SIGNALING_PATHWAY | C22orf29 |
| GO_APOPTOTIC_SIGNALING_PATHWAY | CASP10 |
| GO_APOPTOTIC_SIGNALING_PATHWAY | CASP12 |
| GO_APOPTOTIC_SIGNALING_PATHWAY | CASP2 |
| GO_APOPTOTIC_SIGNALING_PATHWAY | CASP3 |
| GO_APOPTOTIC_SIGNALING_PATHWAY | CASP4 |
| GO_APOPTOTIC_SIGNALING_PATHWAY | CASP8 |
| GO_APOPTOTIC_SIGNALING_PATHWAY | CASP8AP2 |
| GO_APOPTOTIC_SIGNALING_PATHWAY | CASP9 |
| GO_APOPTOTIC_SIGNALING_PATHWAY | CAV1 |
| GO_APOPTOTIC_SIGNALING_PATHWAY | CCK |
| GO_APOPTOTIC_SIGNALING_PATHWAY | CD14 |
| GO_APOPTOTIC_SIGNALING_PATHWAY | CD24 |
| GO_APOPTOTIC_SIGNALING_PATHWAY | CD27 |
| GO_APOPTOTIC_SIGNALING_PATHWAY | CD28 |
| GO_APOPTOTIC_SIGNALING_PATHWAY | CD38 |
| GO_APOPTOTIC_SIGNALING_PATHWAY | CD3E |
| GO_APOPTOTIC_SIGNALING_PATHWAY | CD40 |
| GO_APOPTOTIC_SIGNALING_PATHWAY | CD5 |
| GO_APOPTOTIC_SIGNALING_PATHWAY | CD70 |
| GO_APOPTOTIC_SIGNALING_PATHWAY | CDKN1A |
| GO_APOPTOTIC_SIGNALING_PATHWAY | CEBPB |
| GO_APOPTOTIC_SIGNALING_PATHWAY | CHAC1 |
| GO_APOPTOTIC_SIGNALING_PATHWAY | CHEK2 |
| GO_APOPTOTIC_SIGNALING_PATHWAY | CIB1 |
| GO_APOPTOTIC_SIGNALING_PATHWAY | CIDEB |
| GO_APOPTOTIC_SIGNALING_PATHWAY | CLU |
| GO_APOPTOTIC_SIGNALING_PATHWAY | CRADD |
| GO_APOPTOTIC_SIGNALING_PATHWAY | CRH |
| GO_APOPTOTIC_SIGNALING_PATHWAY | CRIP1 |
| GO_APOPTOTIC_SIGNALING_PATHWAY | CUL1 |
| GO_APOPTOTIC_SIGNALING_PATHWAY | CUL2 |
| GO_APOPTOTIC_SIGNALING_PATHWAY | CUL3 |
| GO_APOPTOTIC_SIGNALING_PATHWAY | CUL4A |
| GO_APOPTOTIC_SIGNALING_PATHWAY | CUL5 |
| GO_APOPTOTIC_SIGNALING_PATHWAY | CYCS |
| GO_APOPTOTIC_SIGNALING_PATHWAY | CYP1B1 |
| GO_APOPTOTIC_SIGNALING_PATHWAY | DAB2IP |
| GO_APOPTOTIC_SIGNALING_PATHWAY | DAP |
| GO_APOPTOTIC_SIGNALING_PATHWAY | DAP3 |
| GO_APOPTOTIC_SIGNALING_PATHWAY | DAPK1 |
| GO_APOPTOTIC_SIGNALING_PATHWAY | DAPK3 |
| GO_APOPTOTIC_SIGNALING_PATHWAY | DAPL1 |
| GO_APOPTOTIC_SIGNALING_PATHWAY | DAXX |
| GO_APOPTOTIC_SIGNALING_PATHWAY | DCC |

| | |
|---|---|
| GO_APOPTOTIC_SIGNALING_PATHWAY | DDIT3 |
| GO_APOPTOTIC_SIGNALING_PATHWAY | DDIT4 |
| GO_APOPTOTIC_SIGNALING_PATHWAY | DDX3X |
| GO_APOPTOTIC_SIGNALING_PATHWAY | DDX47 |
| GO_APOPTOTIC_SIGNALING_PATHWAY | DDX5 |
| GO_APOPTOTIC_SIGNALING_PATHWAY | DEDD |
| GO_APOPTOTIC_SIGNALING_PATHWAY | DEDD2 |
| GO_APOPTOTIC_SIGNALING_PATHWAY | DIABLO |
| GO_APOPTOTIC_SIGNALING_PATHWAY | DIDO1 |
| GO_APOPTOTIC_SIGNALING_PATHWAY | DNAJC10 |
| GO_APOPTOTIC_SIGNALING_PATHWAY | DNM1L |
| GO_APOPTOTIC_SIGNALING_PATHWAY | DPF2 |
| GO_APOPTOTIC_SIGNALING_PATHWAY | DYRK2 |
| GO_APOPTOTIC_SIGNALING_PATHWAY | E2F1 |
| GO_APOPTOTIC_SIGNALING_PATHWAY | E2F2 |
| GO_APOPTOTIC_SIGNALING_PATHWAY | EDA2R |
| GO_APOPTOTIC_SIGNALING_PATHWAY | EP300 |
| GO_APOPTOTIC_SIGNALING_PATHWAY | EPHA2 |
| GO_APOPTOTIC_SIGNALING_PATHWAY | ERBB3 |
| GO_APOPTOTIC_SIGNALING_PATHWAY | ERCC6 |
| GO_APOPTOTIC_SIGNALING_PATHWAY | ERN1 |
| GO_APOPTOTIC_SIGNALING_PATHWAY | ERO1L |
| GO_APOPTOTIC_SIGNALING_PATHWAY | EYA2 |
| GO_APOPTOTIC_SIGNALING_PATHWAY | FADD |
| GO_APOPTOTIC_SIGNALING_PATHWAY | FAS |
| GO_APOPTOTIC_SIGNALING_PATHWAY | FASLG |
| GO_APOPTOTIC_SIGNALING_PATHWAY | FASTK |
| GO_APOPTOTIC_SIGNALING_PATHWAY | FGFR3 |
| GO_APOPTOTIC_SIGNALING_PATHWAY | FHIT |
| GO_APOPTOTIC_SIGNALING_PATHWAY | FIS1 |
| GO_APOPTOTIC_SIGNALING_PATHWAY | FNIP2 |
| GO_APOPTOTIC_SIGNALING_PATHWAY | FOXO3 |
| GO_APOPTOTIC_SIGNALING_PATHWAY | G0S2 |
| GO_APOPTOTIC_SIGNALING_PATHWAY | GABARAP |
| GO_APOPTOTIC_SIGNALING_PATHWAY | GGCT |
| GO_APOPTOTIC_SIGNALING_PATHWAY | GPX1 |
| GO_APOPTOTIC_SIGNALING_PATHWAY | GSK3A |
| GO_APOPTOTIC_SIGNALING_PATHWAY | GSK3B |
| GO_APOPTOTIC_SIGNALING_PATHWAY | HIC1 |
| GO_APOPTOTIC_SIGNALING_PATHWAY | HINT1 |
| GO_APOPTOTIC_SIGNALING_PATHWAY | HIP1 |
| GO_APOPTOTIC_SIGNALING_PATHWAY | HIPK1 |
| GO_APOPTOTIC_SIGNALING_PATHWAY | HIPK2 |
| GO_APOPTOTIC_SIGNALING_PATHWAY | HMOX1 |
| GO_APOPTOTIC_SIGNALING_PATHWAY | HRAS |
| GO_APOPTOTIC_SIGNALING_PATHWAY | HTRA2 |
| GO_APOPTOTIC_SIGNALING_PATHWAY | IFI16 |

| | |
|---|---|
| GO_APOPTOTIC_SIGNALING_PATHWAY | IFI27 |
| GO_APOPTOTIC_SIGNALING_PATHWAY | IFI6 |
| GO_APOPTOTIC_SIGNALING_PATHWAY | IFNG |
| GO_APOPTOTIC_SIGNALING_PATHWAY | IKBKE |
| GO_APOPTOTIC_SIGNALING_PATHWAY | IL12A |
| GO_APOPTOTIC_SIGNALING_PATHWAY | IL1A |
| GO_APOPTOTIC_SIGNALING_PATHWAY | IL1B |
| GO_APOPTOTIC_SIGNALING_PATHWAY | IL2 |
| GO_APOPTOTIC_SIGNALING_PATHWAY | IL33 |
| GO_APOPTOTIC_SIGNALING_PATHWAY | IL4 |
| GO_APOPTOTIC_SIGNALING_PATHWAY | IL6R |
| GO_APOPTOTIC_SIGNALING_PATHWAY | INHBA |
| GO_APOPTOTIC_SIGNALING_PATHWAY | ITGAV |
| GO_APOPTOTIC_SIGNALING_PATHWAY | ITPR1 |
| GO_APOPTOTIC_SIGNALING_PATHWAY | JAK2 |
| GO_APOPTOTIC_SIGNALING_PATHWAY | JUN |
| GO_APOPTOTIC_SIGNALING_PATHWAY | KIAA0141 |
| GO_APOPTOTIC_SIGNALING_PATHWAY | KITLG |
| GO_APOPTOTIC_SIGNALING_PATHWAY | KRT18 |
| GO_APOPTOTIC_SIGNALING_PATHWAY | KRT8 |
| GO_APOPTOTIC_SIGNALING_PATHWAY | LAMP1 |
| GO_APOPTOTIC_SIGNALING_PATHWAY | LGALS12 |
| GO_APOPTOTIC_SIGNALING_PATHWAY | LY96 |
| GO_APOPTOTIC_SIGNALING_PATHWAY | MAEL |
| GO_APOPTOTIC_SIGNALING_PATHWAY | MAP3K5 |
| GO_APOPTOTIC_SIGNALING_PATHWAY | MAPK9 |
| GO_APOPTOTIC_SIGNALING_PATHWAY | MAPT |
| GO_APOPTOTIC_SIGNALING_PATHWAY | MCL1 |
| GO_APOPTOTIC_SIGNALING_PATHWAY | MELK |
| GO_APOPTOTIC_SIGNALING_PATHWAY | MFF |
| GO_APOPTOTIC_SIGNALING_PATHWAY | MKNK2 |
| GO_APOPTOTIC_SIGNALING_PATHWAY | MLH1 |
| GO_APOPTOTIC_SIGNALING_PATHWAY | MLLT11 |
| GO_APOPTOTIC_SIGNALING_PATHWAY | MOAP1 |
| GO_APOPTOTIC_SIGNALING_PATHWAY | MSH2 |
| GO_APOPTOTIC_SIGNALING_PATHWAY | MSH6 |
| GO_APOPTOTIC_SIGNALING_PATHWAY | MYBBP1A |
| GO_APOPTOTIC_SIGNALING_PATHWAY | NBN |
| GO_APOPTOTIC_SIGNALING_PATHWAY | NDUFA13 |
| GO_APOPTOTIC_SIGNALING_PATHWAY | NF1 |
| GO_APOPTOTIC_SIGNALING_PATHWAY | NFATC4 |
| GO_APOPTOTIC_SIGNALING_PATHWAY | NGF |
| GO_APOPTOTIC_SIGNALING_PATHWAY | NGFR |
| GO_APOPTOTIC_SIGNALING_PATHWAY | NGFRAP1 |
| GO_APOPTOTIC_SIGNALING_PATHWAY | NOL3 |
| GO_APOPTOTIC_SIGNALING_PATHWAY | NUPR1 |
| GO_APOPTOTIC_SIGNALING_PATHWAY | P2RX4 |

| | |
|---|---|
| GO_APOPTOTIC_SIGNALING_PATHWAY | P2RX7 |
| GO_APOPTOTIC_SIGNALING_PATHWAY | PARP2 |
| GO_APOPTOTIC_SIGNALING_PATHWAY | PAWR |
| GO_APOPTOTIC_SIGNALING_PATHWAY | PDCD10 |
| GO_APOPTOTIC_SIGNALING_PATHWAY | PDCD6 |
| GO_APOPTOTIC_SIGNALING_PATHWAY | PDK1 |
| GO_APOPTOTIC_SIGNALING_PATHWAY | PDK2 |
| GO_APOPTOTIC_SIGNALING_PATHWAY | PDPK1 |
| GO_APOPTOTIC_SIGNALING_PATHWAY | PERP |
| GO_APOPTOTIC_SIGNALING_PATHWAY | PGAP2 |
| GO_APOPTOTIC_SIGNALING_PATHWAY | PHLDA3 |
| GO_APOPTOTIC_SIGNALING_PATHWAY | PIK3R1 |
| GO_APOPTOTIC_SIGNALING_PATHWAY | PMAIP1 |
| GO_APOPTOTIC_SIGNALING_PATHWAY | PML |
| GO_APOPTOTIC_SIGNALING_PATHWAY | POLB |
| GO_APOPTOTIC_SIGNALING_PATHWAY | PPARD |
| GO_APOPTOTIC_SIGNALING_PATHWAY | PPM1F |
| GO_APOPTOTIC_SIGNALING_PATHWAY | PPP1R13B |
| GO_APOPTOTIC_SIGNALING_PATHWAY | PPP1R15A |
| GO_APOPTOTIC_SIGNALING_PATHWAY | PPP2R5C |
| GO_APOPTOTIC_SIGNALING_PATHWAY | PRKCA |
| GO_APOPTOTIC_SIGNALING_PATHWAY | PRKCD |
| GO_APOPTOTIC_SIGNALING_PATHWAY | PRKDC |
| GO_APOPTOTIC_SIGNALING_PATHWAY | PRODH |
| GO_APOPTOTIC_SIGNALING_PATHWAY | PTGIS |
| GO_APOPTOTIC_SIGNALING_PATHWAY | PTH |
| GO_APOPTOTIC_SIGNALING_PATHWAY | PYCARD |
| GO_APOPTOTIC_SIGNALING_PATHWAY | RAF1 |
| GO_APOPTOTIC_SIGNALING_PATHWAY | RELT |
| GO_APOPTOTIC_SIGNALING_PATHWAY | RHOT1 |
| GO_APOPTOTIC_SIGNALING_PATHWAY | RHOT2 |
| GO_APOPTOTIC_SIGNALING_PATHWAY | RIPK1 |
| GO_APOPTOTIC_SIGNALING_PATHWAY | RIPK3 |
| GO_APOPTOTIC_SIGNALING_PATHWAY | RNF41 |
| GO_APOPTOTIC_SIGNALING_PATHWAY | RPS27L |
| GO_APOPTOTIC_SIGNALING_PATHWAY | RRP8 |
| GO_APOPTOTIC_SIGNALING_PATHWAY | SART1 |
| GO_APOPTOTIC_SIGNALING_PATHWAY | SCN2A |
| GO_APOPTOTIC_SIGNALING_PATHWAY | SELK |
| GO_APOPTOTIC_SIGNALING_PATHWAY | SENP1 |
| GO_APOPTOTIC_SIGNALING_PATHWAY | SFN |
| GO_APOPTOTIC_SIGNALING_PATHWAY | SGPL1 |
| GO_APOPTOTIC_SIGNALING_PATHWAY | SGPP1 |
| GO_APOPTOTIC_SIGNALING_PATHWAY | SHH |
| GO_APOPTOTIC_SIGNALING_PATHWAY | SHISA5 |
| GO_APOPTOTIC_SIGNALING_PATHWAY | SIRT1 |
| GO_APOPTOTIC_SIGNALING_PATHWAY | SIVA1 |

| | |
|---|---|
| GO_APOPTOTIC_SIGNALING_PATHWAY | SMAD3 |
| GO_APOPTOTIC_SIGNALING_PATHWAY | SNW1 |
| GO_APOPTOTIC_SIGNALING_PATHWAY | SOD2 |
| GO_APOPTOTIC_SIGNALING_PATHWAY | SORT1 |
| GO_APOPTOTIC_SIGNALING_PATHWAY | SPN |
| GO_APOPTOTIC_SIGNALING_PATHWAY | SRGN |
| GO_APOPTOTIC_SIGNALING_PATHWAY | SST |
| GO_APOPTOTIC_SIGNALING_PATHWAY | SSTR3 |
| GO_APOPTOTIC_SIGNALING_PATHWAY | ST20 |
| GO_APOPTOTIC_SIGNALING_PATHWAY | STK11 |
| GO_APOPTOTIC_SIGNALING_PATHWAY | STK24 |
| GO_APOPTOTIC_SIGNALING_PATHWAY | STK25 |
| GO_APOPTOTIC_SIGNALING_PATHWAY | TFPT |
| GO_APOPTOTIC_SIGNALING_PATHWAY | TGFB1 |
| GO_APOPTOTIC_SIGNALING_PATHWAY | TGFB2 |
| GO_APOPTOTIC_SIGNALING_PATHWAY | TICAM1 |
| GO_APOPTOTIC_SIGNALING_PATHWAY | TICAM2 |
| GO_APOPTOTIC_SIGNALING_PATHWAY | TIMM50 |
| GO_APOPTOTIC_SIGNALING_PATHWAY | TLR3 |
| GO_APOPTOTIC_SIGNALING_PATHWAY | TLR4 |
| GO_APOPTOTIC_SIGNALING_PATHWAY | TM2D1 |
| GO_APOPTOTIC_SIGNALING_PATHWAY | TMBIM6 |
| GO_APOPTOTIC_SIGNALING_PATHWAY | TMEM109 |
| GO_APOPTOTIC_SIGNALING_PATHWAY | TNF |
| GO_APOPTOTIC_SIGNALING_PATHWAY | TNFRSF10A |
| GO_APOPTOTIC_SIGNALING_PATHWAY | TNFRSF10B |
| GO_APOPTOTIC_SIGNALING_PATHWAY | TNFRSF10C |
| GO_APOPTOTIC_SIGNALING_PATHWAY | TNFRSF10D |
| GO_APOPTOTIC_SIGNALING_PATHWAY | TNFRSF11A |
| GO_APOPTOTIC_SIGNALING_PATHWAY | TNFRSF11B |
| GO_APOPTOTIC_SIGNALING_PATHWAY | TNFRSF12A |
| GO_APOPTOTIC_SIGNALING_PATHWAY | TNFRSF14 |
| GO_APOPTOTIC_SIGNALING_PATHWAY | TNFRSF18 |
| GO_APOPTOTIC_SIGNALING_PATHWAY | TNFRSF1A |
| GO_APOPTOTIC_SIGNALING_PATHWAY | TNFRSF1B |
| GO_APOPTOTIC_SIGNALING_PATHWAY | TNFRSF21 |
| GO_APOPTOTIC_SIGNALING_PATHWAY | TNFRSF25 |
| GO_APOPTOTIC_SIGNALING_PATHWAY | TNFRSF4 |
| GO_APOPTOTIC_SIGNALING_PATHWAY | TNFRSF6B |
| GO_APOPTOTIC_SIGNALING_PATHWAY | TNFRSF8 |
| GO_APOPTOTIC_SIGNALING_PATHWAY | TNFRSF9 |
| GO_APOPTOTIC_SIGNALING_PATHWAY | TNFSF10 |
| GO_APOPTOTIC_SIGNALING_PATHWAY | TNFSF12 |
| GO_APOPTOTIC_SIGNALING_PATHWAY | TOPORS |
| GO_APOPTOTIC_SIGNALING_PATHWAY | TP53 |
| GO_APOPTOTIC_SIGNALING_PATHWAY | TP53BP2 |
| GO_APOPTOTIC_SIGNALING_PATHWAY | TP63 |

| | |
|---|---|
| GO_APOPTOTIC_SIGNALING_PATHWAY | TP73 |
| GO_APOPTOTIC_SIGNALING_PATHWAY | TRADD |
| GO_APOPTOTIC_SIGNALING_PATHWAY | TRAF2 |
| GO_APOPTOTIC_SIGNALING_PATHWAY | TRIB3 |
| GO_APOPTOTIC_SIGNALING_PATHWAY | UACA |
| GO_APOPTOTIC_SIGNALING_PATHWAY | UBE2K |
| GO_APOPTOTIC_SIGNALING_PATHWAY | UBE4B |
| GO_APOPTOTIC_SIGNALING_PATHWAY | USP28 |
| GO_APOPTOTIC_SIGNALING_PATHWAY | WWOX |
| GO_APOPTOTIC_SIGNALING_PATHWAY | XBP1 |
| GO_APOPTOTIC_SIGNALING_PATHWAY | XPA |
| GO_APOPTOTIC_SIGNALING_PATHWAY | ZMAT1 |
| GO_APOPTOTIC_SIGNALING_PATHWAY | ZMAT3 |
| GO_APOPTOTIC_SIGNALING_PATHWAY | ZMAT4 |
| GO_APOPTOTIC_SIGNALING_PATHWAY | ZNF346 |
| GO_APOPTOTIC_SIGNALING_PATHWAY | ZNF385B |
| GO_APOPTOTIC_SIGNALING_PATHWAY | ZNF385C |
| GO_APOPTOTIC_SIGNALING_PATHWAY | ZNF385D |
| GO_APOPTOTIC_SIGNALING_PATHWAY | ZNF622 |
| GO_CELLULAR_RESPONSE_TO_DNA_DAMAGE_STIMULUS | AATF |
| GO_CELLULAR_RESPONSE_TO_DNA_DAMAGE_STIMULUS | ABL1 |
| GO_CELLULAR_RESPONSE_TO_DNA_DAMAGE_STIMULUS | ACD |
| GO_CELLULAR_RESPONSE_TO_DNA_DAMAGE_STIMULUS | ACTL6A |
| GO_CELLULAR_RESPONSE_TO_DNA_DAMAGE_STIMULUS | ACTR5 |
| GO_CELLULAR_RESPONSE_TO_DNA_DAMAGE_STIMULUS | ACTR8 |
| GO_CELLULAR_RESPONSE_TO_DNA_DAMAGE_STIMULUS | AEN |
| GO_CELLULAR_RESPONSE_TO_DNA_DAMAGE_STIMULUS | AKT1 |
| GO_CELLULAR_RESPONSE_TO_DNA_DAMAGE_STIMULUS | ALKBH1 |
| GO_CELLULAR_RESPONSE_TO_DNA_DAMAGE_STIMULUS | ALKBH2 |
| GO_CELLULAR_RESPONSE_TO_DNA_DAMAGE_STIMULUS | ALKBH3 |
| GO_CELLULAR_RESPONSE_TO_DNA_DAMAGE_STIMULUS | ALKBH7 |
| GO_CELLULAR_RESPONSE_TO_DNA_DAMAGE_STIMULUS | ALKBH8 |
| GO_CELLULAR_RESPONSE_TO_DNA_DAMAGE_STIMULUS | ANKRD32 |
| GO_CELLULAR_RESPONSE_TO_DNA_DAMAGE_STIMULUS | APBB1 |
| GO_CELLULAR_RESPONSE_TO_DNA_DAMAGE_STIMULUS | APC |
| GO_CELLULAR_RESPONSE_TO_DNA_DAMAGE_STIMULUS | APEX1 |
| GO_CELLULAR_RESPONSE_TO_DNA_DAMAGE_STIMULUS | APEX2 |
| GO_CELLULAR_RESPONSE_TO_DNA_DAMAGE_STIMULUS | APITD1 |
| GO_CELLULAR_RESPONSE_TO_DNA_DAMAGE_STIMULUS | APLF |
| GO_CELLULAR_RESPONSE_TO_DNA_DAMAGE_STIMULUS | APTX |
| GO_CELLULAR_RESPONSE_TO_DNA_DAMAGE_STIMULUS | AQR |
| GO_CELLULAR_RESPONSE_TO_DNA_DAMAGE_STIMULUS | ARID3A |
| GO_CELLULAR_RESPONSE_TO_DNA_DAMAGE_STIMULUS | ASCC1 |
| GO_CELLULAR_RESPONSE_TO_DNA_DAMAGE_STIMULUS | ASCC2 |
| GO_CELLULAR_RESPONSE_TO_DNA_DAMAGE_STIMULUS | ASCC3 |
| GO_CELLULAR_RESPONSE_TO_DNA_DAMAGE_STIMULUS | ASF1A |
| GO_CELLULAR_RESPONSE_TO_DNA_DAMAGE_STIMULUS | ASH2L |

| | |
|---|---|
| GO_CELLULAR_RESPONSE_TO_DNA_DAMAGE_STIMULUS | ASTE1 |
| GO_CELLULAR_RESPONSE_TO_DNA_DAMAGE_STIMULUS | ATAD5 |
| GO_CELLULAR_RESPONSE_TO_DNA_DAMAGE_STIMULUS | ATF2 |
| GO_CELLULAR_RESPONSE_TO_DNA_DAMAGE_STIMULUS | ATM |
| GO_CELLULAR_RESPONSE_TO_DNA_DAMAGE_STIMULUS | ATMIN |
| GO_CELLULAR_RESPONSE_TO_DNA_DAMAGE_STIMULUS | ATR |
| GO_CELLULAR_RESPONSE_TO_DNA_DAMAGE_STIMULUS | ATRIP |
| GO_CELLULAR_RESPONSE_TO_DNA_DAMAGE_STIMULUS | ATRX |
| GO_CELLULAR_RESPONSE_TO_DNA_DAMAGE_STIMULUS | ATXN3 |
| GO_CELLULAR_RESPONSE_TO_DNA_DAMAGE_STIMULUS | AURKA |
| GO_CELLULAR_RESPONSE_TO_DNA_DAMAGE_STIMULUS | BABAM1 |
| GO_CELLULAR_RESPONSE_TO_DNA_DAMAGE_STIMULUS | BACH1 |
| GO_CELLULAR_RESPONSE_TO_DNA_DAMAGE_STIMULUS | BAD |
| GO_CELLULAR_RESPONSE_TO_DNA_DAMAGE_STIMULUS | BAG6 |
| GO_CELLULAR_RESPONSE_TO_DNA_DAMAGE_STIMULUS | BAK1 |
| GO_CELLULAR_RESPONSE_TO_DNA_DAMAGE_STIMULUS | BARD1 |
| GO_CELLULAR_RESPONSE_TO_DNA_DAMAGE_STIMULUS | BATF |
| GO_CELLULAR_RESPONSE_TO_DNA_DAMAGE_STIMULUS | BAX |
| GO_CELLULAR_RESPONSE_TO_DNA_DAMAGE_STIMULUS | BAZ1B |
| GO_CELLULAR_RESPONSE_TO_DNA_DAMAGE_STIMULUS | BBC3 |
| GO_CELLULAR_RESPONSE_TO_DNA_DAMAGE_STIMULUS | BCCIP |
| GO_CELLULAR_RESPONSE_TO_DNA_DAMAGE_STIMULUS | BCL2 |
| GO_CELLULAR_RESPONSE_TO_DNA_DAMAGE_STIMULUS | BCL2A1 |
| GO_CELLULAR_RESPONSE_TO_DNA_DAMAGE_STIMULUS | BCL2L1 |
| GO_CELLULAR_RESPONSE_TO_DNA_DAMAGE_STIMULUS | BCL2L10 |
| GO_CELLULAR_RESPONSE_TO_DNA_DAMAGE_STIMULUS | BCL2L11 |
| GO_CELLULAR_RESPONSE_TO_DNA_DAMAGE_STIMULUS | BCL2L2 |
| GO_CELLULAR_RESPONSE_TO_DNA_DAMAGE_STIMULUS | BCL3 |
| GO_CELLULAR_RESPONSE_TO_DNA_DAMAGE_STIMULUS | BCL6 |
| GO_CELLULAR_RESPONSE_TO_DNA_DAMAGE_STIMULUS | BID |
| GO_CELLULAR_RESPONSE_TO_DNA_DAMAGE_STIMULUS | BLM |
| GO_CELLULAR_RESPONSE_TO_DNA_DAMAGE_STIMULUS | BOD1L |
| GO_CELLULAR_RESPONSE_TO_DNA_DAMAGE_STIMULUS | BOK |
| GO_CELLULAR_RESPONSE_TO_DNA_DAMAGE_STIMULUS | BRAT1 |
| GO_CELLULAR_RESPONSE_TO_DNA_DAMAGE_STIMULUS | BRCA1 |
| GO_CELLULAR_RESPONSE_TO_DNA_DAMAGE_STIMULUS | BRCA2 |
| GO_CELLULAR_RESPONSE_TO_DNA_DAMAGE_STIMULUS | BRCC3 |
| GO_CELLULAR_RESPONSE_TO_DNA_DAMAGE_STIMULUS | BRD4 |
| GO_CELLULAR_RESPONSE_TO_DNA_DAMAGE_STIMULUS | BRE |
| GO_CELLULAR_RESPONSE_TO_DNA_DAMAGE_STIMULUS | BRIP1 |
| GO_CELLULAR_RESPONSE_TO_DNA_DAMAGE_STIMULUS | BRSK1 |
| GO_CELLULAR_RESPONSE_TO_DNA_DAMAGE_STIMULUS | BTG2 |
| GO_CELLULAR_RESPONSE_TO_DNA_DAMAGE_STIMULUS | C11orf30 |
| GO_CELLULAR_RESPONSE_TO_DNA_DAMAGE_STIMULUS | C12orf32 |
| GO_CELLULAR_RESPONSE_TO_DNA_DAMAGE_STIMULUS | C12orf48 |
| GO_CELLULAR_RESPONSE_TO_DNA_DAMAGE_STIMULUS | C12orf5 |
| GO_CELLULAR_RESPONSE_TO_DNA_DAMAGE_STIMULUS | C13orf15 |

| | |
|---|---|
| GO_CELLULAR_RESPONSE_TO_DNA_DAMAGE_STIMULUS | C15orf42 |
| GO_CELLULAR_RESPONSE_TO_DNA_DAMAGE_STIMULUS | C16orf5 |
| GO_CELLULAR_RESPONSE_TO_DNA_DAMAGE_STIMULUS | C16orf53 |
| GO_CELLULAR_RESPONSE_TO_DNA_DAMAGE_STIMULUS | C16orf73 |
| GO_CELLULAR_RESPONSE_TO_DNA_DAMAGE_STIMULUS | C17orf70 |
| GO_CELLULAR_RESPONSE_TO_DNA_DAMAGE_STIMULUS | C19orf39 |
| GO_CELLULAR_RESPONSE_TO_DNA_DAMAGE_STIMULUS | C19orf40 |
| GO_CELLULAR_RESPONSE_TO_DNA_DAMAGE_STIMULUS | C1orf124 |
| GO_CELLULAR_RESPONSE_TO_DNA_DAMAGE_STIMULUS | C1orf86 |
| GO_CELLULAR_RESPONSE_TO_DNA_DAMAGE_STIMULUS | C20orf29 |
| GO_CELLULAR_RESPONSE_TO_DNA_DAMAGE_STIMULUS | C20orf72 |
| GO_CELLULAR_RESPONSE_TO_DNA_DAMAGE_STIMULUS | C2orf29 |
| GO_CELLULAR_RESPONSE_TO_DNA_DAMAGE_STIMULUS | C6orf211 |
| GO_CELLULAR_RESPONSE_TO_DNA_DAMAGE_STIMULUS | C8orf80 |
| GO_CELLULAR_RESPONSE_TO_DNA_DAMAGE_STIMULUS | C9orf102 |
| GO_CELLULAR_RESPONSE_TO_DNA_DAMAGE_STIMULUS | C9orf142 |
| GO_CELLULAR_RESPONSE_TO_DNA_DAMAGE_STIMULUS | C9orf80 |
| GO_CELLULAR_RESPONSE_TO_DNA_DAMAGE_STIMULUS | CARM1 |
| GO_CELLULAR_RESPONSE_TO_DNA_DAMAGE_STIMULUS | CASP2 |
| GO_CELLULAR_RESPONSE_TO_DNA_DAMAGE_STIMULUS | CASP3 |
| GO_CELLULAR_RESPONSE_TO_DNA_DAMAGE_STIMULUS | CASP9 |
| GO_CELLULAR_RESPONSE_TO_DNA_DAMAGE_STIMULUS | CCDC111 |
| GO_CELLULAR_RESPONSE_TO_DNA_DAMAGE_STIMULUS | CCDC13 |
| GO_CELLULAR_RESPONSE_TO_DNA_DAMAGE_STIMULUS | CCDC155 |
| GO_CELLULAR_RESPONSE_TO_DNA_DAMAGE_STIMULUS | CCNA2 |
| GO_CELLULAR_RESPONSE_TO_DNA_DAMAGE_STIMULUS | CCNB1 |
| GO_CELLULAR_RESPONSE_TO_DNA_DAMAGE_STIMULUS | CCND1 |
| GO_CELLULAR_RESPONSE_TO_DNA_DAMAGE_STIMULUS | CCNH |
| GO_CELLULAR_RESPONSE_TO_DNA_DAMAGE_STIMULUS | CCNK |
| GO_CELLULAR_RESPONSE_TO_DNA_DAMAGE_STIMULUS | CCNO |
| GO_CELLULAR_RESPONSE_TO_DNA_DAMAGE_STIMULUS | CDC14B |
| GO_CELLULAR_RESPONSE_TO_DNA_DAMAGE_STIMULUS | CDC25C |
| GO_CELLULAR_RESPONSE_TO_DNA_DAMAGE_STIMULUS | CDC45 |
| GO_CELLULAR_RESPONSE_TO_DNA_DAMAGE_STIMULUS | CDC5L |
| GO_CELLULAR_RESPONSE_TO_DNA_DAMAGE_STIMULUS | CDC7 |
| GO_CELLULAR_RESPONSE_TO_DNA_DAMAGE_STIMULUS | CDCA5 |
| GO_CELLULAR_RESPONSE_TO_DNA_DAMAGE_STIMULUS | CDK1 |
| GO_CELLULAR_RESPONSE_TO_DNA_DAMAGE_STIMULUS | CDK2 |
| GO_CELLULAR_RESPONSE_TO_DNA_DAMAGE_STIMULUS | CDK3 |
| GO_CELLULAR_RESPONSE_TO_DNA_DAMAGE_STIMULUS | CDK5RAP3 |
| GO_CELLULAR_RESPONSE_TO_DNA_DAMAGE_STIMULUS | CDK7 |
| GO_CELLULAR_RESPONSE_TO_DNA_DAMAGE_STIMULUS | CDK9 |
| GO_CELLULAR_RESPONSE_TO_DNA_DAMAGE_STIMULUS | CDKN1A |
| GO_CELLULAR_RESPONSE_TO_DNA_DAMAGE_STIMULUS | CDKN1B |
| GO_CELLULAR_RESPONSE_TO_DNA_DAMAGE_STIMULUS | CDKN2AIP |
| GO_CELLULAR_RESPONSE_TO_DNA_DAMAGE_STIMULUS | CDKN2D |
| GO_CELLULAR_RESPONSE_TO_DNA_DAMAGE_STIMULUS | CENPJ |

| | |
|---|---|
| GO_CELLULAR_RESPONSE_TO_DNA_DAMAGE_STIMULUS | CEP164 |
| GO_CELLULAR_RESPONSE_TO_DNA_DAMAGE_STIMULUS | CEP63 |
| GO_CELLULAR_RESPONSE_TO_DNA_DAMAGE_STIMULUS | CETN2 |
| GO_CELLULAR_RESPONSE_TO_DNA_DAMAGE_STIMULUS | CHAF1A |
| GO_CELLULAR_RESPONSE_TO_DNA_DAMAGE_STIMULUS | CHAF1B |
| GO_CELLULAR_RESPONSE_TO_DNA_DAMAGE_STIMULUS | CHCHD6 |
| GO_CELLULAR_RESPONSE_TO_DNA_DAMAGE_STIMULUS | CHD1L |
| GO_CELLULAR_RESPONSE_TO_DNA_DAMAGE_STIMULUS | CHD2 |
| GO_CELLULAR_RESPONSE_TO_DNA_DAMAGE_STIMULUS | CHEK1 |
| GO_CELLULAR_RESPONSE_TO_DNA_DAMAGE_STIMULUS | CHEK2 |
| GO_CELLULAR_RESPONSE_TO_DNA_DAMAGE_STIMULUS | CHRNA4 |
| GO_CELLULAR_RESPONSE_TO_DNA_DAMAGE_STIMULUS | CIB1 |
| GO_CELLULAR_RESPONSE_TO_DNA_DAMAGE_STIMULUS | CIDEB |
| GO_CELLULAR_RESPONSE_TO_DNA_DAMAGE_STIMULUS | CINP |
| GO_CELLULAR_RESPONSE_TO_DNA_DAMAGE_STIMULUS | CLOCK |
| GO_CELLULAR_RESPONSE_TO_DNA_DAMAGE_STIMULUS | CLSPN |
| GO_CELLULAR_RESPONSE_TO_DNA_DAMAGE_STIMULUS | CNOT1 |
| GO_CELLULAR_RESPONSE_TO_DNA_DAMAGE_STIMULUS | CNOT10 |
| GO_CELLULAR_RESPONSE_TO_DNA_DAMAGE_STIMULUS | CNOT2 |
| GO_CELLULAR_RESPONSE_TO_DNA_DAMAGE_STIMULUS | CNOT3 |
| GO_CELLULAR_RESPONSE_TO_DNA_DAMAGE_STIMULUS | CNOT4 |
| GO_CELLULAR_RESPONSE_TO_DNA_DAMAGE_STIMULUS | CNOT6 |
| GO_CELLULAR_RESPONSE_TO_DNA_DAMAGE_STIMULUS | CNOT6L |
| GO_CELLULAR_RESPONSE_TO_DNA_DAMAGE_STIMULUS | CNOT7 |
| GO_CELLULAR_RESPONSE_TO_DNA_DAMAGE_STIMULUS | CNOT8 |
| GO_CELLULAR_RESPONSE_TO_DNA_DAMAGE_STIMULUS | COPS2 |
| GO_CELLULAR_RESPONSE_TO_DNA_DAMAGE_STIMULUS | COPS3 |
| GO_CELLULAR_RESPONSE_TO_DNA_DAMAGE_STIMULUS | COPS4 |
| GO_CELLULAR_RESPONSE_TO_DNA_DAMAGE_STIMULUS | COPS5 |
| GO_CELLULAR_RESPONSE_TO_DNA_DAMAGE_STIMULUS | COPS6 |
| GO_CELLULAR_RESPONSE_TO_DNA_DAMAGE_STIMULUS | COPS7A |
| GO_CELLULAR_RESPONSE_TO_DNA_DAMAGE_STIMULUS | COPS7B |
| GO_CELLULAR_RESPONSE_TO_DNA_DAMAGE_STIMULUS | COPS8 |
| GO_CELLULAR_RESPONSE_TO_DNA_DAMAGE_STIMULUS | CRADD |
| GO_CELLULAR_RESPONSE_TO_DNA_DAMAGE_STIMULUS | CRIP1 |
| GO_CELLULAR_RESPONSE_TO_DNA_DAMAGE_STIMULUS | CRY1 |
| GO_CELLULAR_RESPONSE_TO_DNA_DAMAGE_STIMULUS | CRY2 |
| GO_CELLULAR_RESPONSE_TO_DNA_DAMAGE_STIMULUS | CSNK1D |
| GO_CELLULAR_RESPONSE_TO_DNA_DAMAGE_STIMULUS | CSNK1E |
| GO_CELLULAR_RESPONSE_TO_DNA_DAMAGE_STIMULUS | CTC1 |
| GO_CELLULAR_RESPONSE_TO_DNA_DAMAGE_STIMULUS | CTLA4 |
| GO_CELLULAR_RESPONSE_TO_DNA_DAMAGE_STIMULUS | CUL4A |
| GO_CELLULAR_RESPONSE_TO_DNA_DAMAGE_STIMULUS | CUL4B |
| GO_CELLULAR_RESPONSE_TO_DNA_DAMAGE_STIMULUS | DCLRE1A |
| GO_CELLULAR_RESPONSE_TO_DNA_DAMAGE_STIMULUS | DCLRE1B |
| GO_CELLULAR_RESPONSE_TO_DNA_DAMAGE_STIMULUS | DCLRE1C |
| GO_CELLULAR_RESPONSE_TO_DNA_DAMAGE_STIMULUS | DDB1 |

| | |
|---|---|
| GO_CELLULAR_RESPONSE_TO_DNA_DAMAGE_STIMULUS | DDB2 |
| GO_CELLULAR_RESPONSE_TO_DNA_DAMAGE_STIMULUS | DDIT3 |
| GO_CELLULAR_RESPONSE_TO_DNA_DAMAGE_STIMULUS | DDIT4 |
| GO_CELLULAR_RESPONSE_TO_DNA_DAMAGE_STIMULUS | DDX1 |
| GO_CELLULAR_RESPONSE_TO_DNA_DAMAGE_STIMULUS | DDX39A |
| GO_CELLULAR_RESPONSE_TO_DNA_DAMAGE_STIMULUS | DEM1 |
| GO_CELLULAR_RESPONSE_TO_DNA_DAMAGE_STIMULUS | DGKZ |
| GO_CELLULAR_RESPONSE_TO_DNA_DAMAGE_STIMULUS | DMAP1 |
| GO_CELLULAR_RESPONSE_TO_DNA_DAMAGE_STIMULUS | DMC1 |
| GO_CELLULAR_RESPONSE_TO_DNA_DAMAGE_STIMULUS | DNA2 |
| GO_CELLULAR_RESPONSE_TO_DNA_DAMAGE_STIMULUS | DNAJA1 |
| GO_CELLULAR_RESPONSE_TO_DNA_DAMAGE_STIMULUS | DTL |
| GO_CELLULAR_RESPONSE_TO_DNA_DAMAGE_STIMULUS | DTX3L |
| GO_CELLULAR_RESPONSE_TO_DNA_DAMAGE_STIMULUS | DYRK2 |
| GO_CELLULAR_RESPONSE_TO_DNA_DAMAGE_STIMULUS | E2F1 |
| GO_CELLULAR_RESPONSE_TO_DNA_DAMAGE_STIMULUS | E2F4 |
| GO_CELLULAR_RESPONSE_TO_DNA_DAMAGE_STIMULUS | E2F7 |
| GO_CELLULAR_RESPONSE_TO_DNA_DAMAGE_STIMULUS | EEPD1 |
| GO_CELLULAR_RESPONSE_TO_DNA_DAMAGE_STIMULUS | EGLN3 |
| GO_CELLULAR_RESPONSE_TO_DNA_DAMAGE_STIMULUS | EID3 |
| GO_CELLULAR_RESPONSE_TO_DNA_DAMAGE_STIMULUS | EME1 |
| GO_CELLULAR_RESPONSE_TO_DNA_DAMAGE_STIMULUS | EME2 |
| GO_CELLULAR_RESPONSE_TO_DNA_DAMAGE_STIMULUS | ENDOV |
| GO_CELLULAR_RESPONSE_TO_DNA_DAMAGE_STIMULUS | EP300 |
| GO_CELLULAR_RESPONSE_TO_DNA_DAMAGE_STIMULUS | EPC2 |
| GO_CELLULAR_RESPONSE_TO_DNA_DAMAGE_STIMULUS | EPHA2 |
| GO_CELLULAR_RESPONSE_TO_DNA_DAMAGE_STIMULUS | ERCC1 |
| GO_CELLULAR_RESPONSE_TO_DNA_DAMAGE_STIMULUS | ERCC2 |
| GO_CELLULAR_RESPONSE_TO_DNA_DAMAGE_STIMULUS | ERCC3 |
| GO_CELLULAR_RESPONSE_TO_DNA_DAMAGE_STIMULUS | ERCC4 |
| GO_CELLULAR_RESPONSE_TO_DNA_DAMAGE_STIMULUS | ERCC5 |
| GO_CELLULAR_RESPONSE_TO_DNA_DAMAGE_STIMULUS | ERCC6 |
| GO_CELLULAR_RESPONSE_TO_DNA_DAMAGE_STIMULUS | ERCC8 |
| GO_CELLULAR_RESPONSE_TO_DNA_DAMAGE_STIMULUS | ESCO2 |
| GO_CELLULAR_RESPONSE_TO_DNA_DAMAGE_STIMULUS | EXD2 |
| GO_CELLULAR_RESPONSE_TO_DNA_DAMAGE_STIMULUS | EXO1 |
| GO_CELLULAR_RESPONSE_TO_DNA_DAMAGE_STIMULUS | EYA1 |
| GO_CELLULAR_RESPONSE_TO_DNA_DAMAGE_STIMULUS | EYA2 |
| GO_CELLULAR_RESPONSE_TO_DNA_DAMAGE_STIMULUS | EYA3 |
| GO_CELLULAR_RESPONSE_TO_DNA_DAMAGE_STIMULUS | EYA4 |
| GO_CELLULAR_RESPONSE_TO_DNA_DAMAGE_STIMULUS | FAM175A |
| GO_CELLULAR_RESPONSE_TO_DNA_DAMAGE_STIMULUS | FAM178A |
| GO_CELLULAR_RESPONSE_TO_DNA_DAMAGE_STIMULUS | FAN1 |
| GO_CELLULAR_RESPONSE_TO_DNA_DAMAGE_STIMULUS | FANCA |
| GO_CELLULAR_RESPONSE_TO_DNA_DAMAGE_STIMULUS | FANCB |
| GO_CELLULAR_RESPONSE_TO_DNA_DAMAGE_STIMULUS | FANCC |
| GO_CELLULAR_RESPONSE_TO_DNA_DAMAGE_STIMULUS | FANCD2 |

| | |
|---|---|
| GO_CELLULAR_RESPONSE_TO_DNA_DAMAGE_STIMULUS | FANCE |
| GO_CELLULAR_RESPONSE_TO_DNA_DAMAGE_STIMULUS | FANCF |
| GO_CELLULAR_RESPONSE_TO_DNA_DAMAGE_STIMULUS | FANCG |
| GO_CELLULAR_RESPONSE_TO_DNA_DAMAGE_STIMULUS | FANCI |
| GO_CELLULAR_RESPONSE_TO_DNA_DAMAGE_STIMULUS | FANCL |
| GO_CELLULAR_RESPONSE_TO_DNA_DAMAGE_STIMULUS | FANCM |
| GO_CELLULAR_RESPONSE_TO_DNA_DAMAGE_STIMULUS | FBXO18 |
| GO_CELLULAR_RESPONSE_TO_DNA_DAMAGE_STIMULUS | FBXO31 |
| GO_CELLULAR_RESPONSE_TO_DNA_DAMAGE_STIMULUS | FBXO45 |
| GO_CELLULAR_RESPONSE_TO_DNA_DAMAGE_STIMULUS | FBXO6 |
| GO_CELLULAR_RESPONSE_TO_DNA_DAMAGE_STIMULUS | FBXW7 |
| GO_CELLULAR_RESPONSE_TO_DNA_DAMAGE_STIMULUS | FEN1 |
| GO_CELLULAR_RESPONSE_TO_DNA_DAMAGE_STIMULUS | FMN2 |
| GO_CELLULAR_RESPONSE_TO_DNA_DAMAGE_STIMULUS | FNIP2 |
| GO_CELLULAR_RESPONSE_TO_DNA_DAMAGE_STIMULUS | FOXM1 |
| GO_CELLULAR_RESPONSE_TO_DNA_DAMAGE_STIMULUS | FOXN3 |
| GO_CELLULAR_RESPONSE_TO_DNA_DAMAGE_STIMULUS | FOXO1 |
| GO_CELLULAR_RESPONSE_TO_DNA_DAMAGE_STIMULUS | FOXO3 |
| GO_CELLULAR_RESPONSE_TO_DNA_DAMAGE_STIMULUS | FOXO4 |
| GO_CELLULAR_RESPONSE_TO_DNA_DAMAGE_STIMULUS | FTO |
| GO_CELLULAR_RESPONSE_TO_DNA_DAMAGE_STIMULUS | FZR1 |
| GO_CELLULAR_RESPONSE_TO_DNA_DAMAGE_STIMULUS | GADD45A |
| GO_CELLULAR_RESPONSE_TO_DNA_DAMAGE_STIMULUS | GEN1 |
| GO_CELLULAR_RESPONSE_TO_DNA_DAMAGE_STIMULUS | GGN |
| GO_CELLULAR_RESPONSE_TO_DNA_DAMAGE_STIMULUS | GIGYF2 |
| GO_CELLULAR_RESPONSE_TO_DNA_DAMAGE_STIMULUS | GINS2 |
| GO_CELLULAR_RESPONSE_TO_DNA_DAMAGE_STIMULUS | GINS4 |
| GO_CELLULAR_RESPONSE_TO_DNA_DAMAGE_STIMULUS | GML |
| GO_CELLULAR_RESPONSE_TO_DNA_DAMAGE_STIMULUS | GNL1 |
| GO_CELLULAR_RESPONSE_TO_DNA_DAMAGE_STIMULUS | GPS1 |
| GO_CELLULAR_RESPONSE_TO_DNA_DAMAGE_STIMULUS | GRB2 |
| GO_CELLULAR_RESPONSE_TO_DNA_DAMAGE_STIMULUS | GTF2H1 |
| GO_CELLULAR_RESPONSE_TO_DNA_DAMAGE_STIMULUS | GTF2H2 |
| GO_CELLULAR_RESPONSE_TO_DNA_DAMAGE_STIMULUS | GTF2H2C |
| GO_CELLULAR_RESPONSE_TO_DNA_DAMAGE_STIMULUS | GTF2H2D |
| GO_CELLULAR_RESPONSE_TO_DNA_DAMAGE_STIMULUS | GTF2H3 |
| GO_CELLULAR_RESPONSE_TO_DNA_DAMAGE_STIMULUS | GTF2H4 |
| GO_CELLULAR_RESPONSE_TO_DNA_DAMAGE_STIMULUS | GTF2H5 |
| GO_CELLULAR_RESPONSE_TO_DNA_DAMAGE_STIMULUS | GTSE1 |
| GO_CELLULAR_RESPONSE_TO_DNA_DAMAGE_STIMULUS | H2AFX |
| GO_CELLULAR_RESPONSE_TO_DNA_DAMAGE_STIMULUS | HELB |
| GO_CELLULAR_RESPONSE_TO_DNA_DAMAGE_STIMULUS | HELQ |
| GO_CELLULAR_RESPONSE_TO_DNA_DAMAGE_STIMULUS | HERC2 |
| GO_CELLULAR_RESPONSE_TO_DNA_DAMAGE_STIMULUS | HIC1 |
| GO_CELLULAR_RESPONSE_TO_DNA_DAMAGE_STIMULUS | HINFP |
| GO_CELLULAR_RESPONSE_TO_DNA_DAMAGE_STIMULUS | HIPK1 |
| GO_CELLULAR_RESPONSE_TO_DNA_DAMAGE_STIMULUS | HIPK2 |

| | |
|---|---|
| GO_CELLULAR_RESPONSE_TO_DNA_DAMAGE_STIMULUS | HIST1H4A |
| GO_CELLULAR_RESPONSE_TO_DNA_DAMAGE_STIMULUS | HIST1H4B |
| GO_CELLULAR_RESPONSE_TO_DNA_DAMAGE_STIMULUS | HIST1H4C |
| GO_CELLULAR_RESPONSE_TO_DNA_DAMAGE_STIMULUS | HIST1H4D |
| GO_CELLULAR_RESPONSE_TO_DNA_DAMAGE_STIMULUS | HIST1H4E |
| GO_CELLULAR_RESPONSE_TO_DNA_DAMAGE_STIMULUS | HIST1H4F |
| GO_CELLULAR_RESPONSE_TO_DNA_DAMAGE_STIMULUS | HIST1H4H |
| GO_CELLULAR_RESPONSE_TO_DNA_DAMAGE_STIMULUS | HIST1H4I |
| GO_CELLULAR_RESPONSE_TO_DNA_DAMAGE_STIMULUS | HIST1H4J |
| GO_CELLULAR_RESPONSE_TO_DNA_DAMAGE_STIMULUS | HIST1H4K |
| GO_CELLULAR_RESPONSE_TO_DNA_DAMAGE_STIMULUS | HIST1H4L |
| GO_CELLULAR_RESPONSE_TO_DNA_DAMAGE_STIMULUS | HIST2H4A |
| GO_CELLULAR_RESPONSE_TO_DNA_DAMAGE_STIMULUS | HIST2H4B |
| GO_CELLULAR_RESPONSE_TO_DNA_DAMAGE_STIMULUS | HIST3H2A |
| GO_CELLULAR_RESPONSE_TO_DNA_DAMAGE_STIMULUS | HIST3H3 |
| GO_CELLULAR_RESPONSE_TO_DNA_DAMAGE_STIMULUS | HIST4H4 |
| GO_CELLULAR_RESPONSE_TO_DNA_DAMAGE_STIMULUS | HLTF |
| GO_CELLULAR_RESPONSE_TO_DNA_DAMAGE_STIMULUS | HMGA1 |
| GO_CELLULAR_RESPONSE_TO_DNA_DAMAGE_STIMULUS | HMGA2 |
| GO_CELLULAR_RESPONSE_TO_DNA_DAMAGE_STIMULUS | HMGB1 |
| GO_CELLULAR_RESPONSE_TO_DNA_DAMAGE_STIMULUS | HMGB2 |
| GO_CELLULAR_RESPONSE_TO_DNA_DAMAGE_STIMULUS | HMGN1 |
| GO_CELLULAR_RESPONSE_TO_DNA_DAMAGE_STIMULUS | HMOX1 |
| GO_CELLULAR_RESPONSE_TO_DNA_DAMAGE_STIMULUS | HSPA1A |
| GO_CELLULAR_RESPONSE_TO_DNA_DAMAGE_STIMULUS | HTRA2 |
| GO_CELLULAR_RESPONSE_TO_DNA_DAMAGE_STIMULUS | HUS1 |
| GO_CELLULAR_RESPONSE_TO_DNA_DAMAGE_STIMULUS | HUS1B |
| GO_CELLULAR_RESPONSE_TO_DNA_DAMAGE_STIMULUS | HUWE1 |
| GO_CELLULAR_RESPONSE_TO_DNA_DAMAGE_STIMULUS | IFI16 |
| GO_CELLULAR_RESPONSE_TO_DNA_DAMAGE_STIMULUS | IGHMBP2 |
| GO_CELLULAR_RESPONSE_TO_DNA_DAMAGE_STIMULUS | IKBKE |
| GO_CELLULAR_RESPONSE_TO_DNA_DAMAGE_STIMULUS | IKBKG |
| GO_CELLULAR_RESPONSE_TO_DNA_DAMAGE_STIMULUS | IMMP2L |
| GO_CELLULAR_RESPONSE_TO_DNA_DAMAGE_STIMULUS | ING4 |
| GO_CELLULAR_RESPONSE_TO_DNA_DAMAGE_STIMULUS | INO80 |
| GO_CELLULAR_RESPONSE_TO_DNA_DAMAGE_STIMULUS | INO80B |
| GO_CELLULAR_RESPONSE_TO_DNA_DAMAGE_STIMULUS | INO80C |
| GO_CELLULAR_RESPONSE_TO_DNA_DAMAGE_STIMULUS | INO80D |
| GO_CELLULAR_RESPONSE_TO_DNA_DAMAGE_STIMULUS | INO80E |
| GO_CELLULAR_RESPONSE_TO_DNA_DAMAGE_STIMULUS | INTS3 |
| GO_CELLULAR_RESPONSE_TO_DNA_DAMAGE_STIMULUS | INTS7 |
| GO_CELLULAR_RESPONSE_TO_DNA_DAMAGE_STIMULUS | IRF3 |
| GO_CELLULAR_RESPONSE_TO_DNA_DAMAGE_STIMULUS | IRF7 |
| GO_CELLULAR_RESPONSE_TO_DNA_DAMAGE_STIMULUS | ISG15 |
| GO_CELLULAR_RESPONSE_TO_DNA_DAMAGE_STIMULUS | ISY1 |
| GO_CELLULAR_RESPONSE_TO_DNA_DAMAGE_STIMULUS | JMY |
| GO_CELLULAR_RESPONSE_TO_DNA_DAMAGE_STIMULUS | KAT5 |

| | |
|---|---|
| GO_CELLULAR_RESPONSE_TO_DNA_DAMAGE_STIMULUS | KDM2A |
| GO_CELLULAR_RESPONSE_TO_DNA_DAMAGE_STIMULUS | KDM4D |
| GO_CELLULAR_RESPONSE_TO_DNA_DAMAGE_STIMULUS | KIAA0101 |
| GO_CELLULAR_RESPONSE_TO_DNA_DAMAGE_STIMULUS | KIAA0146 |
| GO_CELLULAR_RESPONSE_TO_DNA_DAMAGE_STIMULUS | KIAA0247 |
| GO_CELLULAR_RESPONSE_TO_DNA_DAMAGE_STIMULUS | KIAA0415 |
| GO_CELLULAR_RESPONSE_TO_DNA_DAMAGE_STIMULUS | KIAA0430 |
| GO_CELLULAR_RESPONSE_TO_DNA_DAMAGE_STIMULUS | KIAA1530 |
| GO_CELLULAR_RESPONSE_TO_DNA_DAMAGE_STIMULUS | KIAA1967 |
| GO_CELLULAR_RESPONSE_TO_DNA_DAMAGE_STIMULUS | KIF22 |
| GO_CELLULAR_RESPONSE_TO_DNA_DAMAGE_STIMULUS | KIN |
| GO_CELLULAR_RESPONSE_TO_DNA_DAMAGE_STIMULUS | LIG1 |
| GO_CELLULAR_RESPONSE_TO_DNA_DAMAGE_STIMULUS | LIG3 |
| GO_CELLULAR_RESPONSE_TO_DNA_DAMAGE_STIMULUS | LIG4 |
| GO_CELLULAR_RESPONSE_TO_DNA_DAMAGE_STIMULUS | LOC100133315 |
| GO_CELLULAR_RESPONSE_TO_DNA_DAMAGE_STIMULUS | LOC389493 |
| GO_CELLULAR_RESPONSE_TO_DNA_DAMAGE_STIMULUS | LYN |
| GO_CELLULAR_RESPONSE_TO_DNA_DAMAGE_STIMULUS | MACROD1 |
| GO_CELLULAR_RESPONSE_TO_DNA_DAMAGE_STIMULUS | MACROD2 |
| GO_CELLULAR_RESPONSE_TO_DNA_DAMAGE_STIMULUS | MAD2L2 |
| GO_CELLULAR_RESPONSE_TO_DNA_DAMAGE_STIMULUS | MAEL |
| GO_CELLULAR_RESPONSE_TO_DNA_DAMAGE_STIMULUS | MAP2K6 |
| GO_CELLULAR_RESPONSE_TO_DNA_DAMAGE_STIMULUS | MAPK1 |
| GO_CELLULAR_RESPONSE_TO_DNA_DAMAGE_STIMULUS | MAPK12 |
| GO_CELLULAR_RESPONSE_TO_DNA_DAMAGE_STIMULUS | MAPK14 |
| GO_CELLULAR_RESPONSE_TO_DNA_DAMAGE_STIMULUS | MAPK3 |
| GO_CELLULAR_RESPONSE_TO_DNA_DAMAGE_STIMULUS | MAPKAPK2 |
| GO_CELLULAR_RESPONSE_TO_DNA_DAMAGE_STIMULUS | MASTL |
| GO_CELLULAR_RESPONSE_TO_DNA_DAMAGE_STIMULUS | MBD4 |
| GO_CELLULAR_RESPONSE_TO_DNA_DAMAGE_STIMULUS | MC1R |
| GO_CELLULAR_RESPONSE_TO_DNA_DAMAGE_STIMULUS | MCL1 |
| GO_CELLULAR_RESPONSE_TO_DNA_DAMAGE_STIMULUS | MCM10 |
| GO_CELLULAR_RESPONSE_TO_DNA_DAMAGE_STIMULUS | MCM7 |
| GO_CELLULAR_RESPONSE_TO_DNA_DAMAGE_STIMULUS | MCM8 |
| GO_CELLULAR_RESPONSE_TO_DNA_DAMAGE_STIMULUS | MCM9 |
| GO_CELLULAR_RESPONSE_TO_DNA_DAMAGE_STIMULUS | MCRS1 |
| GO_CELLULAR_RESPONSE_TO_DNA_DAMAGE_STIMULUS | MCTS1 |
| GO_CELLULAR_RESPONSE_TO_DNA_DAMAGE_STIMULUS | MDC1 |
| GO_CELLULAR_RESPONSE_TO_DNA_DAMAGE_STIMULUS | MDM2 |
| GO_CELLULAR_RESPONSE_TO_DNA_DAMAGE_STIMULUS | MDM4 |
| GO_CELLULAR_RESPONSE_TO_DNA_DAMAGE_STIMULUS | MEN1 |
| GO_CELLULAR_RESPONSE_TO_DNA_DAMAGE_STIMULUS | MGMT |
| GO_CELLULAR_RESPONSE_TO_DNA_DAMAGE_STIMULUS | MICA |
| GO_CELLULAR_RESPONSE_TO_DNA_DAMAGE_STIMULUS | MIF |
| GO_CELLULAR_RESPONSE_TO_DNA_DAMAGE_STIMULUS | MLH1 |
| GO_CELLULAR_RESPONSE_TO_DNA_DAMAGE_STIMULUS | MLH3 |
| GO_CELLULAR_RESPONSE_TO_DNA_DAMAGE_STIMULUS | MMS19 |

| | |
|---|---|
| GO_CELLULAR_RESPONSE_TO_DNA_DAMAGE_STIMULUS | MMS22L |
| GO_CELLULAR_RESPONSE_TO_DNA_DAMAGE_STIMULUS | MNAT1 |
| GO_CELLULAR_RESPONSE_TO_DNA_DAMAGE_STIMULUS | MNDA |
| GO_CELLULAR_RESPONSE_TO_DNA_DAMAGE_STIMULUS | MOAP1 |
| GO_CELLULAR_RESPONSE_TO_DNA_DAMAGE_STIMULUS | MORF4L1 |
| GO_CELLULAR_RESPONSE_TO_DNA_DAMAGE_STIMULUS | MORF4L2 |
| GO_CELLULAR_RESPONSE_TO_DNA_DAMAGE_STIMULUS | MPG |
| GO_CELLULAR_RESPONSE_TO_DNA_DAMAGE_STIMULUS | MRE11A |
| GO_CELLULAR_RESPONSE_TO_DNA_DAMAGE_STIMULUS | MRPS11 |
| GO_CELLULAR_RESPONSE_TO_DNA_DAMAGE_STIMULUS | MRPS26 |
| GO_CELLULAR_RESPONSE_TO_DNA_DAMAGE_STIMULUS | MRPS35 |
| GO_CELLULAR_RESPONSE_TO_DNA_DAMAGE_STIMULUS | MRPS9 |
| GO_CELLULAR_RESPONSE_TO_DNA_DAMAGE_STIMULUS | MSH2 |
| GO_CELLULAR_RESPONSE_TO_DNA_DAMAGE_STIMULUS | MSH3 |
| GO_CELLULAR_RESPONSE_TO_DNA_DAMAGE_STIMULUS | MSH4 |
| GO_CELLULAR_RESPONSE_TO_DNA_DAMAGE_STIMULUS | MSH5 |
| GO_CELLULAR_RESPONSE_TO_DNA_DAMAGE_STIMULUS | MSH6 |
| GO_CELLULAR_RESPONSE_TO_DNA_DAMAGE_STIMULUS | MTA1 |
| GO_CELLULAR_RESPONSE_TO_DNA_DAMAGE_STIMULUS | MTOR |
| GO_CELLULAR_RESPONSE_TO_DNA_DAMAGE_STIMULUS | MUC1 |
| GO_CELLULAR_RESPONSE_TO_DNA_DAMAGE_STIMULUS | MUM1 |
| GO_CELLULAR_RESPONSE_TO_DNA_DAMAGE_STIMULUS | MUS81 |
| GO_CELLULAR_RESPONSE_TO_DNA_DAMAGE_STIMULUS | MUTYH |
| GO_CELLULAR_RESPONSE_TO_DNA_DAMAGE_STIMULUS | MYC |
| GO_CELLULAR_RESPONSE_TO_DNA_DAMAGE_STIMULUS | MYO6 |
| GO_CELLULAR_RESPONSE_TO_DNA_DAMAGE_STIMULUS | NBN |
| GO_CELLULAR_RESPONSE_TO_DNA_DAMAGE_STIMULUS | NCOA6 |
| GO_CELLULAR_RESPONSE_TO_DNA_DAMAGE_STIMULUS | NDNL2 |
| GO_CELLULAR_RESPONSE_TO_DNA_DAMAGE_STIMULUS | NDRG1 |
| GO_CELLULAR_RESPONSE_TO_DNA_DAMAGE_STIMULUS | NEIL1 |
| GO_CELLULAR_RESPONSE_TO_DNA_DAMAGE_STIMULUS | NEIL2 |
| GO_CELLULAR_RESPONSE_TO_DNA_DAMAGE_STIMULUS | NEIL3 |
| GO_CELLULAR_RESPONSE_TO_DNA_DAMAGE_STIMULUS | NEK11 |
| GO_CELLULAR_RESPONSE_TO_DNA_DAMAGE_STIMULUS | NEK4 |
| GO_CELLULAR_RESPONSE_TO_DNA_DAMAGE_STIMULUS | NEK6 |
| GO_CELLULAR_RESPONSE_TO_DNA_DAMAGE_STIMULUS | NFATC2 |
| GO_CELLULAR_RESPONSE_TO_DNA_DAMAGE_STIMULUS | NFATC4 |
| GO_CELLULAR_RESPONSE_TO_DNA_DAMAGE_STIMULUS | NFRKB |
| GO_CELLULAR_RESPONSE_TO_DNA_DAMAGE_STIMULUS | NHEJ1 |
| GO_CELLULAR_RESPONSE_TO_DNA_DAMAGE_STIMULUS | NIPBL |
| GO_CELLULAR_RESPONSE_TO_DNA_DAMAGE_STIMULUS | NONO |
| GO_CELLULAR_RESPONSE_TO_DNA_DAMAGE_STIMULUS | NPAS2 |
| GO_CELLULAR_RESPONSE_TO_DNA_DAMAGE_STIMULUS | NPLOC4 |
| GO_CELLULAR_RESPONSE_TO_DNA_DAMAGE_STIMULUS | NPM1 |
| GO_CELLULAR_RESPONSE_TO_DNA_DAMAGE_STIMULUS | NSMCE1 |
| GO_CELLULAR_RESPONSE_TO_DNA_DAMAGE_STIMULUS | NSMCE2 |
| GO_CELLULAR_RESPONSE_TO_DNA_DAMAGE_STIMULUS | NSMCE4A |

| | |
|---|---|
| GO_CELLULAR_RESPONSE_TO_DNA_DAMAGE_STIMULUS | NTHL1 |
| GO_CELLULAR_RESPONSE_TO_DNA_DAMAGE_STIMULUS | NUAK1 |
| GO_CELLULAR_RESPONSE_TO_DNA_DAMAGE_STIMULUS | NUDT1 |
| GO_CELLULAR_RESPONSE_TO_DNA_DAMAGE_STIMULUS | NUPR1 |
| GO_CELLULAR_RESPONSE_TO_DNA_DAMAGE_STIMULUS | OBFC2A |
| GO_CELLULAR_RESPONSE_TO_DNA_DAMAGE_STIMULUS | OBFC2B |
| GO_CELLULAR_RESPONSE_TO_DNA_DAMAGE_STIMULUS | OGG1 |
| GO_CELLULAR_RESPONSE_TO_DNA_DAMAGE_STIMULUS | OTUB1 |
| GO_CELLULAR_RESPONSE_TO_DNA_DAMAGE_STIMULUS | PALB2 |
| GO_CELLULAR_RESPONSE_TO_DNA_DAMAGE_STIMULUS | PAPD7 |
| GO_CELLULAR_RESPONSE_TO_DNA_DAMAGE_STIMULUS | PARP1 |
| GO_CELLULAR_RESPONSE_TO_DNA_DAMAGE_STIMULUS | PARP2 |
| GO_CELLULAR_RESPONSE_TO_DNA_DAMAGE_STIMULUS | PARP3 |
| GO_CELLULAR_RESPONSE_TO_DNA_DAMAGE_STIMULUS | PARP4 |
| GO_CELLULAR_RESPONSE_TO_DNA_DAMAGE_STIMULUS | PARP9 |
| GO_CELLULAR_RESPONSE_TO_DNA_DAMAGE_STIMULUS | PAXIP1 |
| GO_CELLULAR_RESPONSE_TO_DNA_DAMAGE_STIMULUS | PCBP4 |
| GO_CELLULAR_RESPONSE_TO_DNA_DAMAGE_STIMULUS | PCNA |
| GO_CELLULAR_RESPONSE_TO_DNA_DAMAGE_STIMULUS | PEA15 |
| GO_CELLULAR_RESPONSE_TO_DNA_DAMAGE_STIMULUS | PGAP2 |
| GO_CELLULAR_RESPONSE_TO_DNA_DAMAGE_STIMULUS | PHF1 |
| GO_CELLULAR_RESPONSE_TO_DNA_DAMAGE_STIMULUS | PHLDA3 |
| GO_CELLULAR_RESPONSE_TO_DNA_DAMAGE_STIMULUS | PIAS4 |
| GO_CELLULAR_RESPONSE_TO_DNA_DAMAGE_STIMULUS | PIDD |
| GO_CELLULAR_RESPONSE_TO_DNA_DAMAGE_STIMULUS | PIF1 |
| GO_CELLULAR_RESPONSE_TO_DNA_DAMAGE_STIMULUS | PIK3R1 |
| GO_CELLULAR_RESPONSE_TO_DNA_DAMAGE_STIMULUS | PLAGL1 |
| GO_CELLULAR_RESPONSE_TO_DNA_DAMAGE_STIMULUS | PLK1 |
| GO_CELLULAR_RESPONSE_TO_DNA_DAMAGE_STIMULUS | PLK2 |
| GO_CELLULAR_RESPONSE_TO_DNA_DAMAGE_STIMULUS | PLK3 |
| GO_CELLULAR_RESPONSE_TO_DNA_DAMAGE_STIMULUS | PLK5 |
| GO_CELLULAR_RESPONSE_TO_DNA_DAMAGE_STIMULUS | PMAIP1 |
| GO_CELLULAR_RESPONSE_TO_DNA_DAMAGE_STIMULUS | PML |
| GO_CELLULAR_RESPONSE_TO_DNA_DAMAGE_STIMULUS | PMS1 |
| GO_CELLULAR_RESPONSE_TO_DNA_DAMAGE_STIMULUS | PMS2 |
| GO_CELLULAR_RESPONSE_TO_DNA_DAMAGE_STIMULUS | PMS2CL |
| GO_CELLULAR_RESPONSE_TO_DNA_DAMAGE_STIMULUS | PMS2P1 |
| GO_CELLULAR_RESPONSE_TO_DNA_DAMAGE_STIMULUS | PMS2P3 |
| GO_CELLULAR_RESPONSE_TO_DNA_DAMAGE_STIMULUS | PMS2P5 |
| GO_CELLULAR_RESPONSE_TO_DNA_DAMAGE_STIMULUS | PNKP |
| GO_CELLULAR_RESPONSE_TO_DNA_DAMAGE_STIMULUS | POLA1 |
| GO_CELLULAR_RESPONSE_TO_DNA_DAMAGE_STIMULUS | POLB |
| GO_CELLULAR_RESPONSE_TO_DNA_DAMAGE_STIMULUS | POLD1 |
| GO_CELLULAR_RESPONSE_TO_DNA_DAMAGE_STIMULUS | POLD2 |
| GO_CELLULAR_RESPONSE_TO_DNA_DAMAGE_STIMULUS | POLD3 |
| GO_CELLULAR_RESPONSE_TO_DNA_DAMAGE_STIMULUS | POLD4 |
| GO_CELLULAR_RESPONSE_TO_DNA_DAMAGE_STIMULUS | POLE |

| | |
|---|---|
| GO_CELLULAR_RESPONSE_TO_DNA_DAMAGE_STIMULUS | POLE2 |
| GO_CELLULAR_RESPONSE_TO_DNA_DAMAGE_STIMULUS | POLG |
| GO_CELLULAR_RESPONSE_TO_DNA_DAMAGE_STIMULUS | POLG2 |
| GO_CELLULAR_RESPONSE_TO_DNA_DAMAGE_STIMULUS | POLH |
| GO_CELLULAR_RESPONSE_TO_DNA_DAMAGE_STIMULUS | POLI |
| GO_CELLULAR_RESPONSE_TO_DNA_DAMAGE_STIMULUS | POLK |
| GO_CELLULAR_RESPONSE_TO_DNA_DAMAGE_STIMULUS | POLL |
| GO_CELLULAR_RESPONSE_TO_DNA_DAMAGE_STIMULUS | POLM |
| GO_CELLULAR_RESPONSE_TO_DNA_DAMAGE_STIMULUS | POLN |
| GO_CELLULAR_RESPONSE_TO_DNA_DAMAGE_STIMULUS | POLQ |
| GO_CELLULAR_RESPONSE_TO_DNA_DAMAGE_STIMULUS | POLR2A |
| GO_CELLULAR_RESPONSE_TO_DNA_DAMAGE_STIMULUS | POLR2B |
| GO_CELLULAR_RESPONSE_TO_DNA_DAMAGE_STIMULUS | POLR2C |
| GO_CELLULAR_RESPONSE_TO_DNA_DAMAGE_STIMULUS | POLR2D |
| GO_CELLULAR_RESPONSE_TO_DNA_DAMAGE_STIMULUS | POLR2E |
| GO_CELLULAR_RESPONSE_TO_DNA_DAMAGE_STIMULUS | POLR2F |
| GO_CELLULAR_RESPONSE_TO_DNA_DAMAGE_STIMULUS | POLR2G |
| GO_CELLULAR_RESPONSE_TO_DNA_DAMAGE_STIMULUS | POLR2H |
| GO_CELLULAR_RESPONSE_TO_DNA_DAMAGE_STIMULUS | POLR2I |
| GO_CELLULAR_RESPONSE_TO_DNA_DAMAGE_STIMULUS | POLR2J |
| GO_CELLULAR_RESPONSE_TO_DNA_DAMAGE_STIMULUS | POLR2K |
| GO_CELLULAR_RESPONSE_TO_DNA_DAMAGE_STIMULUS | POLR2L |
| GO_CELLULAR_RESPONSE_TO_DNA_DAMAGE_STIMULUS | PPIE |
| GO_CELLULAR_RESPONSE_TO_DNA_DAMAGE_STIMULUS | PPP1R15A |
| GO_CELLULAR_RESPONSE_TO_DNA_DAMAGE_STIMULUS | PPP2R5C |
| GO_CELLULAR_RESPONSE_TO_DNA_DAMAGE_STIMULUS | PPP5C |
| GO_CELLULAR_RESPONSE_TO_DNA_DAMAGE_STIMULUS | PRKDC |
| GO_CELLULAR_RESPONSE_TO_DNA_DAMAGE_STIMULUS | PRMT1 |
| GO_CELLULAR_RESPONSE_TO_DNA_DAMAGE_STIMULUS | PRMT6 |
| GO_CELLULAR_RESPONSE_TO_DNA_DAMAGE_STIMULUS | PRPF19 |
| GO_CELLULAR_RESPONSE_TO_DNA_DAMAGE_STIMULUS | PSEN1 |
| GO_CELLULAR_RESPONSE_TO_DNA_DAMAGE_STIMULUS | PSMD14 |
| GO_CELLULAR_RESPONSE_TO_DNA_DAMAGE_STIMULUS | PSME4 |
| GO_CELLULAR_RESPONSE_TO_DNA_DAMAGE_STIMULUS | PTPN11 |
| GO_CELLULAR_RESPONSE_TO_DNA_DAMAGE_STIMULUS | PTTG1 |
| GO_CELLULAR_RESPONSE_TO_DNA_DAMAGE_STIMULUS | PYCARD |
| GO_CELLULAR_RESPONSE_TO_DNA_DAMAGE_STIMULUS | RAD1 |
| GO_CELLULAR_RESPONSE_TO_DNA_DAMAGE_STIMULUS | RAD17 |
| GO_CELLULAR_RESPONSE_TO_DNA_DAMAGE_STIMULUS | RAD18 |
| GO_CELLULAR_RESPONSE_TO_DNA_DAMAGE_STIMULUS | RAD21 |
| GO_CELLULAR_RESPONSE_TO_DNA_DAMAGE_STIMULUS | RAD21L1 |
| GO_CELLULAR_RESPONSE_TO_DNA_DAMAGE_STIMULUS | RAD23A |
| GO_CELLULAR_RESPONSE_TO_DNA_DAMAGE_STIMULUS | RAD23B |
| GO_CELLULAR_RESPONSE_TO_DNA_DAMAGE_STIMULUS | RAD50 |
| GO_CELLULAR_RESPONSE_TO_DNA_DAMAGE_STIMULUS | RAD51 |
| GO_CELLULAR_RESPONSE_TO_DNA_DAMAGE_STIMULUS | RAD51AP1 |
| GO_CELLULAR_RESPONSE_TO_DNA_DAMAGE_STIMULUS | RAD51B |

| | |
|---|---|
| GO_CELLULAR_RESPONSE_TO_DNA_DAMAGE_STIMULUS | RAD51C |
| GO_CELLULAR_RESPONSE_TO_DNA_DAMAGE_STIMULUS | RAD51D |
| GO_CELLULAR_RESPONSE_TO_DNA_DAMAGE_STIMULUS | RAD52 |
| GO_CELLULAR_RESPONSE_TO_DNA_DAMAGE_STIMULUS | RAD54B |
| GO_CELLULAR_RESPONSE_TO_DNA_DAMAGE_STIMULUS | RAD54L |
| GO_CELLULAR_RESPONSE_TO_DNA_DAMAGE_STIMULUS | RAD9A |
| GO_CELLULAR_RESPONSE_TO_DNA_DAMAGE_STIMULUS | RAD9B |
| GO_CELLULAR_RESPONSE_TO_DNA_DAMAGE_STIMULUS | RASSF1 |
| GO_CELLULAR_RESPONSE_TO_DNA_DAMAGE_STIMULUS | RBBP5 |
| GO_CELLULAR_RESPONSE_TO_DNA_DAMAGE_STIMULUS | RBBP6 |
| GO_CELLULAR_RESPONSE_TO_DNA_DAMAGE_STIMULUS | RBBP8 |
| GO_CELLULAR_RESPONSE_TO_DNA_DAMAGE_STIMULUS | RBL2 |
| GO_CELLULAR_RESPONSE_TO_DNA_DAMAGE_STIMULUS | RBM14 |
| GO_CELLULAR_RESPONSE_TO_DNA_DAMAGE_STIMULUS | RBM38 |
| GO_CELLULAR_RESPONSE_TO_DNA_DAMAGE_STIMULUS | RBX1 |
| GO_CELLULAR_RESPONSE_TO_DNA_DAMAGE_STIMULUS | RCHY1 |
| GO_CELLULAR_RESPONSE_TO_DNA_DAMAGE_STIMULUS | REC8 |
| GO_CELLULAR_RESPONSE_TO_DNA_DAMAGE_STIMULUS | RECQL |
| GO_CELLULAR_RESPONSE_TO_DNA_DAMAGE_STIMULUS | RECQL4 |
| GO_CELLULAR_RESPONSE_TO_DNA_DAMAGE_STIMULUS | RECQL5 |
| GO_CELLULAR_RESPONSE_TO_DNA_DAMAGE_STIMULUS | REV1 |
| GO_CELLULAR_RESPONSE_TO_DNA_DAMAGE_STIMULUS | REV3L |
| GO_CELLULAR_RESPONSE_TO_DNA_DAMAGE_STIMULUS | RFC1 |
| GO_CELLULAR_RESPONSE_TO_DNA_DAMAGE_STIMULUS | RFC2 |
| GO_CELLULAR_RESPONSE_TO_DNA_DAMAGE_STIMULUS | RFC3 |
| GO_CELLULAR_RESPONSE_TO_DNA_DAMAGE_STIMULUS | RFC4 |
| GO_CELLULAR_RESPONSE_TO_DNA_DAMAGE_STIMULUS | RFC5 |
| GO_CELLULAR_RESPONSE_TO_DNA_DAMAGE_STIMULUS | RFWD3 |
| GO_CELLULAR_RESPONSE_TO_DNA_DAMAGE_STIMULUS | RIF1 |
| GO_CELLULAR_RESPONSE_TO_DNA_DAMAGE_STIMULUS | RMI1 |
| GO_CELLULAR_RESPONSE_TO_DNA_DAMAGE_STIMULUS | RMI2 |
| GO_CELLULAR_RESPONSE_TO_DNA_DAMAGE_STIMULUS | RNASEH2A |
| GO_CELLULAR_RESPONSE_TO_DNA_DAMAGE_STIMULUS | RNF111 |
| GO_CELLULAR_RESPONSE_TO_DNA_DAMAGE_STIMULUS | RNF138 |
| GO_CELLULAR_RESPONSE_TO_DNA_DAMAGE_STIMULUS | RNF168 |
| GO_CELLULAR_RESPONSE_TO_DNA_DAMAGE_STIMULUS | RNF169 |
| GO_CELLULAR_RESPONSE_TO_DNA_DAMAGE_STIMULUS | RNF8 |
| GO_CELLULAR_RESPONSE_TO_DNA_DAMAGE_STIMULUS | RPA1 |
| GO_CELLULAR_RESPONSE_TO_DNA_DAMAGE_STIMULUS | RPA2 |
| GO_CELLULAR_RESPONSE_TO_DNA_DAMAGE_STIMULUS | RPA3 |
| GO_CELLULAR_RESPONSE_TO_DNA_DAMAGE_STIMULUS | RPA4 |
| GO_CELLULAR_RESPONSE_TO_DNA_DAMAGE_STIMULUS | RPAIN |
| GO_CELLULAR_RESPONSE_TO_DNA_DAMAGE_STIMULUS | RPS27A |
| GO_CELLULAR_RESPONSE_TO_DNA_DAMAGE_STIMULUS | RPS27L |
| GO_CELLULAR_RESPONSE_TO_DNA_DAMAGE_STIMULUS | RPS3 |
| GO_CELLULAR_RESPONSE_TO_DNA_DAMAGE_STIMULUS | RPS6KA6 |
| GO_CELLULAR_RESPONSE_TO_DNA_DAMAGE_STIMULUS | RQCD1 |

| | |
|---|---|
| GO_CELLULAR_RESPONSE_TO_DNA_DAMAGE_STIMULUS | RRM2B |
| GO_CELLULAR_RESPONSE_TO_DNA_DAMAGE_STIMULUS | RTEL1 |
| GO_CELLULAR_RESPONSE_TO_DNA_DAMAGE_STIMULUS | RUVBL1 |
| GO_CELLULAR_RESPONSE_TO_DNA_DAMAGE_STIMULUS | RUVBL2 |
| GO_CELLULAR_RESPONSE_TO_DNA_DAMAGE_STIMULUS | SESN1 |
| GO_CELLULAR_RESPONSE_TO_DNA_DAMAGE_STIMULUS | SETD2 |
| GO_CELLULAR_RESPONSE_TO_DNA_DAMAGE_STIMULUS | SETD7 |
| GO_CELLULAR_RESPONSE_TO_DNA_DAMAGE_STIMULUS | SETMAR |
| GO_CELLULAR_RESPONSE_TO_DNA_DAMAGE_STIMULUS | SETX |
| GO_CELLULAR_RESPONSE_TO_DNA_DAMAGE_STIMULUS | SFN |
| GO_CELLULAR_RESPONSE_TO_DNA_DAMAGE_STIMULUS | SFPQ |
| GO_CELLULAR_RESPONSE_TO_DNA_DAMAGE_STIMULUS | SFR1 |
| GO_CELLULAR_RESPONSE_TO_DNA_DAMAGE_STIMULUS | SGK1 |
| GO_CELLULAR_RESPONSE_TO_DNA_DAMAGE_STIMULUS | SHFM1 |
| GO_CELLULAR_RESPONSE_TO_DNA_DAMAGE_STIMULUS | SHISA5 |
| GO_CELLULAR_RESPONSE_TO_DNA_DAMAGE_STIMULUS | SHPRH |
| GO_CELLULAR_RESPONSE_TO_DNA_DAMAGE_STIMULUS | SIRT1 |
| GO_CELLULAR_RESPONSE_TO_DNA_DAMAGE_STIMULUS | SIRT4 |
| GO_CELLULAR_RESPONSE_TO_DNA_DAMAGE_STIMULUS | SIRT6 |
| GO_CELLULAR_RESPONSE_TO_DNA_DAMAGE_STIMULUS | SLC30A9 |
| GO_CELLULAR_RESPONSE_TO_DNA_DAMAGE_STIMULUS | SLX1A |
| GO_CELLULAR_RESPONSE_TO_DNA_DAMAGE_STIMULUS | SLX1B |
| GO_CELLULAR_RESPONSE_TO_DNA_DAMAGE_STIMULUS | SLX4 |
| GO_CELLULAR_RESPONSE_TO_DNA_DAMAGE_STIMULUS | SMARCA5 |
| GO_CELLULAR_RESPONSE_TO_DNA_DAMAGE_STIMULUS | SMARCAD1 |
| GO_CELLULAR_RESPONSE_TO_DNA_DAMAGE_STIMULUS | SMARCAL1 |
| GO_CELLULAR_RESPONSE_TO_DNA_DAMAGE_STIMULUS | SMARCB1 |
| GO_CELLULAR_RESPONSE_TO_DNA_DAMAGE_STIMULUS | SMC1A |
| GO_CELLULAR_RESPONSE_TO_DNA_DAMAGE_STIMULUS | SMC3 |
| GO_CELLULAR_RESPONSE_TO_DNA_DAMAGE_STIMULUS | SMC5 |
| GO_CELLULAR_RESPONSE_TO_DNA_DAMAGE_STIMULUS | SMC6 |
| GO_CELLULAR_RESPONSE_TO_DNA_DAMAGE_STIMULUS | SMG1 |
| GO_CELLULAR_RESPONSE_TO_DNA_DAMAGE_STIMULUS | SMUG1 |
| GO_CELLULAR_RESPONSE_TO_DNA_DAMAGE_STIMULUS | SNW1 |
| GO_CELLULAR_RESPONSE_TO_DNA_DAMAGE_STIMULUS | SOX4 |
| GO_CELLULAR_RESPONSE_TO_DNA_DAMAGE_STIMULUS | SP100 |
| GO_CELLULAR_RESPONSE_TO_DNA_DAMAGE_STIMULUS | SPATA18 |
| GO_CELLULAR_RESPONSE_TO_DNA_DAMAGE_STIMULUS | SPATA22 |
| GO_CELLULAR_RESPONSE_TO_DNA_DAMAGE_STIMULUS | SPDYA |
| GO_CELLULAR_RESPONSE_TO_DNA_DAMAGE_STIMULUS | SSRP1 |
| GO_CELLULAR_RESPONSE_TO_DNA_DAMAGE_STIMULUS | ST20 |
| GO_CELLULAR_RESPONSE_TO_DNA_DAMAGE_STIMULUS | STK11 |
| GO_CELLULAR_RESPONSE_TO_DNA_DAMAGE_STIMULUS | STRA13 |
| GO_CELLULAR_RESPONSE_TO_DNA_DAMAGE_STIMULUS | STUB1 |
| GO_CELLULAR_RESPONSE_TO_DNA_DAMAGE_STIMULUS | STXBP4 |
| GO_CELLULAR_RESPONSE_TO_DNA_DAMAGE_STIMULUS | SUMO1 |
| GO_CELLULAR_RESPONSE_TO_DNA_DAMAGE_STIMULUS | SUMO2 |

| | |
|---|---|
| GO_CELLULAR_RESPONSE_TO_DNA_DAMAGE_STIMULUS | SUMO3 |
| GO_CELLULAR_RESPONSE_TO_DNA_DAMAGE_STIMULUS | SUPT16H |
| GO_CELLULAR_RESPONSE_TO_DNA_DAMAGE_STIMULUS | SUV39H1 |
| GO_CELLULAR_RESPONSE_TO_DNA_DAMAGE_STIMULUS | SWI5 |
| GO_CELLULAR_RESPONSE_TO_DNA_DAMAGE_STIMULUS | SYCP1 |
| GO_CELLULAR_RESPONSE_TO_DNA_DAMAGE_STIMULUS | SYF2 |
| GO_CELLULAR_RESPONSE_TO_DNA_DAMAGE_STIMULUS | TAF1 |
| GO_CELLULAR_RESPONSE_TO_DNA_DAMAGE_STIMULUS | TAF9 |
| GO_CELLULAR_RESPONSE_TO_DNA_DAMAGE_STIMULUS | TAOK1 |
| GO_CELLULAR_RESPONSE_TO_DNA_DAMAGE_STIMULUS | TAOK2 |
| GO_CELLULAR_RESPONSE_TO_DNA_DAMAGE_STIMULUS | TAOK3 |
| GO_CELLULAR_RESPONSE_TO_DNA_DAMAGE_STIMULUS | TCEA1 |
| GO_CELLULAR_RESPONSE_TO_DNA_DAMAGE_STIMULUS | TDG |
| GO_CELLULAR_RESPONSE_TO_DNA_DAMAGE_STIMULUS | TDP1 |
| GO_CELLULAR_RESPONSE_TO_DNA_DAMAGE_STIMULUS | TDP2 |
| GO_CELLULAR_RESPONSE_TO_DNA_DAMAGE_STIMULUS | TERF2 |
| GO_CELLULAR_RESPONSE_TO_DNA_DAMAGE_STIMULUS | TERF2IP |
| GO_CELLULAR_RESPONSE_TO_DNA_DAMAGE_STIMULUS | TEX12 |
| GO_CELLULAR_RESPONSE_TO_DNA_DAMAGE_STIMULUS | TEX15 |
| GO_CELLULAR_RESPONSE_TO_DNA_DAMAGE_STIMULUS | TFAP4 |
| GO_CELLULAR_RESPONSE_TO_DNA_DAMAGE_STIMULUS | TFDP1 |
| GO_CELLULAR_RESPONSE_TO_DNA_DAMAGE_STIMULUS | TFDP2 |
| GO_CELLULAR_RESPONSE_TO_DNA_DAMAGE_STIMULUS | TFDP3 |
| GO_CELLULAR_RESPONSE_TO_DNA_DAMAGE_STIMULUS | TFIP11 |
| GO_CELLULAR_RESPONSE_TO_DNA_DAMAGE_STIMULUS | TFPT |
| GO_CELLULAR_RESPONSE_TO_DNA_DAMAGE_STIMULUS | THOC1 |
| GO_CELLULAR_RESPONSE_TO_DNA_DAMAGE_STIMULUS | THOC4 |
| GO_CELLULAR_RESPONSE_TO_DNA_DAMAGE_STIMULUS | TIMELESS |
| GO_CELLULAR_RESPONSE_TO_DNA_DAMAGE_STIMULUS | TIPIN |
| GO_CELLULAR_RESPONSE_TO_DNA_DAMAGE_STIMULUS | TIPRL |
| GO_CELLULAR_RESPONSE_TO_DNA_DAMAGE_STIMULUS | TLK1 |
| GO_CELLULAR_RESPONSE_TO_DNA_DAMAGE_STIMULUS | TLK2 |
| GO_CELLULAR_RESPONSE_TO_DNA_DAMAGE_STIMULUS | TMEM109 |
| GO_CELLULAR_RESPONSE_TO_DNA_DAMAGE_STIMULUS | TNF |
| GO_CELLULAR_RESPONSE_TO_DNA_DAMAGE_STIMULUS | TNFRSF1A |
| GO_CELLULAR_RESPONSE_TO_DNA_DAMAGE_STIMULUS | TNFRSF1B |
| GO_CELLULAR_RESPONSE_TO_DNA_DAMAGE_STIMULUS | TNKS1BP1 |
| GO_CELLULAR_RESPONSE_TO_DNA_DAMAGE_STIMULUS | TNP1 |
| GO_CELLULAR_RESPONSE_TO_DNA_DAMAGE_STIMULUS | TONSL |
| GO_CELLULAR_RESPONSE_TO_DNA_DAMAGE_STIMULUS | TOP2A |
| GO_CELLULAR_RESPONSE_TO_DNA_DAMAGE_STIMULUS | TOP3A |
| GO_CELLULAR_RESPONSE_TO_DNA_DAMAGE_STIMULUS | TOPBP1 |
| GO_CELLULAR_RESPONSE_TO_DNA_DAMAGE_STIMULUS | TOPORS |
| GO_CELLULAR_RESPONSE_TO_DNA_DAMAGE_STIMULUS | TP53 |
| GO_CELLULAR_RESPONSE_TO_DNA_DAMAGE_STIMULUS | TP53BP1 |
| GO_CELLULAR_RESPONSE_TO_DNA_DAMAGE_STIMULUS | TP53TG1 |
| GO_CELLULAR_RESPONSE_TO_DNA_DAMAGE_STIMULUS | TP63 |

| | |
|---|---|
| GO_CELLULAR_RESPONSE_TO_DNA_DAMAGE_STIMULUS | TP73 |
| GO_CELLULAR_RESPONSE_TO_DNA_DAMAGE_STIMULUS | TREX1 |
| GO_CELLULAR_RESPONSE_TO_DNA_DAMAGE_STIMULUS | TREX2 |
| GO_CELLULAR_RESPONSE_TO_DNA_DAMAGE_STIMULUS | TRIAP1 |
| GO_CELLULAR_RESPONSE_TO_DNA_DAMAGE_STIMULUS | TRIM25 |
| GO_CELLULAR_RESPONSE_TO_DNA_DAMAGE_STIMULUS | TRIM28 |
| GO_CELLULAR_RESPONSE_TO_DNA_DAMAGE_STIMULUS | TRIP12 |
| GO_CELLULAR_RESPONSE_TO_DNA_DAMAGE_STIMULUS | TRIP13 |
| GO_CELLULAR_RESPONSE_TO_DNA_DAMAGE_STIMULUS | TRRAP |
| GO_CELLULAR_RESPONSE_TO_DNA_DAMAGE_STIMULUS | TTC5 |
| GO_CELLULAR_RESPONSE_TO_DNA_DAMAGE_STIMULUS | UBA1 |
| GO_CELLULAR_RESPONSE_TO_DNA_DAMAGE_STIMULUS | UBA52 |
| GO_CELLULAR_RESPONSE_TO_DNA_DAMAGE_STIMULUS | UBA7 |
| GO_CELLULAR_RESPONSE_TO_DNA_DAMAGE_STIMULUS | UBB |
| GO_CELLULAR_RESPONSE_TO_DNA_DAMAGE_STIMULUS | UBC |
| GO_CELLULAR_RESPONSE_TO_DNA_DAMAGE_STIMULUS | UBE2A |
| GO_CELLULAR_RESPONSE_TO_DNA_DAMAGE_STIMULUS | UBE2B |
| GO_CELLULAR_RESPONSE_TO_DNA_DAMAGE_STIMULUS | UBE2D3 |
| GO_CELLULAR_RESPONSE_TO_DNA_DAMAGE_STIMULUS | UBE2E2 |
| GO_CELLULAR_RESPONSE_TO_DNA_DAMAGE_STIMULUS | UBE2F |
| GO_CELLULAR_RESPONSE_TO_DNA_DAMAGE_STIMULUS | UBE2I |
| GO_CELLULAR_RESPONSE_TO_DNA_DAMAGE_STIMULUS | UBE2L6 |
| GO_CELLULAR_RESPONSE_TO_DNA_DAMAGE_STIMULUS | UBE2N |
| GO_CELLULAR_RESPONSE_TO_DNA_DAMAGE_STIMULUS | UBE2NL |
| GO_CELLULAR_RESPONSE_TO_DNA_DAMAGE_STIMULUS | UBE2T |
| GO_CELLULAR_RESPONSE_TO_DNA_DAMAGE_STIMULUS | UBE2U |
| GO_CELLULAR_RESPONSE_TO_DNA_DAMAGE_STIMULUS | UBE2V1 |
| GO_CELLULAR_RESPONSE_TO_DNA_DAMAGE_STIMULUS | UBE2V2 |
| GO_CELLULAR_RESPONSE_TO_DNA_DAMAGE_STIMULUS | UBE2W |
| GO_CELLULAR_RESPONSE_TO_DNA_DAMAGE_STIMULUS | UBR5 |
| GO_CELLULAR_RESPONSE_TO_DNA_DAMAGE_STIMULUS | UCHL5 |
| GO_CELLULAR_RESPONSE_TO_DNA_DAMAGE_STIMULUS | UFD1L |
| GO_CELLULAR_RESPONSE_TO_DNA_DAMAGE_STIMULUS | UHRF1 |
| GO_CELLULAR_RESPONSE_TO_DNA_DAMAGE_STIMULUS | UIMC1 |
| GO_CELLULAR_RESPONSE_TO_DNA_DAMAGE_STIMULUS | UNG |
| GO_CELLULAR_RESPONSE_TO_DNA_DAMAGE_STIMULUS | UPF1 |
| GO_CELLULAR_RESPONSE_TO_DNA_DAMAGE_STIMULUS | USP1 |
| GO_CELLULAR_RESPONSE_TO_DNA_DAMAGE_STIMULUS | USP10 |
| GO_CELLULAR_RESPONSE_TO_DNA_DAMAGE_STIMULUS | USP16 |
| GO_CELLULAR_RESPONSE_TO_DNA_DAMAGE_STIMULUS | USP28 |
| GO_CELLULAR_RESPONSE_TO_DNA_DAMAGE_STIMULUS | USP3 |
| GO_CELLULAR_RESPONSE_TO_DNA_DAMAGE_STIMULUS | USP43 |
| GO_CELLULAR_RESPONSE_TO_DNA_DAMAGE_STIMULUS | USP45 |
| GO_CELLULAR_RESPONSE_TO_DNA_DAMAGE_STIMULUS | USP47 |
| GO_CELLULAR_RESPONSE_TO_DNA_DAMAGE_STIMULUS | USP7 |
| GO_CELLULAR_RESPONSE_TO_DNA_DAMAGE_STIMULUS | UVRAG |
| GO_CELLULAR_RESPONSE_TO_DNA_DAMAGE_STIMULUS | VAV3 |

| | |
|---|---|
| GO_CELLULAR_RESPONSE_TO_DNA_DAMAGE_STIMULUS | VCP |
| GO_CELLULAR_RESPONSE_TO_DNA_DAMAGE_STIMULUS | WAC |
| GO_CELLULAR_RESPONSE_TO_DNA_DAMAGE_STIMULUS | WDR33 |
| GO_CELLULAR_RESPONSE_TO_DNA_DAMAGE_STIMULUS | WDR48 |
| GO_CELLULAR_RESPONSE_TO_DNA_DAMAGE_STIMULUS | WDR76 |
| GO_CELLULAR_RESPONSE_TO_DNA_DAMAGE_STIMULUS | WHSC1 |
| GO_CELLULAR_RESPONSE_TO_DNA_DAMAGE_STIMULUS | WNT1 |
| GO_CELLULAR_RESPONSE_TO_DNA_DAMAGE_STIMULUS | WRN |
| GO_CELLULAR_RESPONSE_TO_DNA_DAMAGE_STIMULUS | WRNIP1 |
| GO_CELLULAR_RESPONSE_TO_DNA_DAMAGE_STIMULUS | XAB2 |
| GO_CELLULAR_RESPONSE_TO_DNA_DAMAGE_STIMULUS | XIAP |
| GO_CELLULAR_RESPONSE_TO_DNA_DAMAGE_STIMULUS | XPA |
| GO_CELLULAR_RESPONSE_TO_DNA_DAMAGE_STIMULUS | XPC |
| GO_CELLULAR_RESPONSE_TO_DNA_DAMAGE_STIMULUS | XRCC1 |
| GO_CELLULAR_RESPONSE_TO_DNA_DAMAGE_STIMULUS | XRCC2 |
| GO_CELLULAR_RESPONSE_TO_DNA_DAMAGE_STIMULUS | XRCC3 |
| GO_CELLULAR_RESPONSE_TO_DNA_DAMAGE_STIMULUS | XRCC4 |
| GO_CELLULAR_RESPONSE_TO_DNA_DAMAGE_STIMULUS | XRCC5 |
| GO_CELLULAR_RESPONSE_TO_DNA_DAMAGE_STIMULUS | XRCC6 |
| GO_CELLULAR_RESPONSE_TO_DNA_DAMAGE_STIMULUS | XRCC6BP1 |
| GO_CELLULAR_RESPONSE_TO_DNA_DAMAGE_STIMULUS | YAP1 |
| GO_CELLULAR_RESPONSE_TO_DNA_DAMAGE_STIMULUS | YY1 |
| GO_CELLULAR_RESPONSE_TO_DNA_DAMAGE_STIMULUS | ZAK |
| GO_CELLULAR_RESPONSE_TO_DNA_DAMAGE_STIMULUS | ZBTB1 |
| GO_CELLULAR_RESPONSE_TO_DNA_DAMAGE_STIMULUS | ZBTB32 |
| GO_CELLULAR_RESPONSE_TO_DNA_DAMAGE_STIMULUS | ZBTB38 |
| GO_CELLULAR_RESPONSE_TO_DNA_DAMAGE_STIMULUS | ZBTB4 |
| GO_CELLULAR_RESPONSE_TO_DNA_DAMAGE_STIMULUS | ZBTB40 |
| GO_CELLULAR_RESPONSE_TO_DNA_DAMAGE_STIMULUS | ZFYVE26 |
| GO_CELLULAR_RESPONSE_TO_DNA_DAMAGE_STIMULUS | ZMAT3 |
| GO_CELLULAR_RESPONSE_TO_DNA_DAMAGE_STIMULUS | ZNF385A |
| GO_CELLULAR_RESPONSE_TO_DNA_DAMAGE_STIMULUS | ZNF830 |
| GO_CELLULAR_RESPONSE_TO_DNA_DAMAGE_STIMULUS | ZRANB3 |
| GO_CELLULAR_RESPONSE_TO_DNA_DAMAGE_STIMULUS | ZSWIM7 |
| GO_INNATE_IMMUNE_RESPONSE | ABL1 |
| GO_INNATE_IMMUNE_RESPONSE | ABL2 |
| GO_INNATE_IMMUNE_RESPONSE | ADAM15 |
| GO_INNATE_IMMUNE_RESPONSE | ADAMTS13 |
| GO_INNATE_IMMUNE_RESPONSE | ADAR |
| GO_INNATE_IMMUNE_RESPONSE | ADARB1 |
| GO_INNATE_IMMUNE_RESPONSE | AGER |
| GO_INNATE_IMMUNE_RESPONSE | AIF1 |
| GO_INNATE_IMMUNE_RESPONSE | AIM2 |
| GO_INNATE_IMMUNE_RESPONSE | AKAP8 |
| GO_INNATE_IMMUNE_RESPONSE | AKIRIN2 |
| GO_INNATE_IMMUNE_RESPONSE | ANG |
| GO_INNATE_IMMUNE_RESPONSE | ANKHD1 |

| | |
|---|---|
| GO_INNATE_IMMUNE_RESPONSE | ANKRD17 |
| GO_INNATE_IMMUNE_RESPONSE | ANXA1 |
| GO_INNATE_IMMUNE_RESPONSE | APCS |
| GO_INNATE_IMMUNE_RESPONSE | APOA4 |
| GO_INNATE_IMMUNE_RESPONSE | APOBEC3A |
| GO_INNATE_IMMUNE_RESPONSE | APOBEC3B |
| GO_INNATE_IMMUNE_RESPONSE | APOBEC3C |
| GO_INNATE_IMMUNE_RESPONSE | APOBEC3D |
| GO_INNATE_IMMUNE_RESPONSE | APOBEC3F |
| GO_INNATE_IMMUNE_RESPONSE | APOBEC3G |
| GO_INNATE_IMMUNE_RESPONSE | APOBEC3H |
| GO_INNATE_IMMUNE_RESPONSE | APOL1 |
| GO_INNATE_IMMUNE_RESPONSE | APP |
| GO_INNATE_IMMUNE_RESPONSE | AQP4 |
| GO_INNATE_IMMUNE_RESPONSE | ARHGEF2 |
| GO_INNATE_IMMUNE_RESPONSE | ASS1 |
| GO_INNATE_IMMUNE_RESPONSE | ATG12 |
| GO_INNATE_IMMUNE_RESPONSE | ATG5 |
| GO_INNATE_IMMUNE_RESPONSE | AXL |
| GO_INNATE_IMMUNE_RESPONSE | B2M |
| GO_INNATE_IMMUNE_RESPONSE | BCL10 |
| GO_INNATE_IMMUNE_RESPONSE | BLK |
| GO_INNATE_IMMUNE_RESPONSE | BMX |
| GO_INNATE_IMMUNE_RESPONSE | BPIFA1 |
| GO_INNATE_IMMUNE_RESPONSE | BPIFB1 |
| GO_INNATE_IMMUNE_RESPONSE | BPIFB3 |
| GO_INNATE_IMMUNE_RESPONSE | BST2 |
| GO_INNATE_IMMUNE_RESPONSE | BTK |
| GO_INNATE_IMMUNE_RESPONSE | C19orf29 |
| GO_INNATE_IMMUNE_RESPONSE | C19orf66 |
| GO_INNATE_IMMUNE_RESPONSE | C1QA |
| GO_INNATE_IMMUNE_RESPONSE | C1QB |
| GO_INNATE_IMMUNE_RESPONSE | C1QBP |
| GO_INNATE_IMMUNE_RESPONSE | C1QC |
| GO_INNATE_IMMUNE_RESPONSE | C1R |
| GO_INNATE_IMMUNE_RESPONSE | C1RL |
| GO_INNATE_IMMUNE_RESPONSE | C1S |
| GO_INNATE_IMMUNE_RESPONSE | C2 |
| GO_INNATE_IMMUNE_RESPONSE | C3 |
| GO_INNATE_IMMUNE_RESPONSE | C4A |
| GO_INNATE_IMMUNE_RESPONSE | C4B |
| GO_INNATE_IMMUNE_RESPONSE | C4BPA |
| GO_INNATE_IMMUNE_RESPONSE | C4BPB |
| GO_INNATE_IMMUNE_RESPONSE | C5 |
| GO_INNATE_IMMUNE_RESPONSE | C6 |
| GO_INNATE_IMMUNE_RESPONSE | C6orf57 |
| GO_INNATE_IMMUNE_RESPONSE | C7 |

| | |
|---|---|
| GO_INNATE_IMMUNE_RESPONSE | C8A |
| GO_INNATE_IMMUNE_RESPONSE | C8B |
| GO_INNATE_IMMUNE_RESPONSE | C8G |
| GO_INNATE_IMMUNE_RESPONSE | C9 |
| GO_INNATE_IMMUNE_RESPONSE | CALCA |
| GO_INNATE_IMMUNE_RESPONSE | CALCOCO2 |
| GO_INNATE_IMMUNE_RESPONSE | CAMK2A |
| GO_INNATE_IMMUNE_RESPONSE | CAMK2B |
| GO_INNATE_IMMUNE_RESPONSE | CAMK2D |
| GO_INNATE_IMMUNE_RESPONSE | CAMK2G |
| GO_INNATE_IMMUNE_RESPONSE | CAMP |
| GO_INNATE_IMMUNE_RESPONSE | CAPZA1 |
| GO_INNATE_IMMUNE_RESPONSE | CAPZA2 |
| GO_INNATE_IMMUNE_RESPONSE | CARD9 |
| GO_INNATE_IMMUNE_RESPONSE | CASP4 |
| GO_INNATE_IMMUNE_RESPONSE | CCL1 |
| GO_INNATE_IMMUNE_RESPONSE | CCL11 |
| GO_INNATE_IMMUNE_RESPONSE | CCL13 |
| GO_INNATE_IMMUNE_RESPONSE | CCL14 |
| GO_INNATE_IMMUNE_RESPONSE | CCL15 |
| GO_INNATE_IMMUNE_RESPONSE | CCL16 |
| GO_INNATE_IMMUNE_RESPONSE | CCL17 |
| GO_INNATE_IMMUNE_RESPONSE | CCL18 |
| GO_INNATE_IMMUNE_RESPONSE | CCL19 |
| GO_INNATE_IMMUNE_RESPONSE | CCL2 |
| GO_INNATE_IMMUNE_RESPONSE | CCL20 |
| GO_INNATE_IMMUNE_RESPONSE | CCL21 |
| GO_INNATE_IMMUNE_RESPONSE | CCL22 |
| GO_INNATE_IMMUNE_RESPONSE | CCL23 |
| GO_INNATE_IMMUNE_RESPONSE | CCL24 |
| GO_INNATE_IMMUNE_RESPONSE | CCL25 |
| GO_INNATE_IMMUNE_RESPONSE | CCL26 |
| GO_INNATE_IMMUNE_RESPONSE | CCL3 |
| GO_INNATE_IMMUNE_RESPONSE | CCL3L1 |
| GO_INNATE_IMMUNE_RESPONSE | CCL3L3 |
| GO_INNATE_IMMUNE_RESPONSE | CCL4 |
| GO_INNATE_IMMUNE_RESPONSE | CCL4L2 |
| GO_INNATE_IMMUNE_RESPONSE | CCL5 |
| GO_INNATE_IMMUNE_RESPONSE | CCL7 |
| GO_INNATE_IMMUNE_RESPONSE | CCL8 |
| GO_INNATE_IMMUNE_RESPONSE | CD14 |
| GO_INNATE_IMMUNE_RESPONSE | CD180 |
| GO_INNATE_IMMUNE_RESPONSE | CD1D |
| GO_INNATE_IMMUNE_RESPONSE | CD209 |
| GO_INNATE_IMMUNE_RESPONSE | CD244 |
| GO_INNATE_IMMUNE_RESPONSE | CD300E |
| GO_INNATE_IMMUNE_RESPONSE | CD300LB |

| | |
|---|---|
| GO_INNATE_IMMUNE_RESPONSE | CD44 |
| GO_INNATE_IMMUNE_RESPONSE | CD46 |
| GO_INNATE_IMMUNE_RESPONSE | CD55 |
| GO_INNATE_IMMUNE_RESPONSE | CD58 |
| GO_INNATE_IMMUNE_RESPONSE | CD6 |
| GO_INNATE_IMMUNE_RESPONSE | CD84 |
| GO_INNATE_IMMUNE_RESPONSE | CD86 |
| GO_INNATE_IMMUNE_RESPONSE | CEBPG |
| GO_INNATE_IMMUNE_RESPONSE | CFB |
| GO_INNATE_IMMUNE_RESPONSE | CFD |
| GO_INNATE_IMMUNE_RESPONSE | CFH |
| GO_INNATE_IMMUNE_RESPONSE | CFHR5 |
| GO_INNATE_IMMUNE_RESPONSE | CFI |
| GO_INNATE_IMMUNE_RESPONSE | CFP |
| GO_INNATE_IMMUNE_RESPONSE | CHGA |
| GO_INNATE_IMMUNE_RESPONSE | CHID1 |
| GO_INNATE_IMMUNE_RESPONSE | CHUK |
| GO_INNATE_IMMUNE_RESPONSE | CIITA |
| GO_INNATE_IMMUNE_RESPONSE | CITED1 |
| GO_INNATE_IMMUNE_RESPONSE | CLEC10A |
| GO_INNATE_IMMUNE_RESPONSE | CLEC2A |
| GO_INNATE_IMMUNE_RESPONSE | CLEC4A |
| GO_INNATE_IMMUNE_RESPONSE | CLEC4C |
| GO_INNATE_IMMUNE_RESPONSE | CLEC4D |
| GO_INNATE_IMMUNE_RESPONSE | CLEC4E |
| GO_INNATE_IMMUNE_RESPONSE | CLEC4M |
| GO_INNATE_IMMUNE_RESPONSE | CLEC5A |
| GO_INNATE_IMMUNE_RESPONSE | CLEC6A |
| GO_INNATE_IMMUNE_RESPONSE | CLEC7A |
| GO_INNATE_IMMUNE_RESPONSE | CLU |
| GO_INNATE_IMMUNE_RESPONSE | CNPY3 |
| GO_INNATE_IMMUNE_RESPONSE | COLEC12 |
| GO_INNATE_IMMUNE_RESPONSE | CORO1A |
| GO_INNATE_IMMUNE_RESPONSE | CR1 |
| GO_INNATE_IMMUNE_RESPONSE | CR2 |
| GO_INNATE_IMMUNE_RESPONSE | CRCP |
| GO_INNATE_IMMUNE_RESPONSE | CRISP3 |
| GO_INNATE_IMMUNE_RESPONSE | CSF1 |
| GO_INNATE_IMMUNE_RESPONSE | CSF1R |
| GO_INNATE_IMMUNE_RESPONSE | CSK |
| GO_INNATE_IMMUNE_RESPONSE | CX3CL1 |
| GO_INNATE_IMMUNE_RESPONSE | CXCL16 |
| GO_INNATE_IMMUNE_RESPONSE | CYBA |
| GO_INNATE_IMMUNE_RESPONSE | CYBB |
| GO_INNATE_IMMUNE_RESPONSE | CYP27B1 |
| GO_INNATE_IMMUNE_RESPONSE | DAB2IP |
| GO_INNATE_IMMUNE_RESPONSE | DAK |

| | |
|---|---|
| GO_INNATE_IMMUNE_RESPONSE | DAPK1 |
| GO_INNATE_IMMUNE_RESPONSE | DAPK3 |
| GO_INNATE_IMMUNE_RESPONSE | DDX3X |
| GO_INNATE_IMMUNE_RESPONSE | DDX58 |
| GO_INNATE_IMMUNE_RESPONSE | DDX60 |
| GO_INNATE_IMMUNE_RESPONSE | DEFA1 |
| GO_INNATE_IMMUNE_RESPONSE | DEFA1B |
| GO_INNATE_IMMUNE_RESPONSE | DEFA3 |
| GO_INNATE_IMMUNE_RESPONSE | DEFA4 |
| GO_INNATE_IMMUNE_RESPONSE | DEFA5 |
| GO_INNATE_IMMUNE_RESPONSE | DEFA6 |
| GO_INNATE_IMMUNE_RESPONSE | DEFB1 |
| GO_INNATE_IMMUNE_RESPONSE | DEFB103A |
| GO_INNATE_IMMUNE_RESPONSE | DEFB103B |
| GO_INNATE_IMMUNE_RESPONSE | DEFB104A |
| GO_INNATE_IMMUNE_RESPONSE | DEFB104B |
| GO_INNATE_IMMUNE_RESPONSE | DEFB105A |
| GO_INNATE_IMMUNE_RESPONSE | DEFB105B |
| GO_INNATE_IMMUNE_RESPONSE | DEFB106A |
| GO_INNATE_IMMUNE_RESPONSE | DEFB106B |
| GO_INNATE_IMMUNE_RESPONSE | DEFB108B |
| GO_INNATE_IMMUNE_RESPONSE | DEFB108P1 |
| GO_INNATE_IMMUNE_RESPONSE | DEFB110 |
| GO_INNATE_IMMUNE_RESPONSE | DEFB112 |
| GO_INNATE_IMMUNE_RESPONSE | DEFB115 |
| GO_INNATE_IMMUNE_RESPONSE | DEFB116 |
| GO_INNATE_IMMUNE_RESPONSE | DEFB118 |
| GO_INNATE_IMMUNE_RESPONSE | DEFB119 |
| GO_INNATE_IMMUNE_RESPONSE | DEFB121 |
| GO_INNATE_IMMUNE_RESPONSE | DEFB123 |
| GO_INNATE_IMMUNE_RESPONSE | DEFB124 |
| GO_INNATE_IMMUNE_RESPONSE | DEFB125 |
| GO_INNATE_IMMUNE_RESPONSE | DEFB126 |
| GO_INNATE_IMMUNE_RESPONSE | DEFB127 |
| GO_INNATE_IMMUNE_RESPONSE | DEFB128 |
| GO_INNATE_IMMUNE_RESPONSE | DEFB129 |
| GO_INNATE_IMMUNE_RESPONSE | DEFB131 |
| GO_INNATE_IMMUNE_RESPONSE | DEFB132 |
| GO_INNATE_IMMUNE_RESPONSE | DEFB133 |
| GO_INNATE_IMMUNE_RESPONSE | DEFB134 |
| GO_INNATE_IMMUNE_RESPONSE | DEFB135 |
| GO_INNATE_IMMUNE_RESPONSE | DEFB4A |
| GO_INNATE_IMMUNE_RESPONSE | DHX58 |
| GO_INNATE_IMMUNE_RESPONSE | DMBT1 |
| GO_INNATE_IMMUNE_RESPONSE | DNAJA3 |
| GO_INNATE_IMMUNE_RESPONSE | ECSIT |
| GO_INNATE_IMMUNE_RESPONSE | EDN1 |

| | |
|---|---|
| GO_INNATE_IMMUNE_RESPONSE | EGR1 |
| GO_INNATE_IMMUNE_RESPONSE | EIF2AK2 |
| GO_INNATE_IMMUNE_RESPONSE | ELF4 |
| GO_INNATE_IMMUNE_RESPONSE | EPRS |
| GO_INNATE_IMMUNE_RESPONSE | F12 |
| GO_INNATE_IMMUNE_RESPONSE | F2RL1 |
| GO_INNATE_IMMUNE_RESPONSE | FADD |
| GO_INNATE_IMMUNE_RESPONSE | FAM105B |
| GO_INNATE_IMMUNE_RESPONSE | FAU |
| GO_INNATE_IMMUNE_RESPONSE | FBXO9 |
| GO_INNATE_IMMUNE_RESPONSE | FCER1G |
| GO_INNATE_IMMUNE_RESPONSE | FCGR1A |
| GO_INNATE_IMMUNE_RESPONSE | FCGR1B |
| GO_INNATE_IMMUNE_RESPONSE | FCN1 |
| GO_INNATE_IMMUNE_RESPONSE | FCN2 |
| GO_INNATE_IMMUNE_RESPONSE | FCN3 |
| GO_INNATE_IMMUNE_RESPONSE | FER |
| GO_INNATE_IMMUNE_RESPONSE | FES |
| GO_INNATE_IMMUNE_RESPONSE | FGA |
| GO_INNATE_IMMUNE_RESPONSE | FGB |
| GO_INNATE_IMMUNE_RESPONSE | FGR |
| GO_INNATE_IMMUNE_RESPONSE | FRK |
| GO_INNATE_IMMUNE_RESPONSE | FYN |
| GO_INNATE_IMMUNE_RESPONSE | GAPDH |
| GO_INNATE_IMMUNE_RESPONSE | GATA3 |
| GO_INNATE_IMMUNE_RESPONSE | GBP1 |
| GO_INNATE_IMMUNE_RESPONSE | GBP2 |
| GO_INNATE_IMMUNE_RESPONSE | GBP5 |
| GO_INNATE_IMMUNE_RESPONSE | GBP6 |
| GO_INNATE_IMMUNE_RESPONSE | GCH1 |
| GO_INNATE_IMMUNE_RESPONSE | GPER |
| GO_INNATE_IMMUNE_RESPONSE | GSDMD |
| GO_INNATE_IMMUNE_RESPONSE | GZMB |
| GO_INNATE_IMMUNE_RESPONSE | GZMM |
| GO_INNATE_IMMUNE_RESPONSE | H2BFS |
| GO_INNATE_IMMUNE_RESPONSE | HAVCR2 |
| GO_INNATE_IMMUNE_RESPONSE | HCK |
| GO_INNATE_IMMUNE_RESPONSE | HERC5 |
| GO_INNATE_IMMUNE_RESPONSE | HIST1H2BC |
| GO_INNATE_IMMUNE_RESPONSE | HIST1H2BE |
| GO_INNATE_IMMUNE_RESPONSE | HIST1H2BF |
| GO_INNATE_IMMUNE_RESPONSE | HIST1H2BG |
| GO_INNATE_IMMUNE_RESPONSE | HIST1H2BI |
| GO_INNATE_IMMUNE_RESPONSE | HIST1H2BJ |
| GO_INNATE_IMMUNE_RESPONSE | HIST1H2BK |
| GO_INNATE_IMMUNE_RESPONSE | HIST2H2BE |
| GO_INNATE_IMMUNE_RESPONSE | HLA-A |

| | |
|---|---|
| GO_INNATE_IMMUNE_RESPONSE | HLA-B |
| GO_INNATE_IMMUNE_RESPONSE | HLA-C |
| GO_INNATE_IMMUNE_RESPONSE | HLA-DPA1 |
| GO_INNATE_IMMUNE_RESPONSE | HLA-DPB1 |
| GO_INNATE_IMMUNE_RESPONSE | HLA-DQA1 |
| GO_INNATE_IMMUNE_RESPONSE | HLA-DQA2 |
| GO_INNATE_IMMUNE_RESPONSE | HLA-DQB1 |
| GO_INNATE_IMMUNE_RESPONSE | HLA-DQB2 |
| GO_INNATE_IMMUNE_RESPONSE | HLA-DRA |
| GO_INNATE_IMMUNE_RESPONSE | HLA-DRB1 |
| GO_INNATE_IMMUNE_RESPONSE | HLA-DRB3 |
| GO_INNATE_IMMUNE_RESPONSE | HLA-DRB4 |
| GO_INNATE_IMMUNE_RESPONSE | HLA-DRB5 |
| GO_INNATE_IMMUNE_RESPONSE | HLA-E |
| GO_INNATE_IMMUNE_RESPONSE | HLA-F |
| GO_INNATE_IMMUNE_RESPONSE | HLA-G |
| GO_INNATE_IMMUNE_RESPONSE | HLA-H |
| GO_INNATE_IMMUNE_RESPONSE | HMGB1 |
| GO_INNATE_IMMUNE_RESPONSE | HMGB2 |
| GO_INNATE_IMMUNE_RESPONSE | HMGB3 |
| GO_INNATE_IMMUNE_RESPONSE | ICAM1 |
| GO_INNATE_IMMUNE_RESPONSE | IFI16 |
| GO_INNATE_IMMUNE_RESPONSE | IFI27 |
| GO_INNATE_IMMUNE_RESPONSE | IFI30 |
| GO_INNATE_IMMUNE_RESPONSE | IFI35 |
| GO_INNATE_IMMUNE_RESPONSE | IFI6 |
| GO_INNATE_IMMUNE_RESPONSE | IFIH1 |
| GO_INNATE_IMMUNE_RESPONSE | IFIT1 |
| GO_INNATE_IMMUNE_RESPONSE | IFIT2 |
| GO_INNATE_IMMUNE_RESPONSE | IFIT3 |
| GO_INNATE_IMMUNE_RESPONSE | IFIT5 |
| GO_INNATE_IMMUNE_RESPONSE | IFITM1 |
| GO_INNATE_IMMUNE_RESPONSE | IFITM2 |
| GO_INNATE_IMMUNE_RESPONSE | IFITM3 |
| GO_INNATE_IMMUNE_RESPONSE | IFNA1 |
| GO_INNATE_IMMUNE_RESPONSE | IFNA10 |
| GO_INNATE_IMMUNE_RESPONSE | IFNA13 |
| GO_INNATE_IMMUNE_RESPONSE | IFNA14 |
| GO_INNATE_IMMUNE_RESPONSE | IFNA16 |
| GO_INNATE_IMMUNE_RESPONSE | IFNA17 |
| GO_INNATE_IMMUNE_RESPONSE | IFNA2 |
| GO_INNATE_IMMUNE_RESPONSE | IFNA21 |
| GO_INNATE_IMMUNE_RESPONSE | IFNA4 |
| GO_INNATE_IMMUNE_RESPONSE | IFNA5 |
| GO_INNATE_IMMUNE_RESPONSE | IFNA6 |
| GO_INNATE_IMMUNE_RESPONSE | IFNA7 |
| GO_INNATE_IMMUNE_RESPONSE | IFNA8 |

| | |
|---|---|
| GO_INNATE_IMMUNE_RESPONSE | IFNAR1 |
| GO_INNATE_IMMUNE_RESPONSE | IFNAR2 |
| GO_INNATE_IMMUNE_RESPONSE | IFNB1 |
| GO_INNATE_IMMUNE_RESPONSE | IFNE |
| GO_INNATE_IMMUNE_RESPONSE | IFNG |
| GO_INNATE_IMMUNE_RESPONSE | IFNGR1 |
| GO_INNATE_IMMUNE_RESPONSE | IFNGR2 |
| GO_INNATE_IMMUNE_RESPONSE | IFNW1 |
| GO_INNATE_IMMUNE_RESPONSE | IGHA1 |
| GO_INNATE_IMMUNE_RESPONSE | IGHA2 |
| GO_INNATE_IMMUNE_RESPONSE | IGHD |
| GO_INNATE_IMMUNE_RESPONSE | IGHE |
| GO_INNATE_IMMUNE_RESPONSE | IGHG1 |
| GO_INNATE_IMMUNE_RESPONSE | IGHG2 |
| GO_INNATE_IMMUNE_RESPONSE | IGHG3 |
| GO_INNATE_IMMUNE_RESPONSE | IGHG4 |
| GO_INNATE_IMMUNE_RESPONSE | IGHM |
| GO_INNATE_IMMUNE_RESPONSE | IGHV1OR21-1 |
| GO_INNATE_IMMUNE_RESPONSE | IGHV3-23 |
| GO_INNATE_IMMUNE_RESPONSE | IGHV4OR15-8 |
| GO_INNATE_IMMUNE_RESPONSE | IGJ |
| GO_INNATE_IMMUNE_RESPONSE | IGKC |
| GO_INNATE_IMMUNE_RESPONSE | IGLC1 |
| GO_INNATE_IMMUNE_RESPONSE | IGLC2 |
| GO_INNATE_IMMUNE_RESPONSE | IGLC3 |
| GO_INNATE_IMMUNE_RESPONSE | IGLC6 |
| GO_INNATE_IMMUNE_RESPONSE | IGLC7 |
| GO_INNATE_IMMUNE_RESPONSE | IGLL1 |
| GO_INNATE_IMMUNE_RESPONSE | IGLL5 |
| GO_INNATE_IMMUNE_RESPONSE | IKBKB |
| GO_INNATE_IMMUNE_RESPONSE | IKBKE |
| GO_INNATE_IMMUNE_RESPONSE | IKBKG |
| GO_INNATE_IMMUNE_RESPONSE | IL12B |
| GO_INNATE_IMMUNE_RESPONSE | IL12RB1 |
| GO_INNATE_IMMUNE_RESPONSE | IL1RAP |
| GO_INNATE_IMMUNE_RESPONSE | IL1RL2 |
| GO_INNATE_IMMUNE_RESPONSE | IL23A |
| GO_INNATE_IMMUNE_RESPONSE | IL23R |
| GO_INNATE_IMMUNE_RESPONSE | IL27 |
| GO_INNATE_IMMUNE_RESPONSE | IL28A |
| GO_INNATE_IMMUNE_RESPONSE | IL28B |
| GO_INNATE_IMMUNE_RESPONSE | IL28RA |
| GO_INNATE_IMMUNE_RESPONSE | IL29 |
| GO_INNATE_IMMUNE_RESPONSE | IL34 |
| GO_INNATE_IMMUNE_RESPONSE | IL36A |
| GO_INNATE_IMMUNE_RESPONSE | IL36B |
| GO_INNATE_IMMUNE_RESPONSE | IL36G |

| | |
|---|---|
| GO_INNATE_IMMUNE_RESPONSE | IL36RN |
| GO_INNATE_IMMUNE_RESPONSE | IL4 |
| GO_INNATE_IMMUNE_RESPONSE | IP6K2 |
| GO_INNATE_IMMUNE_RESPONSE | IPO7 |
| GO_INNATE_IMMUNE_RESPONSE | IRAK1 |
| GO_INNATE_IMMUNE_RESPONSE | IRAK4 |
| GO_INNATE_IMMUNE_RESPONSE | IRF1 |
| GO_INNATE_IMMUNE_RESPONSE | IRF2 |
| GO_INNATE_IMMUNE_RESPONSE | IRF3 |
| GO_INNATE_IMMUNE_RESPONSE | IRF4 |
| GO_INNATE_IMMUNE_RESPONSE | IRF5 |
| GO_INNATE_IMMUNE_RESPONSE | IRF6 |
| GO_INNATE_IMMUNE_RESPONSE | IRF7 |
| GO_INNATE_IMMUNE_RESPONSE | IRF8 |
| GO_INNATE_IMMUNE_RESPONSE | IRF9 |
| GO_INNATE_IMMUNE_RESPONSE | IRG1 |
| GO_INNATE_IMMUNE_RESPONSE | IRGM |
| GO_INNATE_IMMUNE_RESPONSE | ISG15 |
| GO_INNATE_IMMUNE_RESPONSE | ISG20 |
| GO_INNATE_IMMUNE_RESPONSE | ITCH |
| GO_INNATE_IMMUNE_RESPONSE | ITK |
| GO_INNATE_IMMUNE_RESPONSE | JAK1 |
| GO_INNATE_IMMUNE_RESPONSE | JAK2 |
| GO_INNATE_IMMUNE_RESPONSE | JAK3 |
| GO_INNATE_IMMUNE_RESPONSE | KIR2DS1 |
| GO_INNATE_IMMUNE_RESPONSE | KIR2DS2 |
| GO_INNATE_IMMUNE_RESPONSE | KIR2DS4 |
| GO_INNATE_IMMUNE_RESPONSE | KIR2DS5 |
| GO_INNATE_IMMUNE_RESPONSE | KIR3DL1 |
| GO_INNATE_IMMUNE_RESPONSE | KIR3DS1 |
| GO_INNATE_IMMUNE_RESPONSE | KLRC2 |
| GO_INNATE_IMMUNE_RESPONSE | KLRC4-KLRK1 |
| GO_INNATE_IMMUNE_RESPONSE | KLRD1 |
| GO_INNATE_IMMUNE_RESPONSE | KLRF2 |
| GO_INNATE_IMMUNE_RESPONSE | KLRG1 |
| GO_INNATE_IMMUNE_RESPONSE | KLRK1 |
| GO_INNATE_IMMUNE_RESPONSE | KRT1 |
| GO_INNATE_IMMUNE_RESPONSE | KRT16 |
| GO_INNATE_IMMUNE_RESPONSE | KYNU |
| GO_INNATE_IMMUNE_RESPONSE | LBP |
| GO_INNATE_IMMUNE_RESPONSE | LCK |
| GO_INNATE_IMMUNE_RESPONSE | LCN2 |
| GO_INNATE_IMMUNE_RESPONSE | LGALS3 |
| GO_INNATE_IMMUNE_RESPONSE | LGALS9 |
| GO_INNATE_IMMUNE_RESPONSE | LGR4 |
| GO_INNATE_IMMUNE_RESPONSE | LILRA5 |
| GO_INNATE_IMMUNE_RESPONSE | LRRC33 |

| | |
|---|---|
| GO_INNATE_IMMUNE_RESPONSE | LTF |
| GO_INNATE_IMMUNE_RESPONSE | LY86 |
| GO_INNATE_IMMUNE_RESPONSE | LY9 |
| GO_INNATE_IMMUNE_RESPONSE | LY96 |
| GO_INNATE_IMMUNE_RESPONSE | LYN |
| GO_INNATE_IMMUNE_RESPONSE | LYST |
| GO_INNATE_IMMUNE_RESPONSE | MALT1 |
| GO_INNATE_IMMUNE_RESPONSE | MAP3K5 |
| GO_INNATE_IMMUNE_RESPONSE | MAP4K2 |
| GO_INNATE_IMMUNE_RESPONSE | MARCO |
| GO_INNATE_IMMUNE_RESPONSE | MASP1 |
| GO_INNATE_IMMUNE_RESPONSE | MASP2 |
| GO_INNATE_IMMUNE_RESPONSE | MATK |
| GO_INNATE_IMMUNE_RESPONSE | MAVS |
| GO_INNATE_IMMUNE_RESPONSE | MB21D1 |
| GO_INNATE_IMMUNE_RESPONSE | MBL2 |
| GO_INNATE_IMMUNE_RESPONSE | MEFV |
| GO_INNATE_IMMUNE_RESPONSE | MICA |
| GO_INNATE_IMMUNE_RESPONSE | MICB |
| GO_INNATE_IMMUNE_RESPONSE | MID1 |
| GO_INNATE_IMMUNE_RESPONSE | MID2 |
| GO_INNATE_IMMUNE_RESPONSE | MIF |
| GO_INNATE_IMMUNE_RESPONSE | MR1 |
| GO_INNATE_IMMUNE_RESPONSE | MRC1 |
| GO_INNATE_IMMUNE_RESPONSE | MST1R |
| GO_INNATE_IMMUNE_RESPONSE | MT2A |
| GO_INNATE_IMMUNE_RESPONSE | MX1 |
| GO_INNATE_IMMUNE_RESPONSE | MX2 |
| GO_INNATE_IMMUNE_RESPONSE | MYD88 |
| GO_INNATE_IMMUNE_RESPONSE | NAIP |
| GO_INNATE_IMMUNE_RESPONSE | NCAM1 |
| GO_INNATE_IMMUNE_RESPONSE | NCF1 |
| GO_INNATE_IMMUNE_RESPONSE | NCF2 |
| GO_INNATE_IMMUNE_RESPONSE | NCR2 |
| GO_INNATE_IMMUNE_RESPONSE | NFKB1 |
| GO_INNATE_IMMUNE_RESPONSE | NFKB2 |
| GO_INNATE_IMMUNE_RESPONSE | NLRC4 |
| GO_INNATE_IMMUNE_RESPONSE | NLRC5 |
| GO_INNATE_IMMUNE_RESPONSE | NLRP1 |
| GO_INNATE_IMMUNE_RESPONSE | NLRP10 |
| GO_INNATE_IMMUNE_RESPONSE | NLRP2 |
| GO_INNATE_IMMUNE_RESPONSE | NLRP2P |
| GO_INNATE_IMMUNE_RESPONSE | NLRP3 |
| GO_INNATE_IMMUNE_RESPONSE | NLRP6 |
| GO_INNATE_IMMUNE_RESPONSE | NLRX1 |
| GO_INNATE_IMMUNE_RESPONSE | NOD1 |
| GO_INNATE_IMMUNE_RESPONSE | NOD2 |

| | |
|---|---|
| GO_INNATE_IMMUNE_RESPONSE | NOS2 |
| GO_INNATE_IMMUNE_RESPONSE | NPY |
| GO_INNATE_IMMUNE_RESPONSE | NR1H4 |
| GO_INNATE_IMMUNE_RESPONSE | NUB1 |
| GO_INNATE_IMMUNE_RESPONSE | OAS1 |
| GO_INNATE_IMMUNE_RESPONSE | OAS2 |
| GO_INNATE_IMMUNE_RESPONSE | OAS3 |
| GO_INNATE_IMMUNE_RESPONSE | OASL |
| GO_INNATE_IMMUNE_RESPONSE | PADI4 |
| GO_INNATE_IMMUNE_RESPONSE | PCBP2 |
| GO_INNATE_IMMUNE_RESPONSE | PGLYRP1 |
| GO_INNATE_IMMUNE_RESPONSE | PGLYRP2 |
| GO_INNATE_IMMUNE_RESPONSE | PGLYRP3 |
| GO_INNATE_IMMUNE_RESPONSE | PGLYRP4 |
| GO_INNATE_IMMUNE_RESPONSE | PIK3CD |
| GO_INNATE_IMMUNE_RESPONSE | PIK3CG |
| GO_INNATE_IMMUNE_RESPONSE | PLA2G1B |
| GO_INNATE_IMMUNE_RESPONSE | PML |
| GO_INNATE_IMMUNE_RESPONSE | POLR3A |
| GO_INNATE_IMMUNE_RESPONSE | POLR3B |
| GO_INNATE_IMMUNE_RESPONSE | POLR3C |
| GO_INNATE_IMMUNE_RESPONSE | POLR3D |
| GO_INNATE_IMMUNE_RESPONSE | POLR3E |
| GO_INNATE_IMMUNE_RESPONSE | POLR3F |
| GO_INNATE_IMMUNE_RESPONSE | POLR3G |
| GO_INNATE_IMMUNE_RESPONSE | POLR3H |
| GO_INNATE_IMMUNE_RESPONSE | POLR3K |
| GO_INNATE_IMMUNE_RESPONSE | PPARG |
| GO_INNATE_IMMUNE_RESPONSE | PPP1R14B |
| GO_INNATE_IMMUNE_RESPONSE | PRDX1 |
| GO_INNATE_IMMUNE_RESPONSE | PRKCD |
| GO_INNATE_IMMUNE_RESPONSE | PRKD1 |
| GO_INNATE_IMMUNE_RESPONSE | PSMB8 |
| GO_INNATE_IMMUNE_RESPONSE | PSTPIP1 |
| GO_INNATE_IMMUNE_RESPONSE | PTAFR |
| GO_INNATE_IMMUNE_RESPONSE | PTK2 |
| GO_INNATE_IMMUNE_RESPONSE | PTK2B |
| GO_INNATE_IMMUNE_RESPONSE | PTK6 |
| GO_INNATE_IMMUNE_RESPONSE | PTPN6 |
| GO_INNATE_IMMUNE_RESPONSE | PTX3 |
| GO_INNATE_IMMUNE_RESPONSE | PYCARD |
| GO_INNATE_IMMUNE_RESPONSE | PYDC1 |
| GO_INNATE_IMMUNE_RESPONSE | PYDC2 |
| GO_INNATE_IMMUNE_RESPONSE | RAB27A |
| GO_INNATE_IMMUNE_RESPONSE | RAET1E |
| GO_INNATE_IMMUNE_RESPONSE | RAET1G |
| GO_INNATE_IMMUNE_RESPONSE | RAET1L |

| | |
|---|---|
| GO_INNATE_IMMUNE_RESPONSE | REL |
| GO_INNATE_IMMUNE_RESPONSE | RELB |
| GO_INNATE_IMMUNE_RESPONSE | RIPK2 |
| GO_INNATE_IMMUNE_RESPONSE | RNASE3 |
| GO_INNATE_IMMUNE_RESPONSE | RNASE7 |
| GO_INNATE_IMMUNE_RESPONSE | RNASEL |
| GO_INNATE_IMMUNE_RESPONSE | RNF135 |
| GO_INNATE_IMMUNE_RESPONSE | RPL13A |
| GO_INNATE_IMMUNE_RESPONSE | RPL39 |
| GO_INNATE_IMMUNE_RESPONSE | RPS27A |
| GO_INNATE_IMMUNE_RESPONSE | RSAD2 |
| GO_INNATE_IMMUNE_RESPONSE | S100A12 |
| GO_INNATE_IMMUNE_RESPONSE | S100A7 |
| GO_INNATE_IMMUNE_RESPONSE | S100A8 |
| GO_INNATE_IMMUNE_RESPONSE | S100A9 |
| GO_INNATE_IMMUNE_RESPONSE | S100B |
| GO_INNATE_IMMUNE_RESPONSE | SAA1 |
| GO_INNATE_IMMUNE_RESPONSE | SAMHD1 |
| GO_INNATE_IMMUNE_RESPONSE | SARM1 |
| GO_INNATE_IMMUNE_RESPONSE | SEC14L1 |
| GO_INNATE_IMMUNE_RESPONSE | SEC61A1 |
| GO_INNATE_IMMUNE_RESPONSE | SEPX1 |
| GO_INNATE_IMMUNE_RESPONSE | SERINC3 |
| GO_INNATE_IMMUNE_RESPONSE | SERINC5 |
| GO_INNATE_IMMUNE_RESPONSE | SERPING1 |
| GO_INNATE_IMMUNE_RESPONSE | SFTPD |
| GO_INNATE_IMMUNE_RESPONSE | SH2D1A |
| GO_INNATE_IMMUNE_RESPONSE | SH2D1B |
| GO_INNATE_IMMUNE_RESPONSE | SHMT2 |
| GO_INNATE_IMMUNE_RESPONSE | SIGLEC14 |
| GO_INNATE_IMMUNE_RESPONSE | SIGLEC15 |
| GO_INNATE_IMMUNE_RESPONSE | SIGLEC16 |
| GO_INNATE_IMMUNE_RESPONSE | SIRPB1 |
| GO_INNATE_IMMUNE_RESPONSE | SIRT2 |
| GO_INNATE_IMMUNE_RESPONSE | SLAMF1 |
| GO_INNATE_IMMUNE_RESPONSE | SLAMF6 |
| GO_INNATE_IMMUNE_RESPONSE | SLAMF7 |
| GO_INNATE_IMMUNE_RESPONSE | SLC11A1 |
| GO_INNATE_IMMUNE_RESPONSE | SLC26A6 |
| GO_INNATE_IMMUNE_RESPONSE | SLC30A8 |
| GO_INNATE_IMMUNE_RESPONSE | SLPI |
| GO_INNATE_IMMUNE_RESPONSE | SNCA |
| GO_INNATE_IMMUNE_RESPONSE | SP100 |
| GO_INNATE_IMMUNE_RESPONSE | SPON2 |
| GO_INNATE_IMMUNE_RESPONSE | SRC |
| GO_INNATE_IMMUNE_RESPONSE | SRMS |
| GO_INNATE_IMMUNE_RESPONSE | SRPK1 |

| | |
|---|---|
| GO_INNATE_IMMUNE_RESPONSE | SRPK2 |
| GO_INNATE_IMMUNE_RESPONSE | SSC5D |
| GO_INNATE_IMMUNE_RESPONSE | STAR |
| GO_INNATE_IMMUNE_RESPONSE | STAT1 |
| GO_INNATE_IMMUNE_RESPONSE | STAT2 |
| GO_INNATE_IMMUNE_RESPONSE | STYK1 |
| GO_INNATE_IMMUNE_RESPONSE | SUSD4 |
| GO_INNATE_IMMUNE_RESPONSE | SYK |
| GO_INNATE_IMMUNE_RESPONSE | SYNCRIP |
| GO_INNATE_IMMUNE_RESPONSE | TAC1 |
| GO_INNATE_IMMUNE_RESPONSE | TBK1 |
| GO_INNATE_IMMUNE_RESPONSE | TBKBP1 |
| GO_INNATE_IMMUNE_RESPONSE | TDGF1 |
| GO_INNATE_IMMUNE_RESPONSE | TEC |
| GO_INNATE_IMMUNE_RESPONSE | TGFB1 |
| GO_INNATE_IMMUNE_RESPONSE | TICAM1 |
| GO_INNATE_IMMUNE_RESPONSE | TICAM2 |
| GO_INNATE_IMMUNE_RESPONSE | TIRAP |
| GO_INNATE_IMMUNE_RESPONSE | TLR1 |
| GO_INNATE_IMMUNE_RESPONSE | TLR10 |
| GO_INNATE_IMMUNE_RESPONSE | TLR2 |
| GO_INNATE_IMMUNE_RESPONSE | TLR3 |
| GO_INNATE_IMMUNE_RESPONSE | TLR4 |
| GO_INNATE_IMMUNE_RESPONSE | TLR5 |
| GO_INNATE_IMMUNE_RESPONSE | TLR6 |
| GO_INNATE_IMMUNE_RESPONSE | TLR7 |
| GO_INNATE_IMMUNE_RESPONSE | TLR8 |
| GO_INNATE_IMMUNE_RESPONSE | TLR9 |
| GO_INNATE_IMMUNE_RESPONSE | TMEM173 |
| GO_INNATE_IMMUNE_RESPONSE | TNFAIP8L2 |
| GO_INNATE_IMMUNE_RESPONSE | TNK1 |
| GO_INNATE_IMMUNE_RESPONSE | TNK2 |
| GO_INNATE_IMMUNE_RESPONSE | TOLLIP |
| GO_INNATE_IMMUNE_RESPONSE | TRAF3 |
| GO_INNATE_IMMUNE_RESPONSE | TRDC |
| GO_INNATE_IMMUNE_RESPONSE | TREM1 |
| GO_INNATE_IMMUNE_RESPONSE | TREM2 |
| GO_INNATE_IMMUNE_RESPONSE | TREML1 |
| GO_INNATE_IMMUNE_RESPONSE | TRIL |
| GO_INNATE_IMMUNE_RESPONSE | TRIM10 |
| GO_INNATE_IMMUNE_RESPONSE | TRIM11 |
| GO_INNATE_IMMUNE_RESPONSE | TRIM13 |
| GO_INNATE_IMMUNE_RESPONSE | TRIM14 |
| GO_INNATE_IMMUNE_RESPONSE | TRIM15 |
| GO_INNATE_IMMUNE_RESPONSE | TRIM21 |
| GO_INNATE_IMMUNE_RESPONSE | TRIM22 |
| GO_INNATE_IMMUNE_RESPONSE | TRIM25 |

| | |
|---|---|
| GO_INNATE_IMMUNE_RESPONSE | TRIM26 |
| GO_INNATE_IMMUNE_RESPONSE | TRIM27 |
| GO_INNATE_IMMUNE_RESPONSE | TRIM28 |
| GO_INNATE_IMMUNE_RESPONSE | TRIM31 |
| GO_INNATE_IMMUNE_RESPONSE | TRIM32 |
| GO_INNATE_IMMUNE_RESPONSE | TRIM34 |
| GO_INNATE_IMMUNE_RESPONSE | TRIM35 |
| GO_INNATE_IMMUNE_RESPONSE | TRIM38 |
| GO_INNATE_IMMUNE_RESPONSE | TRIM4 |
| GO_INNATE_IMMUNE_RESPONSE | TRIM5 |
| GO_INNATE_IMMUNE_RESPONSE | TRIM56 |
| GO_INNATE_IMMUNE_RESPONSE | TRIM62 |
| GO_INNATE_IMMUNE_RESPONSE | TRIM68 |
| GO_INNATE_IMMUNE_RESPONSE | TRIM8 |
| GO_INNATE_IMMUNE_RESPONSE | TUBB |
| GO_INNATE_IMMUNE_RESPONSE | TUBB4B |
| GO_INNATE_IMMUNE_RESPONSE | TYK2 |
| GO_INNATE_IMMUNE_RESPONSE | TYROBP |
| GO_INNATE_IMMUNE_RESPONSE | UBA52 |
| GO_INNATE_IMMUNE_RESPONSE | UBB |
| GO_INNATE_IMMUNE_RESPONSE | UBC |
| GO_INNATE_IMMUNE_RESPONSE | UBD |
| GO_INNATE_IMMUNE_RESPONSE | ULBP1 |
| GO_INNATE_IMMUNE_RESPONSE | ULBP2 |
| GO_INNATE_IMMUNE_RESPONSE | ULBP3 |
| GO_INNATE_IMMUNE_RESPONSE | UNC13D |
| GO_INNATE_IMMUNE_RESPONSE | UNC93B1 |
| GO_INNATE_IMMUNE_RESPONSE | VAMP2 |
| GO_INNATE_IMMUNE_RESPONSE | VAMP7 |
| GO_INNATE_IMMUNE_RESPONSE | VCAM1 |
| GO_INNATE_IMMUNE_RESPONSE | VIP |
| GO_INNATE_IMMUNE_RESPONSE | VNN1 |
| GO_INNATE_IMMUNE_RESPONSE | VSIG4 |
| GO_INNATE_IMMUNE_RESPONSE | WNT5A |
| GO_INNATE_IMMUNE_RESPONSE | XAF1 |
| GO_INNATE_IMMUNE_RESPONSE | XCL1 |
| GO_INNATE_IMMUNE_RESPONSE | XCL2 |
| GO_INNATE_IMMUNE_RESPONSE | YES1 |
| GO_INNATE_IMMUNE_RESPONSE | ZAP70 |
| GO_INNATE_IMMUNE_RESPONSE | ZBP1 |
| GO_INNATE_IMMUNE_RESPONSE | ZBTB1 |
| GO_INNATE_IMMUNE_RESPONSE | ZC3HAV1 |
