## Supplementary material for "Human iPSC modeling reveals mutation-specific responses to gene therapy in Best disease": SI Data File C

SI Data File C: Ranked off-target sites for sgRNAs used in this study

**Off-target for A146K sgRNA**

| Sequence | PAM | Score | Gene | Chromosome | Strand | Position | Mismatches | On-target |
|---|---|---|---|---|---|---|---|---|
| CTGCGGTGCTGACGCTGCGC | AGG | 5.146703297 | ENSG00000167995 | chr11 | -1 | 61955893 | 2 | FALSE |
| CCTGTGTCCTGACGCTGCGC | GGG | 1.442307692 | | chr20 | -1 | 50246956 | 4 | FALSE |
| CTTTGTTCATGACGCTGCGC | AGG | 0.896991699 | ENSG00000048471 | chr16 | -1 | 12524752 | 3 | FALSE |
| CTTTTGTGCCGACGTTGCGC | GGG | 0.762558049 | | chr4 | 1 | 54658402 | 3 | FALSE |
| CCGCGGTGCTGACGCTGCTC | AGG | 0.611313191 | ENSG00000090316 | chr4 | -1 | 1309855 | 4 | FALSE |
| CTGTAGTGCAGAGGCTGCGC | TGG | 0.582322973 | | chr2 | 1 | 47237604 | 4 | FALSE |
| TTTTGGTGCTGAAGCTGCCC | AAG | 0.54756383 | | chr8 | -1 | 101982967 | 3 | FALSE |
| CCATGGGGCTGGCGCTGCGC | AAG | 0.543948597 | ENSG00000266074 | chr17 | 1 | 81461477 | 4 | FALSE |
| CCCTGCTGCTGGCGCTGCGC | TGG | 0.477428291 | ENSG00000170128 | chr1 | 1 | 200873893 | 4 | FALSE |
| CTCTGCTGCCGCCGCTGCGC | GAG | 0.431824255 | ENSG00000143502 | chr1 | 1 | 223364152 | 4 | FALSE |
| CGTTGGTGGCGACGCTGCCC | CGG | 0.348889334 | ENSG00000204666 | chr19 | 1 | 50050818 | 4 | FALSE |
| CTGCGGTGCGGACGCTGGGC | CAG | 0.346494543 | ENSG00000167608 | chr19 | -1 | 54164551 | 4 | FALSE |
| CTCCGGTGCTGATGCTGCGG | GAG | 0.343553885 | ENSG00000138311 | chr10 | -1 | 62374409 | 4 | FALSE |
| CTTTGGTGCAGACGCTCCCC | GAG | 0.332111303 | | chr10 | 1 | 15762893 | 3 | FALSE |
| CTTTCAGGCTTACGCTGCGC | GGG | 0.330770661 | ENSG00000273079 | chr12 | 1 | 13563279 | 4 | FALSE |
| CTGTGGTCCTGCCGCTGCTC | TGG | 0.30076609 | ENSG00000073060 | chr12 | -1 | 124787421 | 4 | FALSE |
| CTTTGGAGAACACGCTGCGC | TAG | 0.293406436 | | chr5 | -1 | 14191003 | 4 | FALSE |
| CTGTGGTGCTGATGCTGGGC | CAG | 0.287073004 | | chr3 | 1 | 139829324 | 3 | FALSE |
| CCTTGGTGCCGACGCTGCAG | AAG | 0.271059097 | ENSG00000112782 | chr6 | -1 | 46014623 | 4 | FALSE |
| CTTTGTTGGAGCCGCTGCGC | GGG | 0.241590158 | ENSG00000111058 | chr12 | -1 | 80937330 | 4 | FALSE |
| CAGTGGTGGTGACGCTGAGC | AGG | 0.237659842 | | chr2 | -1 | 23667620 | 4 | FALSE |
| ATTTCTTGCTGACGCTGGGC | AGG | 0.233392956 | | chr9 | -1 | 35982279 | 4 | FALSE |
| GTTTGGAGCTGCTGCTGCGC | AGG | 0.228221001 | ENSG00000132382 | chr17 | 1 | 4544546 | 4 | FALSE |
| TTTTGATGCTGAGGCTGCGG | CGG | 0.221542787 | | chr7 | 1 | 36242063 | 4 | FALSE |
| GTCTGGTGCTGACGATCCGC | GAG | 0.21965615 | ENSG00000108465 | chr17 | 1 | 47973578 | 4 | FALSE |
| CTGTGGTGCGGAAGCTGCCC | AGG | 0.215507231 | | chr5 | -1 | 140677455 | 4 | FALSE |
| CCTGGGGGCTGACGCGGCGC | CGG | 0.214619615 | ENSG00000117322 | chr1 | -1 | 207454238 | 4 | FALSE |
| CCTTGCTGCTGATGCTGCGG | AGG | 0.213387838 | | chr15 | -1 | 74832344 | 4 | FALSE |
| CTTCAGTGCTTACGCTGTGC | CAG | 0.205041667 | | chr10 | -1 | 46442968 | 4 | FALSE |
| CCTGGGTGCTGGCGCTGGGC | TGG | 0.198577457 | | chr17 | 1 | 39635341 | 4 | FALSE |
| CCTTGGCGCTGATGCTTCGC | AGG | 0.19810534 | ENSG00000131759 | chr17 | -1 | 40355405 | 4 | FALSE |
| CTTTGGTGGGGCCGCTGCGG | TGG | 0.191301516 | ENSG00000279685 | chr17 | 1 | 45897419 | 4 | FALSE |
| CTGTGATGCTGGCGCTGCCC | GGG | 0.186075653 | | chr2 | 1 | 144508184 | 4 | FALSE |
| CTTGGGCGCTGAGGCTTCGC | AGG | 0.185914242 | | chr4 | 1 | 1168570 | 4 | FALSE |
| CTGTTGTGCTGACGATGCTC | TGG | 0.175464972 | | chr18 | 1 | 2857771 | 4 | FALSE |
| CTTTGCTACTGTGGCTGCGC | AAG | 0.174629839 | | chr4 | 1 | 71438039 | 4 | FALSE |
| CTTTGGGGCTGACGCAGCGG | AGG | 0.171411114 | ENSG00000072952 | chr11 | -1 | 10652138 | 3 | FALSE |
| ATGTGGTGCTGAGGCTGAGC | AAG | 0.163459897 | | chr11 | -1 | 109641742 | 4 | FALSE |
| CCTGGGTGCTGAGGCTGGGC | CAG | 0.158025 | ENSG00000139880 | chr14 | 1 | 23055784 | 4 | FALSE |
| GTTTGGGGCTGACGTTGCTC | CAG | 0.127583658 | | chr19 | -1 | 17997337 | 4 | FALSE |
| CTGTGGGGCTGGCGCTGAGC | CGG | 0.125055252 | ENSG00000135100 | chr12 | -1 | 120999611 | 4 | FALSE |
| TTTTGGTGCTGACTCTGAGC | AGG | 0.124133333 | | chr2 | -1 | 45393365 | 3 | FALSE |
| GTTTGGTGCTGCAGCTGCAC | CGG | 0.122096565 | ENSG00000125457 | chr17 | -1 | 75266944 | 4 | FALSE |
| CTCTGGAGCTGACGCTGGGG | GAG | 0.121791994 | | chr14 | -1 | 94370120 | 4 | FALSE |
| CTCTGGAGCTGACGCTGGGG | GAG | 0.121791994 | | chr14 | -1 | 94361639 | 4 | FALSE |
| TTTTGCTGCTGCCGCTGTGC | TGG | 0.1215445 | | chr5 | 1 | 179212942 | 4 | FALSE |
| CATTGGTGCAGACACTGCGG | GGG | 0.120628554 | | chr4 | 1 | 8581829 | 4 | FALSE |
| CTTTGGTGCTGGCGCTGGGT | GGG | 0.115238422 | ENSG00000258986 | chr14 | -1 | 104594792 | 3 | FALSE |
| CCTTGGTGATGAGGGTGCGC | AGG | 0.11344589 | ENSG00000052749 | chr10 | 1 | 97373604 | 4 | FALSE |
| ATGTGGTGCTGACTCTGCAC | AGG | 0.106364551 | | chr13 | -1 | 51223383 | 4 | FALSE |

**Off-target for N296H sgRNA** **Off-target for N296H sgRNA**

| Sequence | PAM | Score | Gene | Chromosome | Strand | Position | Mismatches | On-target |
|---|---|---|---|---|---|---|---|---|
| CTTCATCATCTCCAAAGGGG | AAG | 6.69014085 | | chr9 | -1 | 90546396 | 2 | FALSE |
| CAGCATGCTCTCCAAAGGGG | AAG | 4.04915253 | | chr18 | 1 | 37665117 | 2 | FALSE |
| CATGCTCCTCTCCAAAGGGC | AGG | 1.68216561 | | chr5 | 1 | 172749869 | 3 | FALSE |
| CATAATCATCTCCAAAGGGC | TAG | 1.68216561 | | chrX | 1 | 79627241 | 3 | FALSE |
| CACCCTCCTCCCCAAAGGGG | CAG | 1.56823077 | | chr2 | 1 | 74347940 | 3 | FALSE |
| GAATATCGTCTCCAAAGGGG | TAG | 1.45752075 | | chr4 | 1 | 164866636 | 4 | FALSE |
| CATTATCCTACCCAAAGGGG | AGG | 1.41367467 | | chr5 | -1 | 37917393 | 3 | FALSE |
| CGTCATCCTGTACAAAGGGG | AGG | 1.39992 | | chr13 | -1 | 32175536 | 3 | FALSE |
| CTCCTTCCTTTCCAAAGGGG | CAG | 1.38846679 | ENSG000001 | chr14 | -1 | 21405293 | 4 | FALSE |
| ATACCTCCTCTCCAAAGGGG | TAG | 1.35622587 | | chr3 | 1 | 113192849 | 4 | FALSE |
| CAGCATTCTCCCCAAAGGGG | CAG | 1.07110162 | ENSG000000 | chr17 | -1 | 48057103 | 3 | FALSE |
| CAACAACCTCTCCAAAGGGA | AGG | 1.05733942 | | chr15 | -1 | 62599769 | 3 | FALSE |
| CATCCTTCTCACCAAAGGGG | AAG | 1.01297257 | | chr16 | -1 | 86594199 | 3 | FALSE |
| AAAAATTCTCTCCAAAGGGG | AAG | 0.97130481 | | chr5 | 1 | 54721830 | 4 | FALSE |
| CAGCGTGTTCTCCAAAGGGG | CAG | 0.95582585 | | chr5 | 1 | 14191020 | 4 | FALSE |
| GCTCCTCCACTCCAAAGGGG | CAG | 0.94228896 | | chr17 | 1 | 49558343 | 4 | FALSE |
| AGTCATCATCTCCAAAGGGC | AGG | 0.93431604 | | chr5 | -1 | 10046165 | 4 | FALSE |
| CTTCATTCTCTCCAAAGGAG | TAG | 0.91448658 | ENSG000000 | chr11 | -1 | 75798251 | 3 | FALSE |
| AGTCATCCTGTCCAAAGGGA | AAG | 0.88271165 | | chr18 | -1 | 51768352 | 4 | FALSE |
| GATACTCCTCTCCAAAGGGC | AAG | 0.87903107 | | chr15 | 1 | 66408966 | 4 | FALSE |
| CATTCTCTGCTCCAAAGGGG | GAG | 0.87417169 | | chr10 | -1 | 90779223 | 4 | FALSE |
| GCACATCCTCTCCAAAGGGA | AGG | 0.86672463 | | chr9 | 1 | 88700535 | 4 | FALSE |
| CTTAGTCCTCTCCAAAGGGC | GAG | 0.84889286 | | chr16 | -1 | 54391594 | 4 | FALSE |
| TCTCATCTTCTACAAAGGGG | GAG | 0.84673913 | | chrX | -1 | 149295005 | 4 | FALSE |
| CATTTGCCTATCCAAAGGGG | GAG | 0.80366106 | | chr8 | 1 | 65982123 | 4 | FALSE |
| CATCGTCTCTTCCAAAGGGG | GGG | 0.79238308 | ENSG000001 | chr16 | -1 | 87956795 | 4 | FALSE |
| CACCCTCCTTCCCAAAGGGG | CAG | 0.79096733 | | chr2 | 1 | 74347104 | 4 | FALSE |
| CATTTTCCTAGCCAAAGGGG | TAG | 0.78154064 | ENSG000001 | chr11 | -1 | 47109311 | 4 | FALSE |
| CACAATCATCTACAAAGGGG | CAG | 0.77498274 | | chr13 | 1 | 60717346 | 4 | FALSE |
| CTTTATCCTTTACAAAGGGG | AAG | 0.75787923 | | chr17 | 1 | 68574649 | 4 | FALSE |
| CTTCCTCATCTCCAAAAGGG | AGG | 0.72569444 | | chr8 | 1 | 140074446 | 4 | FALSE |
| CTTCGTCCTCTCCAATGGGG | AGG | 0.62967245 | | chr2 | -1 | 136402449 | 3 | FALSE |
| AAGCATCCTATCAAAAGGGG | AGG | 0.62914291 | | chr20 | -1 | 58404478 | 4 | FALSE |
| CTGCATCCTATCTAAAGGGG | CAG | 0.61072897 | | chr5 | -1 | 173795919 | 4 | FALSE |
| TATCTTTCTCTCCAAAGGGT | GGG | 0.60756838 | | chr2 | -1 | 229408453 | 4 | FALSE |
| CACCTTCTTCTCCAAAGGAG | CAG | 0.60463217 | | chr17 | 1 | 35998658 | 4 | FALSE |
| TATCATATTCTCCAAAGGGC | AAG | 0.60037822 | | chr11 | 1 | 63668189 | 4 | FALSE |
| GATCTATCTCTCCAAAGGGG | AAG | 0.59598317 | | chr18 | -1 | 57046457 | 4 | FALSE |
| GATCACTCTTTCCAAAGGGG | AGG | 0.59204552 | | chr12 | 1 | 83596582 | 4 | FALSE |
| CACTATCCTGTCAAAAGGGG | AAG | 0.58779079 | | chr12 | 1 | 79892649 | 4 | FALSE |
| CCTCATCCTCTCAAAAAGGG | CAG | 0.5719 | | chr1 | -1 | 245981290 | 3 | FALSE |
| CATCATCTTCCCCAAAGGAG | AAG | 0.56204315 | ENSG000000 | chr17 | 1 | 4960280 | 3 | FALSE |
| GATAATGCTCTTCAAAGGGG | TAG | 0.56203204 | | chr8 | -1 | 87976851 | 4 | FALSE |
| CAACAAACTTTCCAAAGGGG | CAG | 0.55469015 | | chr3 | 1 | 184027869 | 4 | FALSE |
| CAACATCCTCTCCAAAGGAA | TGG | 0.55051556 | | chr3 | 1 | 113407849 | 3 | FALSE |
| CATGATGTTCTCCAAAGGGC | TGG | 0.5425878 | ENSG000002 | chr3 | 1 | 15475283 | 4 | FALSE |
| CAGCAGCACCTCCAAAGGGG | TGG | 0.52998343 | | chr20 | -1 | 24296403 | 4 | FALSE |
| CCTCATCACCACCAAAGGGG | AGG | 0.5275378 | | chr16 | 1 | 31128217 | 4 | FALSE |
| CATCAGATTGTCCAAAGGGG | AAG | 0.52346882 | | chr2 | 1 | 205279527 | 4 | FALSE |
| CCTCAGCATCTCCAAAGGGA | AGG | 0.51951752 | ENSG000001 | chr9 | -1 | 87707435 | 4 | FALSE |

**Off-target for R218C sgRNA**

| Sequence | PAM | Score | Gene | Chromosome | Strand | Position | Mismatches | On-target |
|---|---|---|---|---|---|---|---|---|
| GTGTCCACACTGAGTACGCA | AGG | 19.6 | ENSG00000167995 | chr11 | -1 | 61957403 | 1 | FALSE |
| ATGTCCACACTGAGTACACC | TGG | 10.425 | | chr7 | 1 | 10314123 | 2 | FALSE |
| GTGTGTAAACTGAGTACACA | AGG | 1.46807152 | | chr10 | -1 | 21984870 | 3 | FALSE |
| ATGCACACAGTGAGTACACA | GAG | 1.432778384 | | chr17 | -1 | 48247156 | 4 | FALSE |
| GAAGCCAGACTGAGTACACA | GAG | 1.422115385 | | chr20 | 1 | 48165409 | 4 | FALSE |
| GTGTTCACCCTGAGTACACT | TAG | 0.977970303 | | chr2 | 1 | 111459080 | 3 | FALSE |
| GTGTTCACCCTGAGTACACT | TAG | 0.977970303 | | chr2 | -1 | 87491157 | 3 | FALSE |
| CTTTCCACACTGAGCACACA | GGG | 0.967381888 | | chr9 | 1 | 109316353 | 3 | FALSE |
| ATTCCCACCCTGAGTACACA | TAG | 0.9132825 | | chr9 | 1 | 120468910 | 4 | FALSE |
| GAATCCATACTGAGTACACT | GAG | 0.877104566 | | chr7 | -1 | 115541230 | 4 | FALSE |
| GTGGGGAGACTGAGTACACA | CAG | 0.832166988 | | chr6 | 1 | 2149828 | 4 | FALSE |
| GTCTCCAGGGTGAGTACACA | CAG | 0.820192107 | | chr15 | -1 | 72867267 | 4 | FALSE |
| GTGATTACAATGAGTACACA | AAG | 0.803661058 | | chr6 | 1 | 131742399 | 4 | FALSE |
| GCATGCACACTCAGTACACA | CGG | 0.781996154 | | chr7 | -1 | 151819824 | 4 | FALSE |
| ATGTCCAGAGTGAGTAAACA | GAG | 0.682815709 | | chr4 | -1 | 121016770 | 4 | FALSE |
| CTGGCCACACTGAGTCCACA | GGG | 0.66020202 | | chr2 | 1 | 101414147 | 3 | FALSE |
| GGCCCCACACTGAGTACAGA | TAG | 0.611313191 | | chr7 | -1 | 134144846 | 4 | FALSE |
| GTCCTCACACTGGGTACACA | GAG | 0.609590078 | | chr22 | 1 | 43615713 | 4 | FALSE |
| CTGCCTACTCTGAGTACACA | GAG | 0.565191389 | | chr5 | 1 | 144359039 | 4 | FALSE |
| GTGTACTCTGTGAGTACACA | TGG | 0.545509974 | | chr10 | -1 | 48556885 | 4 | FALSE |
| GTGTACAGAATGAGTACAGA | TGG | 0.524636897 | | chr4 | 1 | 121633689 | 4 | FALSE |
| GAGTCCACACTGTGTACAGA | GGG | 0.51816443 | | chr4 | -1 | 7581742 | 3 | FALSE |
| TTTTCCACACTGATTACACA | CAG | 0.51406372 | ENSG00000198131 | chr19 | -1 | 58261673 | 3 | FALSE |
| GTCTCCTCACTGAGTACCCA | GGG | 0.506643053 | | chr20 | -1 | 59381089 | 3 | FALSE |
| CAGTCCAGACTGAGAACACA | AAG | 0.50515873 | | chr2 | 1 | 30924911 | 4 | FALSE |
| TGGTTCACACTGAGGACACA | CAG | 0.489615385 | | chr2 | 1 | 1373795 | 4 | FALSE |
| ATGAACACACTGAGCACACA | TGG | 0.479773869 | ENSG00000231508 | chr10 | 1 | 97393296 | 4 | FALSE |
| GTCACCCCACTGAGTAAACA | GAG | 0.473672978 | | chr10 | 1 | 49351882 | 4 | FALSE |
| GTGTTCACACTGTGTACAAA | CAG | 0.446280347 | | chr3 | -1 | 168181081 | 3 | FALSE |
| GTGTACAGACTGACTACACA | CAG | 0.443035994 | | chr20 | -1 | 15444572 | 3 | FALSE |
| GTTTGCACACAAAGTACACA | CAG | 0.437971408 | | chr10 | 1 | 63488170 | 4 | FALSE |
| GTGTACACAATGATTACACA | AGG | 0.40803615 | | chr8 | 1 | 113125227 | 3 | FALSE |
| CTGTCCAGGCTGTGTACACA | CAG | 0.399231309 | | chr16 | -1 | 80654292 | 4 | FALSE |
| CTCTCCACACTGAGTAGACC | TGG | 0.394359707 | | chr1 | -1 | 175504570 | 4 | FALSE |
| GTTTCAACAGTGAGTAGACA | GGG | 0.390436322 | | chr5 | 1 | 179960628 | 4 | FALSE |
| TTCTCCACACTAAGTAAACA | GGG | 0.38910025 | | chr2 | 1 | 34814827 | 4 | FALSE |
| GTTTCCACAAGCAGTACACA | GAG | 0.385757183 | | chr7 | -1 | 21421854 | 4 | FALSE |
| GTCTCCATCCTGAGTACAGA | GAG | 0.361528006 | ENSG00000151348 | chr11 | 1 | 44124841 | 4 | FALSE |
| GTGTACAGGCTGTGTACACA | AAG | 0.35845875 | | chr11 | -1 | 103603253 | 4 | FALSE |
| GTGTGTTCGCTGAGTACACA | GAG | 0.347274102 | | chr16 | 1 | 10566772 | 4 | FALSE |
| GTATCTTCACTGAGTACACT | AGG | 0.33439929 | | chr5 | -1 | 39906309 | 4 | FALSE |
| TTGTCCACAGAGCGTACACA | GGG | 0.333991947 | ENSG00000197989 | chr1 | -1 | 28582390 | 4 | FALSE |
| GTGACCACACTGGGCACACA | TGG | 0.333435533 | | chr2 | 1 | 131536773 | 3 | FALSE |
| GTGACCACACTGGGCACACA | TGG | 0.333435533 | | chr2 | 1 | 131202725 | 3 | FALSE |
| GTGACCACACTGGGCACACA | TGG | 0.333435533 | | chr2 | -1 | 130136194 | 3 | FALSE |
| GTGTATGCACAGAGTACACA | CAG | 0.330770661 | | chr10 | -1 | 105461678 | 4 | FALSE |
| GTGTACACTGTGTGTACACA | GGG | 0.330140509 | | chr2 | 1 | 207287915 | 4 | FALSE |
| GTATACACACTCGGTACACA | CAG | 0.316975285 | | chr19 | -1 | 46319193 | 4 | FALSE |
| CTCTACACACTGAGTCCACA | CAG | 0.316320157 | | chr14 | 1 | 94544044 | 4 | FALSE |
| CTGTCAGCACTGAGTATACA | GGG | 0.303074972 | | chr13 | -1 | 23985245 | 4 | FALSE |

**Off-target for AAVS1 sgRNA**

| Sequence | PAM | Score | Gene | Chromosome | Strand | Position | Mismatches | On-target |
|---|---|---|---|---|---|---|---|---|
| GGGGCCACTAGGGACAGGAT | TGG | 100 | | chr19 | -1 | 55115755 | 0 | TRUE |
| GGAGACATTAGGGACAGGAT | AAG | 2.548843537 | | chr10 | 1 | 119439186 | 3 | FALSE |
| GAGGGCTCTAGGGACAGGAT | GAG | 1.765578231 | | chr9 | -1 | 90086414 | 3 | FALSE |
| GGCCCCACTAGGGACAGGAC | GAG | 1.747668456 | | chr7 | 1 | 122844142 | 3 | FALSE |
| GGGACCATCAGGGACAGGAT | GGG | 1.579455782 | | chr6 | 1 | 36797704 | 3 | FALSE |
| CAGGGCACTGGGGACAGGAT | CAG | 1.45825 | | chr14 | 1 | 92970608 | 4 | FALSE |
| GGGGCCAGTGGGGACAGGAG | CAG | 1.286963492 | | chr11 | 1 | 2127694 | 3 | FALSE |
| GGGGCCAGTGGGGACAGGAA | GGG | 1.286963492 | | chr2 | -1 | 231959836 | 3 | FALSE |
| GGGGCCAATTAGGACAGGAT | GGG | 1.240350575 | | chr13 | 1 | 105960580 | 3 | FALSE |
| GGGGTCACTGGGGACAAGAT | TGG | 1.188203704 | | chr15 | -1 | 45535701 | 3 | FALSE |
| TGGGCCACTATGGACAGGAA | TGG | 1.102071429 | | chr12 | -1 | 108187905 | 3 | FALSE |
| GGGACCACTGGGCACAGGAT | CGG | 1.059798592 | | chr15 | -1 | 25222754 | 3 | FALSE |
| GGGGATGCTAGGGACAGGAT | GAG | 0.972872986 | | chr2 | 1 | 204531421 | 3 | FALSE |
| ACTGCCTCTAGGGACAGGAT | AGG | 0.971304808 | | chr15 | -1 | 36995662 | 4 | FALSE |
| AGTGCCAACAGGGACAGGAT | GGG | 0.945468667 | | chr4 | 1 | 22172000 | 4 | FALSE |
| TCGCCCACCAGGGACAGGAT | CAG | 0.934200644 | ENSG00000140983 | chr16 | -1 | 668379 | 4 | FALSE |
| GTGTCCAAGAGGGACAGGAT | GGG | 0.92625 | | chr1 | -1 | 154829128 | 4 | FALSE |
| GCAGCCAGGAGGGACAGGAT | GGG | 0.921121835 | | chr12 | 1 | 49891231 | 4 | FALSE |
| GAGGGCAGCAGGGACAGGAT | GGG | 0.918433544 | | chr12 | 1 | 131827329 | 4 | FALSE |
| GGCCCCAAGAGGGACAGGAT | GAG | 0.890545176 | | chr8 | 1 | 141604148 | 4 | FALSE |
| GGTTCCAGCAGGGACAGGAT | CAG | 0.890545176 | ENSG00000113303 | chr5 | 1 | 180947683 | 4 | FALSE |
| GGAGCCAGTAGGGAGAGGAT | AGG | 0.885487125 | | chr16 | 1 | 33884580 | 3 | FALSE |
| GGAGCCAGTAGGGAGAGGAT | AGG | 0.885487125 | | chr16 | -1 | 32969375 | 3 | FALSE |
| GTGGCCAGCTGGGACAGGAT | AGG | 0.85307625 | | chr9 | 1 | 34350240 | 4 | FALSE |
| GGAGGGAGTAGGGACAGGAT | GAG | 0.839975543 | | chr11 | 1 | 113401727 | 4 | FALSE |
| GGGGGAACTAGTGACAGGAT | AGG | 0.823222707 | | chr20 | -1 | 43709929 | 3 | FALSE |
| GAACCTACTAGGGACAGGAT | GAG | 0.820516651 | | chr6 | -1 | 157966901 | 4 | FALSE |
| GAGGCCACCAGGGACAGGCT | GGG | 0.818083893 | | chr5 | -1 | 170084415 | 3 | FALSE |
| GGGCACAGTAGGGACAGGAA | GAG | 0.811782787 | | chr8 | -1 | 17472023 | 4 | FALSE |
| GAGGCCAGTGGGGACAGGAC | AGG | 0.790868822 | | chrX | -1 | 150758233 | 4 | FALSE |
| GGGGATATTGGGGACAGGAT | TGG | 0.79085371 | | chr4 | -1 | 47212824 | 4 | FALSE |
| GGAGGCACTGGTGACAGGAT | GAG | 0.720218458 | ENSG00000223995 | chrX | -1 | 128838711 | 4 | FALSE |
| GGGGACAGTGGGGACAGGAG | GGG | 0.716336682 | | chr8 | -1 | 608459 | 4 | FALSE |
| GGTGCCACTAGGCACAGGAG | CGG | 0.676347693 | | chr8 | 1 | 143802967 | 3 | FALSE |
| GCGGCCAATGGGGACATGAT | GGG | 0.661366001 | ENSG00000063978 | chr4 | 1 | 2469079 | 4 | FALSE |
| GAGGACAGTAGGGACAGGTT | AAG | 0.634004237 | | chr18 | 1 | 8749311 | 4 | FALSE |
| CAGGCCCCTAGGGACAGGAG | CAG | 0.630210675 | | chr10 | 1 | 48519097 | 4 | FALSE |
| GCAGCCCCAAGGGACAGGAT | GGG | 0.623771948 | ENSG00000166126 | chr14 | -1 | 102932045 | 4 | FALSE |
| AGGGGCACTGGGGACAGGCT | TGG | 0.618883046 | ENSG00000126461 | chr19 | -1 | 49658572 | 4 | FALSE |
| GCTGCCACTGGGTACAGGAT | CAG | 0.610728971 | | chr9 | -1 | 89227532 | 4 | FALSE |
| AGGGCCCTTATGGACAGGAT | GGG | 0.605569086 | ENSG00000174951 | chr19 | -1 | 48749680 | 4 | FALSE |
| TGGGCCAGTGGGGACAGGGT | GGG | 0.597418895 | | chr2 | -1 | 121052619 | 4 | FALSE |
| GGGGCTTCTAAGGACAGGAT | GGG | 0.592837031 | | chr19 | -1 | 16064183 | 3 | FALSE |
| GAGGGCCCTAGGGACAGGAG | AGG | 0.58649664 | | chr12 | -1 | 125092753 | 4 | FALSE |
| GGTGGCTCTAGGGACAGGAA | GAG | 0.558901809 | | chr3 | 1 | 46366887 | 4 | FALSE |
| GAGGCCGCTGGGGACAGGAC | GGG | 0.546481106 | | chr16 | -1 | 715574 | 4 | FALSE |
| GGGCCCTATAGGGACAGGAA | AGG | 0.542587801 | | chr3 | -1 | 48569139 | 4 | FALSE |
| GGTGCCACCAGGGAGAGGAT | GGG | 0.541032634 | | chr22 | -1 | 44303224 | 3 | FALSE |
| GGAGCCACCAGGGAAAGGAT | GAG | 0.541032634 | | chr13 | -1 | 99628492 | 3 | FALSE |
| ATGGCCACTAAGGACAGGAA | AGG | 0.538880515 | | chr12 | 1 | 107092504 | 4 | FALSE |
