## Supplementary material for "Human iPSC modeling reveals mutation-specific responses to gene therapy in Best disease": SI Data File D

**Table S4: Analysis of additional adBD mutations for amenability to allele specific editing or scarless base editing.** For each adBD mutation tested in this study, and additional representative adBD mutations, a search was performed for unique sgRNAs that overlap with the adBD mutation locus. An additional 23 reported adBD mutations were selected as representative examples using the ClinVar database (https://www.ncbi.nlm.nih.gov/clinvar?term=607854[MIM]) or the Retina International database and were limited to mutations in amino acids not known to have direct involvement in ion binding. Representative mutations were selected from the 5' end of the BEST1 gene as mutations towards the 5’ end are more likely to introduce indels and early premature termination codons that trigger nonsense mediated decay (Popp and Maquat, 2016). The sgRNA search was performed using an online CRISPR tool (benchling.com) and was limited to 20 nucleotide sgRNAs with an "NGG" PAM site. The human reference genome (GRCh38) was used to evaluate uniqueness. Specificity Scores (Doench, Fusi et al., 2016) and Efficiency Score (Hsu et al., 2013) are reported for each sgRNA. Additionally, a score for CRISPR activity at the WT BEST1 locus (WT BEST1 Score) is also listed if it was reported in the Specificity Score output. A composite allele specific editing (ASE) score was calculated using the following formula: ASE score = (Spec. Score/100)*(Efficiency Score/100)*((100-(WT BEST1 Score))/100). In this study, sgRNAs with an ASE score ≥ 0.34 were found to enable mutant allele specific editing while sgRNAs with an ASE score ≤ 0.26 did not enable allele specific editing. Candidate sgRNAs with an ASE score ≥ 0.34 are highlighted in green, while candidate sgRNas with an ASE score of < 0.34 and > 0.26 are highlighted in yellow. A search was also performed for adBD mutations amenable to scarless base editing (Komor et al., 2016, Fig 2b). Again, a search for candidate sgRNAs was performed using an online CRISPR tool (benchling.com) and was limited to 20 nucleotide sgRNAs with an "NGG" PAM site. A sgRNA was considered amendable to scarless base editing if the predicted editing outcome corrected the missense mutation to WT without introducing any other amino acid substitutions.

1=True, 0=False

| BEST1 Mutation | Location on Chr 11 | WT | MT | sgRNA candidates | Position | Strand | Sequence | PAM | Specificity Score | Efficiency Score | Comp eff at WT | ASE score | Position | Guide | Guide Strand | Edited 5'->3' Sequence | Edit Scores | Efficiency Score | Amenable to scarless repair? |
|---|---|---|---|---|---|---|---|---|---|---|---|---|---|---|---|---|---|---|---|
| A146K | 61955906 | CATCCTGCGCAGCGTCAGCACCGCAGTCTACAAGCGCTTCCCCAGC | CATCCTGCGCAGCGTCAGCACCaaAGTCTACAAGCGCTTCCCCAGC | 2 | 61955893 | -1 | CTttGGTGCTGACGCTGCGC | AGG | 84.7 | 51.6 | 5.1 | 0.41 | 61955893 | CTttGGTGCTGACGCTGCGC | -1 | GCGCAGCGTCAGCACCaaAA | 0.8 | 84.6932259 | 0 |
| | | | | | 61955910 | -1 | GGGGAAGCGCTTGTAGACTt | tGG | 84.8 | 53.3 | 41.7 | 0.26 | 61955910 | GGGGAAGCGCTTGTAGACTt | -1 | aAGTCTACAAGCACTTCCCC | 2.8 | 84.7773398 | 0 |
| BEST1 Mutation | Location | WT | MT | sgRNA candidates | Position | Strand | Sequence | PAM | Specificity Score | Efficiency Score | Comp eff at WT | ASE score | Position | Guide | Guide Strand | Edited 5'->3' Sequence | Edit Scores | Efficiency Score | scarless repair |
| R218C | 61957402 | CCCCCAGGAGATGAACACCTTGCGTACTCAGTGTGGACACCTGTA | CCCCCAGGAGATGAACACCTTGtGTACTCAGTGTGGACACCTGTA | 6 | 61957386 | -1 | aCAAGGTGTTCATCTCCTGG | GGG | 58.6 | 68.5 | 100 | 0.00 | 61957386 | aCAAGGTGTTCATCTCCTGG | -1 | CCAGGAGATGAACACCTTAt | 2.5 | 58.5713416 | 0 |
| | | | | | 61957387 | -1 | CaCAAGGTGTTCATCTCCTG | GGG | 61.7 | 59.3 | 100 | 0.00 | 61957387 | CaCAAGGTGTTCATCTCCTG | -1 | CAGGAGATGAACACCTTAtA | 5.6, 0.8 | 61.6679562 | 0 |
| | | | | | 61957388 | -1 | ACaCAAGGTGTTCATCTCCT | GGG | 62.7 | 58.1 | 98.6 | 0.01 | 61957388 | ACaCAAGGTGTTCATCTCCT | -1 | AGGAGATGAACACCTTAtAT | 11.6, 2.5 | 62.6638307 | 0 |
| | | | | | 61957389 | -1 | TACaCAAGGTGTTCATCTCC | TGG | 72.0 | 44.6 | 100 | 0.00 | 61957389 | TACaCAAGGTGTTCATCTCC | -1 | GGAGATGAACACCTTAtATA | 21.9, 5.6 | 71.955753 | 0 |
| | | | | | 61957403 | -1 | GTGTCCACACTGAGTACaCA | AGG | 63.3 | 67.2 | 19.6 | 0.34 | 61957410 | AACACCTTGtGTACTCAGTG | 1 | AATATTTTGtGTACTCAGTG | 5.6, 21.9, 21.4 | 73.6558533 | 0 |
| | | | | | 61957410 | 1 | AACACCTTGtGTACTCAGTG | TGG | 73.7 | 72.1 | 92.1 | 0.04 | 61957403 | GTGTCCACACTGAGTACaCA | -1 | TGtGTACTCAGTATAAACAC | 6.8, 21.4, 24.2 | 63.3051198 | 0 |
| BEST1 Mutation | Location | WT | MT | sgRNA candidates | Position | Strand | Sequence | PAM | Specificity Score | Efficiency Score | Comp eff at WT | ASE score | Position | Guide | Guide Strand | Edited 5'->3' Sequence | Edit Scores | Efficiency Score | scarless repair |
| N296H | 61959516 | CAAGGTGGCAGAGCAGCTCATCAACCCCTTTGGAGAGGATGATGA | CAAGGTGGCAGAGCAGCTCATCcACCCCTTTGGAGAGGATGATGA | 3 | 61959521 | 1 | GAGCAGCTCATCcACCCCTT | TGG | 54.8 | 54.3 | 38.7 | 0.18 | 61959521 | GAGCAGCTCATCcACCCCTT | 1 | GAGTAGTTCATCcACCCCTT | 0.8, 17.0 | 54.8376243 | 0 |
| | | | | | 61959521 | -1 | CATCATCCTCTCCAAAGGGG | TgG | 54.0 | 64.6 | 0 | 0.35 | 61959526 | GCTCATCcACCCCTTTGGAG | 1 | GTTTATTtACCCCTTTGGAG | 0.4, 21.7, 25.2, 8.7 | 71.1233097 | 0 |
| | | | | | 61959526 | 1 | GCTCATCcACCCCTTTGGAG | AGG | 71.1 | 65.1 | 100 | 0.00 | 61959521 | CATCATCCTCTCCAAAGGGG | -1 | CCCCTTTGGAGAAAATAATA | 8.7, 25.2, 21.7, 4.1 | 53.9815098 | 0 |

1=True, 0=False

| BEST1 Mutation | Location | WT | MT | sgRNA candidates | Position | Strand | Sequence | PAM | Specificity Score | Efficiency Score | Comp eff at WT | ASE score | Position | Guide | Guide Strand | Edited 5'->3' Sequence | Edit Scores | Efficiency Score | scarless repair |
|---|---|---|---|---|---|---|---|---|---|---|---|---|---|---|---|---|---|---|---|
| T6R | 61951823 | CTGGCCATGACCATCACTTACACAAGCCAAGTGGCTAATGCCCGC | CTGGCCATGACCATCACTTACAgAAGCCAAGTGGCTAATGCCCGC | 3 | 61951811 | -1 | GCTTcTGTAAGTGATGGTCA | TGG | 64.0 | 60.8 | 100 | 0.00 | 61951811 | GCTTcTGTAAGTGATGGTCA | -1 | TGACCATCACTTACAaAAAC | 24.2, 0.4 | 64.0067307 | 0 |
| | | | | | 61951817 | -1 | CACTTGGCTTcTGTAAGTGA | TGG | 62.8 | 58.8 | 55.5 | 0.16 | 61951817 | CACTTGGCTTcTGTAAGTGA | -1 | TCACTTACAgAAACCAAATA | 2.8, 5.6, 0.5 | 62.7517108 | 0 |
| | | | | | 61951829 | 1 | CATCACTTACAgAAGCCAAG | TGG | 54.7 | 66.0 | 49.2 | 0.18 | 61951829 | CATCACTTACAgAAGCCAAG | 1 | TATTATTTACAgAAGCCAAG | 0.5, 21.7, 13.5 | 54.7476308 | 0 |
| BEST1 Mutation | Location | WT | MT | sgRNA candidates | Position | Strand | Sequence | PAM | Specificity Score | Efficiency Score | Comp eff at WT | ASE score | Position | Guide | Guide Strand | Edited 5'->3' Sequence | Edit Scores | Efficiency Score | scarless repair |
| T6Lys | 61951823 | CTGGCCATGACCATCACTTACACAAGCCAAGTGGCTAATGCCCGC | CTGGCCATGACCATCACTTACAaAAGCCAAGTGGCTAATGCCCGC | 3 | 61951811 | -1 | GCTTtTGTAAGTGATGGTCA | TGG | 64.7 | 57.0 | 100 | 0.00 | 61951811 | GCTTtTGTAAGTGATGGTCA | -1 | TGACCATCACTTACAaAAAC | 0.4 | 64.6836068 | 0 |
| | | | | | 61951817 | -1 | CACTTGGCTTtTGTAAGTGA | TGG | 60.3 | 51.6 | 55.5 | 0.14 | 61951817 | CACTTGGCTTtTGTAAGTGA | -1 | TCACTTACAaAAACCAAATA | 2.8, 5.6, 0.5 | 60.2761609 | 0 |
| | | | | | 61951829 | 1 | CATCACTTACAaAAGCCAAG | TGG | 54.0 | 66.3 | 49.2 | 0.18 | 61951829 | CATCACTTACAaAAGCCAAG | 1 | TATTATTTACAaAAGCCAAG | 0.5, 21.7, 13.5 | 53.9877777 | 0 |
| BEST1 Mutation | Location | WT | MT | sgRNA candidates | Position | Strand | Sequence | PAM | Specificity Score | Efficiency Score | Comp eff at WT | ASE score | Position | Guide | Guide Strand | Edited 5'->3' Sequence | Edit Scores | Efficiency Score | scarless repair |
| S16F | 61951853 | GTGGCTAATGCCCGCTTAGGCTCCTTCTCCCGCCTGCTGCTGTGC | GTGGCTAATGCCCGCTTAGGCTtCTTCTCCCGCCTGCTGCTGTGC | 2 | 61951847 | -1 | GCGGGAGAAGaAGCCTAAGC | GGG | 68.8 | 63.1 | 55.5 | 0.19 | 61951847 | GCGGGAGAAGaAGCCTAAGC | -1 | GCTTAGGCTtCTTCTCCCAC | 0.4 | 68.8070338 | 0 |
| | | | | | 61951848 | -1 | GGCGGGAGAAGaAGCCTAAG | CGG | 58.7 | 63.0 | 49.2 | 0.19 | 61951848 | GGCGGGAGAAGaAGCCTAAG | -1 | CTTAGGCTtCTTCTCCCACC | 0.5 | 58.670214 | 0 |
| BEST1 Mutation | Location | WT | MT | sgRNA candidates | Position | Strand | Sequence | PAM | Specificity Score | Efficiency Score | Comp eff at WT | ASE score | Position | Guide | Guide Strand | Edited 5'->3' Sequence | Edit Scores | Efficiency Score | scarless repair |
| F17Ser | 61951856 | GCTAATGCCCGCTTAGGCTCCTTCTCCCGCCTGCTGCTGTGCTGG | GCTAATGCCCGCTTAGGCTCCTcCTCCCGCCTGCTGCTGTGCTGG | 6 | 61951847 | -1 | GCGGGAGgAGGAGCCTAAGC | GGG | 63.2 | 54.9 | 100 | 0.00 | 61951847 | GCGGGAGgAGGAGCCTAAGC | -1 | GCTTAGGCTCCTcCTCCCAC | 0.4 | 63.1755395 | 0 |
| | | | | | 61951848 | -1 | GGCGGGAGgAGGAGCCTAAG | CGG | 55.7 | 54.9 | 61.1 | 0.12 | 61951848 | GGCGGGAGgAGGAGCCTAAG | -1 | CTTAGGCTCCTcCTCCCACC | 0.5 | 55.7273867 | 0 |
| | | | | | 61951859 | -1 | GCACAGCAGCAGGCGGGAGg | AGG | 43.9 | 48.8 | 41.7 | 0.12 | 61951859 | GCACAGCAGCAGGCGGGAGg | -1 | cCTCCCGCCTGCTACTATAC | 17.0, 11.6, 0.4 | 43.902817 | 0 |
| | | | | | 61951862 | -1 | CCAGCACAGCAGCAGGCGGG | AGg | 50.5 | 60.2 | 0 | 0.30 | 61951873 | cCTCCCGCCTGCTGCTGTGC | 1 | tTTTTTGTCTGCTGCTGTGC | 4.1, 11.0, 21.7, 20.3, 21.4, 2.8 | 50.1175395 | 0 |
| | | | | | 61951873 | 1 | cCTCCCGCCTGCTGCTGTGC | TGG | 50.1 | 46.9 | 100 | 0.00 | 61951862 | CCAGCACAGCAGCAGGCGGG | -1 | CCCGCCTGCTGCTATACTAA | 19.0, 4.6, 11.0, 0.2 | 50.5497598 | 0 |
| BEST1 Mutation | Location | WT | MT | sgRNA candidates | Position | Strand | Sequence | PAM | Specificity Score | Efficiency Score | Comp eff at WT | ASE score | Position | Guide | Guide Strand | Edited 5'->3' Sequence | Edit Scores | Efficiency Score | scarless repair |
| F17Cys | 61951856 | GCTAATGCCCGCTTAGGCTCCTTCTCCCGCCTGCTGCTGTGCTGG | GCTAATGCCCGCTTAGGCTCCTgCTCCCGCCTGCTGCTGTGCTGG | 4 | 61951847 | -1 | GCGGGAGcAGGAGCCTAAGC | GGG | 62.5 | 56.5 | 100 | 0.00 | 61951847 | GCGGGAGcAGGAGCCTAAGC | -1 | GCTTAGGCTCCTaCTCCCAC | 2.8, 0.4 | 62.4850404 | 0 |
| | | | | | 61951848 | -1 | GGCGGGAGcAGGAGCCTAAG | CGG | 62.1 | 53.9 | 61.1 | 0.13 | 61951848 | GGCGGGAGcAGGAGCCTAAG | -1 | CTTAGGCTCCTgCTCCCACC | 0.5 | 62.1269373 | 0 |
| | | | | | 61951859 | -1 | GCACAGCAGCAGGCGGGAGc | AGG | 45.6 | 45.3 | 41.7 | 0.12 | 61951859 | GCACAGCAGCAGGCGGGAGc | -1 | gCTCCCGCCTGCTACTATAC | 17.0, 11.6, 0.4 | 45.5637514 | 0 |
| | | | | | 61951873 | 1 | gCTCCCGCCTGCTGCTGTGC | TGG | 50.7 | 48.8 | 100 | 0.00 | 61951873 | gCTCCCGCCTGCTGCTGTGC | 1 | gTTTTTGTCTGCTGCTGTGC | 0.4, 21.7, 20.3, 21.4, 2.8 | 50.6667529 | 0 |
| BEST1 Mutation | Location | WT | MT | sgRNA candidates | Position | Strand | Sequence | PAM | Specificity Score | Efficiency Score | Comp eff at WT | ASE score | Position | Guide | Guide Strand | Edited 5'->3' Sequence | Edit Scores | Efficiency Score | scarless repair |
| R19Leu | 61951862 | GCCCGCTTAGGCTCCTTCTCCCGCCTGCTGCTGTGCTGGCGGGGC | GCCCGCTTAGGCTCCTTCTCCCtCCTGCTGCTGTGCTGGCGGGGC | 9 | 61951847 | -1 | GaGGGAGAAGGAGCCTAAGC | GGG | 58.0 | 55.6 | 100 | 0.00 | 61951859 | GCACAGCAGCAGGaGGGAGA | -1 | TCTCCCtCCTGCTACTATAC | 17.0, 11.6, 0.4 | 30.2381882 | 0 |
| | | | | | 61951848 | -1 | GGaGGGAGAAGGAGCCTAAG | CGG | 50.7 | 60.2 | 98.6 | 0.00 | 61951873 | TCTCCCtCCTGCTGCTGTGC | 1 | TTTTTTtTCTGCTGCTGTGC | 6.4, 21.7, 20.3, 21.4, 25.4 | 39.594357 | 0 |
| | | | | | 61951859 | -1 | GCACAGCAGCAGGaGGGAGA | AGG | 30.2 | 55.9 | 14.9 | 0.14 | 61951876 | CCCtCCTGCTGCTGTGCTGG | 1 | TTTtTTTGCTGCTGTGCTGG | 4.1, 11.0, 5.7, 24.2, 21.4 | 35.7999893 | 0 |
| | | | | | 61951865 | -1 | CCGCCAGCACAGCAGCAGGa | GGG | 33.5 | 61.6 | 41.7 | 0.12 | 61951877 | CCtCCTGCTGCTGTGCTGGC | 1 | TTtTTTGTTGCTGTGCTGGC | 0.5, 11.0, 21.7, 20.3, 2.8 | 40.4423267 | 0 |
| | | | | | 61951866 | -1 | CCCGCCAGCACAGCAGCAGG | aGG | 33.1 | 53.8 | 100 | 0.00 | 61951878 | CtCCTGCTGCTGTGCTGGCG | 1 | TtTTTGTTGCTGTGCTGGCG | 0.5, 16.0, 9.3, 17.0 | 39.8455569 | 0 |
| | | | | | 61951873 | 1 | TCTCCCtCCTGCTGCTGTGC | TGG | 39.6 | 47.1 | 68.3 | 0.06 | 61951865 | CCGCCAGCACAGCAGCAGGa | -1 | tCCTGCTGCTGTACTAACAA | 2.8, 20.3, 0.8, 11.0, 0.5 | 33.5048104 | 0 |
| | | | | | 61951876 | 1 | CCCtCCTGCTGCTGTGCTGG | CGG | 35.8 | 44.4 | 100 | 0.00 | 61951866 | CCCGCCAGCACAGCAGCAGG | -1 | CCTGCTGCTGTGCTAACAAA | 21.4, 4.6, 5.7, 11.0, 0.5 | 33.1159357 | 0 |
| | | | | | 61951877 | 1 | CCtCCTGCTGCTGTGCTGGC | GGG | 40.4 | 38.0 | 98.6 | 0.00 | | | | | | | 0 |
| | | | | | 61951878 | 1 | CtCCTGCTGCTGTGCTGGCG | GGG | 39.8 | 49.0 | 100 | 0.00 | | | | | | | 0 |
| BEST1 Mutation | Location | WT | MT | sgRNA candidates | Position | Strand | Sequence | PAM | Specificity Score | Efficiency Score | Comp eff at WT | ASE score | Position | Guide | Guide Strand | Edited 5'->3' Sequence | Edit Scores | Efficiency Score | scarless repair |
| L20Val | 61951864 | CCGCTTAGGCTCCTTCTCCCGCCTGCTGCTGTGCTGGCGGGGCAG | CCGCTTAGGCTCCTTCTCCCGCgTGCTGCTGTGCTGGCGGGGCAG | 8 | 61951848 | -1 | cGCGGGAGAAGGAGCCTAAG | CGG | 67.9 | 62.0 | 100 | 0.00 | 61951848 | cGCGGGAGAAGGAGCCTAAG | -1 | CTTAGGCTCCTTCTCCCACa | 0.5, 0.8 | 67.8539998 | 0 |
| | | | | | 61951859 | -1 | GCACAGCAGCAcGCGGGAGA | AGG | 71.7 | 52.0 | 49.2 | 0.19 | 61951859 | GCACAGCAGCAcGCGGGAGA | -1 | TCTCCCGCgTGCTACTATAC | 17.0, 11.6, 0.4 | 71.6655249 | 0 |
| | | | | | 61951865 | -1 | CCGCCAGCACAGCAGCAcGC | GGG | 66.4 | 63.9 | 19.6 | 0.34 | 61951873 | TCTCCCGCgTGCTGCTGTGC | 1 | TTTTTTGTgTGCTGCTGTGC | 6.4, 21.7, 20.3, 21.4, 2.8 | 68.8838656 | 0 |
| | | | | | 61951866 | -1 | CCCGCCAGCACAGCAGCAcG | CGG | 54.4 | 56.7 | 31.5 | 0.21 | 61951876 | CCCGCgTGCTGCTGTGCTGG | 1 | TTTGTgTGCTGCTGTGCTGG | 4.1, 11.0, 5.7, 4.6 | 51.199476 | 0 |
| | | | | | 61951873 | 1 | TCTCCCGCgTGCTGCTGTGC | TGG | 68.9 | 49.9 | 61.1 | 0.13 | 61951877 | CCGCgTGCTGCTGTGCTGGC | 1 | TTGTgTGTTGCTGTGCTGGC | 0.5, 11.0, 0.8, 2.8 | 56.1718796 | 0 |
| | | | | | 61951876 | 1 | CCCGCgTGCTGCTGTGCTGG | CGG | 51.2 | 46.6 | 60.5 | 0.09 | 61951878 | CGCgTGCTGCTGTGCTGGCG | 1 | TGTgTGTTGCTGTGCTGGCG | 0.5, 0.5, 17.0 | 58.7480413 | 0 |
| | | | | | 61951877 | 1 | CCGCgTGCTGCTGTGCTGGC | GGG | 56.2 | 39.3 | 100 | 0.00 | 61951865 | CCGCCAGCACAGCAGCAcGC | -1 | GCgTGCTGCTGTACTAACAA | 2.8, 20.3, 0.8, 11.0, 0.5 | 66.4396565 | 0 |
| | | | | | 61951878 | 1 | CGCgTGCTGCTGTGCTGGCG | GGG | 58.7 | 52.3 | 100 | 0.00 | 61951866 | CCCGCCAGCACAGCAGCAcG | -1 | CgTGCTGCTGTGCTAACAAA | 21.4, 4.6, 5.7, 11.0, 0.5 | 54.4099256 | 0 |
| BEST1 Mutation | Location | WT | MT | sgRNA candidates | Position | Strand | Sequence | PAM | Specificity Score | Efficiency Score | Comp eff at WT | ASE score | Position | Guide | Guide Strand | Edited 5'->3' Sequence | Edit Scores | Efficiency Score | scarless repair |
| L21V | 61951867 | CTTAGGCTCCTTCTCCCGCCTGCTGCTGTGCTGGCGGGGCAGCAT | CTTAGGCTCCTTCTCCCGCCTGgTGCTGTGCTGGCGGGGCAGCAT | 9 | 61951859 | -1 | GCACAGCAcCAGGCGGGAGA | AGG | 56.6 | 51.5 | 61.1 | 0.11 | 61951862 | CTTAGGCTCCTTCTCCCGCC | 1 | TTTAGGTTCCTTCTCCCGCC | 0.2, 17.0 | 66.7581703 | 0 |
| | | | | | 61951862 | 1 | CTTAGGCTCCTTCTCCCGCC | TGg | 66.8 | 49.5 | 0 | 0.33 | 61951859 | GCACAGCAcCAGGCGGGAGA | -1 | TCTCCCGCCTGgTACTATAC | 17.0, 11.6, 0.4 | 56.5597944 | 0 |
| | | | | | 61951865 | -1 | CCGCCAGCACAGCAcCAGGC | GGG | 45.4 | 55.3 | 26.8 | 0.18 | 61951873 | TCTCCCGCCTGgTGCTGTGC | 1 | TTTTTTGTCTGgTGCTGTGC | 6.4, 21.7, 20.3, 21.4, 2.8 | 60.149867 | 0 |
| | | | | | 61951866 | -1 | CCCGCCAGCACAGCAcCAGG | CGG | 42.8 | 63.4 | 17.2 | 0.22 | 61951876 | CCCGCCTGgTGCTGTGCTGG | 1 | TTTGTTTGgTGCTGTGCTGG | 4.1, 11.0, 5.7, 4.6, 21.4 | 30.6659013 | 0 |
| | | | | | 61951869 | -1 | TGCCCCGCCAGCACAGCAcC | AGG | 58.0 | 56.4 | 31.5 | 0.22 | 61951877 | CCGCCTGgTGCTGTGCTGGC | 1 | TTGTTTGgTGCTGTGCTGGC | 0.5, 11.0, 0.8, 20.3 | 53.1706576 | 0 |
| | | | | | 61951873 | 1 | TCTCCCGCCTGgTGCTGTGC | TGG | 60.1 | 49.0 | 49.2 | 0.15 | 61951878 | CGCCTGgTGCTGTGCTGGCG | 1 | TGTTTGgTGCTGTGCTGGCG | 0.5, 0.5, 9.3 | 56.8500979 | 0 |
| | | | | | 61951876 | 1 | CCCGCCTGgTGCTGTGCTGG | CGG | 30.7 | 46.2 | 61.1 | 0.06 | 61951865 | CCGCCAGCACAGCAcCAGGC | -1 | GCCTGgTGCTGTACTAACAA | 2.8, 20.3, 0.8, 11.0, 0.5 | 45.3887965 | 0 |
| | | | | | 61951877 | 1 | CCGCCTGgTGCTGTGCTGGC | GGG | 53.2 | 38.8 | 100 | 0.00 | 61951866 | CCCGCCAGCACAGCAcCAGG | -1 | CCTGgTGCTGTGCTAACAAA | 21.4, 4.6, 5.7, 11.0, 0.5 | 42.7804395 | 0 |
| | | | | | 61951878 | 1 | CGCCTGgTGCTGTGCTGGCG | GGG | 56.9 | 50.6 | 68.3 | 0.09 | 61951869 | TGCCCCGCCAGCACAGCAcC | -1 | GgTGCTGTGCTGACAAAACA | 2.8, 21.4, 20.3, 9.3, 0.5 | 58.0362665 | 0 |
| BEST1 Mutation | Location | WT | MT | sgRNA candidates | Position | Strand | Sequence | PAM | Specificity Score | Efficiency Score | Comp eff at WT | ASE score | Position | Guide | Guide Strand | Edited 5'->3' Sequence | Edit Scores | Efficiency Score | scarless repair |
| W24Cys | 61951878 | TCTCCCGCCTGCTGCTGTGCTGGCGGGGCAGCATCTACAAGCTGC | TCTCCCGCCTGCTGCTGTGCTGtCGGGGCAGCATCTACAAGCTGC | 6 | 61951865 | -1 | CCGaCAGCACAGCAGCAGGC | GGG | 39.8 | 56.8 | 100 | 0.00 | 61951876 | CCCGCCTGCTGCTGTGCTGt | 1 | TTTGTTTGCTGCTGTGCTGt | 4.1, 11.0, 5.7, 4.6, 21.4 | 40.8617291 | 0 |
| | | | | | 61951866 | -1 | CCCGaCAGCACAGCAGCAGG | CGG | 37.5 | 55.5 | 100 | 0.00 | 61951877 | CCGCCTGCTGCTGTGCTGtC | 1 | TTGTTTGTTGCTGTGCTGtC | 0.5, 11.0, 0.8, 20.3, 2.8 | 45.2560924 | 0 |
| | | | | | 61951869 | -1 | TGCCCCGaCAGCACAGCAGC | AGG | 34.3 | 55.4 | 100 | 0.00 | 61951878 | CGCCTGCTGCTGTGCTGtCG | 1 | TGTTTGTTGCTGTGCTGtCG | 0.5, 0.5, 9.3, 17.0 | 32.2670165 | 0 |
| | | | | | 61951876 | 1 | CCCGCCTGCTGCTGTGCTGt | CGG | 40.9 | 37.5 | 41.7 | 0.09 | 61951865 | CCGaCAGCACAGCAGCAGGC | -1 | GCCTGCTGCTGTACTAtCAA | 2.8, 21.9, 11.0, 0.5 | 39.828909 | 0 |
| | | | | | 61951877 | 1 | CCGCCTGCTGCTGTGCTGtC | GGG | 45.3 | 34.6 | 31.5 | 0.11 | 61951866 | CCCGaCAGCACAGCAGCAGG | -1 | CCTGCTGCTGTGCTAtCAAA | 13.5, 5.7, 11.0, 0.5 | 37.4842685 | 0 |
| | | | | | 61951878 | 1 | CGCCTGCTGCTGTGCTGtCG | GGG | 32.3 | 51.5 | 19.6 | 0.13 | 61951869 | TGCCCCGaCAGCACAGCAGC | -1 | GCTGCTGTGCTGtCAAAACA | 21.4, 20.3, 9.3, 0.5 | 34.3365711 | 0 |
| BEST1 Mutation | Location | WT | MT | sgRNA candidates | Position | Strand | Sequence | PAM | Specificity Score | Efficiency Score | Comp eff at WT | ASE score | Position | Guide | Guide Strand | Edited 5'->3' Sequence | Edit Scores | Efficiency Score | scarless repair |
| R25W | 61951879 | CTCCCGCCTGCTGCTGTGCTGGCGGGGCAGCATCTACAAGCTGCT | CTCCCGCCTGCTGCTGTGCTGGtGGGGCAGCATCTACAAGCTGCT | 6 | 61951865 | -1 | CCaCCAGCACAGCAGCAGGC | GGG | 37.8 | 57.3 | 98.6 | 0.00 | 61951876 | CCCGCCTGCTGCTGTGCTGG | 1 | TTTGTTTGCTGCTGTGCTGG | 4.1, 11.0, 5.7, 4.6, 21.4 | 38.6654473 | 0 |
| | | | | | 61951866 | -1 | CCCaCCAGCACAGCAGCAGG | CGG | 33.9 | 59.3 | 100 | 0.00 | 61951877 | CCGCCTGCTGCTGTGCTGGt | 1 | TTGTTTGTTGCTGTGCTGGt | 0.5, 11.0, 0.8, 20.3, 2.8 | 50.6872525 | 0 |
| | | | | | 61951869 | -1 | TGCCCCaCCAGCACAGCAGC | AGG | 46.9 | 55.5 | 68.3 | 0.08 | 61951878 | CGCCTGCTGCTGTGCTGGtG | 1 | TGTTTGTTGCTGTGCTGGtG | 0.5, 0.5, 9.3, 17.0 | 47.4547521 | 0 |
| | | | | | 61951876 | 1 | CCCGCCTGCTGCTGTGCTGG | tGG | 38.7 | 43.0 | 100 | 0.00 | 61951865 | CCaCCAGCACAGCAGCAGGC | -1 | GCCTGCTGCTGTACTAAtAA | 2.8, 20.3, 11.6, 11.0, 0.5 | 37.8354974 | 0 |
| | | | | | 61951877 | 1 | CCGCCTGCTGCTGTGCTGGt | GGG | 50.7 | 44.1 | 41.7 | 0.13 | 61951866 | CCCaCCAGCACAGCAGCAGG | -1 | CCTGCTGCTGTGCTAAtAAA | 21.4, 21.9, 5.7, 11.0, 0.5 | 33.8913824 | 0 |
| | | | | | 61951878 | 1 | CGCCTGCTGCTGTGCTGGtG | GGG | 47.5 | 52.4 | 31.5 | 0.17 | 61951869 | TGCCCCaCCAGCACAGCAGC | -1 | GCTGCTGTGCTGAtAAAACA | 6.8, 21.4, 20.3, 9.3, 0.5 | 46.8580089 | 0 |
| BEST1 Mutation | Location | WT | MT | sgRNA candidates | Position | Strand | Sequence | PAM | Specificity Score | Efficiency Score | Comp eff at WT | ASE score | Position | Guide | Guide Strand | Edited 5'->3' Sequence | Edit Scores | Efficiency Score | scarless repair |
| R25Gln | 61951880 | TCCCGCCTGCTGCTGTGCTGGCGGGGCAGCATCTACAAGCTGCTA | TCCCGCCTGCTGCTGTGCTGGCaGGGCAGCATCTACAAGCTGCTA | 5 | 61951865 | -1 | CtGCCAGCACAGCAGCAGGC | GGG | 17.8 | 55.6 | 100 | 0.00 | 61951877 | CCGCCTGCTGCTGTGCTGGC | 1 | TTGTTTGTTGCTGTGCTGGC | 0.5, 11.0, 0.8, 20.3, 2.8 | 45.3607453 | 0 |
| | | | | | 61951866 | -1 | CCtGCCAGCACAGCAGCAGG | CGG | 27.8 | 55.5 | 98.6 | 0.00 | 61951878 | CGCCTGCTGCTGTGCTGGCa | 1 | TGTTTGTTGCTGTGCTGGCa | 0.5, 0.5, 9.3, 17.0 | 48.1550154 | 0 |
| | | | | | 61951869 | -1 | TGCCCtGCCAGCACAGCAGC | AGG | 48.3 | 51.0 | 60.5 | 0.10 | 61951865 | CtGCCAGCACAGCAGCAGGC | -1 | GCCTGCTGCTGTACTAACaA | 2.8, 20.3, 0.8, 0.5 | 17.8296713 | 0 |
| | | | | | 61951877 | 1 | CCGCCTGCTGCTGTGCTGGC | aGG | 45.4 | 34.4 | 100 | 0.00 | 61951866 | CCtGCCAGCACAGCAGCAGG | -1 | CCTGCTGCTGTGCTAACaAA | 21.4, 4.6, 11.0, 0.5 | 27.8460332 | 0 |
| | | | | | 61951878 | 1 | CGCCTGCTGCTGTGCTGGCa | GGG | 48.2 | 49.9 | 41.7 | 0.14 | 61951869 | TGCCCtGCCAGCACAGCAGC | -1 | GCTGCTGTGCTGACaAAACA | 2.8, 20.3, 9.3, 0.5 | 48.2605659 | 0 |
| BEST1 Mutation | Location | WT | MT | sgRNA candidates | Position | Strand | Sequence | PAM | Specificity Score | Efficiency Score | Comp eff at WT | ASE score | Position | Guide | Guide Strand | Edited 5'->3' Sequence | Edit Scores | Efficiency Score | scarless repair |
| S27Arg | 61951887 | TGCTGCTGTGCTGGCGGGGCAGCATCTACAAGCTGCTATATGGCG | TGCTGCTGTGCTGGCGGGGCAGgATCTACAAGCTGCTATATGGCG | 2 | 61951882 | 1 | TGCTGCTGTGCTGGCGGGGC | AGg | 54.1 | 37.5 | 0 | 0.20 | 61951882 | TGCTGCTGTGCTGGCGGGGC | 1 | TGTTGTTGTGCTGGCGGGGC | 0.5, 13.4 | 54.1495982 | 0 |
| | | | | | 61951902 | 1 | AGgATCTACAAGCTGCTATA | TGG | 75.4 | 44.1 | 98.6 | 0.00 | 61951902 | AGgATCTACAAGCTGCTATA | 1 | AGgATTTACAAGCTGCTATA | 26.6 | 75.3829423 | 0 |
| BEST1 Mutation | Location | WT | MT | sgRNA candidates | Position | Strand | Sequence | PAM | Specificity Score | Efficiency Score | Comp eff at WT | ASE score | Position | Guide | Guide Strand | Edited 5'->3' Sequence | Edit Scores | Efficiency Score | scarless repair |
| Y29His | 61951891 | GCTGTGCTGGCGGGGCAGCATCTACAAGCTGCTATATGGCGAGTT | GCTGTGCTGGCGGGGCAGCATCcACAAGCTGCTATATGGCGAGTT | 2 | 61951896 | -1 | ACTCGCCATATAGCAGCTTG | TgG | 77.5 | 55.5 | 100 | 0.00 | 61951902 | AGCATCcACAAGCTGCTATA | 1 | AGTATTtACAAGCTGCTATA | 0.5, 26.6, 22.4 | 75.824827 | 1 |

| | | | | | | | | | | | | | | | | | | | |
|---|---|---|---|---|---|---|---|---|---|---|---|---|---|---|---|---|---|---|---|
| | | | | | 61951902 | 1 | AGCATCcACAAGCTGCTATA | TGG | 75.8 | 41.4 | 68.3 | 0.10 | 61951896 | ACTCGCCATATAGCAGCTTG | -1 | CAAGCTGCTATATAACAAAT | 22.4, 13.4, 21.7, 2.5 | 77.5443088 | 0 |
| BEST1 Mutation | Location | WT | MT | sgRNA candidates | Position | Strand | Sequence | PAM | Specificity Score | Efficiency Score | Comp eff at WT | ASE score | Position | Guide | Guide Strand | Edited 5'->3' Sequence | Edit Scores | Efficiency Score | scarless repair |
| K30Arg | 61951895 | TGCTGGCGGGGCAGCATCTACAAGCTGCTATATGGCGAGTTCTTA | TGCTGGCGGGGCAGCATCTACAgGCTGCTATATGGCGAGTTCTTA | 2 | 61951891 | 1 | GCTGGCGGGGCAGCATCTAC | AgG | 79.4 | 46.6 | 100 | 0.00 | 61951891 | GCTGGCGGGGCAGCATCTAC | 1 | GTTGGTGGGGCAGCATCTAC | 0.4, 13.4 | 79.4233066 | 0 |
| | | | | | 61951902 | 1 | AGCATCTACAgGCTGCTATA | TGG | 74.4 | 41.4 | 55.5 | 0.14 | 61951902 | AGCATCTACAgGCTGCTATA | 1 | AGTATTTACAgGCTGCTATA | 0.5, 26.6 | 74.3911217 | 0 |
| BEST1 Mutation | Location | WT | MT | sgRNA candidates | Position | Strand | Sequence | PAM | Specificity Score | Efficiency Score | Comp eff at WT | ASE score | Position | Guide | Guide Strand | Edited 5'->3' Sequence | Edit Scores | Efficiency Score | scarless repair |
| K30N | 61951896 | GCTGGCGGGGCAGCATCTACAAGCTGCTATATGGCGAGTTCTTAA | GCTGGCGGGGCAGCATCTACAAcCTGCTATATGGCGAGTTCTTAA | 2 | 61951902 | 1 | AGCATCTACAAcCTGCTATA | TGG | 75.2 | 40.7 | 49.2 | 0.16 | 61951902 | AGCATCTACAAcCTGCTATA | 1 | AGTATTTACAAcCTGCTATA | 0.5, 26.6 | 75.1580543 | 0 |
| | | | | | 61951902 | -1 | TTAAGAACTCGCCATATAGC | AGg | 76.9 | 55.5 | 0 | 0.43 | 61951902 | TTAAGAACTCGCCATATAGC | -1 | GCTATATGGCGAATTCTTAA | 6.8 | 76.9444828 | 0 |
| BEST1 Mutation | Location | WT | MT | sgRNA candidates | Position | Strand | Sequence | PAM | Specificity Score | Efficiency Score | Comp eff at WT | ASE score | Position | Guide | Guide Strand | Edited 5'->3' Sequence | Edit Scores | Efficiency Score | scarless repair |
| L35K | 61951909 | CATCTACAAGCTGCTATATGGCCAGTTCTTAATCTTCCTGCTCTG | CATCTACAAGCTGCTATATGGCaAGTTCTTAATCTTCCTGCTCTG | 0 | N/A | N/A | N/A | N/A | N/A | N/A | N/A | N/A | N/A | N/A | N/A | N/A | N/A | N/A | N/A |
| BEST1 Mutation | Location | WT | MT | sgRNA candidates | Position | Strand | Sequence | PAM | Specificity Score | Efficiency Score | Comp eff at WT | ASE score | Position | Guide | Guide Strand | Edited 5'->3' Sequence | Edit Scores | Efficiency Score | scarless repair |
| L41P | 61951928 | GGCGAGTTCTTAATCTTCCTGCTCTGCTACTACATCATCCGCTTT | GGCGAGTTCTTAATCTTCCTGCcCTGCTACTACATCATCCGCTTT | 3 | 61951929 | -1 | GGATGATGTAGTAGCAGgGC | AGG | 59.7 | 54.9 | 19.6 | 0.26 | 61951933 | AAGCGGATGATGTAGTAGCA | -1 | TGCTACTACATCATCCACTT | 0.8 | 78.6157351 | 0 |
| | | | | | 61951933 | -1 | AAGCGGATGATGTAGTAGCA | GgG | 78.6 | 66.0 | 100 | 0.00 | 61951934 | AAAGCGGATGATGTAGTAGC | -1 | GCTACTACATCATCCACTTT | 4.6 | 82.039877 | 0 |
| BEST1 Mutation | Location | WT | MT | sgRNA candidates | Position | Strand | Sequence | PAM | Specificity Score | Efficiency Score | Comp eff at WT | ASE score | Position | Guide | Guide Strand | Edited 5'->3' Sequence | Edit Scores | Efficiency Score | scarless repair |
| Q58Leu | 61955127 | aGGCTGGCCCTCACGGAAGAACAACAGCTGATGTTTGAGAAACTG | aGGCTGGCCCTCACGGAAGAACtACAGCTGATGTTTGAGAAACTG | 2 | Position | Strand | Sequence | PAM | Specificity Score | Efficiency Score | | | Position | Guide | Guide Strand | Edited 5'->3' Sequence | Edit Scores | Efficiency Score | |
| | | | | | 61955118 | -1 | CAGCTGTaGTTCTTCCGTGA | GGG | 85.6 | 67.1 | 100 | 0.00 | 61955118 | CAGCTGTaGTTCTTCCGTGA | -1 | TCACGGAAGAACtACAACTA | 0.8, 4.1 | 85.6357741 | 0 |
| | | | | | 61955119 | -1 | TCAGCTGTaGTTCTTCCGTG | AGG | 81.6 | 69.8 | 61.1 | 0.22 | 61955119 | TCAGCTGTaGTTCTTCCGTG | -1 | CACGGAAGAACtACAACTAA | 4.6, 6.4 | 81.6166894 | 0 |
| BEST1 Mutation | Location | WT | MT | sgRNA candidates | Position | Strand | Sequence | PAM | Specificity Score | Efficiency Score | Comp eff at WT | ASE score | Position | Guide | Guide Strand | Edited 5'->3' Sequence | Edit Scores | Efficiency Score | scarless repair |
| T91I | 61955742 | TTCTACGTGACGCTGGTCGTGACCCGCTGGTGGAACCAGTACGAG | TTCTACGTGACGCTGGTCGTGAtCCGCTGGTGGAACCAGTACGAG | 2 | 61955744 | 1 | TGACGCTGGTCGTGAtCCGC | TGG | 93.7 | 53.3 | 17.2 | 0.41 | 61955744 | TGACGCTGGTCGTGAtCCGC | 1 | TGATGTTGGTCGTGAtCCGC | 11.6, 13.4 | 93.665206 | 0 |
| | | | | | 61955747 | 1 | CGCTGGTCGTGAtCCGCTGG | TGG | 91.3 | 54.5 | 38.7 | 0.31 | 61955747 | CGCTGGTCGTGAtCCGCTGG | 1 | TGTTGGTTGTGAtCCGCTGG | 0.8, 0.5, 25.4 | 91.3186015 | 0 |
| BEST1 Mutation | Location | WT | MT | sgRNA candidates | Position | Strand | Sequence | PAM | Specificity Score | Efficiency Score | Comp eff at WT | ASE score | Position | Guide | Guide Strand | Edited 5'->3' Sequence | Edit Scores | Efficiency Score | scarless repair |
| R92G | 61955744 | CTACGTGACGCTGGTCGTGACCCGCTGGTGGAACCAGTACGAGAA | CTACGTGACGCTGGTCGTGACCgGCTGGTGGAACCAGTACGAGAA | 4 | 61955740 | 1 | TACGTGACGCTGGTCGTGAC | CgG | 95.3 | 56.5 | 0 | 0.54 | 61955740 | TACGTGACGCTGGTCGTGAC | 1 | TATGTGATGCTGGTCGTGAC | 5.6, 6.8 | 95.2968945 | 0 |
| | | | | | 61955744 | 1 | TGACGCTGGTCGTGACCgGC | TGG | 90.1 | 53.1 | 19.6 | 0.38 | 61955744 | TGACGCTGGTCGTGACCgGC | 1 | TGATGTTGGTCGTGACCgGC | 11.6, 13.4 | 90.0697165 | 0 |
| | | | | | 61955747 | 1 | CGCTGGTCGTGACCgGCTGG | TGG | 85.2 | 52.7 | 26.8 | 0.33 | 61955747 | CGCTGGTCGTGACCgGCTGG | 1 | TGTTGGTTGTGACCgGCTGG | 0.8, 0.5, 25.4 | 85.1688349 | 0 |
| | | | | | 61955748 | -1 | CTCGTACTGGTTCCACCAGC | cGG | 75.2 | 55.3 | 100 | 0.00 | 61955748 | CTCGTACTGGTTCCACCAGC | -1 | GCTGGTGGAACCAATACAAA | 19.0, 16.0, 4.1 | 75.1890487 | 0 |
| BEST1 Mutation | Location | WT | MT | sgRNA candidates | Position | Strand | Sequence | PAM | Specificity Score | Efficiency Score | Comp eff at WT | ASE score | Position | Guide | Guide Strand | Edited 5'->3' Sequence | Edit Scores | Efficiency Score | scarless repair |
| R92C | 61955744 | CTACGTGACGCTGGTCGTGACCCGCTGGTGGAACCAGTACGAGAA | CTACGTGACGCTGGTCGTGACCtGCTGGTGGAACCAGTACGAGAA | 3 | 61955744 | 1 | TGACGCTGGTCGTGACCtGC | TGG | 81.2 | 54.2 | 19.6 | 0.35 | 61955744 | TGACGCTGGTCGTGACCtGC | 1 | TGATGTTGGTCGTGACCtGC | 11.6, 13.4 | 81.1858422 | 0 |
| | | | | | 61955747 | 1 | CGCTGGTCGTGACCtGCTGG | TGG | 71.1 | 53.7 | 26.8 | 0.28 | 61955747 | CGCTGGTCGTGACCtGCTGG | 1 | TGTTGGTTGTGACCtGCTGG | 0.8, 0.5, 25.4 | 71.0970537 | 0 |
| | | | | | 61955748 | -1 | CTCGTACTGGTTCCACCAGC | aGG | 75.2 | 53.5 | 100 | 0.00 | 61955748 | CTCGTACTGGTTCCACCAGC | -1 | GCTGGTGGAACCAATACAAA | 19.0, 16.0, 4.1 | 75.1890487 | 0 |
| BEST1 Mutation | Location | WT | MT | sgRNA candidates | Position | Strand | Sequence | PAM | Specificity Score | Efficiency Score | Comp eff at WT | ASE score | Position | Guide | Guide Strand | Edited 5'->3' Sequence | Edit Scores | Efficiency Score | scarless repair |
| R92S | 61955744 | CTACGTGACGCTGGTCGTGACCCGCTGGTGGAACCAGTACGAGAA | CTACGTGACGCTGGTCGTGACCaGCTGGTGGAACCAGTACGAGAA | 3 | 61955744 | 1 | TGACGCTGGTCGTGACCaGC | TGG | 79.5 | 54.7 | 19.6 | 0.35 | 61955744 | TGACGCTGGTCGTGACCaGC | 1 | TGATGTTGGTCGTGACCaGC | 11.6, 13.4 | 79.4690404 | 0 |
| | | | | | 61955747 | 1 | CGCTGGTCGTGACCaGCTGG | TGG | 75.5 | 56.2 | 26.8 | 0.31 | 61955747 | CGCTGGTCGTGACCaGCTGG | 1 | TGTTGGTTGTGACCaGCTGG | 0.8, 0.5, 25.4 | 75.495335 | 0 |
| | | | | | 61955748 | -1 | CTCGTACTGGTTCCACCAGC | tGG | 75.2 | 47.5 | 100 | 0.00 | 61955748 | CTCGTACTGGTTCCACCAGC | -1 | GCTGGTGGAACCAATACAAA | 19.0, 16.0, 4.1 | 75.1890487 | 0 |
| BEST1 Mutation | Location | WT | MT | sgRNA candidates | Position | Strand | Sequence | PAM | Specificity Score | Efficiency Score | Comp eff at WT | ASE score | Position | Guide | Guide Strand | Edited 5'->3' Sequence | Edit Scores | Efficiency Score | scarless repair |
| R92H | 61955745 | TACGTGACGCTGGTCGTGACCCGCTGGTGGAACCAGTACGAGAAC | TACGTGACGCTGGTCGTGACCCaCTGGTGGAACCAGTACGAGAAC | 4 | 61955744 | 1 | TGACGCTGGTCGTGACCCaC | TGG | 87.8 | 55.0 | 31.5 | 0.33 | 61955744 | TGACGCTGGTCGTGACCCaC | 1 | TGATGTTGGTCGTGACCCaC | 11.6, 13.4 | 87.8034991 | 0 |
| | | | | | 61955747 | 1 | CGCTGGTCGTGACCCaCTGG | TGG | 78.3 | 57.9 | 17.2 | 0.38 | 61955747 | CGCTGGTCGTGACCCaCTGG | 1 | TGTTGGTTGTGACCCaCTGG | 0.8, 0.5, 25.4 | 78.2885309 | 0 |
| | | | | | 61955748 | -1 | CTCGTACTGGTTCCACCAGt | GGG | 82.7 | 58.4 | 41.7 | 0.28 | 61955748 | CTCGTACTGGTTCCACCAGt | -1 | aCTGGTGGAACCAATACAAA | 19.0, 16.0, 4.1 | 82.693454 | 0 |
| | | | | | 61955749 | -1 | TCTCGTACTGGTTCCACCAG | tGG | 76.3 | 63.5 | 100 | 0.00 | 61955749 | TCTCGTACTGGTTCCACCAG | -1 | CTGGTGGAACCAATACAAAA | 6.8, 21.7, 6.4 | 76.2936189 | 0 |
